# Functional carbohydrate-active enzymes acquired by horizontal gene transfer from plants in the whitefly *Bemisia tabaci*

**DOI:** 10.1101/2024.06.03.597214

**Authors:** D Colinet, M Haon, E Drula, M Boyer, S Grisel, C Belliardo, GD Koutsovoulos, JG Berrin, EGJ Danchin

**Affiliations:** Institut Sophia Agrobiotech, INRAE, Université Côte d’Azur, CNRS, Sophia Antipolis, France; INRAE, Aix Marseille Univ., BBF, Biodiversité et Biotechnologie Fongiques, Marseille, France; INRAE, Aix Marseille Univ., 3PE Platform, Marseille, France; Architecture et Fonction des Macromolécules Biologiques (AFMB), CNRS, Aix Marseille Université, Marseille, France

**Keywords:** horizontal gene transfer, genome evolution, carbohydrate active enzymes, functional glucanases, plant feeding insects

## Abstract

Carbohydrate-active enzymes (CAZymes) involved in the degradation of plant cell walls and/or the assimilation of plant carbohydrates for energy uptake are widely distributed in microorganisms. In contrast, they are less frequent in animals, although there are exceptions, including examples of CAZymes acquired by horizontal gene transfer (HGT) from bacteria or fungi in several of phytophagous arthropods and plant-parasitic nematodes. Although the whitefly *Bemisia tabaci* is a major agricultural pest, knowledge of HGT-acquired CAZymes in this phloem-feeding insect of the Hemiptera order (subfamily Aleyrodinae) is still lacking. We performed a comprehensive and accurate detection of HGT candidates in *B. tabaci* and identified 136 HGT events, 14 of which corresponding to CAZymes. The *B. tabaci* HGT-acquired CAZymes were not only of bacterial or fungal origin, but some were also acquired from plants. Biochemical analysis revealed that members of the glycoside hydrolase families 17 (GH17) and 152 (GH152) acquired from plants are functional beta-glucanases with different substrate specificities, suggesting distinct roles. These two CAZymes are the first characterized GH17 and GH152 glucanases in an animal. We identified a lower number of HGT events in the related Aleyrodinae *Trialeurodes vaporariorum*, with only three HGT-acquired CAZymes, including a GH152 glucanase, with phylogenetic analysis suggesting a unique HGT event in the ancestor of the Aleyrodinae. Another GH152 CAZyme, most likely independently acquired from plants, was also identified in two plant cell-feeding insects of the Thysanoptera order, highlighting the importance of plant-acquired CAZymes in the biology of piercing-sucking insects.

**Significance statement:** Carbohydrate-active enzymes (CAZymes) are crucial for sugar metabolism. Those involved in plant cell wall degradation are usually absent from animal genomes. In this study, we explored CAZyme repertoires in the genomes of several insects: the phloem-feeding whitefly *Bemisia tabaci*, a major agricultural pest, and the related greenhouse whitefly *Trialeurodes vaporariorum*, as well as two Thysanoptera species that feed on plant cell contents. We identified several cases of CAZymes acquired from plant via horizontal gene transfer in the genome of these insects. Notably, we showed that two *B. tabaci* CAZymes of plant origin function as glucanases with distinct substrate specificities, potentially helping the insect to overcome plant defenses. Overall, these findings enhance our understanding of how the ability to feed on plants evolved in insects.

## Introduction

Horizontal gene transfer (HGT) can be defined as the transfer of genetic material between organisms without vertical transmission from parent to offspring. HGT allows for the transfer of genes between different species, regardless of their evolutionary distance. This phenomenon has been extensively studied in bacteria, for which the supporting mechanisms have been comprehensively documented (Arnold et al. 2022). Although less common than in bacteria, it is becoming increasingly evident that HGT has had a significant impact on the evolution of multicellular eukaryotic genomes, driving functional novelty. In recent years, several HGT events have been described between bacteria or viruses and eukaryotes, and even between different eukaryotic organisms (Keeling and Palmer 2008; Crisp et al. 2015; Soucy et al. 2015; Drezen et al. 2017; Husnik and McCutcheon 2018; Chen et al. 2021). Among eukaryotes, HGT has played an important role in the evolution of the ability of many animals to feed on plants, including phytophagous arthropods (Kirsch et al. 2014; Nakabachi 2015; Wybouw et al. 2016; Husnik and McCutcheon 2018) and plant-parasitic nematodes (Haegeman et al. 2011; Danchin and Rosso 2012; Husnik and McCutcheon 2018; Lai et al. 2023). In particular, a number of genes encoding carbohydrate-active enzymes (CAZymes) involved in plant cell wall degradation have been described as acquired via HGT in arthropods and nematodes.

The plant cell wall is a dynamic structure that serves multiple functions, including protection from biotic stresses. It is composed of a network of high molecular weight polysaccharides, including β-1,4 glucans (cellulose), β-1,4 heteroxylans, β-1,4 glucomannans, xyloglucan, or mixed β-1,3/β-1,4 glucans (hemicelluloses), the latter being mainly restricted to Poales, and the heteropolysaccharide pectin (Perrot et al. 2022). The first plant cell wall degrading enzymes (PCWDEs) identified in an animal were ß-1,4-endoglucanases (or cellulases), which belong to the glycoside hydrolase 5 (GH5) family of CAZymes (Drula et al. 2022) and were presumably horizontally acquired from bacteria by plant-parasitic nematodes (Smant et al. 1998). Since then, numerous examples of CAZymes acquired by HGT from bacterial or fungal donors, and presumed or shown to be active on plant cell wall polysaccharides, have been described in plant-parasitic nematodes (Danchin et al. 2010; Haegeman et al. 2011; Danchin and Rosso 2012; Palomares-Rius et al. 2014; Vicente et al. 2019), but also in phytophagous arthropods (Pauchet et al. 2010; Pauchet and Heckel 2013; Kirsch et al. 2014; Pauchet et al. 2014; Evangelista et al. 2015; Antony et al. 2017; Faddeeva-Vakhrusheva et al. 2017; Busch et al. 2019; Shin et al. 2021; Le et al. 2022; Shin and Pauchet 2023). HGT-acquired CAZymes identified in plant-parasitic nematodes and phytophagous arthropods also include enzymes directly involved in plant carbohydrate assimilation for energy metabolism (Acuña et al. 2012; Sun et al. 2013; Danchin et al. 2016; Dai et al. 2021).

The whitefly *Bemisia tabaci* (Hemiptera: Aleyrodinae) is a major agricultural pest. It feeds on phloem sap, causing damage directly through feeding with its piercing-sucking mouthparts and indirectly through the transmission of numerous plant pathogenic viruses (Navas-Castillo et al. 2011). *B. tabaci* is a highly polyphagous species complex of more than 30 cryptic species, among which the genome of the Middle East-Asia Minor 1 (MEAM1), Mediterranean (MED), and SSA-East and Central Africa (SSA-ECA) pests have been recently released (Chen et al. 2016; Xie et al. 2017; Chen et al. 2019). HGT appears to be widespread in *B. tabaci,* with numerous genes reported to have been transferred not only from bacteria or fungi (Chen et al. 2016; Li et al. 2022), but also from plants (Gilbert and Maumus 2022; Li et al. 2022), making *B. tabaci* the first animal species documented to have acquired genes of plant origin (Lapadula et al. 2020). Among these horizontally acquired genes, functional evidence supports the involvement of genes of bacterial origin in B vitamin synthesis (Ren et al. 2020) and nitrogen metabolism (Yang et al. 2024), genes of fungal origin in the ferredoxin-mediated suppression of plant defenses (Wang et al. 2023), and genes of plant origin in processes such as detoxification of plant toxic compounds (Xia et al. 2021) or reproduction (Gong et al. 2023). We performed a careful mining of previous analyses (Chen et al. 2016; Gilbert and Maumus 2022; Li et al. 2022), suggesting the possible acquisition of several CAZymes of different origin in *B. tabaci*. However, none of these enzymes have been biochemically characterized and their possible role in the biology of the insect remains unknown. Furthermore, comprehensive and accurate detection of HGT-acquired CAZymes in *B. tabaci* is still lacking, leaving a knowledge gap. Phloem feeders are thought to rely on CAZymes to facilitate stylet penetration through plant cell walls into a phloem sieve element, but also to counteract the sieve element occlusion defense mechanism consisting of callose (β-1,3-glucan) deposition (Silva-Sanzana et al. 2020; Walker 2022). CAZymes are also needed to transform sugars taken up from the plant, mainly sucrose, into fructose for energy metabolism. Finally, some phloem feeders depend on CAZymes to overcome the high osmolarity of the phloem through osmoregulation (Douglas 2006).

In this work, we used comprehensive phylogenetic analyses to identify HGT events in *B. tabaci* MEAM1 that are specific to the Aleyrodinae, and we performed further characterization with a focus on CAZymes. Biochemical analysis revealed that two of the plant-acquired CAZymes, belonging to the GH17 and GH152 families, are functional beta-glucanases with different substrate specificities, suggesting distinct functional roles. We then performed a comparative analysis with the related greenhouse whitefly *Trialeurodes vaporariorum* (Hemiptera: Aleyrodinae), which differs from *B. tabaci* in various aspects such as host plant range or virus transmission (Byrne and Bellows 1991; Fiallo-Olivé et al. 2020. Finally, we expanded our analysis beyond the Aleyrodinae to detect HGT in two other piercing-sucking insects, *Frankliniella occidentalis* and *Thrips palmi* (Thysanoptera: Thripinae), which feed on the contents of plant cells. Overall, our findings suggest that the acquisition of CAZymes by HGT has had a significant impact on the biology of piercing-sucking insects, particularly in the case of *B. tabaci*.

## Results and Discussion

### The genome of B. tabaci contains numerous genes of non-animal origin

Our first objective was to perform a comprehensive and accurate phylogenetic detection of HGT candidates in *B. tabaci,* a member of the Aleyrodinae subfamily in the Hemiptera order, and to compare the results obtained with the literature. The proteome completeness estimated by BUSCO was compared between the three *B. tabaci* cryptic species, MEAM1, MED and SSA-ECA, for which genome and predicted proteome data are available (Supplementary Table 1). With 95.6% of arthropoda BUSCO proteins found to be complete, *B. tabaci* MEAM1 had the most complete proteome, compared to less than 90% for the other two *B. tabaci* cryptic species (Supplementary Table 1). Therefore, the search for candidate HGTs was performed for *B. tabaci* MEAM1 as a reference.

Of the 15,662 *B. tabaci* MEAM1 predicted proteins, 686 returned an AHS greater than 0, meaning that they are more similar to non-animal than to animal proteins. From these predicted proteins, AvP identified 357 as possible HGT candidates specific to Aleyrodinae. The vast majority of these (255 out of 357) were validated according to the criteria described in the Material and Methods section and were from potential bacterial, fungal, or plant (Viridiplantae) donors (Table 1 and Supplementary Table 2). These 255 validated HGT candidates correspond to at least 136 distinct HGT events (Supplementary Table 2). For each of these events, homologs were found in at least one of the other two *B. tabaci* cryptic species, MED and SSA-ECA, making the possibility of contamination unlikely (Supplementary Table 2). Further supporting the lack of contamination, the local score calculated by AvP was greater than 0 for each of the 255 validated HGT candidates. This result (136 events for 255 genes) also suggests that most of the candidate HGTs were inserted into multiple independent genomic regions and did not undergo massive cis-duplication (Supplementary Table 2).

**Table 1.** Overall comparison of HGT candidate search results.

|  | *Bemisia tabaci*<br>MEAM1 | *Trialeurodes*<br>*vaporariorum* | *Frankliniella*<br>*occidentalis* | *Thrips palmi* |
| --- | --- | --- | --- | --- |
| Number of protein-coding genes | 15,662 | 18,275 | 15,678 | 14,332 |
| **Number of predicted proteins with AHS > 0** | **686** | **910** | **281** | **233** |
| **Number of HGT candidates detected by AvP** | **357** | **182** | **50** | **57** |
| - Bacteria | 147 | 87 | 15 | 6 |
| - Fungi | 93 | 10 | 10 | 29 |
| - Viridiplantae | 78 | 15 | 7 | 9 |
| - Other | 39 | 70 | 18 | 13 |
| **Number of validated HGT candidates** | **255** | **75** | **23** | **20** |
| - Bacteria | 113 | 61 | 10 | 6 |
| - Fungi | 88 | 8 | 3 | 2 |
| - Viridiplantae | 53 | 5 | 1 | 3 |
| - Complex: Bacteria or Fungi | 1 | 1 | 10 | 9 |
| **Number of validated HGT events** | **136** | **32** | **8** | **8** |
| - Bacteria | 70 | 24 | 5 | 4 |
| - Fungi | 38 | 4 | 1 | 1 |
| - Viridiplantae | 27 | 3 | 1 | 2 |
| - Complex: Bacteria or Fungi | 1 | 1 | 1 | 1 |

We compared the HGT candidates of bacterial, fungal and plant origin we found in *B. tabaci* MEAM1 with data from the literature. Most of the HGT candidates described in the literature for *B. tabaci* MEAM1 were confirmed by our study suggesting that our approach is comprehensive despite the stringent filters applied (Figure 1). Of the 93 HGT candidates of bacterial origin described in the literature (Chen et al. 2016; Li et al. 2022), 11 were not found in our work (Figure 1 and Supplementary Table 3). Of these, one was no longer present in version 1.2 of the *B. tabaci* MEAM1 proteome, one was rejected because its sequence identity with the donor sequences was below 30%, and one was considered non-HGT as animal sequences were present in the sister and ancestral sister branches (Supplementary Table 3). The remaining eight were found to have an AHS score equal to or less than 0, indicating a higher similarity to animals than to non-animals in the NR database and thus unlikely acquisition via HGT (Supplementary Table 3). On the other hand, we identified 31 new phylogenetically supported HGT candidates with bacteria as potential donors not yet described in the literature (Figure 1 and Supplementary Table 2). 17 of these new HGT candidates belong to the AAA-ATPase-like domain-containing protein family, a large family of ATPases associated with various cellular activities. All but one of them group with previously identified HGT candidates in *B. tabaci* MEAM1, suggesting that they originate from the same HGT events but were overlooked in previous analyses. The remaining one would correspond to a new, independent HGT event of a member of the AAA-ATPase-like domain-containing protein family in *B. tabaci* (Supplementary Table 2). Horizontal gene transfer from bacteria has been described for members of this protein family in two other phloem-feeding insects of the order Hemiptera, but belonging to the superfamilies Psylloidea and Coccoidea, separated from Aleyrodinae by more than 250 million years (Myr) (Misof et al. 2014), with a possible role in mediating the interaction with endosymbionts (Sloan et al. 2014).

**Figure 1.**
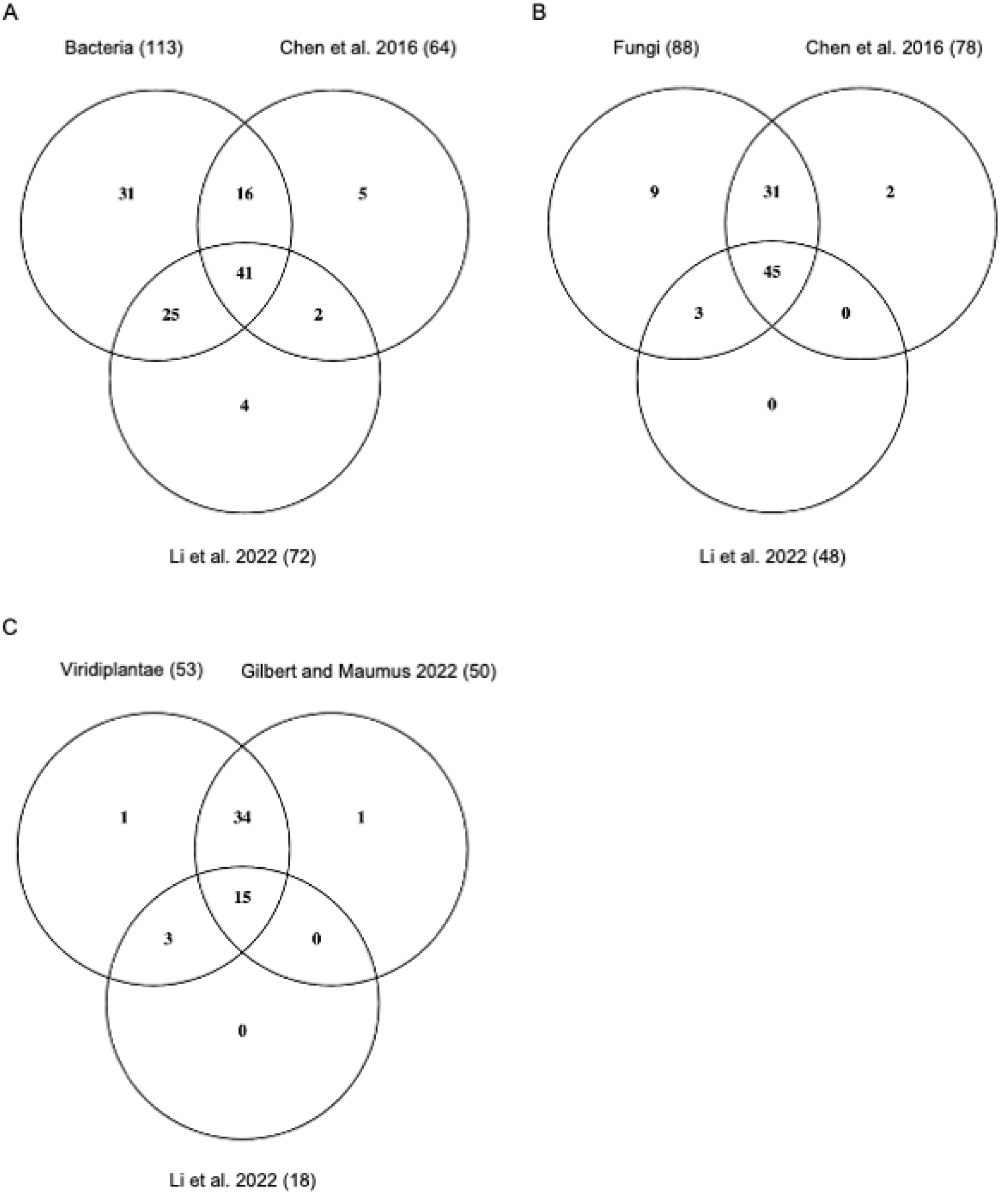
Comparison of the number of HGT candidates found for *B. tabaci* MEAM1 with bacteria (A), fungi (B) or plant (C) as donors with data from the literature.

Of the 81 HGT candidates of fungal origin described in the literature (Chen et al. 2016), only two were not retrieved in our work (Figure 1). For one of these two HGT candidates, no similarities were found in the NR database, preventing the search for a possible HGT origin (Supplementary Table 3). For the second candidate, classified as HGT_complex in our study, the HGT donor could not be unambiguously determined because the sister branch consists of bacterial sequences and the ancestral sister branch consists of fungal sequences (Supplementary Table 3). On the other hand, we identified nine new phylogenetically confirmed HGT candidates with fungi as potential donors (Figure 1 and Supplementary Table 2). Of these nine new HGT candidates, five cluster with HGT candidates already described in the literature, suggesting that they originated from the same HGT events as the latter (Supplementary Table 2). The other four would correspond to previously undescribed HGT acquisitions of two different genes encoding replication factor A protein 1 and a protein of unknown function, respectively (Supplementary Table 2).

Finally, of the 53 HGT candidates of plant origin described in the literature (Lapadula et al. 2020; Xia et al. 2021; Gilbert and Maumus 2022; Li et al. 2022), only one was not found in our work (Figure 1). No significant similarities were found in the NR database for this HGT candidate, which prevented the search for a possible HGT origin (Supplementary Table 3). In the work of Gilbert and Maumus (2022), this HGT candidate was identified as a member of a large family of omega-6 fatty acid desaturases, all derived from a single HGT event. The other 14 members of this family were successfully found in our work (Supplementary Table 2). On the other hand, we identified one new HGT candidate acquired from plants (Figure 1 and Supplementary Table 2). This new HGT candidate belongs to the HXXXD family of acyl-transferases. Other members of this family have already been described as HGT candidates for *B. tabaci* MEAM1 (Gilbert and Maumus 2022; Li et al. 2022). This new case would correspond to an independent HGT event in the same family (Supplementary Table 2).

Overall, our approach to identify HGT candidates using AvP appeared to be not only reliable, but also more exhaustive than previous approaches used for bacterial and fungal donors (Chen et al. 2016; Li et al. 2022) (Figure 1). For HGT candidates with plants as donors, our approach is comparable to that of Gilbert and Maumus (2022), and both of them are more comprehensive than that of Li et al. (2022) (Figure 1).

### B. tabaci has acquired CAZymes from bacteria, fungi, and plant donors via multiple HGT events

We identified significantly overrepresented GO terms among validated HGT candidates from potential bacterial, fungal, or plant donors in *B. tabaci* based on their InterPro domain annotation (Supplementary Table 4). This revealed a significant enrichment of GO terms related to carbohydrate metabolism and plant cell wall degradation, such as hydrolase activity, acting on glycosyl bonds (GO:0016798), pectinesterase activity (GO:0030599), or cell wall modification (GO:0042545). This suggests horizontal acquisition of a substantial proportion of Carbohydrate-active enzymes (CAZymes) in *B. tabaci.* We therefore investigated the relationships between HGT candidates and CAZymes, as well as possible donors, in more detail. A total of 433 *B. tabaci* MEAM1 proteins were predicted to be CAZymes (Supplementary Table 5). These include two members of the Glycoside Hydrolase (GH) family GH13, subfamily 17 (GH13_17), which have been shown to catalyze the detoxification of glucosinolates produced by plants against herbivores (Malka et al. 2020), but were not detected as having been acquired by HGT.

We identified 25 CAZymes likely acquired by HGT via 14 independent events (Table 2 and Supplementary Figure 1). Three CAZymes were predicted to originate from bacteria (corresponding to two HGT events), twelve from fungi (corresponding to four HGT events), and nine from plants (corresponding to seven HGT events). For the last CAZyme HGT candidate, it was not possible to decipher whether the origin was bacterial or fungal (Table 2 and Supplementary Figure 1).

**Table 2.** Horizontally acquired CAZymes in *B. tabaci* and *T. vaporariorum*.

| CAZy |  | Origin of donor sequences | *B. tabaci* |  | *T. vaporariorum* |  |
| --- | --- | --- | --- | --- | --- | --- |
| Description | Definition |  | HGT event | Sequence name | HGT event | Sequence name |
| GH32 | Glycoside Hydrolase Family 32 protein | Bacteria | BtaB53 | Bta00773<br>Bta04549 | - | - |
| CBM50 | Carbohydrate-Binding Module Family 50 | Bacteria | BtaB62 | Bta15714 | - | - |
| GH13 | Glycoside Hydrolase Family 13 protein | Bacteria | - | - | TvaB10 | Tv_04079-RA |
| GH49 | Glycoside Hydrolase Family 49 protein | Fungi | BtaF06 | Bta02250 | - | - |
| GH71 | Glycoside Hydrolase Family 71 protein | Fungi | BtaF15 | Bta02866 | - | - |
| CBM32-AA5_2 | Carbohydrate-Binding Module Family 32 / Galactose oxidase | Fungi | BtaF18 | Bta01072<br>Bta03567<br>Bta05090<br>Bta05091<br>Bta05092<br>Bta05093<br>Bta07691<br>Bta09066<br>Bta12453 | - | - |
| GH30_3 | Glycoside Hydrolase Family 30 | Fungi | BtaF20 | Bta02658 | - | - |
| GH152 | Glycoside Hydrolase Family 152 protein | Viridiplantae | BtaV01 | Bta13961 | TvaV01 | Tv_15928-RA |
| GT10 | Glycosyltransferase Family 10 protein | Viridiplantae | BtaV10 | Bta08851 | - | - |
| GH17 | Glycoside Hydrolase Family 17 protein | Viridiplantae | BtaV12 | Bta06115<br>Bta06118 | - | - |
| EXPN | Distantly related to plant expansins | Viridiplantae | BtaV13 | Bta15239 | - | - |
| CE8 | Carbohydrate Esterase Family 8 protein | Viridiplantae | BtaV24 | Bta11221<br>Bta11222 | - | - |
| GT61 | Glycosyltransferase Family 61 protein | Viridiplantae | BtaV25 | Bta14885 | - | - |
| GT17 | Glycosyltransferase Family 17 protein | Viridiplantae | BtaV27 | Bta15659 | - | - |
| PL1_4 | Polysaccharide Lyase<br>Family 1 protein | Complex:<br>Bacteria or<br>Fungi | BtaC01 | Bta02634 | TvaC01 | Tv_05105-RA |

Two of the CAZymes of bacterial origin belong to the GH32 family and would correspond to a single unique HGT event according to the AvP results (Table 2 and Supplementary Figure 1). Enzymatic activities in the GH32 family include invertase (or ß-fructofuranosidase) activity, which catalyzes the hydrolysis of sucrose into fructose and glucose. Independent horizontal acquisition events of ß-fructofuranosidase genes from bacterial donors have been evidenced for several animal species with a role in assimilation of plant carbohydrates (Nakabachi 2015; Danchin et al. 2016; Wybouw et al. 2016; Dai et al. 2021). Acquisition of these GH32 CAZymes from bacteria would have allowed *B. tabaci* to metabolize host-derived sucrose, which is abundant in the phloem on which it feeds. The third protein of bacterial origin classified as a CAZyme contains a carbohydrate-binding module of the CBM50 family, but no associated catalytic module (Table 2 and Supplementary Figure 1).

Among the CAZymes of fungal origin, three belong to the GH30, GH49 and GH71 families, respectively (Table 2 and Supplementary Figure 1). Interestingly, GH30 enzymes, presumably acquired via HGT, have been found in numerous plant-parasitic nematodes (Danchin et al. 2010), with xylanase activity confirmed in root-knot nematodes (Mitreva-Dautova et al. 2006). However, although xylanase enzymes have been found in other insects, none of those characterized belong to the GH30 family (Padilla-Hurtado et al. 2012; Pauchet and Heckel 2013; Pauchet et al. 2014; Vega et al. 2015). This suggests a convergent acquisition of xylanases from different sources. The GH49 enzyme may be active on plant arabinan (http://www.cazy.org/; Drula et al. 2022), although arabinanase activity has never been confirmed so far in any eukaryote in this GH family. Finally, the only known activity to date in the GH71 family is α-1,3-glucanase, which has been characterized for this family only in fungi. A multigenic family of nine proteins containing a carbohydrate binding module (CBM32) in tandem with a galactose oxidase module (AA5_2) was also found to be horizontally acquired from fungi (Table 2 and Supplementary Figure 1). The biological role of fungal AA5_2 galactose oxidases is not known but other members of this subfamily displaying alcohol oxidase activity play a role in the mechanism of plant penetration in fungal phytopathogens (Bissaro et al. 2022). To the best of our knowledge, there have been no reports of horizontal acquisition of genes containing this combination of modules (AA5_2-CBM32) in animals.

The HGT candidate whose bacterial or fungal origin could not be determined from the phylogeny belongs to the PL1_4 family (Table 2 and Supplementary Figure 1). PL1_4 enzymes are pectin lyases involved in the degradation of pectin and are usually found in bacteria or fungi (http://www.cazy.org/; Drula et al. 2022). Sequences from a few other insects, including the Thripinae *F. occidentalis* and *T. palmi*, are found in the phylogeny, in different branches for some (Supplementary Figure 1). This will be further investigated in the last part of the Results and Discussion section.

As far as we know, no CAZyme acquired by HGT from a plant donor has yet been fully described and characterized in an animal. Here, we identified a total of nine *B. tabaci* MEAM1 CAZymes, representing seven HGT events of plant origin (Table 2 and Supplementary Figure 1). Of these, three belong to the GlycosylTransferase (GT) families GT10, GT17 and GT61. Of the remaining six, three belong to the GH17 and GH152 families, two belong to the Carbohydrate Esterase 8 (CE8) family, and one is distantly related to plant expansins. The possible functions of some of these CAZymes are described in more detail below.

### A GH17 CAZyme acquired by HGT from plants in B. tabaci is a functional β-1,3-glucanase

The *B. tabaci* MEAM1 proteins Bta06115 and Bta06118 share similarities with plant ß-glucanases belonging to the GH17 family (Table 2 and Supplementary Table 2). The GH17 family is well described, with over 50 enzymes biochemically characterized (http://www.cazy.org/; Drula et al. 2022). The GH17 family includes two major groups of ß-glucanases, endo-β-1,3-glucanases, which hydrolyze internal β-1,3 glycosidic linkages in β-1,3-glucans, and endo-β-1,3-1,4-glucanases, which hydrolyze internal β-1,4 glycosidic linkages in mixed β-1,3-1,4-glucans (http://www.cazy.org/; Drula et al. 2022). Members of the GH17 family are found in bacteria, fungi and plants, but to our knowledge, they have not been documented in animals so far. Accordingly, homologs of the *B. tabaci* MEAM1 Bta06115 and Bta06118 proteins were identified only in *B. tabaci* MED and SSA-ECA, and no other organism besides plants (Figure 2A and Supplementary Table 2). The close proximity of both Bta06115 and Bta06118 genes on the same scaffold, together with the bootstrap-supported grouping of all *B. tabaci* GH17 sequences (Figure 2A), suggests that they originated from a single HGT event from a plant donor followed by a tandem duplication. The bootstrap-supported grouping of *B. tabaci* MED and SSA-ECA proteins with each of *B. tabaci* MEAM1 Bta06115 and Bta06118 proteins, respectively (Figure 2A), further suggests that the duplication occurred before the divergence of the three cryptic species, which is estimated to have occurred between approximately 5 Myr ago (Chen et al. 2019) and 40 Myr ago (Mugerwa et al. 2018). *B. tabaci* MEAM1 Bta06115 and Bta06118 were surrounded by *bona fide* insect genes not acquired by HGT (Supplementary Table 2), and synteny was quite conserved in the genomic region of the Bta06115 and Bta06118 genes in *B. tabaci* MED (Supplementary Figure 2A), ruling out the possibility of contamination.

**Figure 2.**
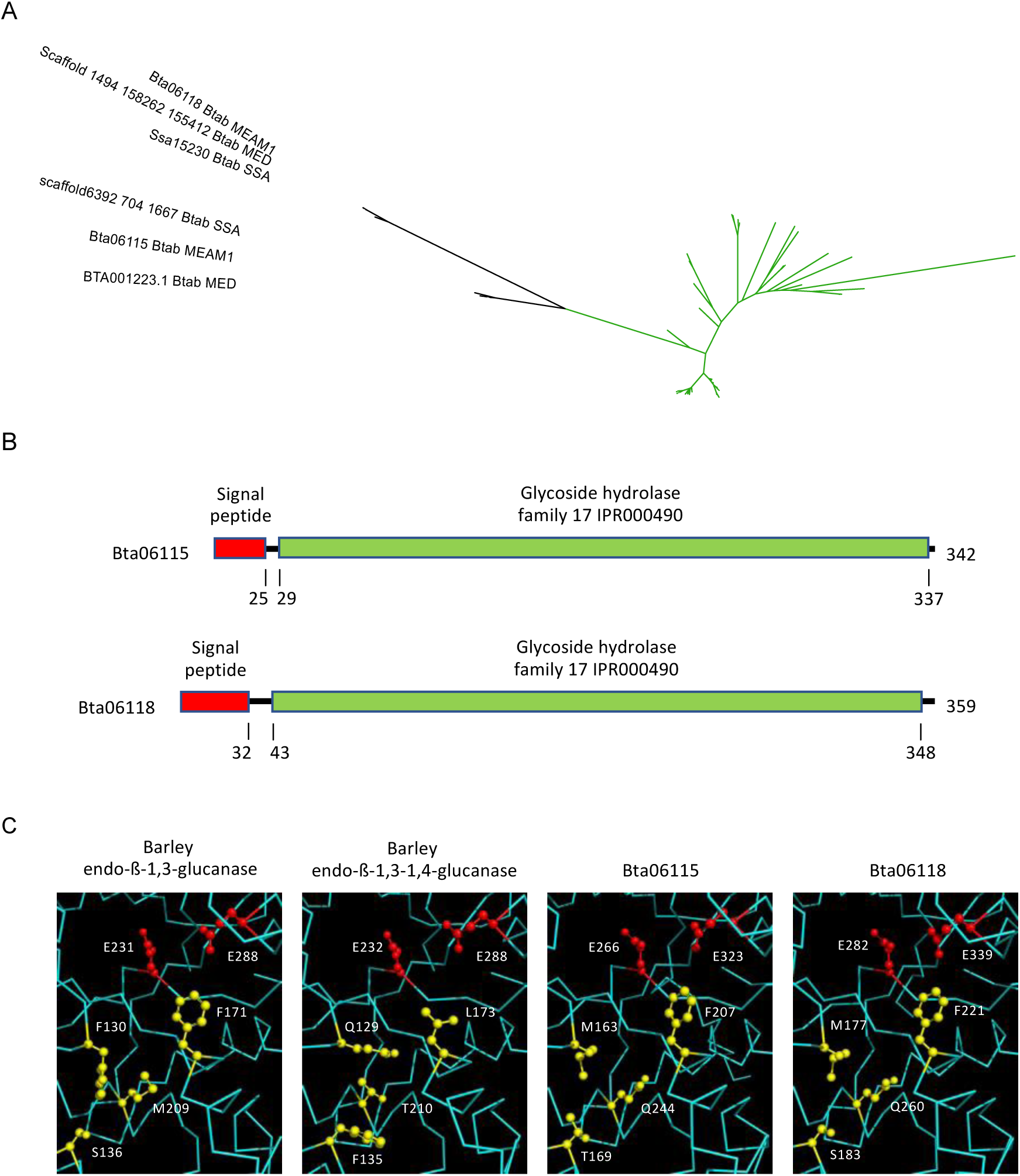
Analysis of the GH17 family proteins acquired by HGT in *B. tabaci*. (A) Phylogenetic tree with *B. tabaci* MEAM1, MED and SSA sequences shown in black. Green branches represent plant sequences. Gray circles indicate nodes with support values greater than or equal to 80% and 95% for SH-aLRT and UFboot, respectively. (B) Domain organization of the *B. tabaci* MEAM1 Bta06115 and Bta06118 proteins. (C) Comparison of the two catalytic glutamate residues (red) and specific residues that determine substrate specificity (yellow) in the catalytic cleft between the crystal structures of barley endo-ß-1,3-glucanase and endo-ß-1,3-1,4-glucanase and the predicted structures of *B. tabaci* MEAM1 Bta06115 and Bta06118 proteins.

Both *B. tabaci* MEAM1 Bta06115 and Bta06118 proteins consist of a GH17 domain, preceded by a signal peptide, suggesting that they are secreted (Figure 2B). The predicted structures of the Bta06115 and Bta06118 proteins (without the peptide signal) were obtained using AlphaFold2 and aligned with the crystal structure of barley endo-ß-1,3-glucanase and endo-ß-1,3-1,4-glucanase (Varghese et al. 1994). The RMSD values for the alpha carbon atoms ranged from 0.629 Å (for all 255 atoms) to 0.767 Å (for all 278 atoms) between the Bta06118 and barley endo-ß-1,3-1,4-glucanase structures and the Bta06115 and barley endo-ß-1,3-glucanase structures, respectively, indicating a high degree of structural similarity (Supplementary Figure 3). Accordingly, the predicted structures of Bta06115 and Bta06118, along with the crystal structures of barley endo-ß-1,3-glucanase and barley endo-ß-1,3-1,4-glucanase, exhibit the typical (α/ß)_8_ TIM barrel motif shared by GH17 glucanases (Supplementary Figure 3). This motif consists of an inner crown of eight ß-strands linked by loops to an outer crown of eight α-helices (Varghese et al. 1994; Receveur-Bréchot et al. 2006). The two glutamic acid catalytic residues that constitute the active site of GH17 ß-glucanases (Varghese et al. 1994) are conserved in the predicted structures of Bta06115 and Bta06118, suggesting the functionality of the *B. tabaci* MEAM1 GH17 proteins (Figure 2C).

Bta06115 was heterologously expressed in *P. pastoris* and purified to homogeneity. To investigate substrate preferences of this putative glucanase, we first assayed Bta06115 on cellohexaose (G6; β-1,4-gluco-oligosaccharides) and laminarihexaose (L6; β-1,3-gluco-oligosaccharides) as substrates (Figure 3A). Enzyme assays revealed that Bta06115 was able to efficiently cleave the β-1,3-linkages found in L6, producing L2 and glucose (G1), suggesting exo-type activity. Although very weak activity was detected on the β-1,4-linkages found in G6, no activity was detected on amorphous cellulose. We then tested the enzyme on two different mixed-linked β-glucans (i.e. lichenan and barley glucan), displaying a 1,3/1,4 mean ratio of 1:2 and 1:3, respectively, as found in cereals (Figure 3B). Significant activity was detected on lichenan, which has a low ratio of β-1,4-linkages, while no significant cleavage was observed on barley β-glucan, which contains a high ratio of β-1,4-linkages. Overall, we conclude that the GH17 Bta06115 is a functional β-1,3-glucanase active on both β-1,3-glucans and mixed β-1,3-1,4-glucans, but only with a low ratio of β-1,4-linkages.

**Figure 3.**
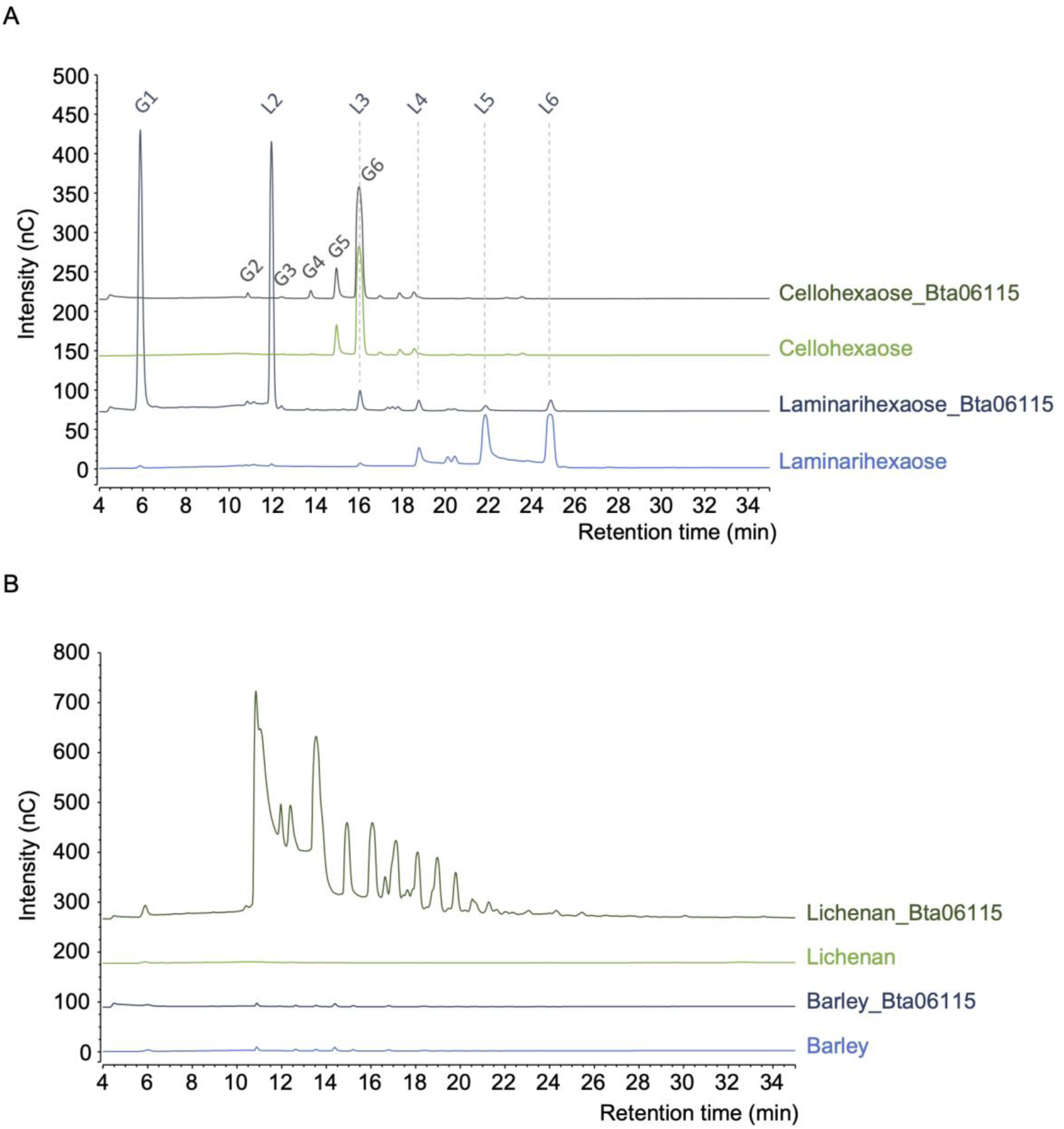
Functional analysis of the GH17 family protein Bta06115 acquired by HGT in *B. tabaci*. The graphs show HPAEC-PAD chromatograms of reaction products released by the GH17 Bta06115 from cellohexaose (G6) and laminarihexaose (L6) (A) and from barley glucan and lichenan (B). G1 (L1): glucose; G2: cellobiose; G3: cellotriose; G4: cellotetraose; G5: cellopentaose; G6: cellohexaose; L2: laminaribiose; L3: laminaritriose; L4: laminaritetraose; L5: laminaripentaose; L6: laminarihexaose.

Accordingly, comparison of the specific residues that determine substrate specificity confirms that Bta06115 and Bta06118 are ß-1,3-glucanases (Figure 2C). In barley endo-ß-1,3-1,4-glucanase, residues Q129 and F135 block the catalytic cleft and prevent the ß-1,3-glucans from binding, while the short T210 leaves sufficient sufficient space for mixed β-1,3-1,4-glucans (Varghese et al. 1994). Residues F130 and S136, which replace Q129 and F135 in barley endo-ß-1,3-glucanase, provide space to accommodate ß-1,3-glucans (Varghese et al. 1994), as would M163 and T169 in Bta06115 and M177 and S183 in Bta06118 (Figure 2C). In addition, the replacement of T210 by M209 in barley endo-ß-1,3-glucanase represents a barrier to a mixed β-1,3-1,4-glucan with a high ratio of β-1,4-linkages (Varghese et al. 1994). Residues Q244 and Q260 would form a similar barrier in Bta06115 and Bta06118 respectively (Figure 2C).

### A functional GH152 ß-1,3-1,4-glucanase was acquired by HGT from a plant donor in B. tabaci

The *B. tabaci* MEAM1 Bta13961 protein shares similarities with plant ß-glucanases belonging to the GH152 family (Table 2 and Supplementary Table 2). The GH152 family is much less described than the GH17 family (http://www.cazy.org/; Drula et al. 2022), with only one biochemically characterized enzyme from a filamentous fungi (Sakamoto et al. 2006). In our Diamond homology search, homologs of the *B. tabaci* MEAM1 Bta13961 protein were identified in *B. tabaci* MED and SSA-ECA, but not in any organism other than plants (Supplementary Table 2 and Figure 4A). The monophyletic grouping of *B. tabaci* sequences was robust, suggesting horizontal acquisition from a plant donor prior to the divergence of the three *B. tabaci* cryptic species (Figure 4A). The *B. tabaci* MEAM1 Bta13961 was surrounded by non-HGT genes (Supplementary Table 2), and synteny was quite conserved in the genomic region of the Bta13961 gene in *B. tabaci* SSA (Figure 4B and Supplementary Figure 2B), ruling out the possibility of contamination. The Bta13961 protein shares similarities to thaumatin-like (TL) proteins found in plants (Supplementary Table 2). It consists of a signal peptide followed by a thaumatin family domain that covers most, if not all, of the mature peptide (Figure 4C), typical of TL proteins (De Jesús-Pires et al. 2020). Commonly found in plants, TL proteins include pathogenesis-related proteins involved in plant defense against pathogens (De Jesús-Pires et al. 2020). One mechanism of action of some of these TL proteins in plant defense would be their demonstrated endo-ß-1,3-glucanase activity, which would degrade the cell walls of pathogenic fungi (Grenier et al. 1999). TL proteins have also been identified in animals, although no evidence of HGT has yet been demonstrated, and in fungi, including a protein from *Lentinula edodes* with demonstrated ß-1,3-glucanase activity (Sakamoto et al. 2006) and classified as the only characterized member of the GH152 family (http://www.cazy.org/; Drula et al. 2022).

**Figure 4.**
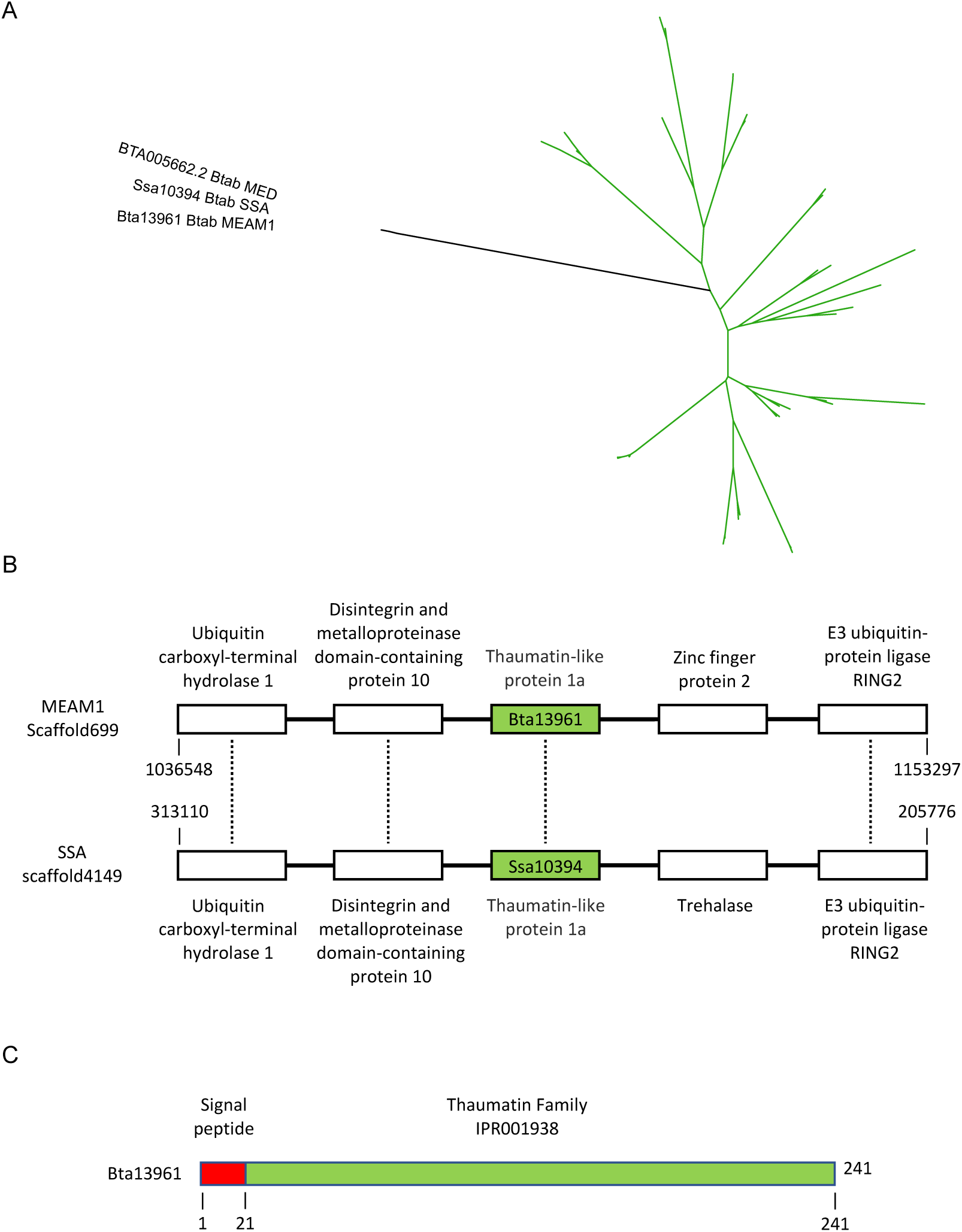
Analysis of the GH152 family protein Bta13961 acquired by HGT in *B. tabaci*. (A) Phylogenetic tree with *B. tabaci* MEAM1, MED and SSA sequences shown in black. Green branches represent plant sequences. Gray circles indicate nodes with support values greater than or equal to 80% and 95% for SH-aLRT and UFboot, respectively. (B) Comparative map of the genomic region of the gene encoding the GH152 family protein in *B. tabaci* MEAM1 (Bta13961) and SSA (Ssa10394) Each rectangle represents a *B. tabaci* gene. (C) Domain organization of the *B. tabaci* MEAM1 Bta13961 protein.

The predicted structure of the Bta13961 protein (without the peptide signal) was obtained using AlphaFold2 and aligned with the crystal structure of the TL protein NP24-I from tomato (Ghosh and Chakrabarti 2008). The RMSD values for the alpha carbon atoms was 0.693 Å (for all 157 atoms), indicating a high degree of structural similarity (Figure 5A). The REDDD motif that form the acidic cleft responsible for the β-1,3-glucanase activity of TL proteins and that contain the two presumed catalytic residues (De Jesús-Pires et al. 2020) were conserved in the Bta13961 structure, suggesting the functionality of the *B. tabaci* MEAM1 Bta13961 protein (Figure 5A).

**Figure 5.**
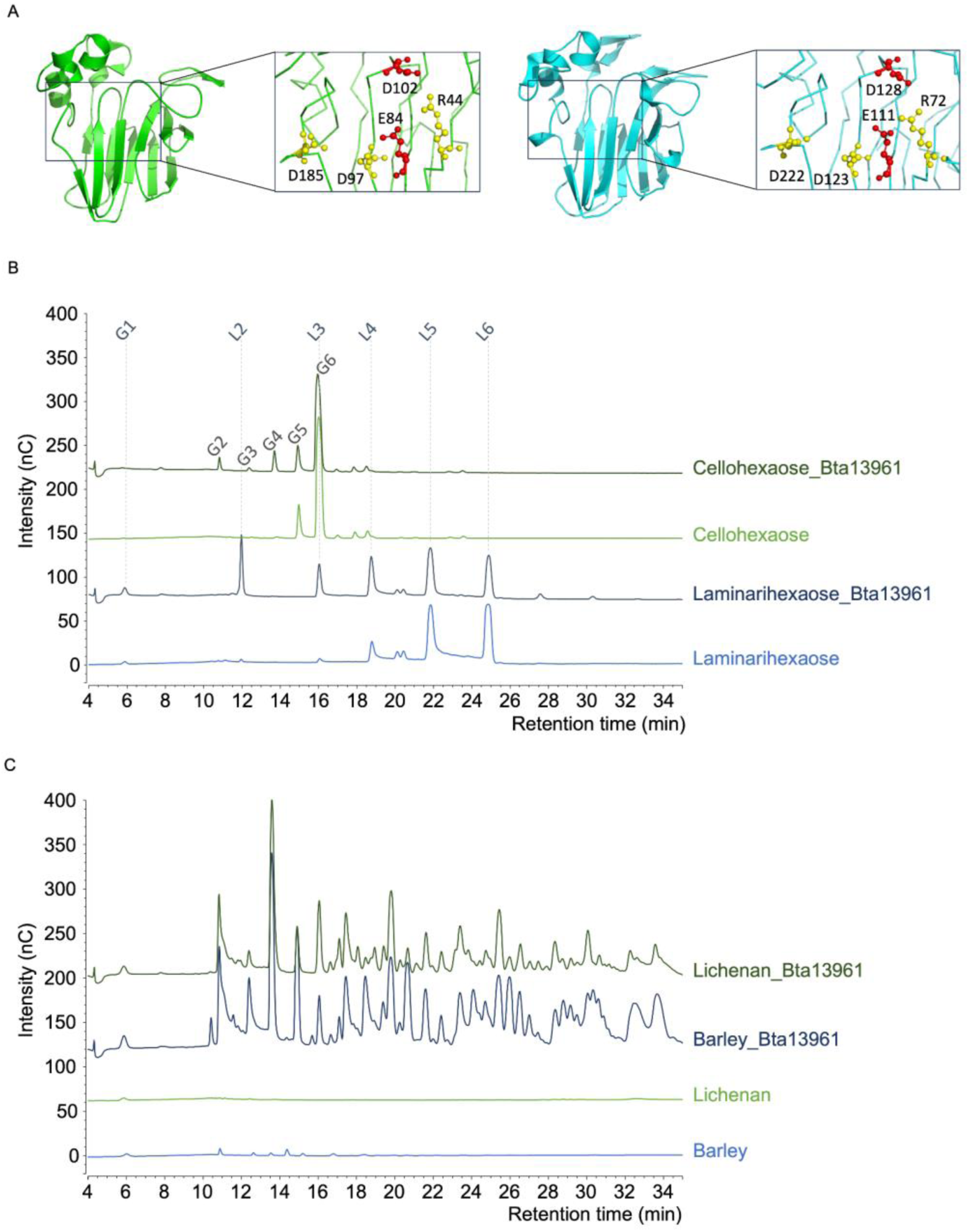
Structural and functional analysis of the GH152 family protein Bta13961 acquired by HGT in *B. tabaci*. (A) Comparison of the crystal structure of tomato TL protein NP24-I (green) with the predicted structure of *B. tabaci* MEAM1 Bta13961 (cyan). Amino acids of the REDDD motif are colored red for the two putative catalytic residues and yellow for the other three. (B and C) HPAEC-PAD chromatograms of reaction products released by the GH152 Bta13961 from cellohexaose (G6) and laminarihexaose (L6) (B) and from barley glucan and lichenan (C). G1 (L1): glucose; G2: cellobiose; G3: cellotriose; G4: cellotetraose; G5: cellopentaose; G6: cellohexaose; L2: laminaribiose; L3: laminaritriose; L4: laminaritetraose; L5: laminaripentaose; L6: laminarihexaose.

To investigate the substrate specificity of Bta13961, we used the same approach as for the GH17 Bta06115. After successful expression in *P. pastoris* and purification, we first demonstrated that the enzyme was active on both β-1,3 and β-1,4 oligosaccharides using G6 or L6 as substrates (Figure 5B). The activity was significant, but in both cases, the substrates were not fully consumed, meaning that they are not the preferred substrates. Furthermore, no activity was detected on amorphous cellulose. Enzyme activity was much more significant on β-1,3/β-1,4 glucans, Bta13961 being able to cleave both mixed-linked β-glucans (barley glucan and lichenan) with similar efficiency (Figure 5C). These results clearly indicate a different behavior of the GH152 Bta13961 compared to the GH17 Bta06115. Indeed, the 1,3/1,4 mean ratio in β-glucans did not impact enzyme activity and the products released (most probably β-1,3/β-1,4 oligosaccharides) were much longer as compared to the products released by Bta06115. Overall, these results indicated that the GH152 Bta13961 is a functional endo-β-1,3-1,4-glucanase able to cleave both β-1,3 and β-1,4 linkages. Interestingly, only β-1,3-glucanase activity has been described for the sole GH152 biochemically characterized to date (http://www.cazy.org/; Drula et al. 2022).

### A CE8 candidate pectin methylesterases was horizontally transferred from a plant donor in B. tabaci

The *B. tabaci* MEAM1 Bta11221 and Bta11222 genes encode proteins with similarities to pectin methylesterases (PMEs) found in plants (Supplementary Table 2). PMEs are enzymes of the CE8 family of CAZymes that catalyze the demethoxylation of pectin, a major component of the plant cell wall (Pelloux et al. 2007; Jolie et al. 2010). PMEs are ubiquitous enzymes in plants, but are also found in bacterial and fungal phytopathogens with a role in breaking down the plant cell wall to allow infection. PMEs have also been identified in some arthropods (Pauchet et al. 2010; Evangelista et al. 2015; Antony et al. 2017; Faddeeva-Vakhrusheva et al. 2017) and in a plant parasitic nematode (Vicente et al. 2019), and were likely horizontally acquired from bacteria.

Homologs of the *B. tabaci* MEAM1 Bta11221 and Bta1122 proteins were identified in *B. tabaci* MED and SSA-ECA, but not in any organism other than plants (Supplementary Table 2 and Figure 6A). The robust monophyletic grouping of all *B. tabaci* sequences suggests horizontal acquisition from a plant donor prior to the divergence of the three *B. tabaci* cryptic species (Supplementary Table 2 and Figure 6A). The *B. tabaci* MEAM1 Bta11221 and Bta1122 genes were surrounded by genes conserved in other arthropods and animals, which were therefore not horizontally acquired (Supplementary Table 2). Furthermore, synteny was conserved in the genomic region of the Bta11221 and Bta1122 genes in *B. tabaci* SSA (Figure 6B and Supplementary Figure 2C), ruling out the possibility of contamination.

**Figure 6.**
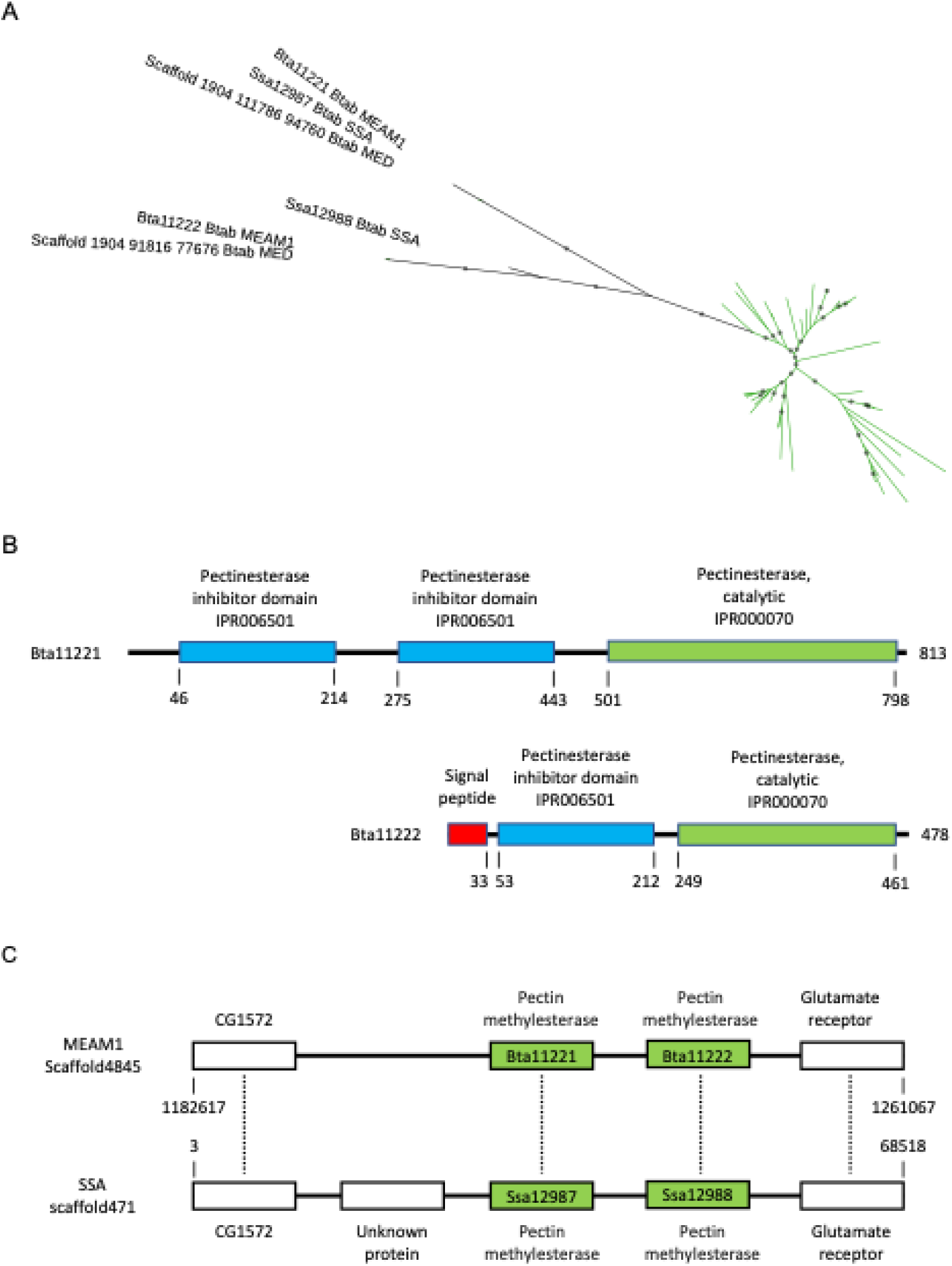
Analysis of the pectin methylesterases acquired by HGT in *B. tabaci*. (A) Phylogenetic tree with *B. tabaci* MEAM1, MED and SSA sequences shown in black. Green branches represent plant sequences. Gray circles indicate nodes with support values greater than or equal to 80% and 95% for SH-aLRT and UFboot, respectively. (B) Domain organization of the *B. tabaci* MEAM1 Bta11221 and Bta11222 proteins. (C) Comparative map of the genomic region of the gene encoding the pectin methylesterases in *B. tabaci* MEAM1 (Bta11221 and Bta11222) and SSA (Ssa12987 and Ssa12988). Each rectangle represents a *B. tabaci* gene.

The two *B. tabaci* PME genes are consecutive in *B. tabaci* MEAM1 and SSA-ECA genomes, suggesting that the HGT event was followed by a tandem duplication before the divergence of the two cryptic species (Figure 6B). Another possibility is that the two genes were organized in tandem in the plant donor and were transferred in the same HGT event. However, this is not consistent with the bootstrap-supported grouping of *B. tabaci* PME proteins into a single monophyletic group (Figure 6A). Both *B. tabaci* MEAM1 Bta11221 and Bta1122 proteins consist of a pectinesterase catalytic domain, preceded by one or two pectinesterase inhibitory domains and, in the case of Bta11222, a signal peptide (Figure 6C). This domain organization, in which the active part of the protein is later cleaved from the N-terminal inhibitory region, is typical for group 2 PMEs found exclusively in plants (Pelloux et al. 2007; Jolie et al. 2010). This is consistent with the hypothesis of an HGT event from a plant donor.

The predicted structures of the PME active part of the Bta11221 and Bta1122 proteins were obtained using AlphaFold2 and aligned with the crystal structure of carrot PME, the first to be solved in plants (Johansson et al. 2002). The resulting RMSD values calculated for the alpha carbon atoms were 0.613 Å (for 285 out of 297 atoms) and 0.529 Å (for 171 out of 210 atoms), respectively, indicating a high level of structural similarity (Supplementary Figure 4). The five key residues that were identified in the active site of carrot PME (Johansson et al. 2002; Jolie et al. 2010) are conserved in the predicted structure of Bta11221 and Bta11222 (Supplementary Figure 4). These observations suggest that the Bta11221 and Bta11222 proteins are functional PMEs.

### An expansin-related protein was acquired horizontally from a plant donor in B. tabaci

Expansins are enigmatic proteins that promote loosening and extension of the plant cell wall, although they lack enzymatic activity (Cosgrove 2015). Canonical expansins are characterized by the presence of two domains, an N-terminal six-stranded double-psi beta-barrel (DPBB), which is structurally related to GH45, and a C-terminal CBM63. The *B. tabaci* MEAM1 Bta15239 gene encodes a protein with similarities to GH45 endoglucanases (EG45-like domain-containing proteins) from plants (Supplementary Table 2). These are expansin-related proteins that contain the N-terminal expansin-like DPBB domain, but not the C-terminal CBM63 domain found in canonical expansins (Cosgrove 2015). Consistently, the *B. tabaci* MEAM1 Bta15239 protein consists of a single DPBB domain preceded by a signal peptide (Figure 7A). Homologs of the *B. tabaci* MEAM1 Bta15239 protein were identified in the *B. tabaci* MED and SSA-ECA cryptic species, but not in any organism other than plants (Supplementary Table 2 and Figure 7B). The *B. tabaci* MEAM1 Bta15239 gene was surrounded by non-HGT genes (Supplementary Table 2), and synteny was conserved in the genomic region of the Bta15239 gene in *B. tabaci* SSA (Figure 7C and Supplementary Figure 2D), ruling out the possibility of contamination.

**Figure 7.**
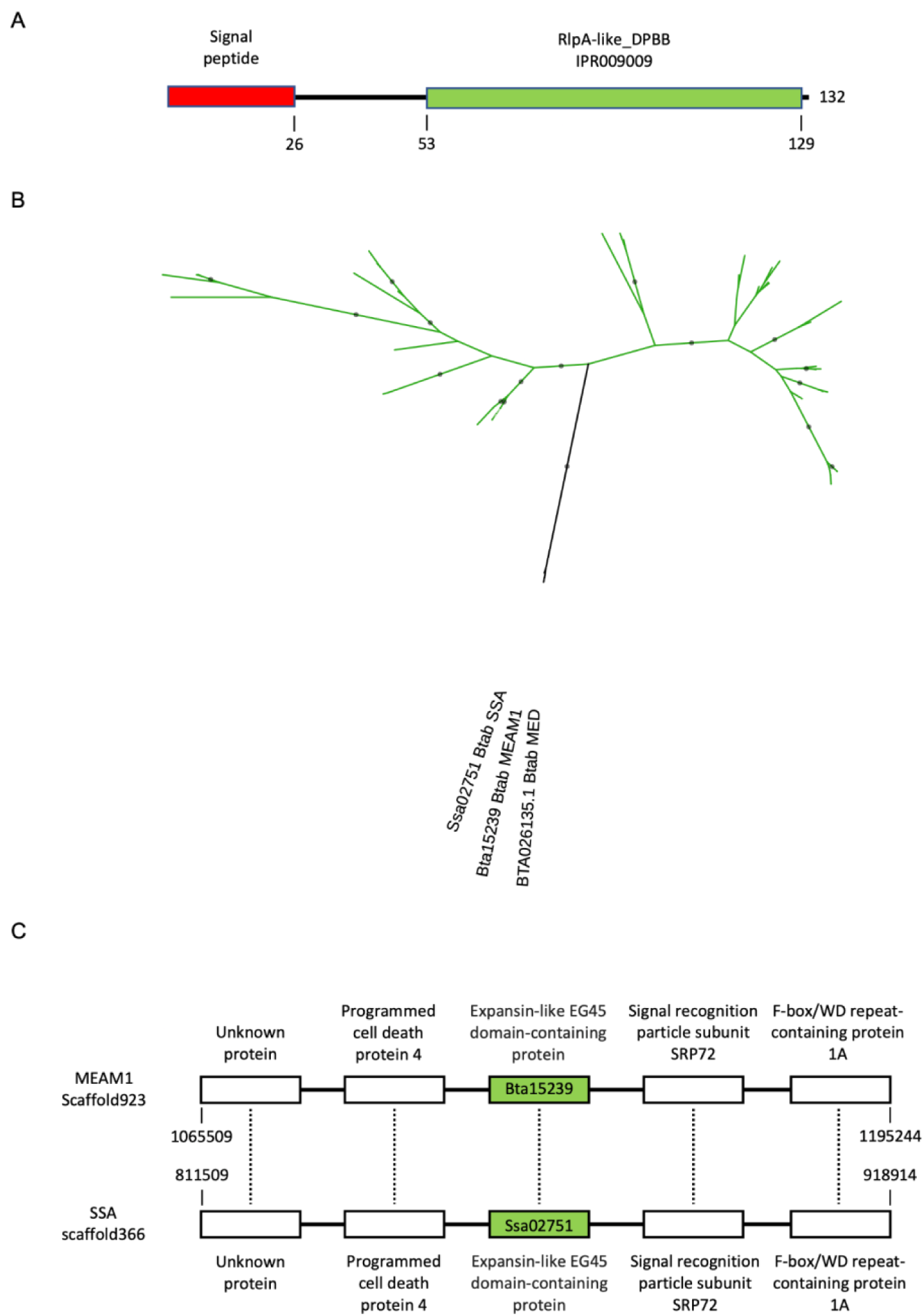
Analysis of the expansin-related EG45 domain-containing protein acquired by HGT in *B. tabaci*. (A) Domain organization of the *B. tabaci* MEAM1 Bta15239 protein. (B) Phylogenetic tree with *B. tabaci* MEAM1, MED and SSA sequences shown in black. Green branches represent plant sequences. Gray circles indicate nodes with support values greater than or equal to 80% and 95% for SH-aLRT and UFboot, respectively. (C) Comparative map of the genomic region of the gene encoding the expansin-related EG45 domain-containing protein in *B. tabaci* MEAM1 (Bta15239) and SSA (Ssa02751). Each rectangle represents a *B. tabaci* gene.

Expansin-like and expansin-related proteins are present in a large number of plant pathogens among bacteria and fungi with reported roles in plant colonization and virulence (Georgelis et al. 2015; Cosgrove 2017). Evolutionary analyses suggest that these microbial proteins originated from multiple horizontal gene transfers from plants as well as between microbes (Georgelis et al. 2015; Cosgrove 2017). In animals, expansin-like and expansin-related proteins, probably of bacterial origin, have been described as effectors of parasitism in several plant parasitic nematodes (Danchin et al. 2010). Expansins are thought to act by loosening the interactions between plant cell wall components to facilitate the action of various enzymes also secreted by the nematodes (Qin et al. 2004). However, there is no evidence of cell wall loosening activity for single domain DPBB proteins, which are likely to have a function distinct from that of canonical expansins (Cosgrove 2015).

### Much less horizontally acquired CAZymes were found in T. vaporariorum compared to B. tabaci

Next, we investigated whether HGT events have made similar contributions to the biology and genome evolution of *T. vaporariorum,* another species in the Aleyrodinae subfamily in the Hemiptera order, which is separated from *B. tabaci* by approximately 80 to 90 Myr (Misof et al. 2014). In their study, Gilbert and Maumus investigated whether some homologs of the *B. tabaci* MEAM1 horizontally acquired genes from plants were conserved in *T. vaporariorum* (Gilbert and Maumus 2022). However, they did not perform a comprehensive search for horizontally acquired genes in the latter. We used AvP to identify HGT candidates in *T. vaporariorum* and compared the results with those obtained in *B. tabaci*. Of the 18,725 predicted proteins in *T. vaporariorum*, 182 proteins were identified as likely HGT candidates using AvP (Table 1 and Supplementary Table 6). Of these, 75 candidates specific to Aleyrodinae have been validated with potential donors from bacteria, fungi, or plants, representing less than one-third of the number of validated HGT candidates for *B. tabaci*. The main difference between the two species is in the number of HGT candidates from fungi (8 for *T. vaporariorum* compared to 88 for *B. tabaci*) or plants (5 for *T. vaporariorum* compared to 53 for *B. tabaci*) potential donors (Table 1 and Supplementary Table 6). Viruses have been proposed to act as vectors for HGT in eukaryotes (Gilbert and Cordeaux, 2017). Therefore, a possible explanation for the difference in the number of HGT candidates from plants between the two Aleyrodinae species could be their different capacities for viral transmission. *B. tabaci* has been reported to transmit a large number of viruses belonging to at least six different genera, of which the genus *Begomovirus* is by far the most important, whereas the number of viruses transmitted by *T. vaporariorum* is more limited and mainly restricted to criniviruses and torradoviruses (Fiallo-Olivé et al. 2020).

The 75 HGT candidates that were validated in *T. vaporariorum* correspond to at least 32 distinct HGT events (Supplementary Table 6). Of these events, 23 showed clustering between sequences of *B. tabaci* and *T. vaporariorum*, as determined by Orthofinder. For each of these 23 events, we combined the groups defined by AvP for both *B. tabaci* and *T. vaporariorum*. Phylogenies were then inferred according to the procedure described in the Material and Methods section. The sequences of *B. tabaci* and *T. vaporariorum* formed robust monophyletic clades in 11 of the combined groups, suggesting HGT events that occurred before the two species diverged (Supplementary Figure 5). One of these groups includes Bta02634, classified as a member of the PL1_4 pectin lyase CAZyme family, and Tv_05105-RA, identified as its ortholog by Orthofinder (Table 2). The origin of these genes would correspond to an HGT event in the ancestor of Aleyrodinae, although it could not not be clearly determined from the phylogeny whether the donor belongs to bacteria or fungi (Supplementary Figure 1).

Sequences of *B. tabaci* and *T. vaporariorum* formed a monophyletic clade in three of the remaining 12 combined groups, albeit with lower bootstrap support values (Supplementary Table 7). In the remaining nine combined groups, there was no monophyletic clade grouping both *B. tabaci* and *T. vaporariorum* sequences (Supplementary Figure 6). Alternative topology tests for monophyly of Aleyrodinae sequences proved inconclusive for seven of them (Supplementary Table 7). Therefore, it is difficult to conclude with confidence that there is a single ancestral HGT event for each of these 11 combined groups. One of these groups involves Bta13961, classified as a member of the GH152 CAZyme family acquired from plants and demonstrated to be functional as a glucanase (see above), and Tv_15928-RA, identified as its ortholog through Orthofinder. The two sequences shared 42.2% identity and 53.5% similarity at the protein level, but no signal peptide was found for Tv_15928-RA, in contrast to the *B. tabaci* GH152 sequence. Their predicted structures were aligned with an RMSD of 0.586 Å, indicating structural similarity (Figure 8A and Supplementary Figure 1). Despite their similarity, the GH152 sequences of *B. tabaci* and *T. vaporariorum* were found in two different clades in the phylogeny obtained for the combined group (Figure 8B). Furthermore, the synteny of neighboring genes of Bta13961 in *B. tabaci* and Tv_15928-RA in *T. vaporariorum* was not conserved between the two species, indicating that the HGT occurred at different genomic locations (Supplementary Figure 2E). These findings suggest that the GH152 sequences in *B. tabaci* and *T. vaporariorum* originated from two distinct HGT events, although an alternative topology test, constrained to support the monophyly of Aleyrodinae sequences, was not significantly less likely (Supplementary Table 7). Interestingly, the group formed by Tv_15928-RA and two plant sequences included a sequence from *T. palmi*, which belongs to the Thysanoptera order, an outgroup of Hemiptera (Figure 8B). This candidate case of HGT in thrips will be further investigated in the next section.

**Figure 8.**
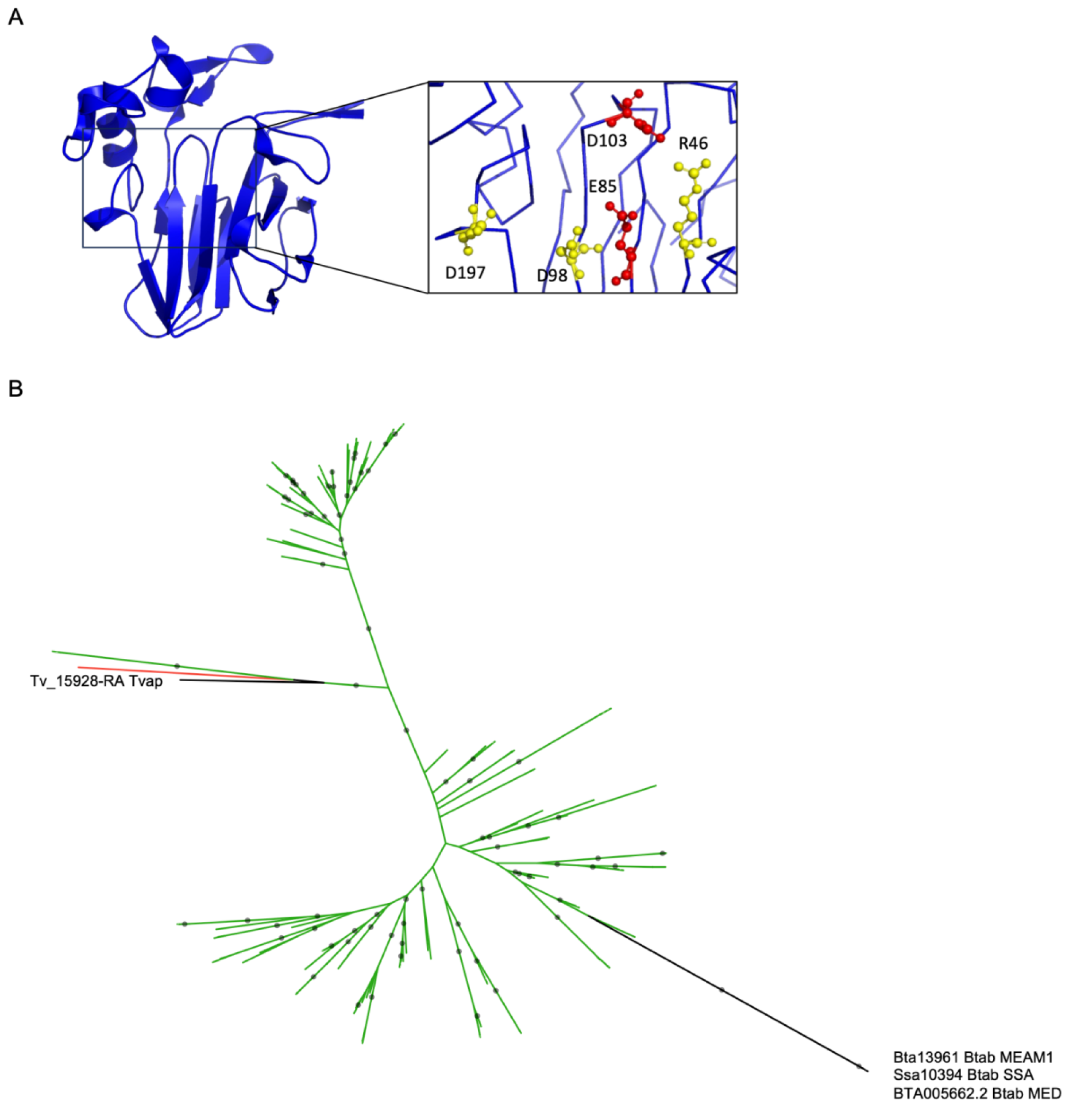
Analysis of the GH152 family protein acquired by HGT in *B. tabaci* and *T. vaporariorum*. (A) Predicted structure of *T. vaporariorum* Tv_15928-RA colored in blue. Amino acids of the REDDD motif are colored red for the two putative catalytic residues and yellow for the other three. (B) Phylogenetic tree showing *B. tabaci* and *T. vaporariorum* sequences in black. Green branches represent plant sequences, and the red branch corresponds to an animal sequence (*T. palmi* of the Thysanoptera order, accession number XP_034247376.1). Gray circles indicate nodes with support values greater than or equal to 80% and 95% for SH-aLRT and UFboot, respectively.

Among the 32 HGT events in *T. vaporariorum,* all nine for which no homologs were found in *B. tabaci* have a potential bacterial origin (Supplementary Table 6). Six of these HGT events involve genes that encode proteins from the AAA-ATPase-like domain-containing protein family. As previously mentioned, various HGT events that involve this large family of ATPases have already been identified in *B. tabaci*, and some of them are common to *T. vaporariorum* (Supplementary Table 2). The six HGT events that are unique to *T. vaporariorum* and that involve AAA-ATPase-like domain-containing proteins may correspond to new, independent HGT events of other members of this protein family. Among the three other HGT events unique to *T. vaporariorum*, one concerns the Tv_04079-RA gene, which codes for a CAZyme belonging to the α-glucosidase GH13 family (Supplementary Table 6). This *T. vaporariorum*-specific CAZyme is not related to the two *B. tabaci* GH13 CAZymes of the subfamily 17, which have been shown to catalyze the detoxification of glucosinolates produced by plants against herbivores (Malka et al. 2020), but were not identified as HGT candidates. Unfortunately, the sequence of the *T. vaporariorum* GH13 CAZyme acquired horizontally from bacteria is only partial and it is not known to which subfamily it belongs. Overall, only three of the 363 predicted *T. vaporariorum* CAZymes (Supplementary Tables 5 and 6), one PL1_4, one GH152 and one GH13, would have been acquired by HGT, compared to 25 (out of 433) in *B. tabaci* (Supplementary Tables 2 and 5). This is consistent with the GO analysis, which showed no significant enrichment of GO terms related to carbohydrate metabolism and plant cell wall degradation among HGT candidates for *T. vaporariorum* (Supplementary Table 8).

### Few HGT events in two piercing-sucking insects of the Thysanoptera order, but involving CAZymes with possible activity on plant carbohydrates

Since our results indicate that a member of the GH152 family has been acquired from plants not only in the Aleyrodinae *B. tabaci* and *T. vaporariorum*, but also in *T. palmi* (see above), we used the AvP-based approach to identify potential HGT candidates in *T. palmi* and its related species *F. occidentalis*. Both species are cell-content feeding members of the Thripinae subfamily in the Thysanoptera order, which is the closest outgroup to Hemiptera, although the divergence between these two orders probably occurred more than 300 Myr ago (Misof et al. 2014). The number of HGT candidates specific to Thysanoptera found using AvP was low in *F. occidentalis* and *T. palmi* compared to *B. tabaci* and *T. vaporariorum* (Table 1). Only 23 candidates were validated for *F. occidentalis* and only 20 for *T. palmi*. The validated HGT candidates correspond to at least eight HGT events in both *F. occidentalis* and *T. palmi*, seven of which are shared by both species (Supplementary Tables 9 and 10). Among the eight HGT events in *F. occidentalis*, we found the three that have been described in the literature with bacteria as potential donors (Rotenberg et al. 2020), further suggesting that our approach is reliable.

The analysis of overrepresented GO terms among HGT candidates in *F. occidentalis* and *T. palmi* showed a significant enrichment in GO terms related to carbohydrate metabolism and plant cell wall degradation (Supplementary Table 11). These GO terms include hydrolase activity, hydrolyzing O-glycosyl compounds (GO:0004553), cellulase activity (GO:0008810), or carbohydrate metabolic process (GO:0005975). Accordingly, five out of eight HGT events in *F. occidentalis* and six out of eight in *T. palmi* correspond to CAZymes that are predicted to originate from bacteria, fungi, and plants (Table 3, Supplementary Table 5 and Supplementary Figure 6).

**Table 3.** Horizontally acquired CAZymes in *F. occidentalis* and *T. palmi*.

| CAZy |  | Origin of donor sequences | *F. occidentalis* |  | *T. palmi* |  |
| --- | --- | --- | --- | --- | --- | --- |
| Description | Definition |  | HGT event | Sequence accession | HGT event | Sequence accession |
| GH32 | Glycoside Hydrolase Family 32 protein | Bacteria | FocB01 | XP_026273122<br>XP_026287851 | TpaB01 | XP_034233420<br>XP_034232383 |
| GH5_8 | Glycoside Hydrolase Family 5 protein | Bacteria | FocB02 | XP_026276666<br>XP_026285289<br>XP_026285291 | TpaB02 | XP_034242052<br>XP_034242578 |
| GH45 | Glycoside Hydrolase Family 45 protein | Fungi | FocF01 | XP_026287984<br>XP_026289264<br>XP_026291890 | TpaF01 | XP_034236588<br>XP_034233350 |
| GH152 | Glycoside Hydrolase Family 152 protein | Viridiplantae | FocV01 | XP_026290383 | TpaV01 | XP_034247376<br>XP_034247673 |
| CBM43-CBM43-CBM43-CBM43 | Carbohydrate-Binding Module Family 43 protein | Viridiplantae | - | - | TpaV02 | XP_034245819 |
| PL1_4 | Polysaccharide Lyase Family 1 protein | Complex: Bacteria or Fungi | FocC01 | XP_026274115<br>XP_026279513<br>XP_026279528<br>XP_026280402<br>XP_026285066<br>XP_026285067<br>XP_026285883<br>XP_026289641<br>XP_026292396<br>XP_026292397 | TpaC01 | XP_034235640<br>XP_034237191<br>XP_034239440<br>XP_034241038<br>XP_034248354<br>XP_034248552<br>XP_034253769<br>XP_034253771<br>XP_034254938 |

One of these HGT events corresponding to CAZymes is unique to *T. palmi* and involves a protein of plant origin that contains four carbohydrate-binding modules of the CBM43 family but no associated catalytic module, suggesting that the protein does not directly digest carbohydrates (Table 3 and Supplementary Figure 7). The other HGT events corresponding to CAZymes are shared by the two Thripinae species and concern the GH152 family, the GH32 family, subfamily 8 of the GH5 family (GH5_8), the GH45 family, and subfamily 4 of the PL1 family (PL1_4) (Table 3 and Supplementary Tables 9 and 10).

Our results confirm that members of the GH152 family of endoglucanases have been acquired from plants not only in the Aleyrodinae *B. tabaci* and *T. vaporariorum* (see above), but also in the Thripinae *T. palmi* and *F. occidentalis* (Table 3). In the phylogeny performed on the AvP-defined Thripinae group, the unique sequence found in *F. occidentalis* and the two sequences found in *T. palmi* form a highly-supported monophyletic group, suggesting that they originated from a single HGT event (Supplementary Figure 7). In the phylogeny performed after combining the AvP-defined Aleyrodinae and Thripinae groups, the Aleyrodinae and Thripinae sequences were not clearly separated into distinct clades (Supplementary Figure 7). In addition, the alternative topology constraining the grouping of Aleyrodinae and Thripinae sequences into a single clade was not significantly less likely. Thus, the hypothesis of a single HGT event in the common ancestor of both Aleyrodinae and Thripinae, rather than two independent HGT events, could not be rejected on the basis of the phylogeny alone. However, the hypothesis of a single HGT event would require numerous subsequent loss events. Indeed, the last common ancestor of both Hemiptera and Thysanoptera dates back more than 300 Myr (Misof et al. 2014), and no orthologs of the GH152 that we identified as acquired by HGT from plants are found in the numerous other species from these orders (particularly Hemiptera) represented at the sequence level in public libraries. Our results thus suggest a convergent acquisition of GH152 family glucanases in the Aleyrodinae *B. tabaci* and *T. vaporariorum* on the one hand, and the Thripinae *F. occidentalis* and *T. palmi* on the other.

Two sequences, which are predicted to have a bacterial origin and are found in both *F. occidentalis* and *T. palmi,* belong to the GH32 family. This family includes several horizontally-acquired genes in various animal species that are involved in plant carbohydrate assimilation, including the metabolism of host-derived sucrose (Nakabachi 2015; Danchin et al. 2016; Wybouw et al. 2016; Dai et al. 2021). The four *F. occidentalis* and *T. palmi* sequences did not form a monophyletic group, suggesting that they resulted from two distinct HGT events (Supplementary Figure 7). However, the alternative topology test using a constrained topology supporting the monophyly of these four sequences was not significantly less likely, and the hypothesis of a single event cannot be rejected. It is worth noting that members of the GH32 family have also been identified as HGT candidates in *B. tabaci* (see above). However, they form a completely distinct clade in a phylogeny performed after combining the AvP-defined *B. tabaci* and Thripinae groups (Supplementary Figure 7), with different bacteria as donors for *B. tabaci* (phylum Pseudomonodata) and for the Thripinae (phylum Actinomycetota). In addition, the alternative topology constraining all Thripinae and *B. tabaci* sequences into one clade was significantly less likely (Supplementary Figure 7). This, together with the absence of homologous sequences in other members of the Thysanoptera and Hemiptera, suggests two independent and convergent acquisition events.

Three sequences found in *F. occidentalis* and two in *T. palmi*, all predicted to originate from bacteria, belong to the GH5_8 family of endomannanases (Table 3). All five sequences form a highly-supported monophyletic group, indicating that they originated from a single HGT event (Supplementary Figure 7). GH5_8 enzymes, which degrade mannan, have reportedly been acquired by several other arthropods through HGT, possibly playing a role in plant cell wall degradation (Acuña et al. 2012; Vega et al. 2015; Shin et al. 2021).

Three sequences found in *F. occidentalis* and two in *T. palmi*, all predicted to originate from fungi, belong to the GH45 family of endoglucanases capable of hydrolyzing cellulose (Table 3). All five sequences constitute a monophyletic group supported by bootstrap, suggesting a single HGT event origin (Supplementary Figure 7). Sequences from other arthropods, as well as from nematodes and rotifers, appear in the phylogeny. Although they also share similarities with GH45 proteins from fungi, they form distinct clades in the phylogeny (Supplementary Figure 7). This was not unexpected, as it has been shown that GH45 enzymes, with cellulase activity characterized in some cases, were probably acquired by HGT independently in various phytophagous and phytoparasitic organisms (Palomares-Rius et al. 2014; Szydlowski et al. 2015; Busch et al. 2019).

The last HGT event corresponding to a CAZyme and shared by the two Thripinae species concerns 10 sequences found in *F. occidentalis* and nine in *T. palmi* belonging to the PL1_4 family of pectin lyases (Table 3). All sequences form a highly-supported monophyletic group, indicating that they originated from a single HGT event, although we could not clearly determine from the phylogeny whether the donor belonged to bacteria or fungi (Supplementary Figure 7). It is worth noting that a member of this family has been identified as an HGT candidate in *B. tabaci* and *T. vaporariorum* (see above), and that sequences from two other insects, belonging to Hemiptera and Hymenoptera, are found in the phylogeny (Supplementary Figure 7). In the phylogeny performed after combining the AvP-defined Aleyrodinae and Thripinae groups, the Hemiptera and Thysanoptera sequences were not separated into distinct clades, in contrast to the Hymenoptera sequences (Supplementary Figure 7). The alternative topology, which constrains all animal sequences to be grouped in a single clade, was not significantly less likely (Supplementary Figure 7). The hypothesis of a single ancestral HGT event could therefore not be rejected, although it would require numerous subsequent loss events. Indeed, the last common ancestor of Hemiptera, Hymenoptera, and Thysanoptera dates back approximately to 360 Myr (Misof et al. 2014), and most of the descendant species lack this gene while being represented at the sequence level in public libraries.

## Conclusion

In the present study, we used the AvP software package (Koutsovoulos et al. 2022) followed by manual validation to perform a comprehensive and accurate phylogenetic detection of HGT candidates from non-animal donors in the related phloem-feeding insects *B. tabaci* MEAM1 and *T. vaporariorum*, members of the subfamily Aleyrodinae of the order Hemiptera. With a total of 255 genes likely acquired by HGT (corresponding to 136 distinct acquisition events), our results confirm findings from previous studies indicating that a large number of genes have been acquired by HGT from bacterial, fungal, but also plant donors in *B. tabaci* (Chen et al. 2016; Gilbert and Maumus 2022; Li et al. 2022). In contrast, the number of HGT candidates in the related species *T. vaporariorum* was substantially lower, with 75 HGT candidates corresponding to a total of 32 HGT events, of which only three represented gene acquisition from plants (compared to 27 for *B. tabaci*). The majority of HGT events found in *T. vaporariorum* appear to have occurred in the common ancestor of the Aleyrodinae, whereas the majority of HGT events found in *B. tabaci* MEAM1, including the plant-acquired BtPMaT1 gene involved in detoxification of plant toxins (Xia et al. 2021), are more recent and occurred in the common ancestor of the three cryptic species of *B. tabaci*. An even lower number of HGT candidates was obtained from a similar automated HGT detection and manual validation analysis performed on the plant cell-feeding insects, *F. occidentalis* (23 HGT candidates corresponding to eight HGT events) and *T. palmi* (20 HGT candidates corresponding to eight HGT events), members of the subfamily Thripinae of the order Thysanoptera. Most of the HGT events are shared between *F. occidentalis* and *T. palmi* and probably occurred in the common ancestor of the Thripinae, in contrast to the Aleyrodinae, for which most of the HGT events are unique to *B. tabaci*. The much higher number of HGT candidates in *B. tabaci* compared to the three other species analyzed is consistent with the results of a recent study of 218 insect species, in which *B. tabaci* had by far the highest number of HGT-acquired genes (Li et al. 2022). An open question is whether variations in viral transmision capacity might explain why the number of HGT-acquired genes, especially from plants, is so high in *B. tabaci* compared to *T. vaporariorum*, but also to other piercing-sucking insect species.

In this study we focused on the horizontal acquisition of CAZymes, given their significance in phytophagous arthropods (Kirsch et al. 2014; Nakabachi 2015; Wybouw et al. 2016; Husnik and McCutcheon 2018). A total of 25 HGT-acquired CAZymes (corresponding to 14 HGT events) were identified for *B. tabaci*, with potential roles including plant cell wall degradation to facilitate stylet penetration into a phloem sieve element or assimilation of carbohydrates found in phloem. In contrast, only three CAZymes acquired by HGT were found in the related *T. vaporariorum*, one of which would correspond to a HGT event unique to this species. This indicates that HGT contributed more to the composition of CAZymes in *B. tabaci* than in *T. vaporariorum*. The 14 HGT events corresponding to CAZymes identified for *B. tabaci* involved not only bacteria or fungi as potential donors, as is typically described, but also plants for as many as seven of them. A functional analysis was conducted on two of the plant-acquired *B. tabaci* CAZymes, belonging to the GH17 and GH152 families. Both CAZymes were found to be functional glucanases, exhibiting different activities suggesting distinct functional roles. The GH17 enzyme is capable of cleaving ß-1,3-linkages and may be involved in the hydrolysis of callose, which is composed of β-1,3-glucan (Silva-Sanzana et al. 2020; Walker 2022). The occlusion of the punctured sieve element by the deposition of callose is one of the mechanisms by which plants defend themselves against phloem-feeding insects. The GH152 enzyme exhibited ß-1,3-1,4-glucanase activity, capable of cleaving both β-1,3 and β-1,4 linkages. Both GH17 and GH152 enzymes were active on mixed-linked β-glucans, but only GH152 was able to cleave the β-1,3/β-1,4 glucan with the higher ratio of β-1,4-linkages. Mixed-linked β-glucans are typically found in Poales including maize, one of the many plant hosts *of B. tabaci* (Quintela et al. 2016).

A member of the GH152 family was also identified as having been acquired from plants in *T. vaporariorum*, suggesting a unique HGT event in the ancestor of the Aleyrodinae. However, our results do not exclude the possibility that it originated from a distinct HGT event. It is noteworthy that a GH152 CAZyme with a plant as a potential donor was identified as well in the Thripinae *F. occidentalis* and *T. palmi*, although it was probably acquired independently. This suggests a convergent acquisition of a GH152 CAZyme in the unrelated Aleyrodinae and Thripinae, indicating the potential importance of this enzyme for piercing-sucking insects. Although the number of HGT events identified for *F. occidentalis* and *T. palmi* was relatively low compared to the Aleyrodinae, the majority of them corresponded to the acquisition of CAZymes, several of which had predicted functions important in the plant-insect interaction. This indicates that HGT-acquired CAZymes have biological significance not only for Aleyrodinae, and particularly *B. tabaci*, but also for other piercing-sucking insects, including the Thripinae *F. occidentalis* and *T. palmi*.

## Material and Methods

### Data used and quality control

Genome and predicted proteome data from the *Bemisia tabaci* MEAM1 (v1.2) (Chen et al. 2016), MED (v1.0) (Xie et al. 2017), and SSA-ECA (v1.0) (Chen et al. 2019) cryptic species and from *Trialeurodes vaporariorum* (v1.0) (Xie et al. 2020) were obtained from the Whitefly Genome Database (http://www.whiteflygenomics.org).

Genome and predicted proteome data from *Frankliniella occidentalis* (assembly GCF_000697945.2_Focc_2.1) and *Thrips palmi* (assembly GCF_012932325.1_TpBJ-2018v1) were obtained from the NCBI database (https://www.ncbi.nlm.nih.gov/).

Proteome completeness was compared between *B. tabaci* MEAM1, MED and SSA-ECA cryptic species, *T. vaporariorum*, *F. occidentalis* and *T. palmi* using BUSCO (v5) according to the Arthropoda Odb10 dataset (Manni et al. 2021).

### Orthogroup inference

Orthogroups (i) between the *B. tabaci* MEAM1, MED and SSA-ECA cryptic species and *T. vaporariorum* and (ii) between *F. occidentalis* and *T. palmi* were defined using Orthofinder (v2) with default parameters (Emms and Kelly 2015; Emms and Kelly 2019).

### Functional annotation

All the predicted proteins from the *B. tabaci* MEAM1, *T. vaporariorum*, *F. occidentalis* and *T. palmi* genomes were analyzed using InterProScan (v5) to identify conserved protein domains (Jones et al. 2014). The -iprlookup and -goterms options were used to assign Gene Ontology (GO) terms from the identified InterPro domains. Signal peptide prediction was obtained from the Phobius tool integrated within InterProScan (v5).

### Detection of HGT candidates

A homology search against the NCBI non-redundant (NR) protein database was performed for the *B. tabaci* MEAM1, *T. vaporariorum*, *F. occidentalis* and *T. palmi* proteomes using DIAMOND (v2) (Buchfink et al. 2021). The DIAMOND search was performed in the more sensitive mode with an e-value threshold of 1.0e^−3^ and a maximum number of hits of 500.

The DIAMOND homology search results were submitted to AvP (Koutsovoulos et al. 2022) to calculate the Aggregate Hit Score (AHS) (Koutsovoulos et al. 2022) for each query sequence based on the normalized sum of the scores of the best animal (NCBI:txid33208) and non-animal hits. This AHS metric was shown to be less sensitive to contamination and taxonomic assignment errors in databases than the classical Alien Index (AI) (Koutsovoulos et al. 2022), which only considers the single best animal and single best non-animal hits. Proteins with an AHS greater than 0 have a higher similarity to non-animal than to animal hits in the NR database. Self-hits to Aleyrodinae (NCBI:txid33379) were ignored for *B. tabaci* MEAM1 and *T. vaporariorum.* Self-hits to Thysanoptera (NCBI:txid30262) have been ignored for *F. occidentalis* and *T. palmi*.

We again used AvP for automatic phylogenetic detection of HGT candidates among proteins with AHS above 0. The first step of AvP was to cluster the query sequences based on the percentage of shared hits in the DIAMOND homology search result (default 70% overlap). This clustering method was supplemented with orthogroups inferred by Orthofinder (see above) and shared Pfam domains as determined by InterProScan (see above). When analyzing the AVP results, we retained the most comprehensive clustering method, for which only one HGT event was predicted from the obtained phylogeny. All protein sequences with significant hits in the DIAMOND homology search result were retrieved from the NR database by AvP and aligned using MAFFT (v7) with the --auto option (Katoh and Standley 2013) for each group. The second step of AvP was to infer the phylogeny for each group and to detect HGT candidates according to the species found in the sister branch of the query sequence and the ancestral sister branch. FastTree (v2) with the default parameters (Price et al. 2010) was preferred to IQ-TREE (v2) (Minh et al. 2020) for phylogeny inference in this initial step to improve the speed of the analysis. In a third step, AvP classified the HGT candidates according to their putative origin. The fourth step of AvP was to infer a constrained topology in which the query sequence(s) and the other animal sequence(s), if present, form a single monophyletic group and to determine whether the topology supporting HGT is significantly more likely than the constrained alternative one using an approximately unbiased (AU) test (Shimodaira 2002). Finally, AvP analyzed the genomic environment of each HGT candidate gene and calculated a local score ranging from -1 to +1, with -1 for a HGT candidate gene surrounded by other HGT candidate genes (indicating a possible contamination, duplications after an HGT event, or multiple HGTs) or +1 for a HGT candidate gene surrounded by genes that are likely to be vertically inherited (suggesting that the contamination hypothesis can be ruled out). The local score was not reported if the number of genes in the scaffold (including the HGT candidate) was less than 5.

### Validation of HGT candidates

A manual analysis of the AvP results was performed, and HGT candidates were not further considered if at least one of the following criteria was met:

(i) The total number of donor sequences was less than 3 in the sister branch plus the ancestral sister branch (if present).
(ii) The HGT-supporting topology was not significantly more likely than the constrained alternative topology in which the query was forced to group with all animal sequences from NR. However, HGT candidates were retained if these animal sequences were either (i) taxonomically misannotated, likely due to contamination by a donor species or (ii) suspected to have originated from an HGT event themselves based on BLASTP results performed at NCBI against NR (https://www.ncbi.nlm.nih.gov/).
(iii) At least one donor sequence in the DIAMOND homology search results shared more than 70% identity with the HGT candidate sequence and either (i) no homologous sequence was found in the closely related cryptic species for *B. tabaci* MEAM1, suggesting contamination by a donor species, or (ii) the donor sequences were suspected to be misannotated, probably in the case of Viridiplantae due to contamination of the donor species by the query species (or a related insect species), based on BLASTP results performed at NCBI against NR (https://www.ncbi.nlm.nih.gov/).
(iv) The identity between the donor sequences and the HGT candidate sequence in the DIAMOND homology search results was less than 30%. Searching for homologous proteins and building a phylogenetic tree below this value is problematic due to the low quality of the alignments.
(v) The average alignment length in the DIAMOND homology search results was less than 100 amino acids with an average query coverage of less than 50%.
(vi) The local score calculated by AvP from the genomic environment of the HGT candidate was less than 0, and there was no indication of duplication after an HGT event or of multiple HGTs, indicating a possible contamination.

### Phylogeny reconstruction for validated HGT candidates

For each HGT candidate detected by AvP and satisfying all the above-mentioned criteria, we reconstructed a maximum likelihood phylogeny with IQ-TREE (v2) (Minh et al. 2020) to improve accuracy. Before reconstructing the phylogeny, the AvP-defined groups were refined by the following three preliminary steps:

(i) We searched for homologous sequences of each HGT candidate validated for *B. tabaci* MEAM1 in the cryptic species MED and SSA and included them in the groups formed by AvP. The search for homologous sequence(s) in the cryptic species MED and SSA was first performed in the orthogroups determined by Orthofinder (see above). If no homologous sequence was found, we performed a protein-to-genome comparison using exonerate (v2) with a score threshold of 500 (Slater and Birney 2005). The *B. tabaci* MEAM1 HGT candidate protein sequence was used as the query and the genome of each cryptic species as the database. When a significant hit was found, the protein sequence was inferred from the cryptic species genomic sequence. The homologous sequence(s) found for the cryptic MED and SSA species were added to the *B. tabaci* MEAM1 groups previously defined by AvP.
(ii) We combined the groups defined by AvP for *B. tabaci* MEAM1 and *T. vaporariorum* on the one hand and *F. occidentalis* and *T. palmi* on the other hand, where the sequences of the two related species were in the same orthogroup defined by Orthofinder. If no homologous sequence was found for a validated HGT candidate in the related species, a protein-to-genome comparison was performed using exonerate (v2) as described above.
(iii) Animal sequences suspected to be misannotated, probably due to contamination, based on BLASTP results performed at NCBI against NR (https://www.ncbi.nlm.nih.gov/) were removed from the groups.

For each group, a CD-HIT analysis was then performed with an identity threshold of 100% to remove redundancy (Li and Godzik 2006). Sequences were aligned using MAFFT (v7) with the --auto option (Katoh and Standley 2013). Poorly aligned regions were removed using trimal (v1.4) with the - automated1 option (Capella-Gutiérrez et al. 2009). Phylogenies were inferred using IQ-TREE (v2) (Minh et al. 2020) with automated model selection (Kalyaanamoorthy et al. 2017). Support values were based on a Shimodaira–Hasegawa approximate likelihood ratio test (SH-aLRT) (Guindon et al. 2010) combined with an ultrafast bootstrap (UFboot) approximation with 1,000 replicates (Hoang et al. 2018). Only support values greater than or equal to 80% and 95% for SH-aLRT and UFboot, respectively, were considered. Phylogenies were visualized using iTol (Letunic and Bork 2021).

When *B. tabaci* MEAM1 and *T. vaporariorum* sequences on the one hand and *F. occidentalis* and *T. palmi* sequences on the other hand did not form monophyletic groups, we forced a constrained topology supporting monophyly of animal sequences and determined whether the unconstrained topology was significantly more likely than the constrained alternative one using an approximately unbiased (AU) test (Shimodaira 2002) with IQ-TREE (v2).

### Identification of overrepresented GO terms among validated HGT candidates

Identification of overrepresented Gene Ontology (GO) terms among validated HGT candidates was performed using a hypergeometric test as implemented in func (Prüfer et al. 2007) within the R package GOfuncR with a family-wise error rate threshold of 0.05. The -refine option was used to eliminate redundancy between GO terms and to keep only representative terms.

### Detection of encoded Carbohydrate-active enzymes (CAZymes)

All the predicted proteins from the, *T. vaporariorum*, *F. occidentalis* and *T. palmi* genomes were compared with entries in the CAZy database (Drula et al., 2022). A homemade pipeline combining the BlastP (https://blast.ncbi.nlm.nih.gov/Blast.cgi) and HMMER3 (http://hmmer.janelia.org/) tools was used to compare protein models with the sequences of the CAZy modules. Proteins with E-values less than 0.1 were further screened by a combination of BlastP searches against libraries generated from the sequences of the catalytic and non-catalytic modules. HMMER3 was used to search against a collection of custom Hidden Markov Model (HMM) profiles constructed for each CAZy family. Expert curators then performed manual inspection to resolve borderline cases.

### Structural analysis

Structure prediction was performed using the ColabFold v1.5.2 implementation of AlphaFold2 (Mirdita et al. 2022) at https://colab.research.google.com/github/sokrypton/ColabFold/blob/main/AlphaFold2.ipynb with the following parameters: num_relax=1 and template_mode=pdb100. Secondary structure assignment was performed with the DSSP program (https://www3.cmbi.umcn.nl/xssp/) (Kabsch and Sander 1983). Visualization of predicted protein structures, alignment of the alpha carbon atoms with crystallized structures, and calculation of the root-mean-square deviation (RMSD) value were performed using PyMOL (http://sourceforge.net/projects/pymol/).

### Synteny analysis

Synteny analysis was performed using MCScanX (Wang et al. 2012), with additional manual inspection.

### Recombinant production of CAZymes

The recombinant proteins were produced using the in-house 3PE Platform (*Pichia Pastoris* Protein Express; www.platform3pe.com/) as described in Haon et al. (2015). The sequences of the genes coding for the GH17 Bta06115 and the GH152 Bta13961 from *B. tabaci* were synthesized after codon optimization for expression in *Pichia pastoris* and inserted into a modified pPICZαA vector using *Xho*I and *Xba*I restriction sites in frame with the α secretion factor at the N terminus (i.e., without native signal peptide) and with a (His)_6_-tag at the C terminus (without *c*-myc epitope) (Genewiz, Leipzig, Germany). Transformation of competent *P. pastoris* X33 cells (Invitrogen, Carlsbad, California, USA) was performed by electroporation with PmeI-linearized pPICZαA recombinant plasmids as described in Haon et al. (2015). Zeocin-resistant transformants were then screened for protein production.

### Heterologous protein production in flasks

The best-producing transformants (GH17 Bta06115 and GH152 Bta13961) were grown in 2 L BMGY medium (10 g.L^-1^ glycerol, 10 g.L^-1^ yeast extract, 20 g.L^-1^ peptone, 3.4 g.L^-1^ YNB, 10 g.L^-1^ ammonium sulfate, 100 mM phosphate buffer pH 6 and 0.2 g.L^−1^ of biotin) at 30°C and 200 rpm to an optical density at 600 nm of 2-6. Expression was induced by transferring cells into 400 mL of BMMY media at 20 °C in an orbital shaker (200 rpm) for another 3 days. Each day, the medium was supplemented with 3% (v/v) methanol. The cells were harvested by centrifugation, and just before purification, the pH was adjusted to 7.8 and was filtered on 0.45-µm membrane (Millipore, Burlington, Massachusetts, USA).

### Purification by affinity chromatography

Filtered culture supernatant was loaded onto a 20 ml HisPrep FF 16/10 column (Cytiva, Vélizy-Villacoublay, France) equilibrated with buffer A (Tris-HCl 50 mM pH 7.8, NaCl 150 mM, imidazole 10 mM) that was connected to an Äkta pure (Cytiva). (His)_6_-tagged recombinant proteins were eluted with buffer B (Tris-HCl 50 mM pH 7.8, NaCl 150 mM, imidazole 250 mM). Fractions containing recombinant enzymes were pooled, concentrated, and dialyzed against Tris-HCl 50 mM pH 7.8, NaCl 150 mM.

Concentration of purified protein was determined by absorption at 280 nm using a Nanodrop ND-2000 spectrophotometer (Thermo Fisher Scientific) with calculated molecular mass and molar extinction coefficients derived from the sequences. Proteins were loaded onto a 10% Tris-glycine precast SDS-PAGE gel (BioRad, Gemenos, France) which was stained with Coomassie Blue. The molecular mass under denaturing conditions was determined with reference standard proteins (Page Ruler Prestained Protein Ladder, Thermo Fisher Scientific, Waltham, MA, USA).

### Functional enzymatic assays

Enzyme assays (final reaction volume 200 µL) were performed at 30°C in 2 mL Eppendorf tubes, incubated in a thermomixer (Eppendorf, Hamburg, Germany) at 1000 rpm for 24 h in sodium phosphate buffer (50 mM, pH 7.0). Polysaccharides (Barley β-glucan, Lichenan) (MEGAZYME, Bray, Ireland) were used at 10 mg.mL^-1^, and oligosaccharides (laminarihexaose, cellohexaose) (MEGAZYME) at a concentration of 0.5 mM. Reactions were initiated by the addition of Bta06115 and Bta13961 at a final concentration of 8 µM. Reactions were stopped by heating at 100°C for 5 min, centrifuged (12,000*xg*, 2 min, 4°C), and diluted 10-fold in milliQ H_2_O before injection onto the HPAEC column.

### HPAEC-PAD analyses

The detection method is performed using a high-performance anion-exchange chromatography (HPAEC) coupled with pulsed amperometric detection (PAD) (DIONEX ICS6000 system, Thermo Fisher Scientific). The system is equipped with a CarboPac-PA1 guard column (2 x 50 mm) and a CarboPac-PA1 column (2 x 250 mm) kept at 30°C. Elution was carried out at a flow rate of 0.25 mL.min^-1^ and 25 µL of samples was injected. The eluents used were 100 mM NaOH (eluent A) and NaOAc (1M) in 100 mM NaOH (eluent B). The initial conditions were set to 100% eluent A, and the following gradient was applied: 0 to 10 min, 0 to 10% B; 10 to 35 min, 10 to 35% B (linear gradient); 35 to 40 min, 30 to 100% B (curve 6); 40 to 41 min, 100 to 0% B; 41 to 50 min, 100% A. Integration was performed using the Chromeleon 7.2.10 data software.

## Acknowledgements

We are grateful to the BIG bioinformatics platform from the PlantBios infrastructure for providing facilities and technical support (ISC PlantBios, https://doi.org/10.15454/qyey-ar89) and to the genotoul bioinformatics platform Toulouse Occitanie (Bioinfo Genotoul, https://doi.org/10.15454/1.5572369328961167E12) for providing computing resources. Part of the work described was performed using services provided by the 3PE platform, a member of IBISBA-FR (https://doi.org/10.15454/08BX-VJ91; www.ibisba.fr), the French node of the European research infrastructure, EU-IBISBA (www.ibisba.eu).

## Data and resource availability

Phylogenetic trees generated by IQ-TREE for validated HGT candidates from potential bacterial, fungal, or viridiplantae donors for *B. tabaci*, *T. vaporariorum*, *F. occidentalis*, and *T. palmi* are available as supplementary datasets online at Data INRAE: https://doi.org/10.57745/6GC9WA

## Author contributions

Conceived and designed the study: DC, JGB, EGJD Performed experiments: DC, MH, MB, SG, GDK Analyzed the data: DC, ED, MB, SG, JGB, EGJD Contributed analysis tools: CB Wrote the manuscript: DC, EGJD, with contribution from JGB All authors read and approved the final version of the manuscript.

