## Supplementary Figures for "Functional carbohydrate-active enzymes acquired by horizontal gene transfer from plants in the whitefly *Bemisia tabaci*"

**Supplementary Figure 1.**
Phylogeny inferred for the HGT events corresponding to CAZymes in *B. tabaci* and *T. vaporariorum.* Gray circles indicate nodes with support values greater than or equal to 80% and 95% for SH-aLRT and UFboot, respectively.

**Donor: Bacteria**
**BtaB53: GH32**

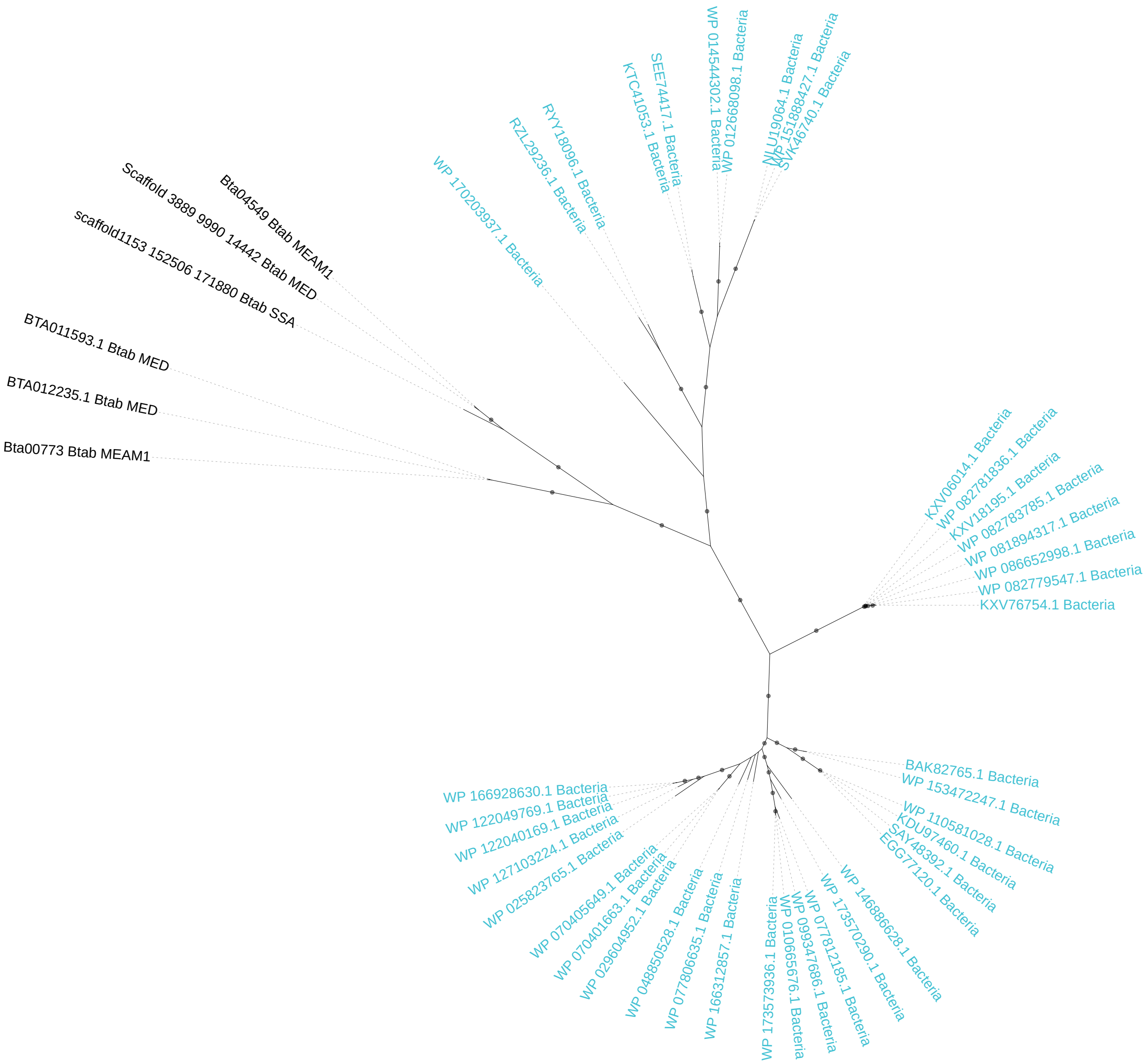

**Donor: Bacteria**
**BtaB62: CBM50**

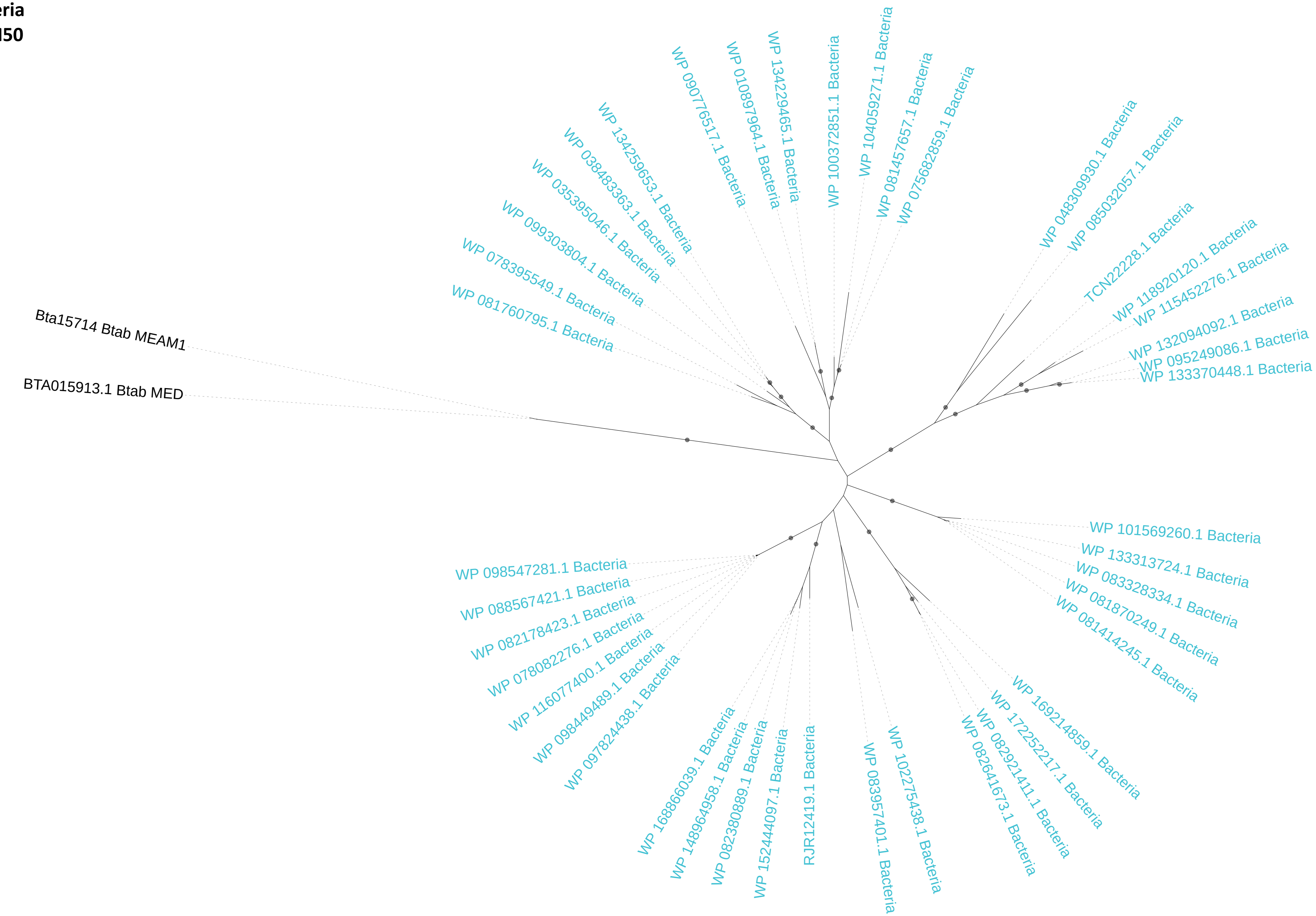

**Donor: Bacteria**
**TvaB10: GH13**

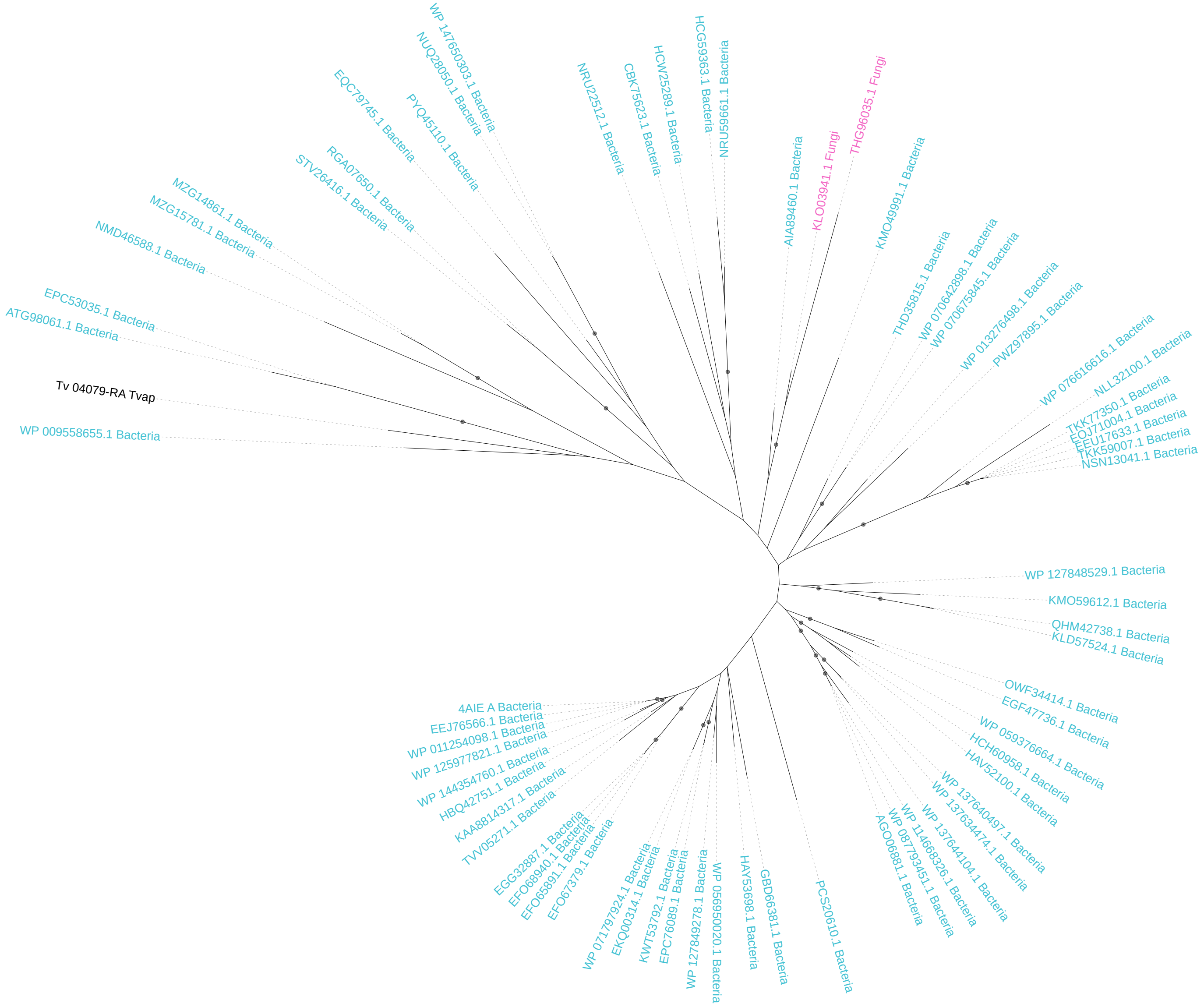

**Donor: Fungi**
**BtaF06: GH49**

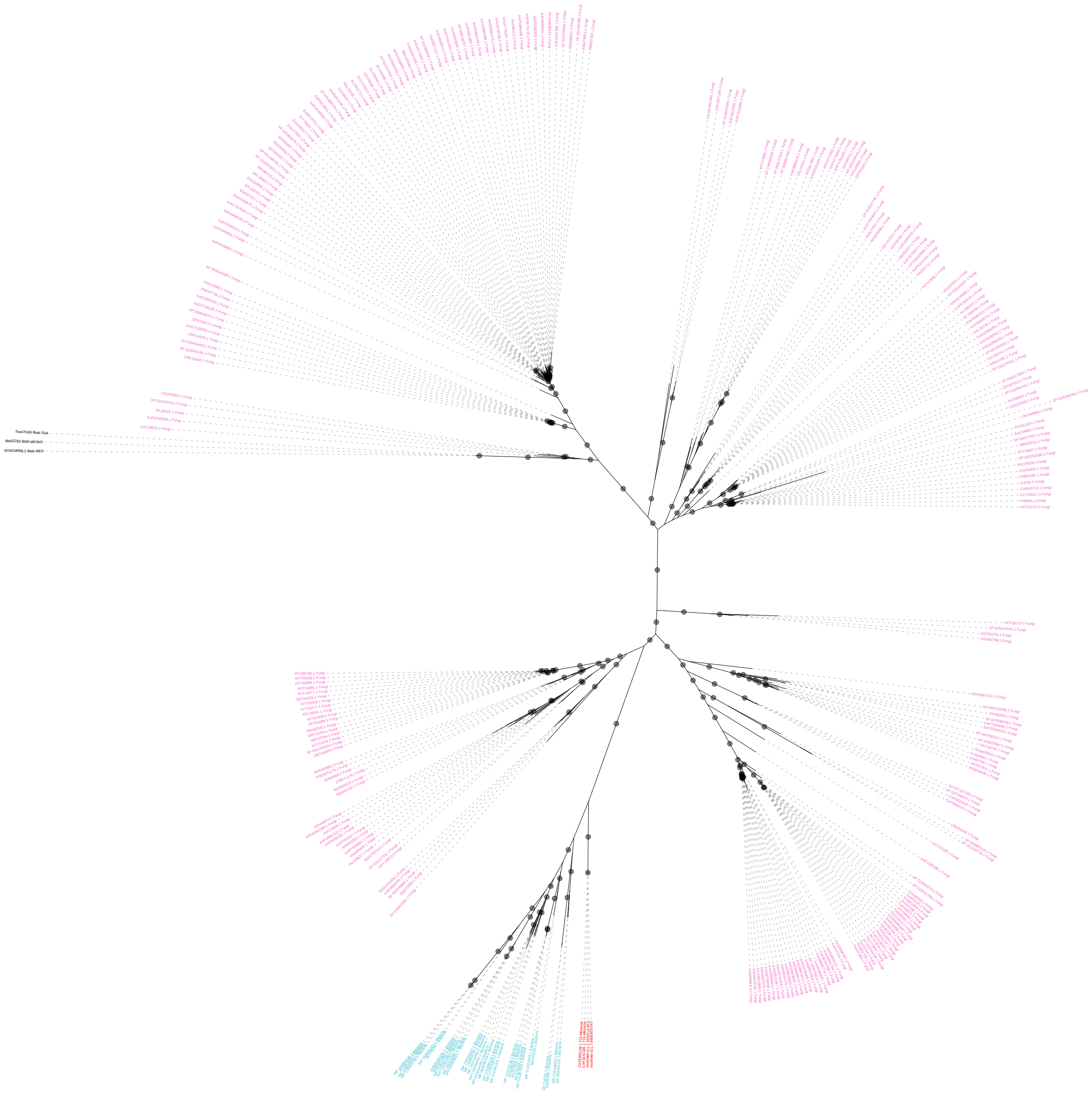

**Donor: Fungi**
**BtaF15: GH71**

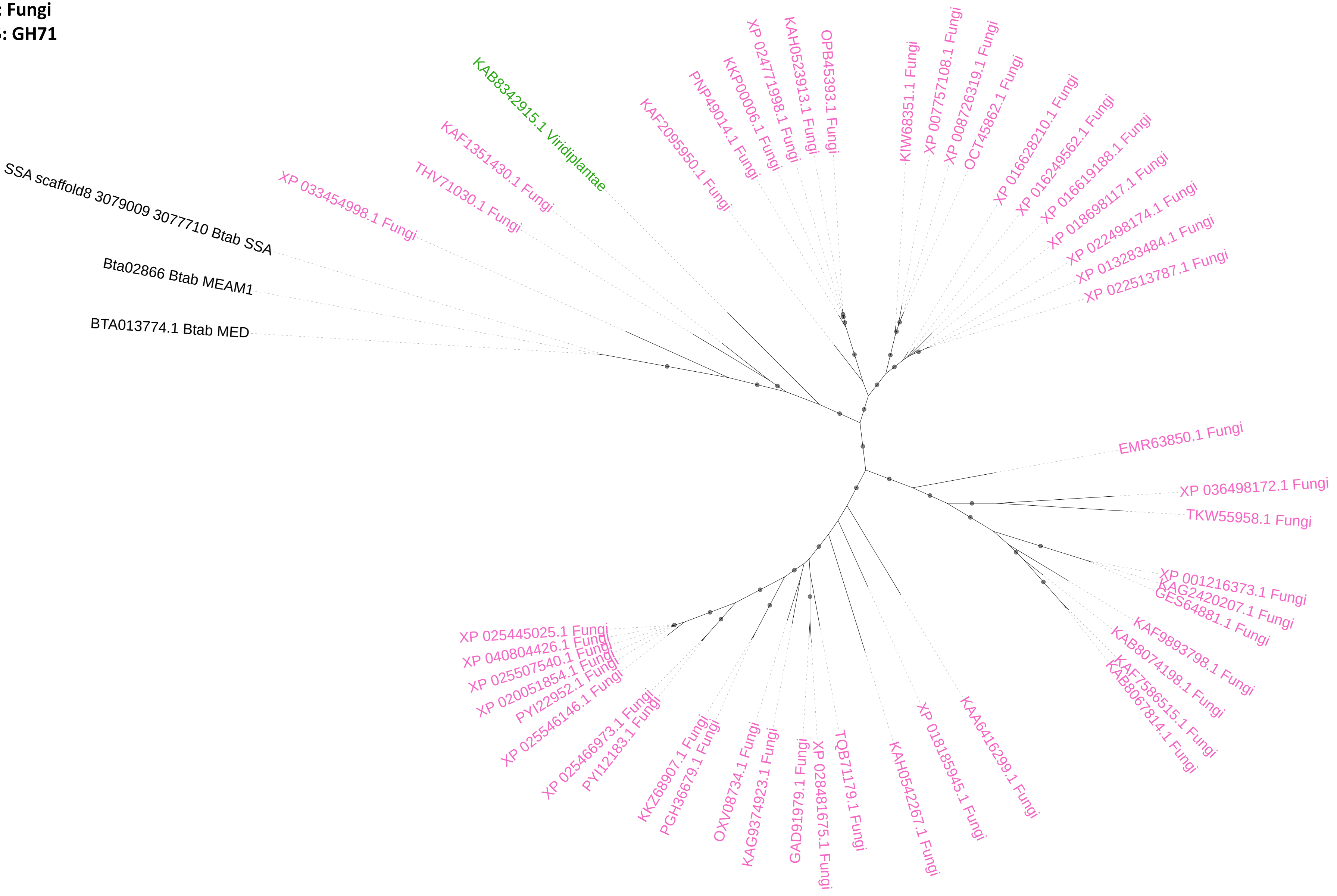

**Donor: Fungi**
**BtaF18: CBM32-AA5_2**

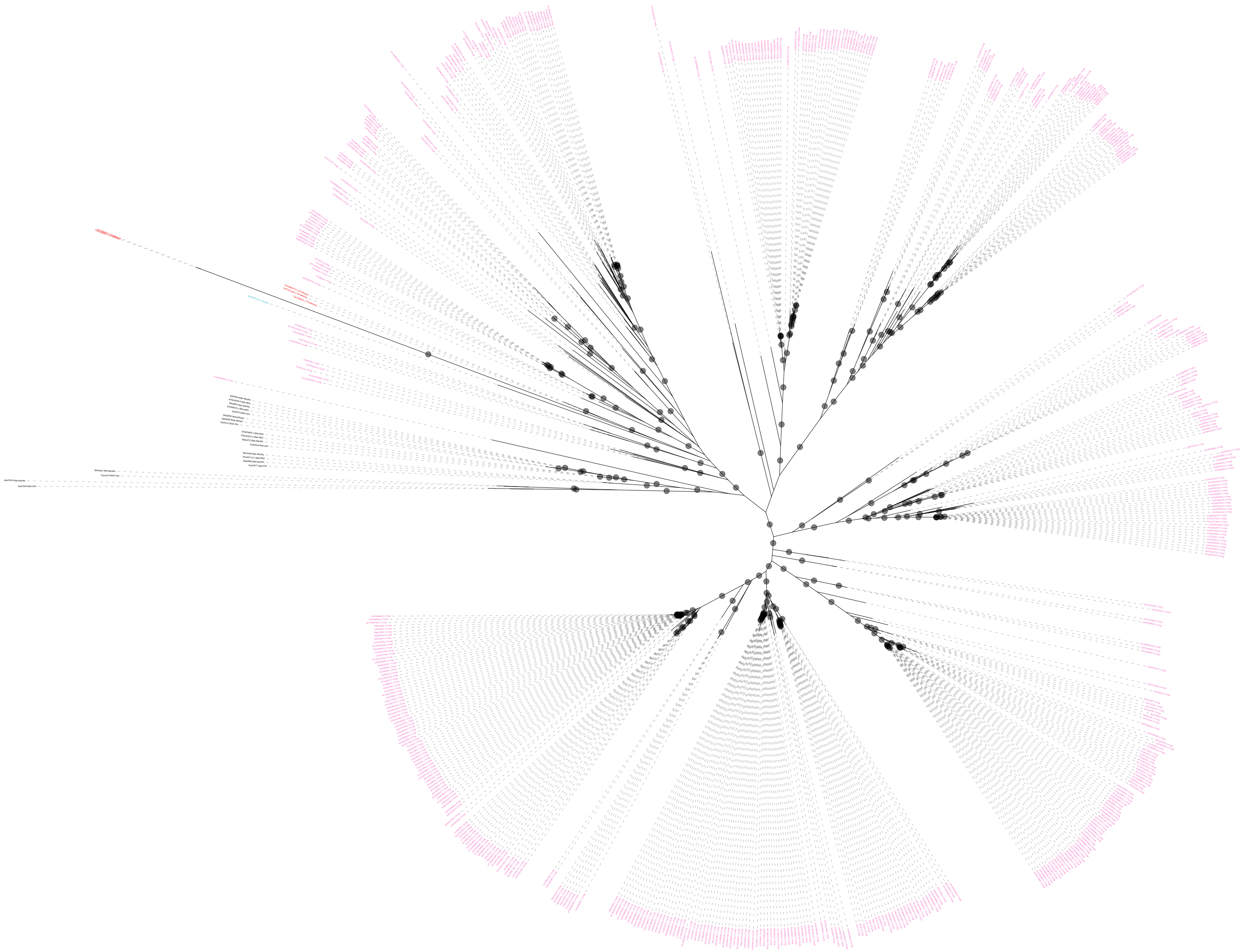

**Donor: Fungi**
**BtaF20: GH30_3**

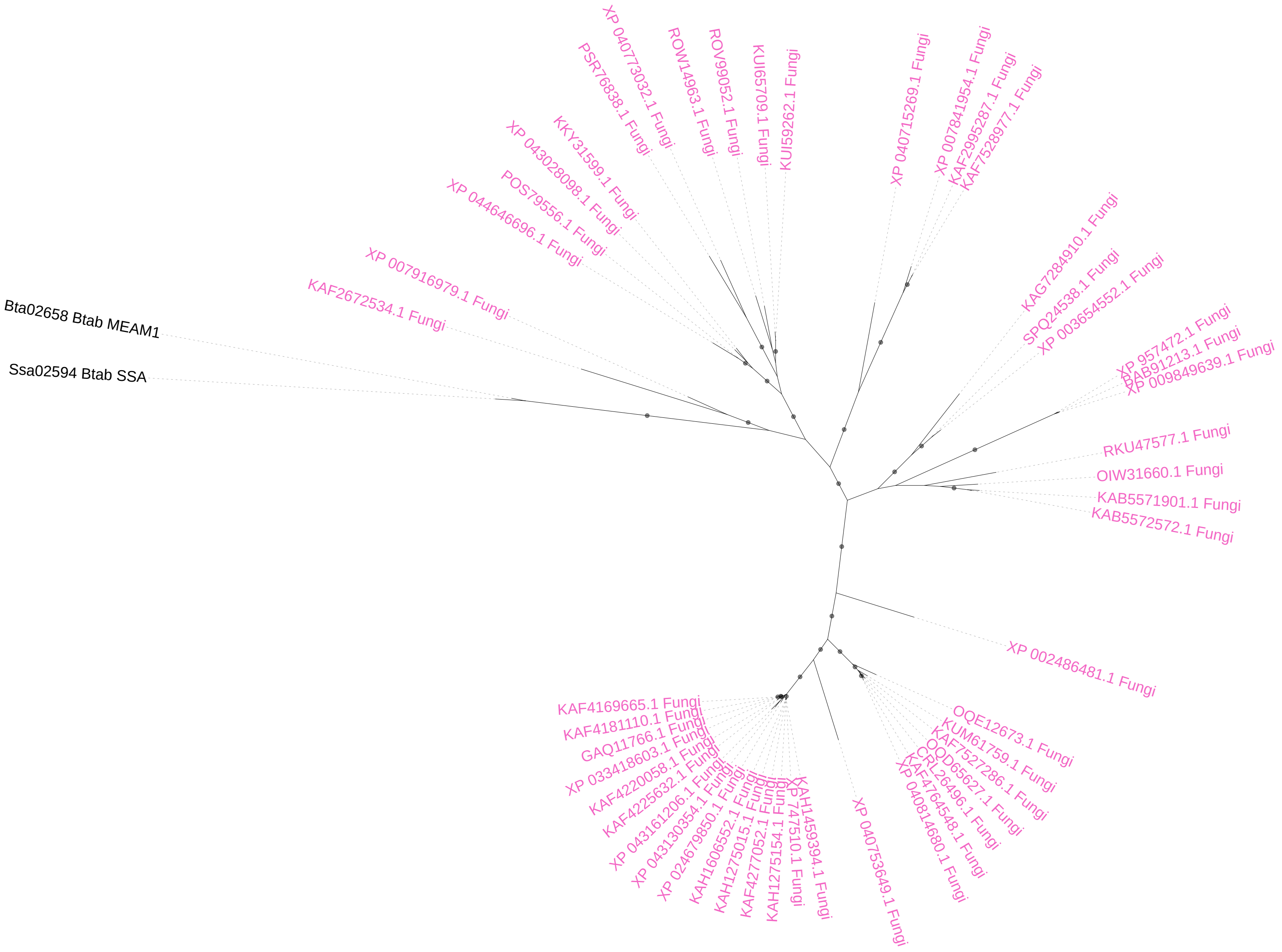

**Donor: Viridiplantae**
**BtaV01_TvaV01: GH152**

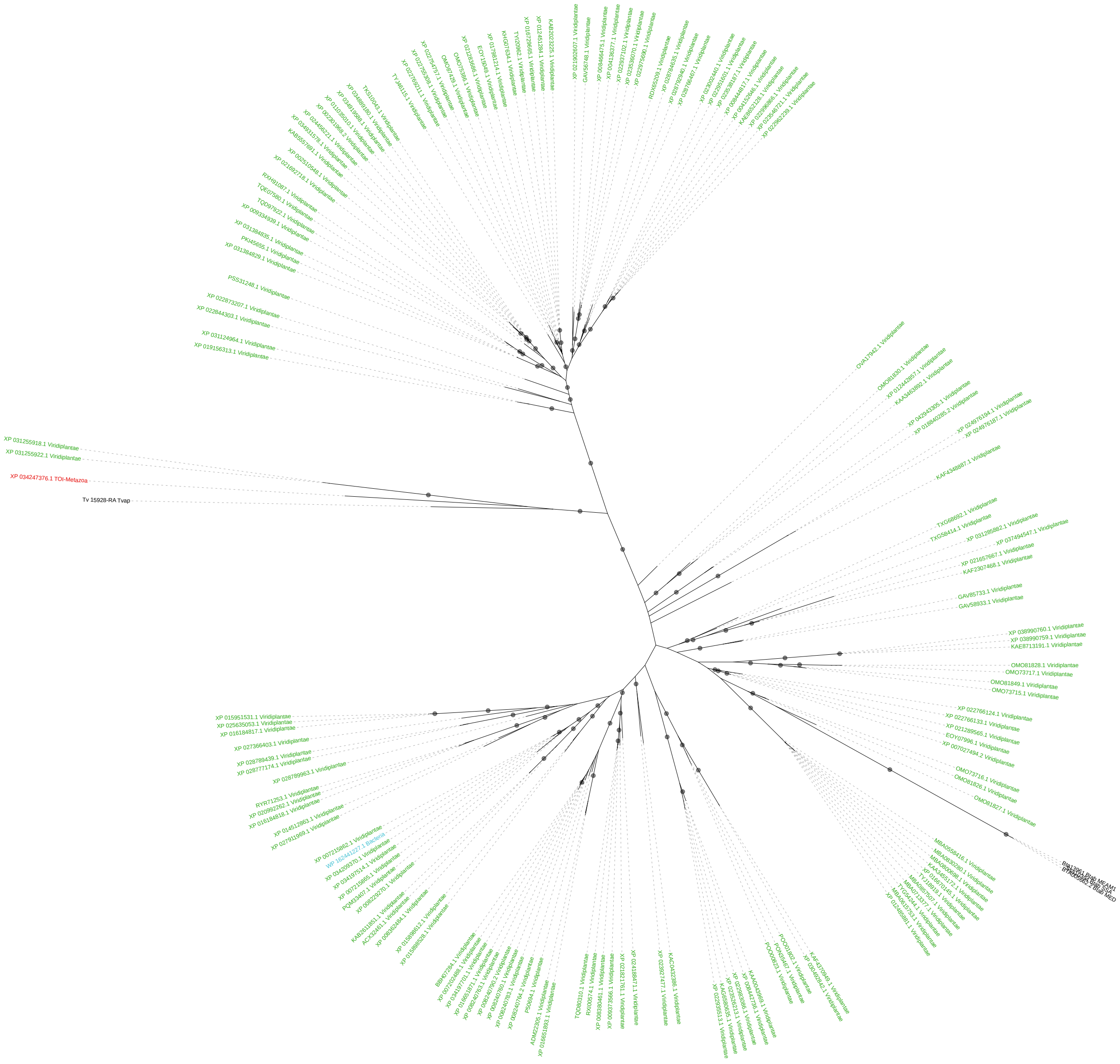

**Donor: Viridiplantae**
**BtaV10: GT10**

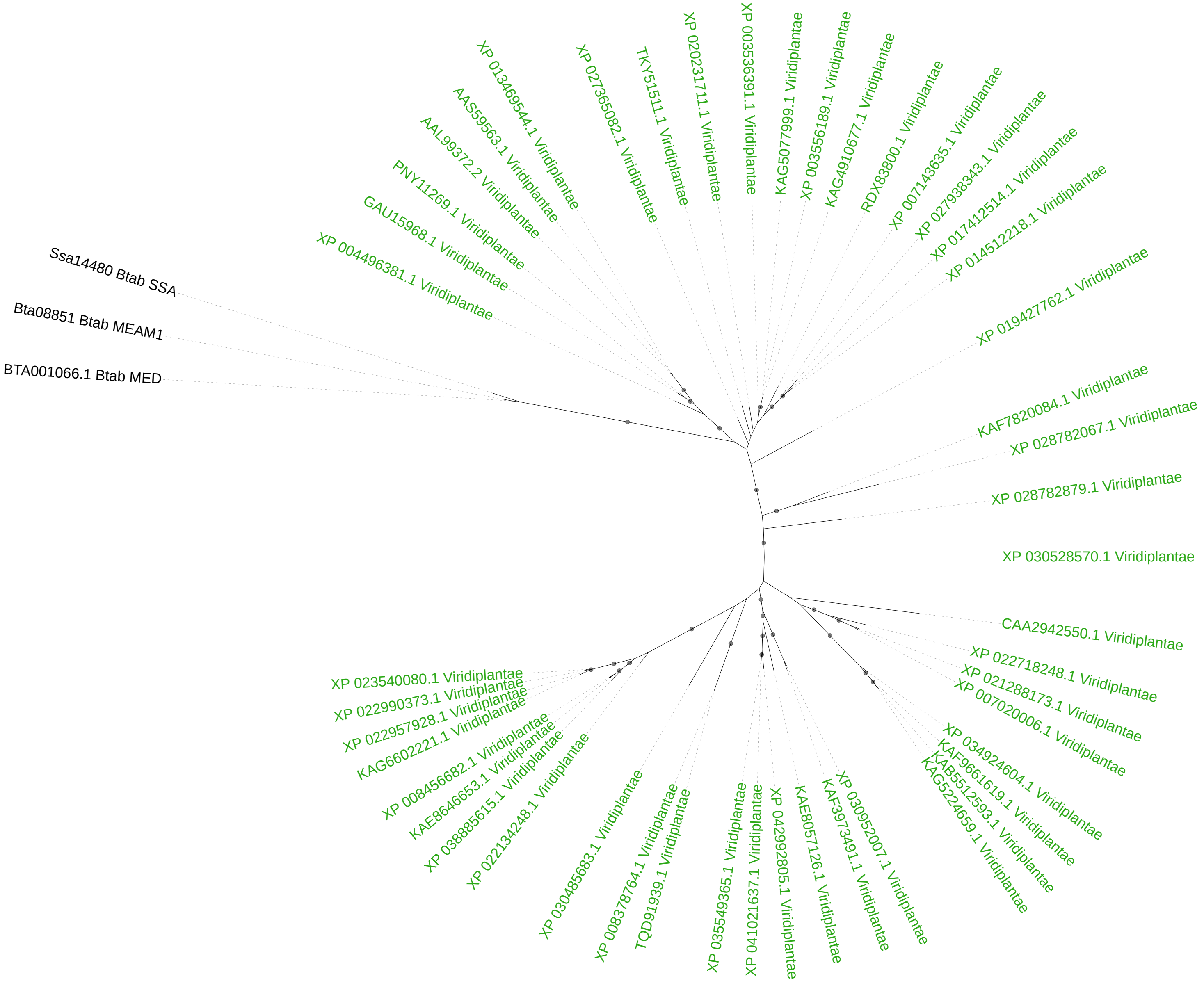

**Donor: Viridiplantae**
**BtaV12: GH17**

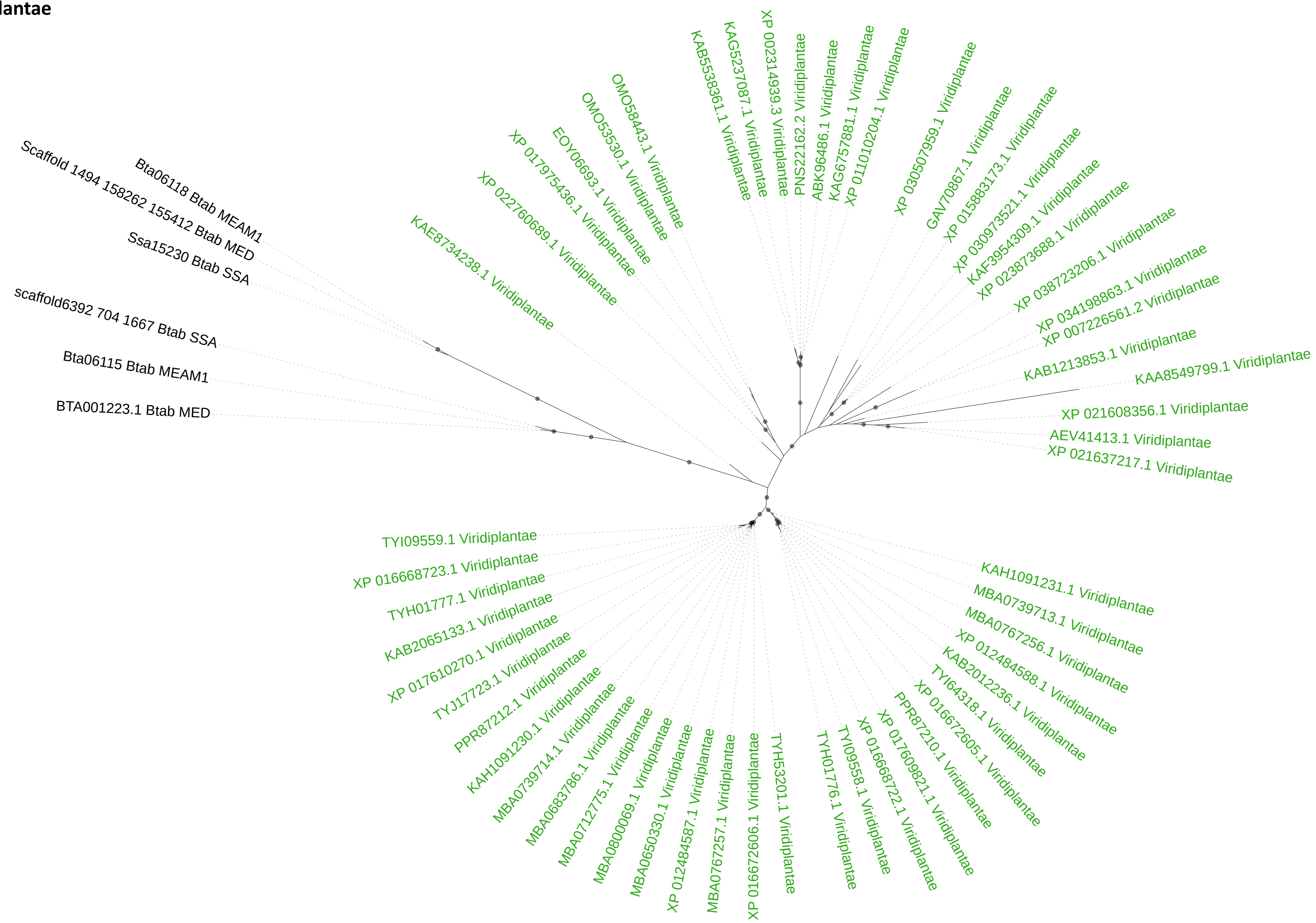

**Donor: Viridiplantae**
**BtaV13: EXPN**

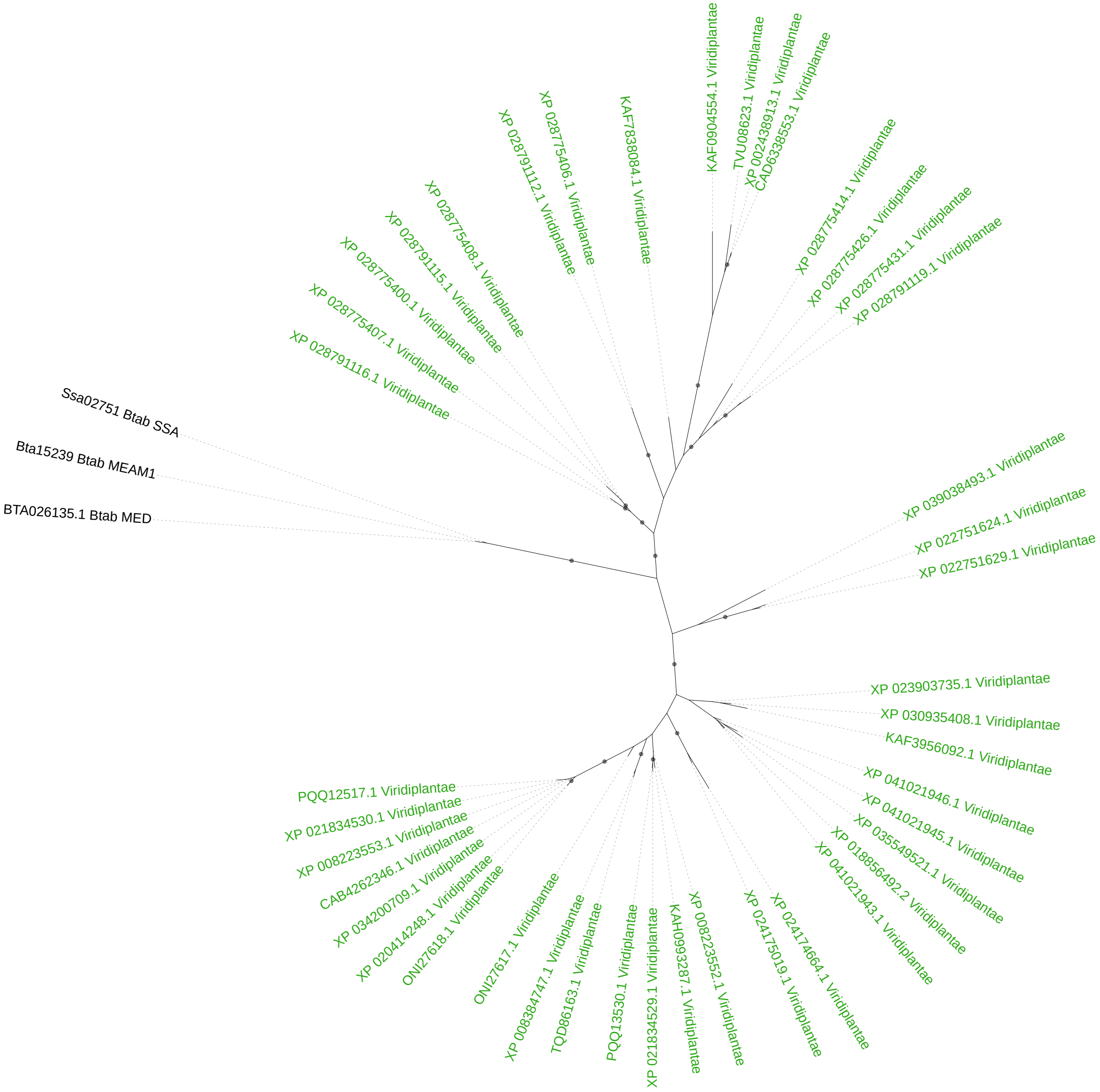

**Donor: Viridiplantae**
**BtaV24: CE8**

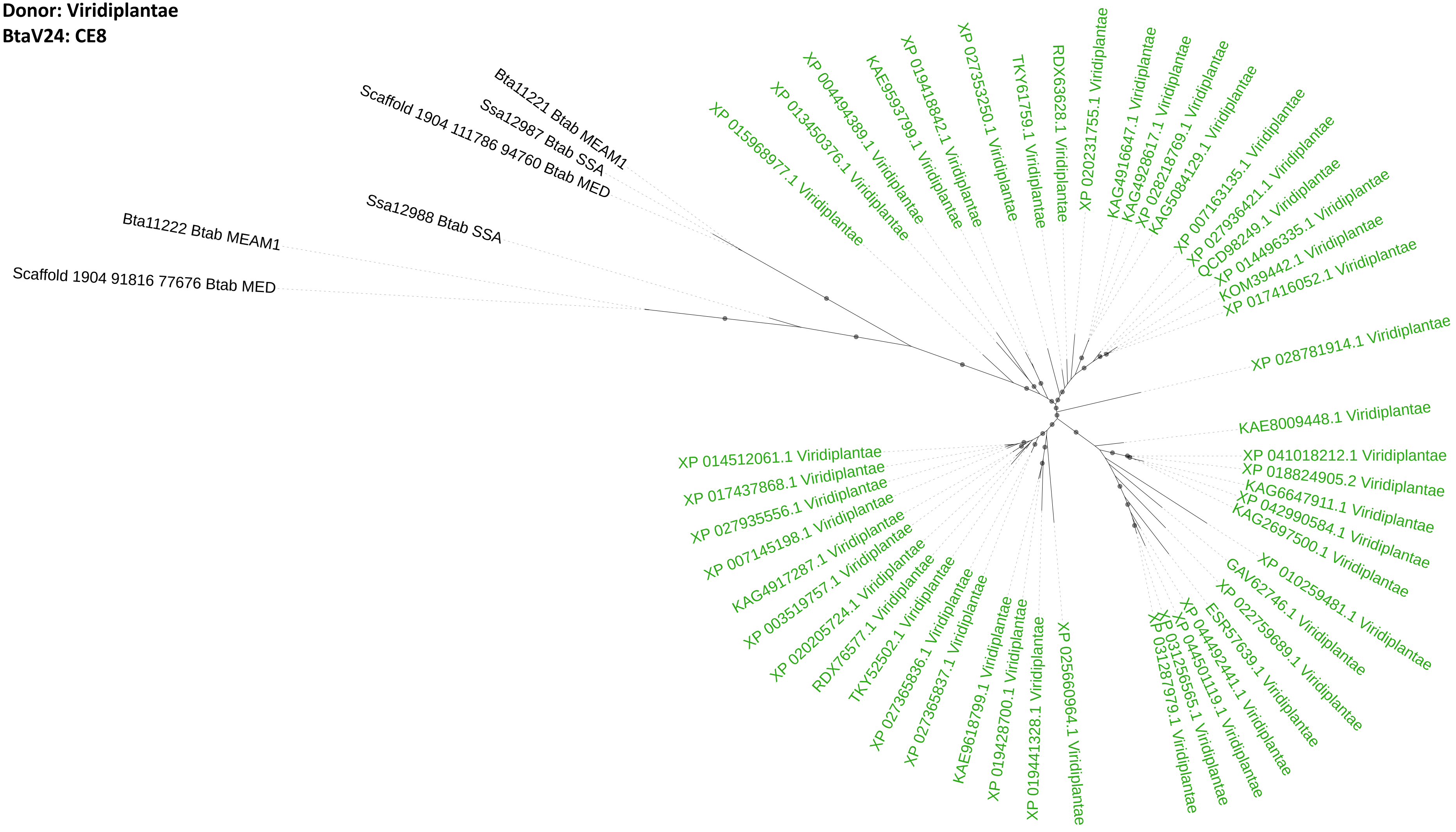

**Donor: Viridiplantae**
**BtaV25: GT61**

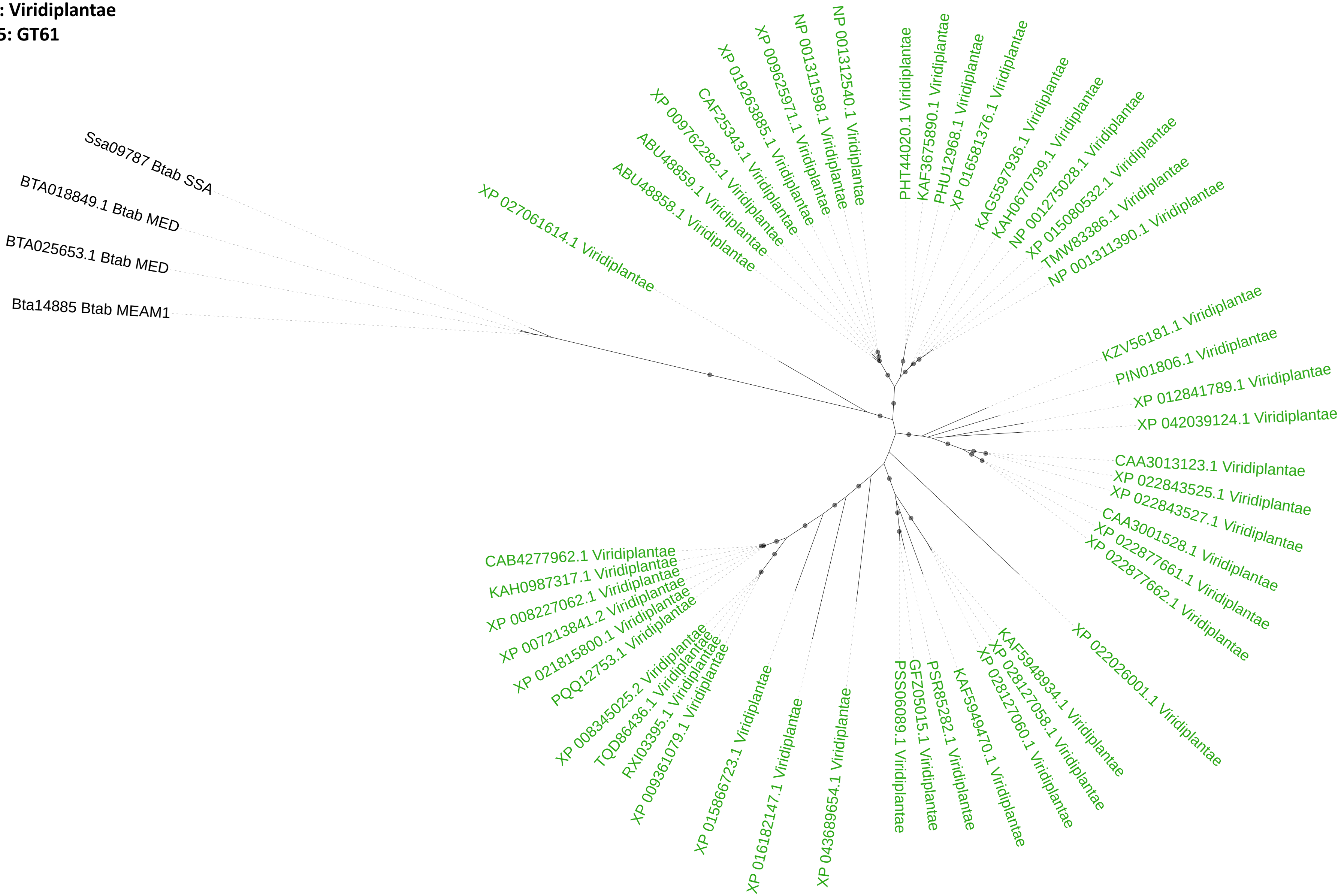

**Donor: Viridiplantae**
**BtaV27: GT17**

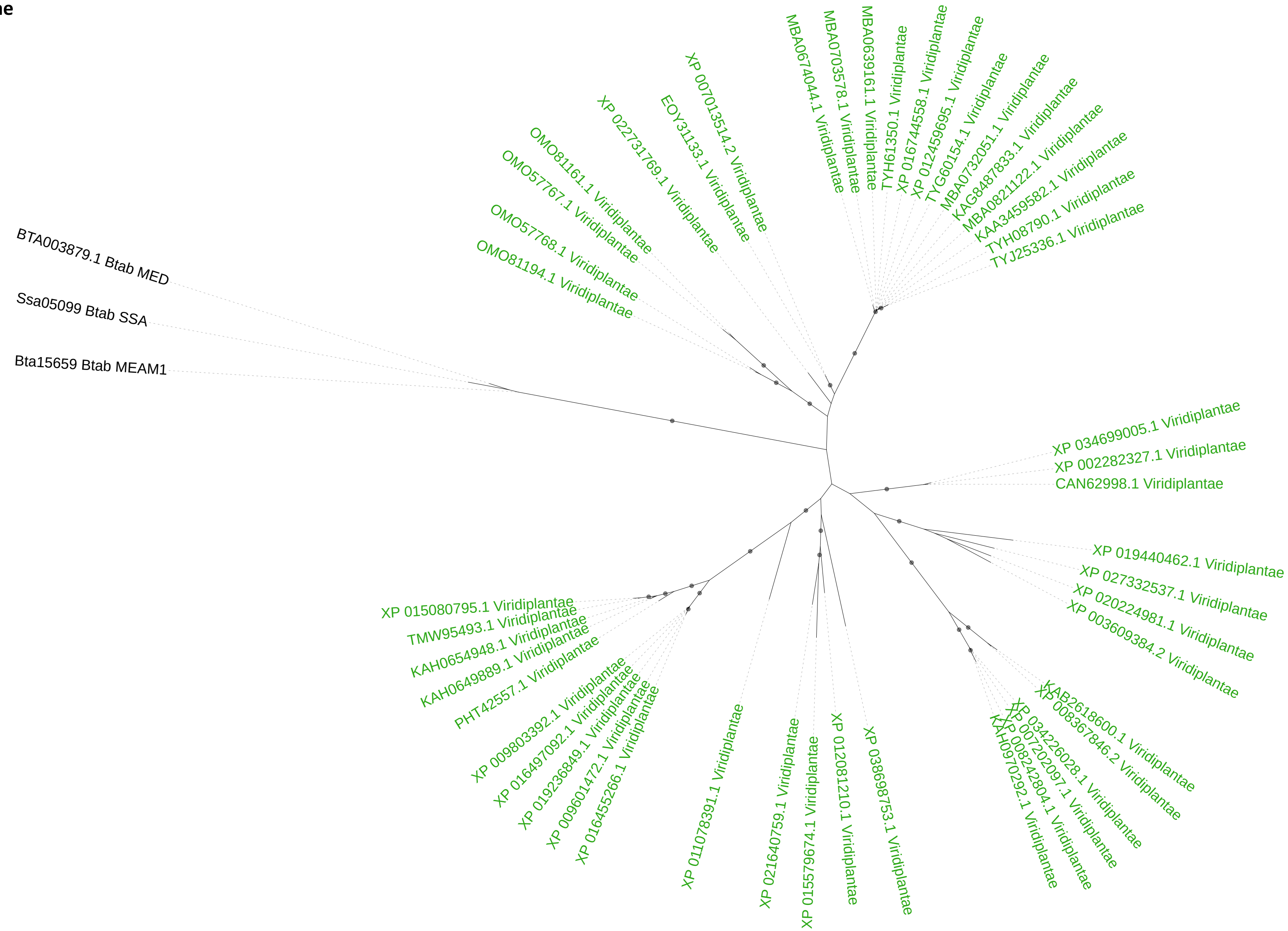

**Donor: Bacteria or Fungi**
**BtaC01_TvaC01: PL1_4**

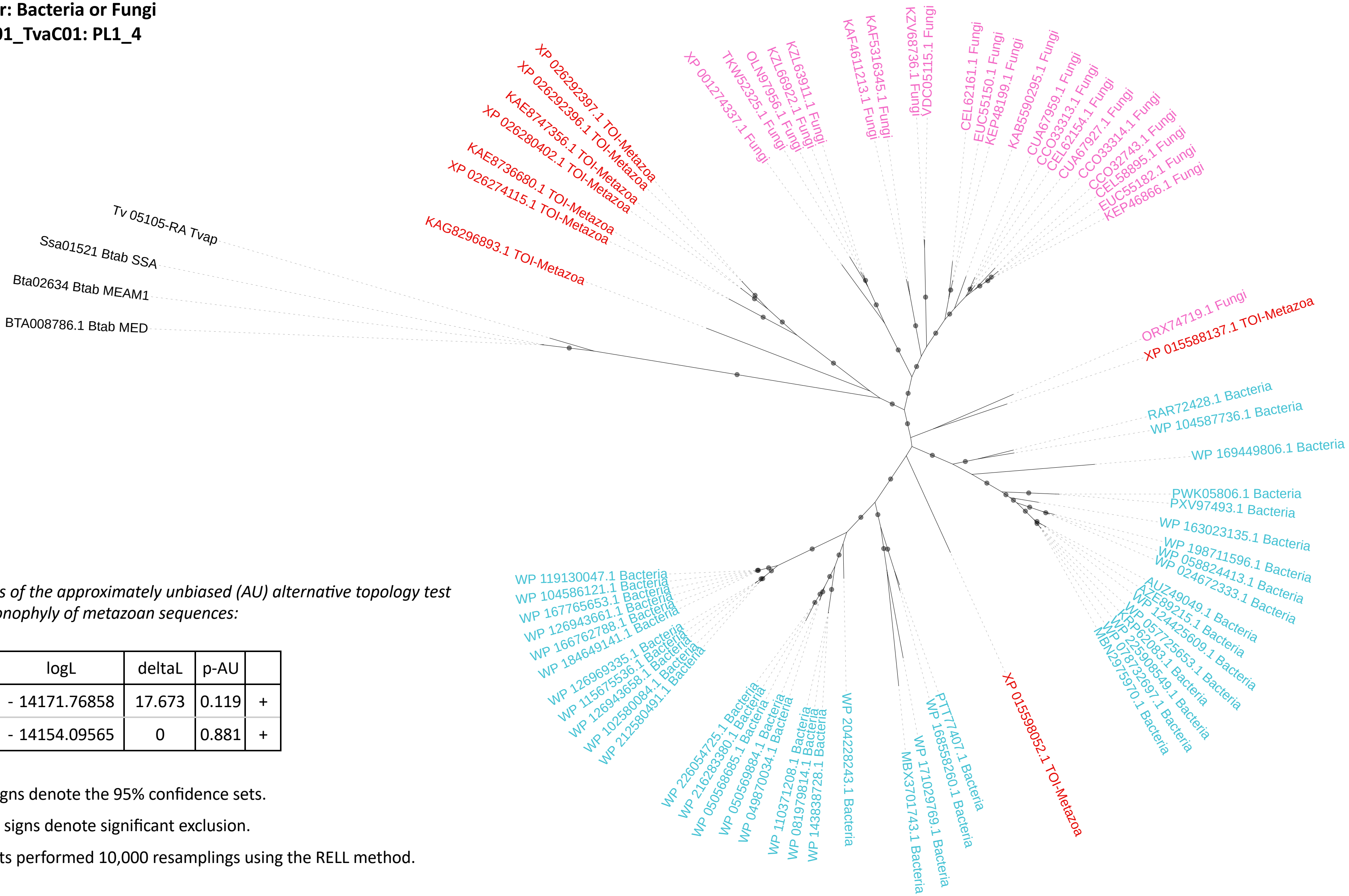


*Results of the approximately unbiased (AU) alternative topology test for monophyly of metazoan sequences:*

| Tree | logL | deltaL | p-AU | |
|---|---|---|---|---|
| 1 | - 14171.76858 | 17.673 | 0.119 | + |
| 2 | - 14154.09565 | 0 | 0.881 | + |

Plus signs denote the 95% confidence sets.

Minus signs denote significant exclusion.

All tests performed 10,000 resamplings using the RELL method.

**Supplementary Figure 2A.** Synteny analysis performed between the *B. tabaci* MEAM1, *B. tabaci* MED and *B. tabaci* SSA genomes in the region corresponding to the *B. tabaci* MEAM1 Bta06115 and Bta06118 genes. Accession numbers in green correspond to the *B. tabaci* MEAM1 Bta06115 and Bta06118 genes and their orthologs in *B. tabaci* MED and *B. tabaci* SSA.

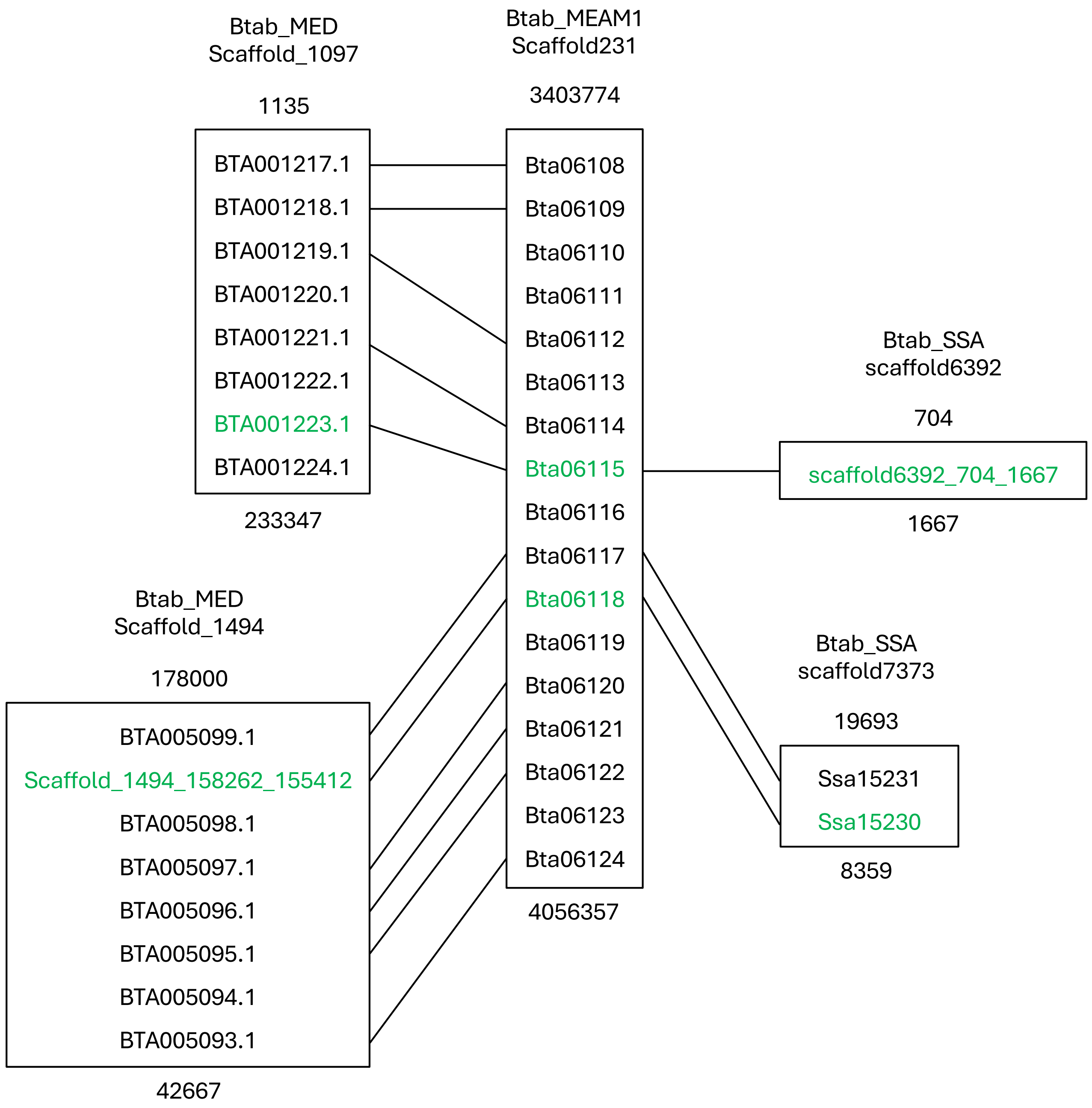

**Supplementary Figure 2B.** Synteny analysis performed between the *B. tabaci* MEAM1, *B. tabaci* MED and *B. tabaci* SSA genomes in the region corresponding to the *B. tabaci* MEAM1 Bta13961 gene. Accession numbers in green correspond to the *B. tabaci* MEAM1 Bta13961 gene and its orthologs in *B. tabaci* MED and *B. tabaci* SSA.

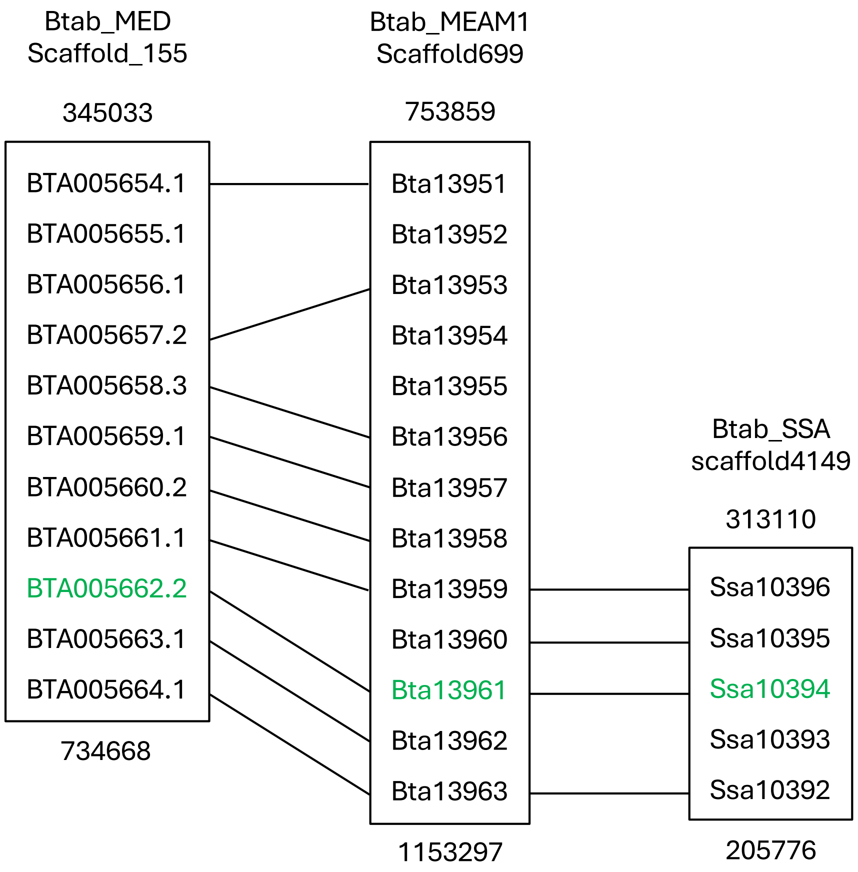

**Supplementary Figure 2C.** Synteny analysis performed between the *B. tabaci* MEAM1, *B. tabaci* MED and *B. tabaci* SSA genomes in the region corresponding to the *B. tabaci* MEAM1 Bta11221 and Bta11222 genes. Accession numbers in green correspond to the *B. tabaci* MEAM1 Bta11221 and Bta11222 genes and their orthologs in *B. tabaci* MED and *B. tabaci* SSA.

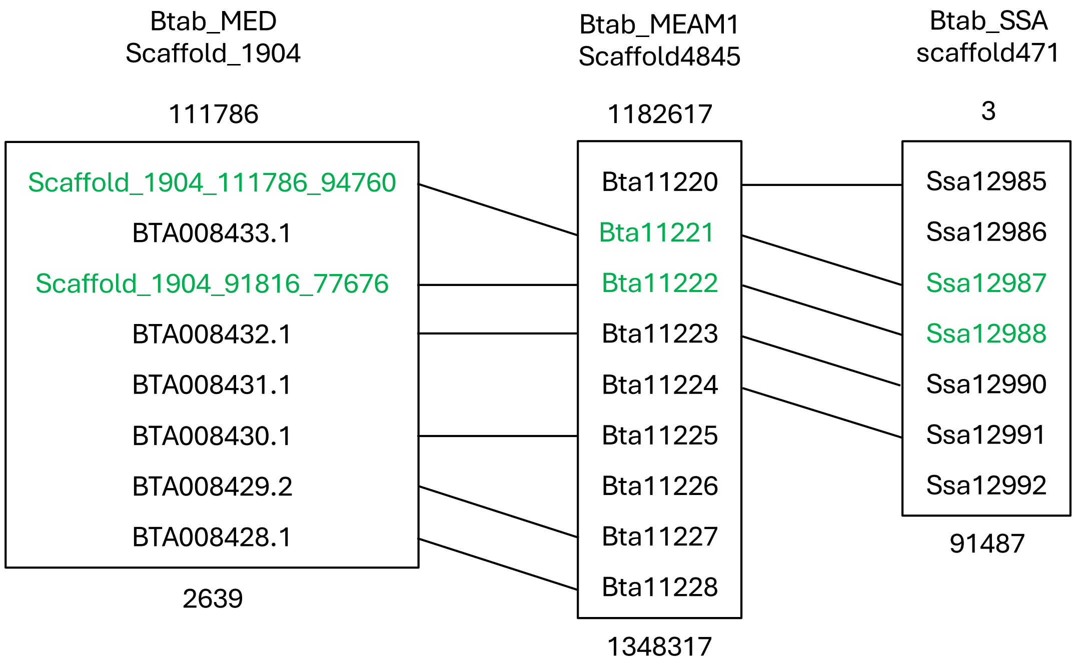

**Supplementary Figure 2D.** Synteny analysis performed between the *B. tabaci* MEAM1, *B. tabaci* MED and *B. tabaci* SSA genomes in the region corresponding to the *B. tabaci* MEAM1 Bta15239 gene. Accession numbers in green correspond to the *B. tabaci* MEAM1 Bta15239 gene and its orthologs in *B. tabaci* MED and *B. tabaci* SSA*.*

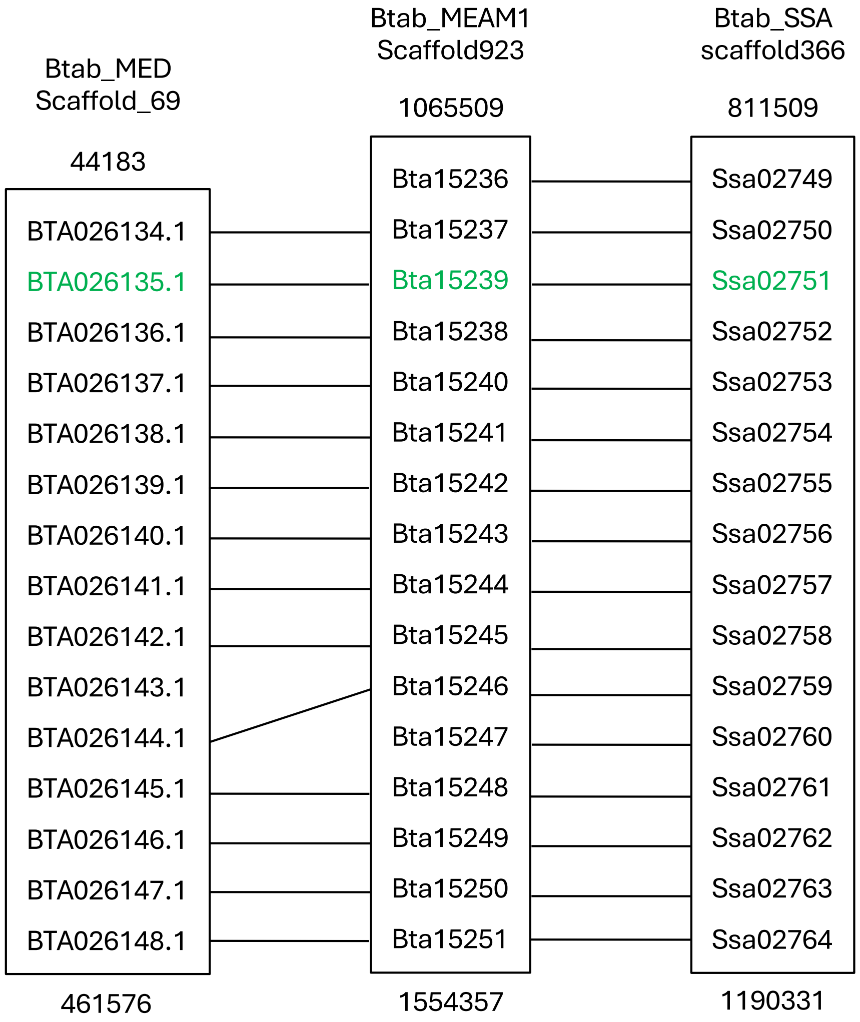

**Supplementary Figure 2E.** Synteny analysis performed between the *B. tabaci* MEAM1 and *T. vaporariorum* genomes in the regions corresponding to the *B. tabaci* MEAM1 Bta13961 gene and to the *T. vaporariorum* Tv_15928-RA gene, respectively. The accession number in green corresponds to the *B. tabaci* MEAM1 Bta13961 gene. The accession number in red corresponds to the *T. vaporariorum* Tv_15928-RA gene.

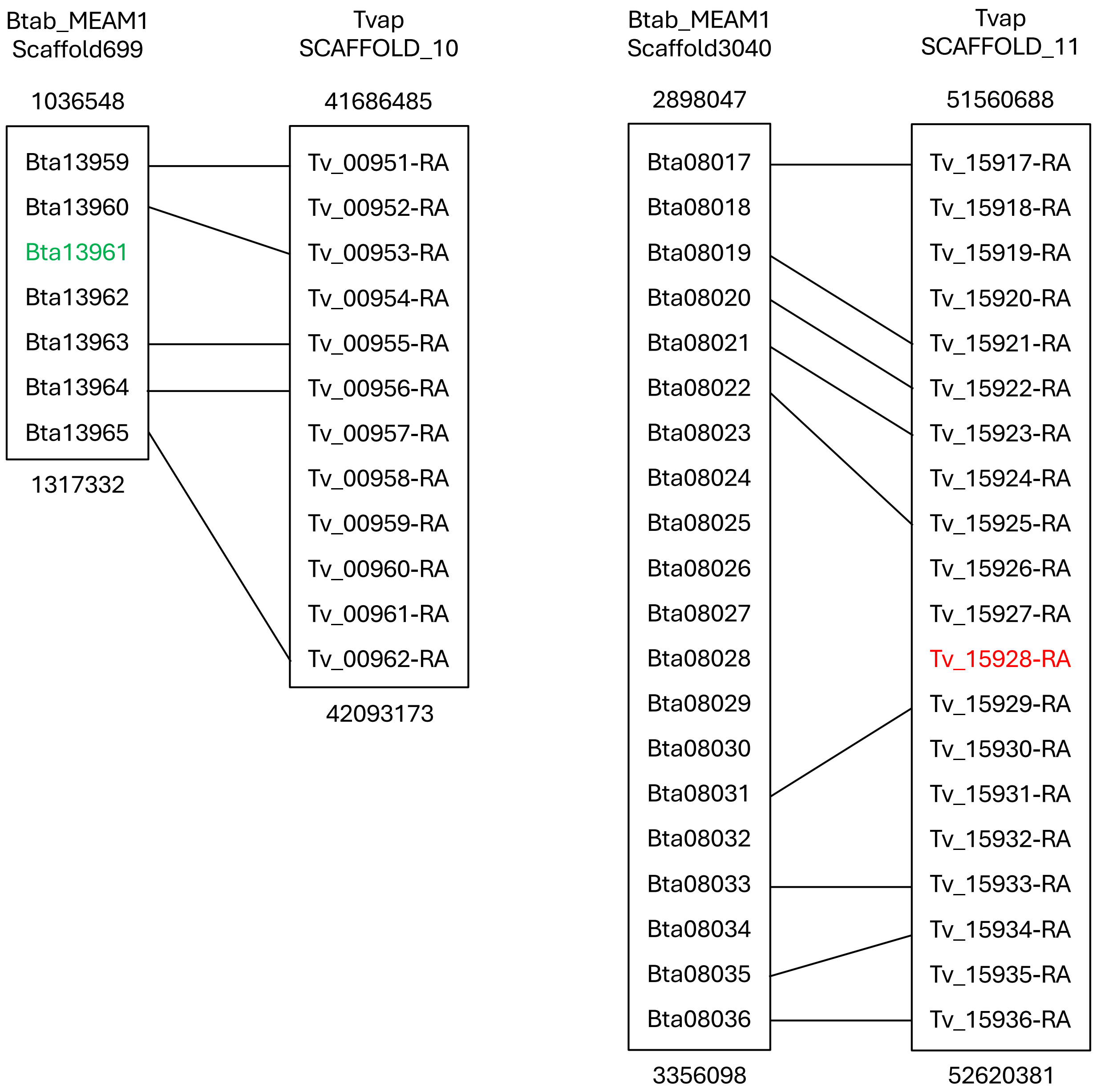

**Supplementary Figure 3.** Comparison of the predicted structures of Bta06115 and Bta06118 proteins with the known structures of barley (1-3)-ß-glucanase and barley (1- 3,1-4)-ß-glucanase. The $\alpha$-helices are colored red, ß-sheets are yellow, and loops are green.

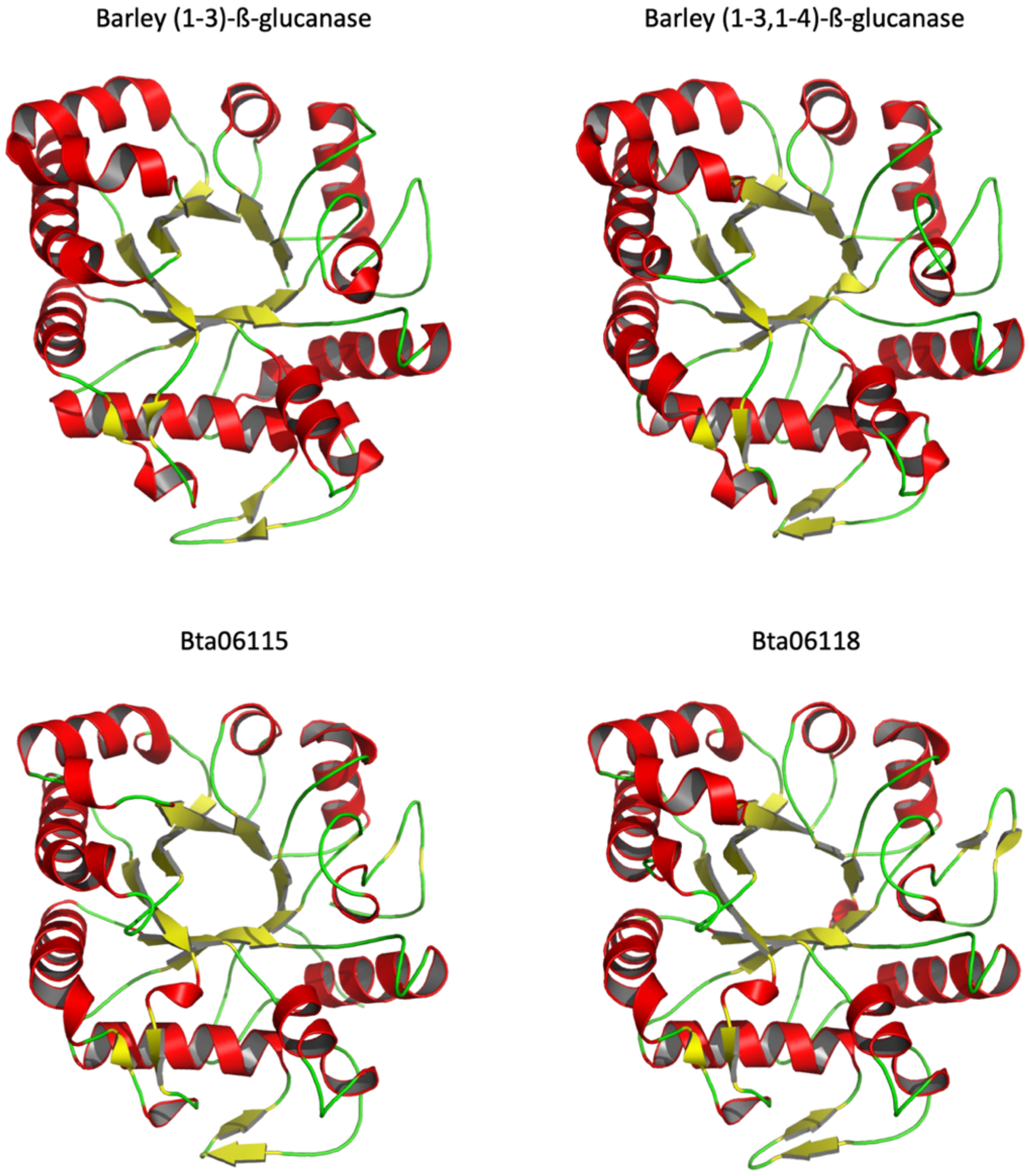

**Supplementary Figure 4.** Comparison of the predicted structures of the pectinesterase active part of Bta11221 and Bta11222 proteins with the known structure of carrot PME. (A) Alignment of Bta11221 (blue) and carrot PME (orange) structures. (B) Alignment of Bta11222 (green) and carrot PME (orange) structures. (C) Comparison of the active site for carrot PME, Bta11221 and Bta11222. The five catalytically important residues for carrot PME (Gln113, Gln135, Asp136, Asp157 and Arg225) and the corresponding residues in Bta11221 and Bta11222 are shown as balls and sticks.

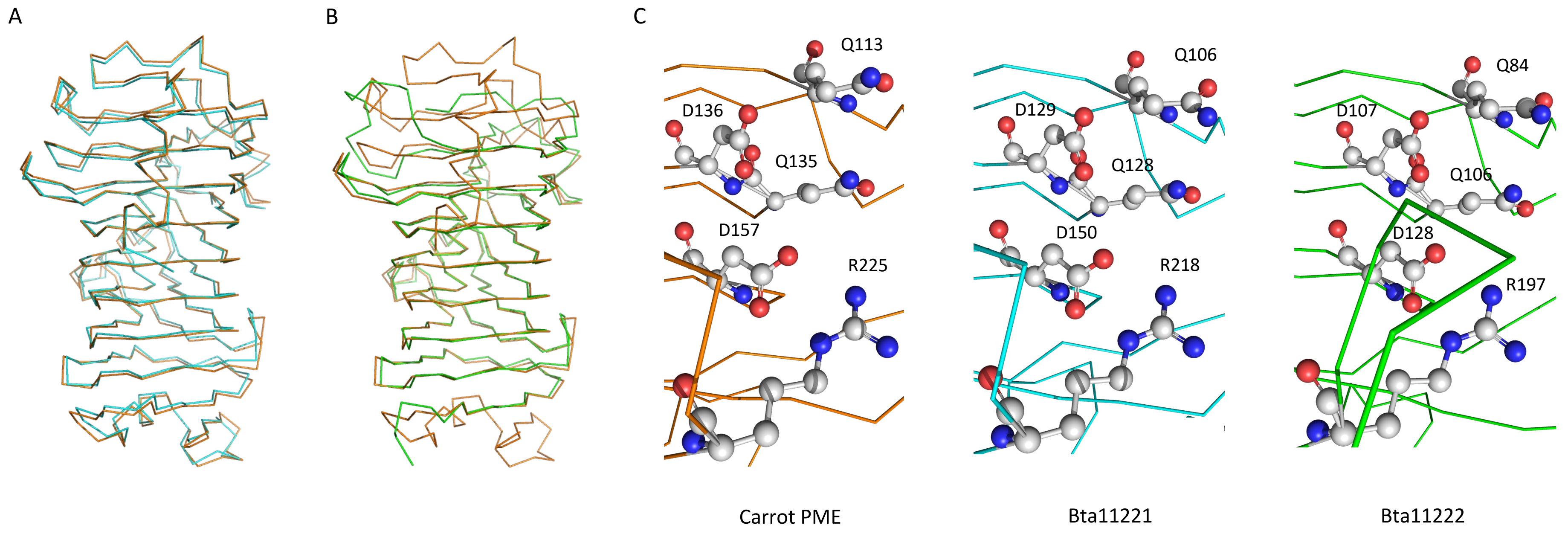

**Supplementary Figure 5.**
Phylogeny inferred for the 11 combined groups in which the sequences of *B. tabaci* and *T. vaporariorum* formed monophyletic clades with support values greater than or equal to 80% and 95% for SH-aLRT and UFboot, respectively.

**Donor: Bacteria**

**TvaB06_BtaB09**

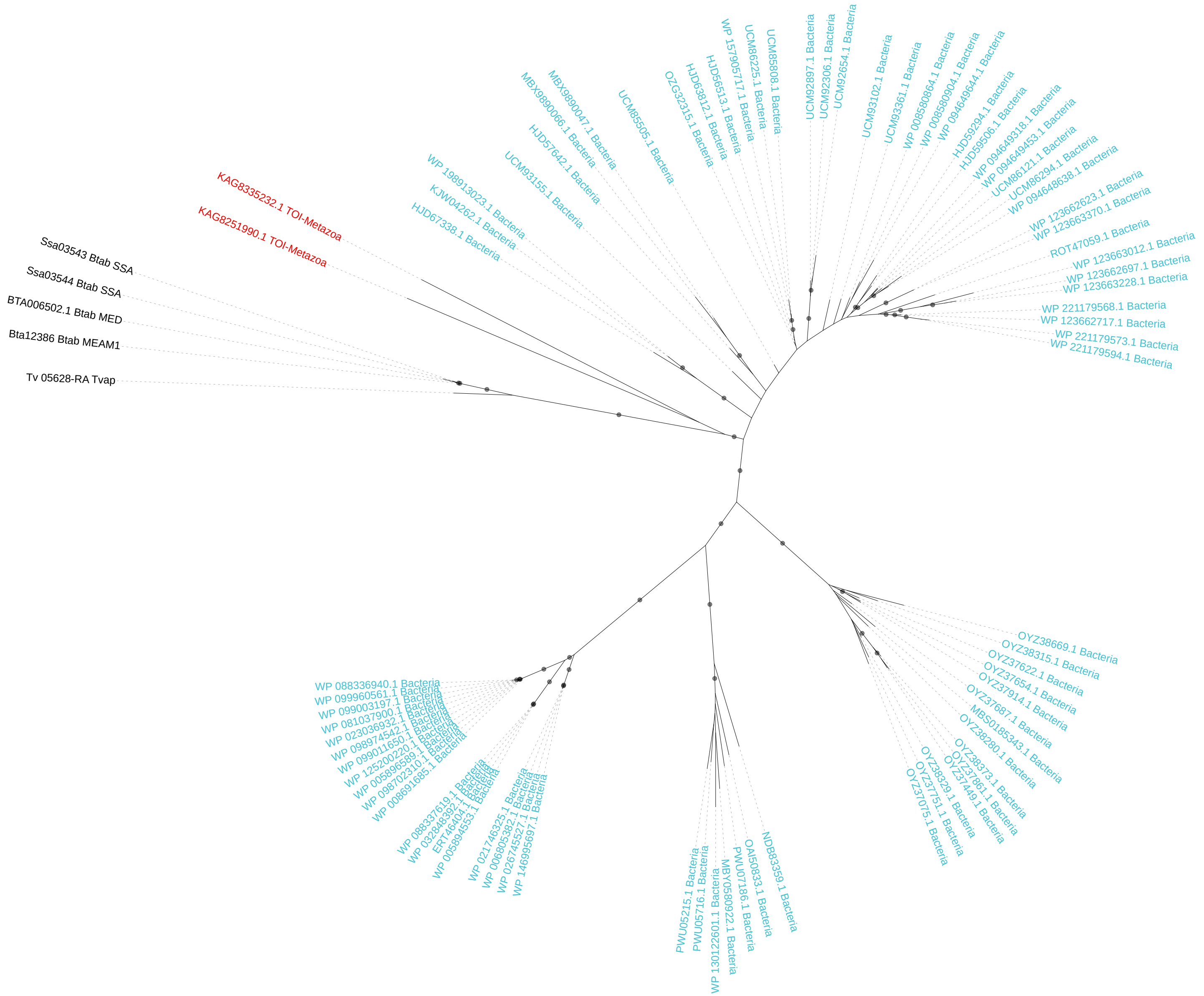

**Donor: Bacteria**

**TvaB07_BtaB36**

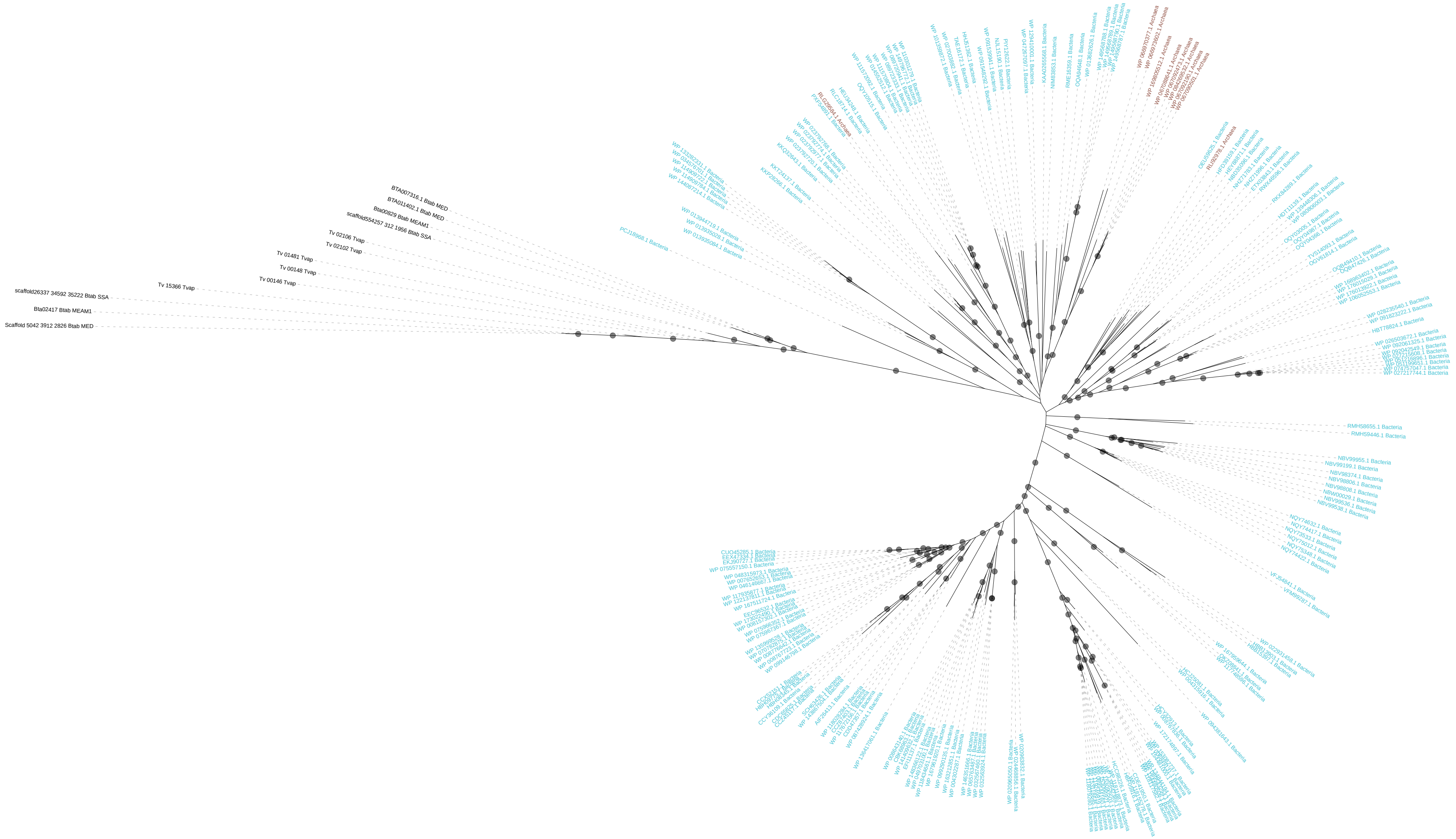

**Donor: Bacteria**
**TvaB16_BtaB45**

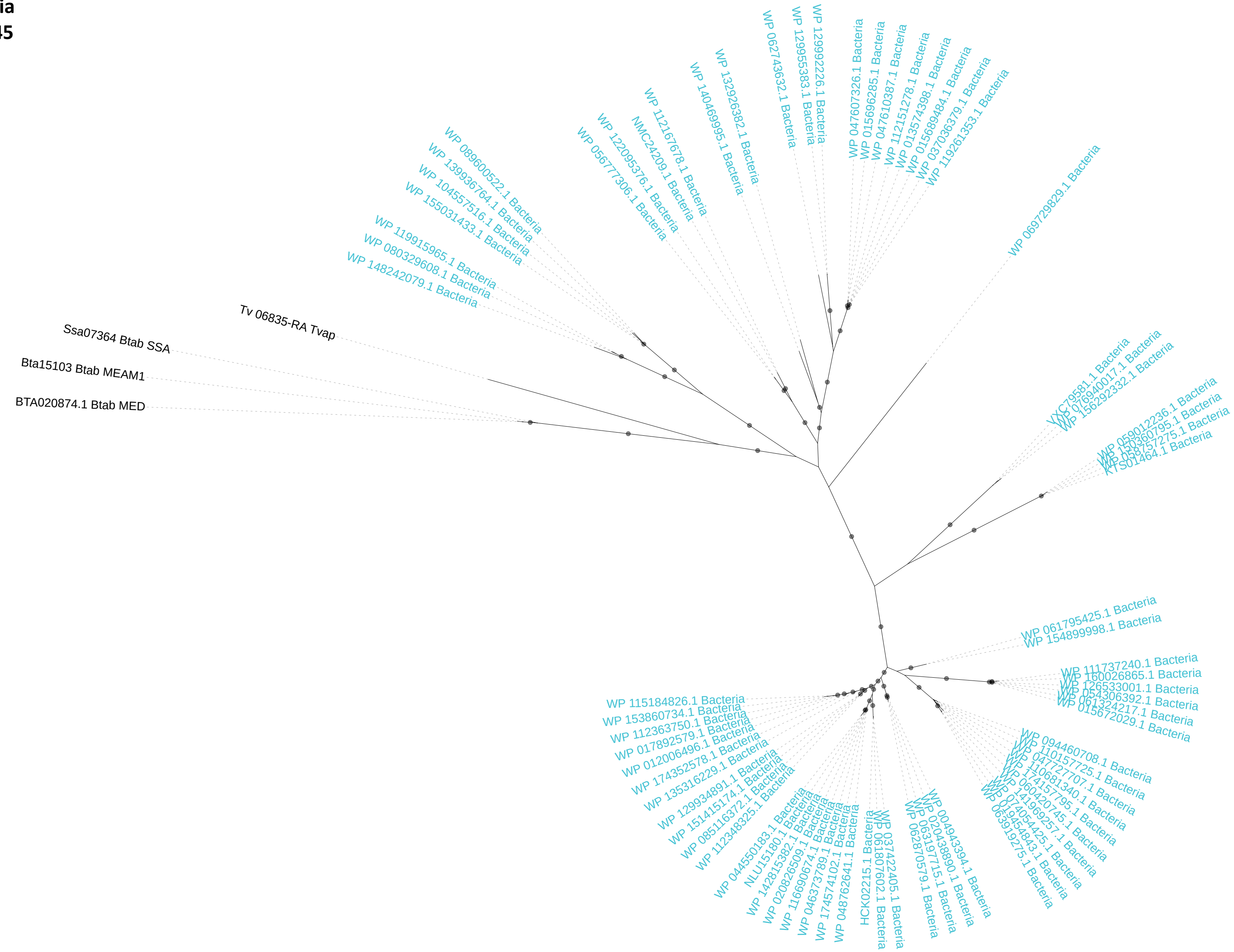

**Donor: Bacteria**
**TvaB17_BtaB65**

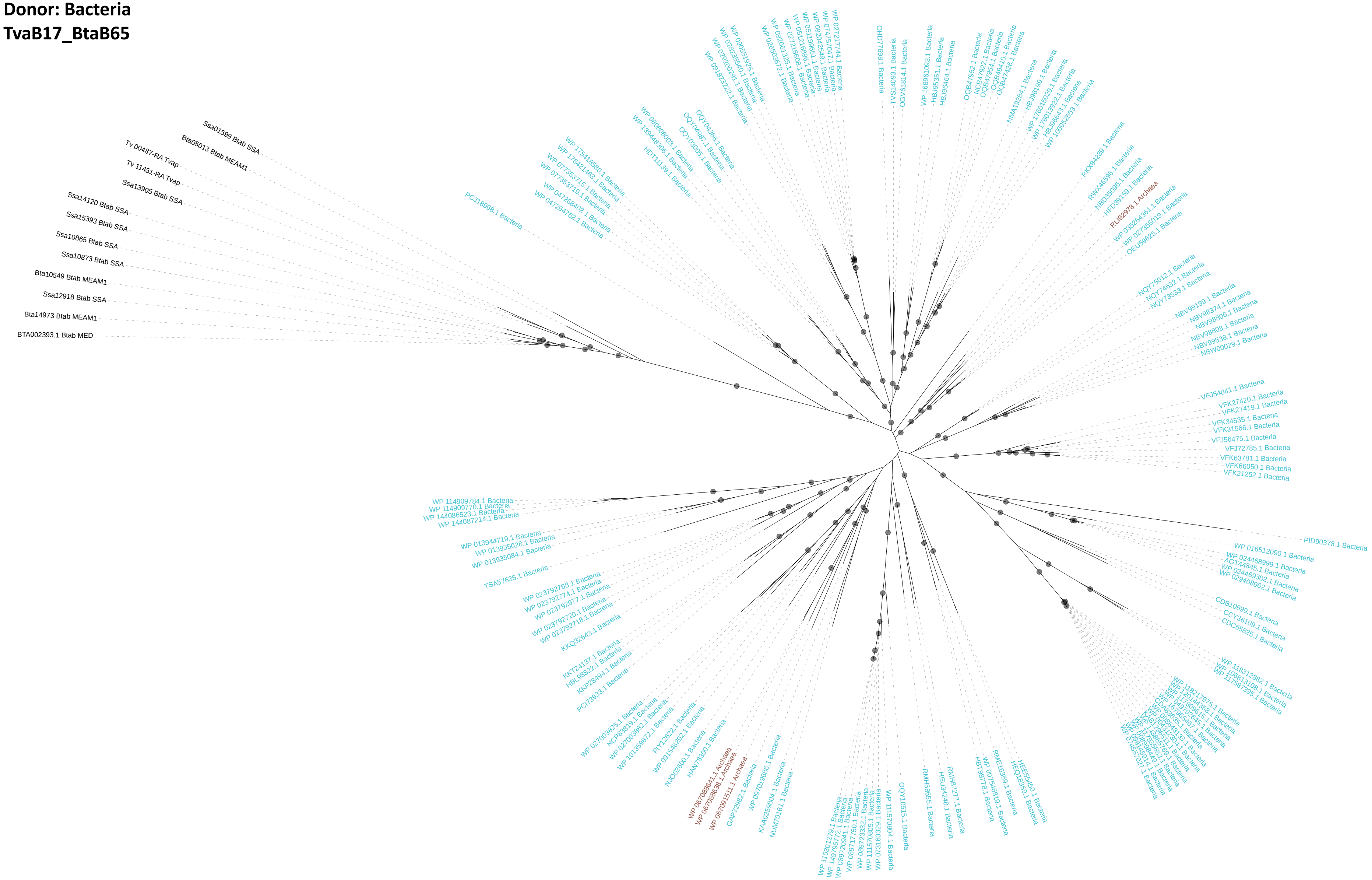

**Donor: Bacteria**
**TvaB19_BtaB01**

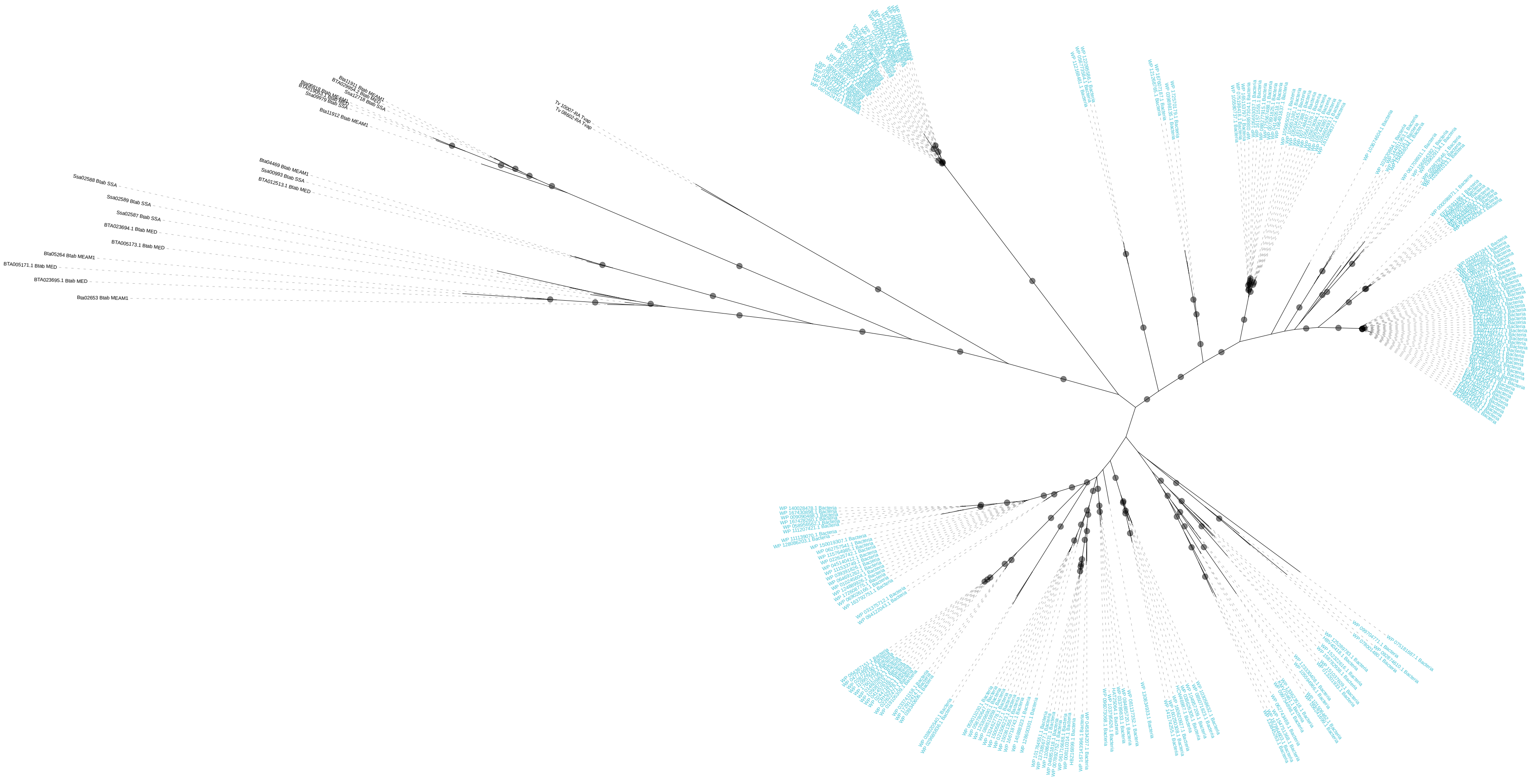

**Donor: Bacteria**
**TvaB20_BtaB11**

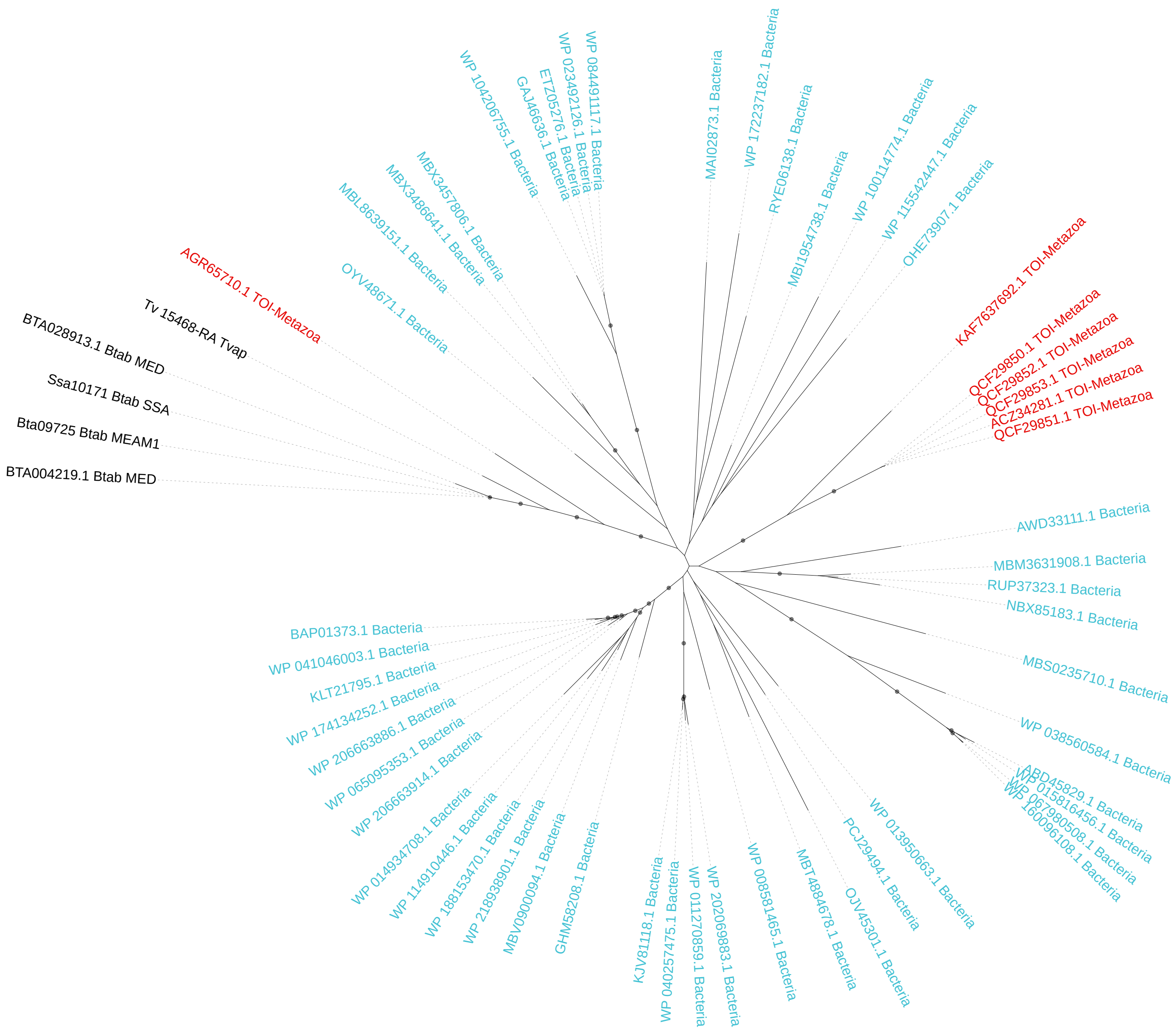

**Donor: Bacteria**
**TvaB22_BtaB35**

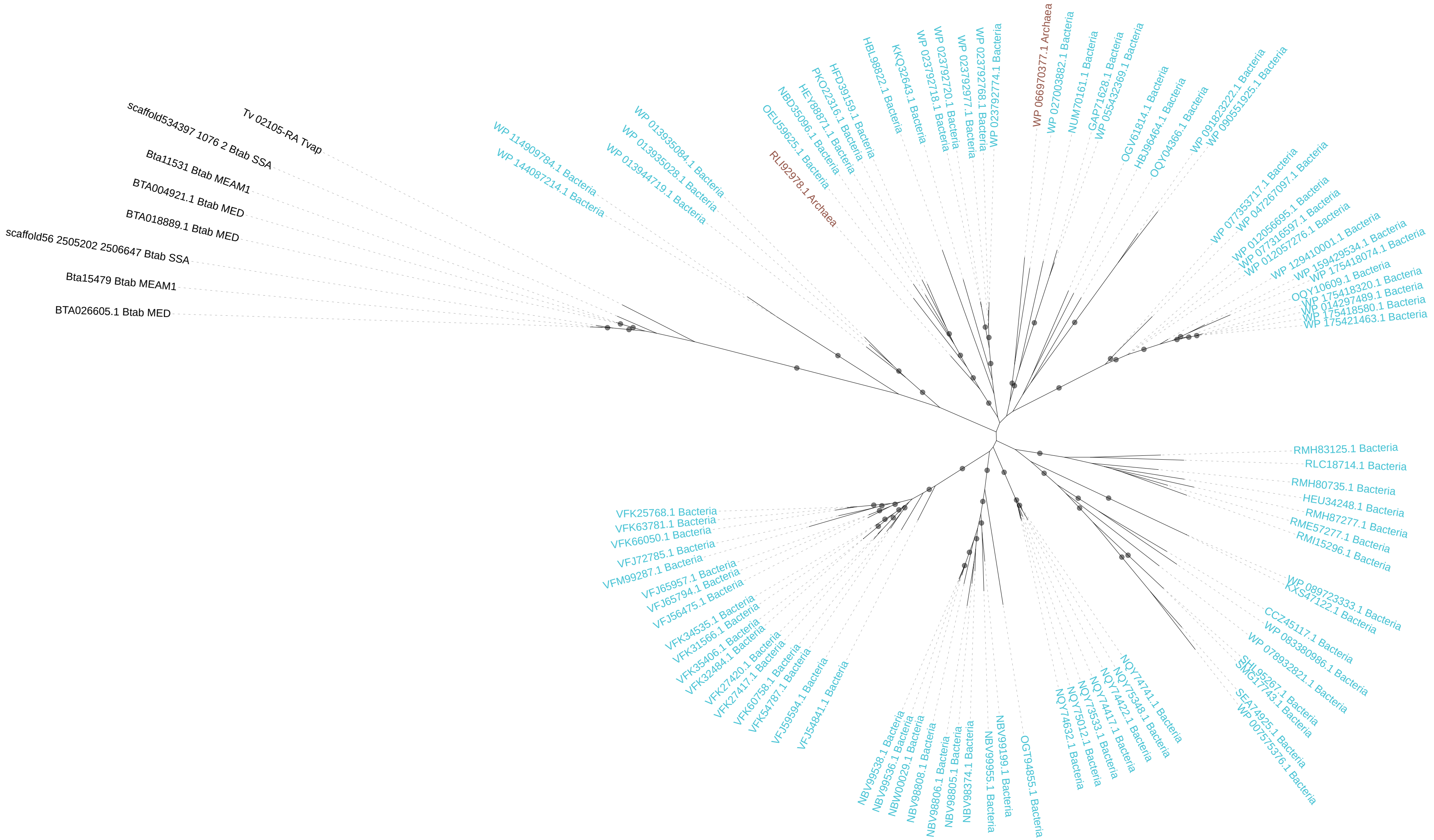

**Donor: Fungi**
**TvaF01_BtaF22**

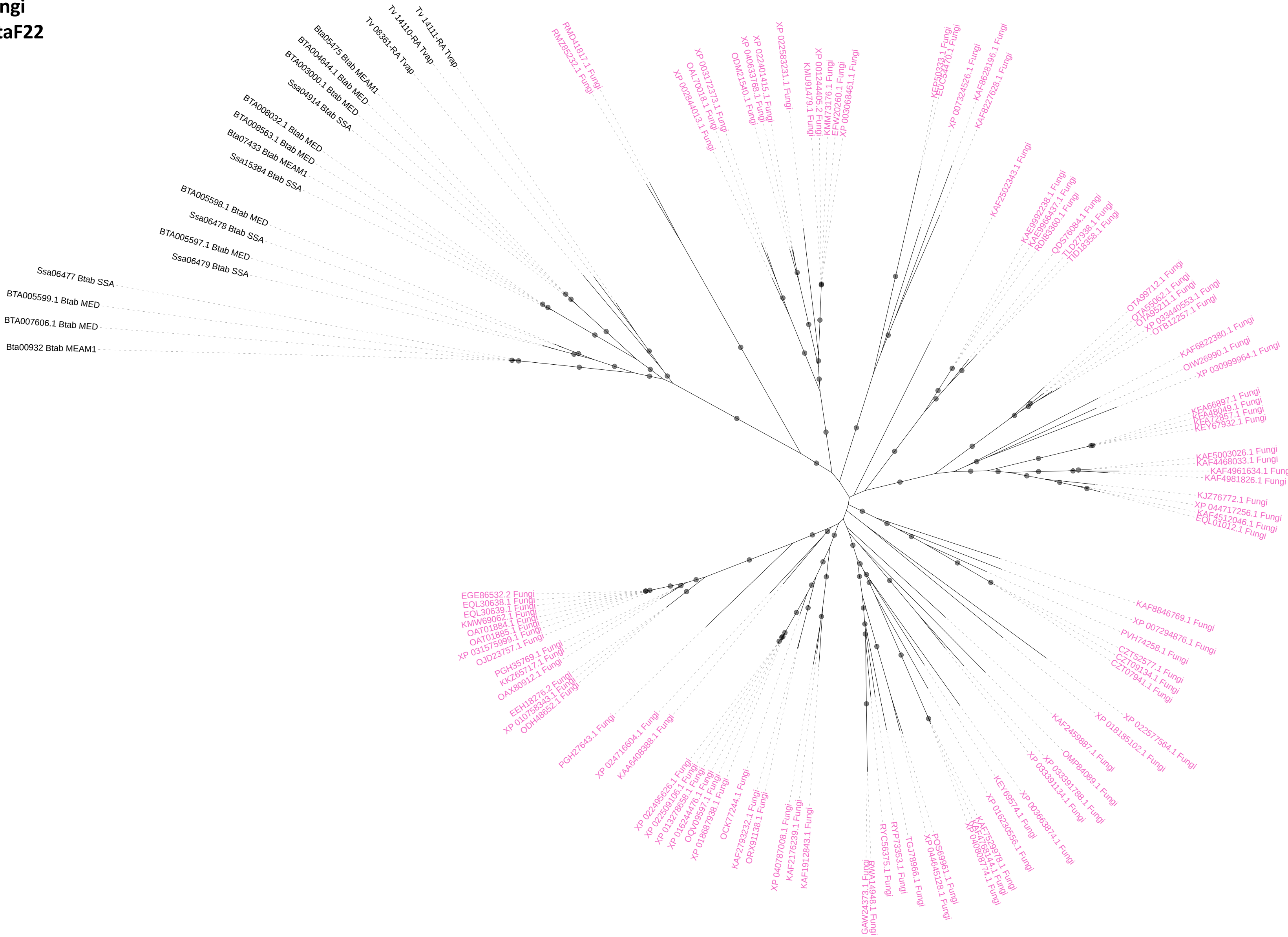

**Donor: Fungi**
**TvaF02_BtaF37**

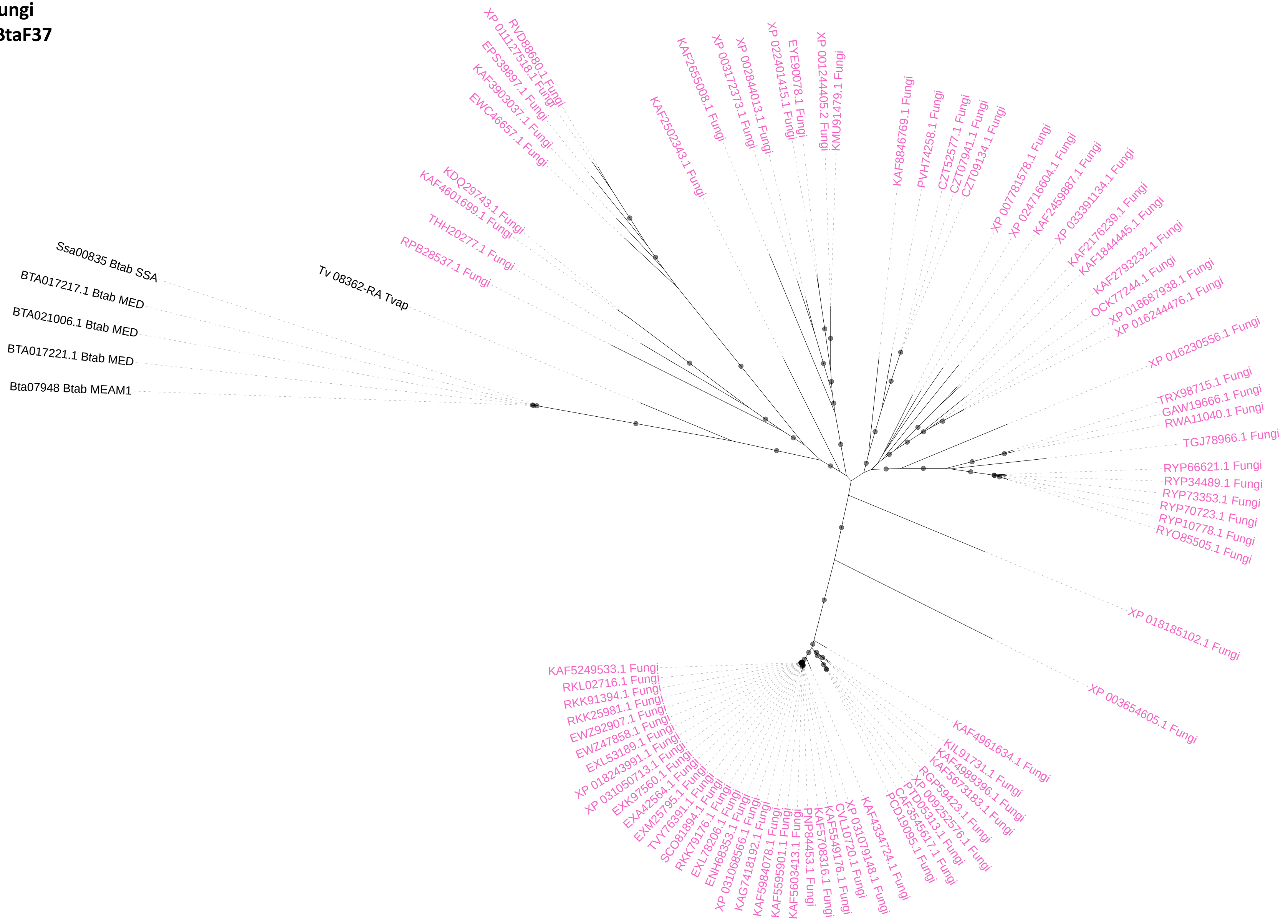

**Donor: Fungi**
**TvaF03_BtaF26**

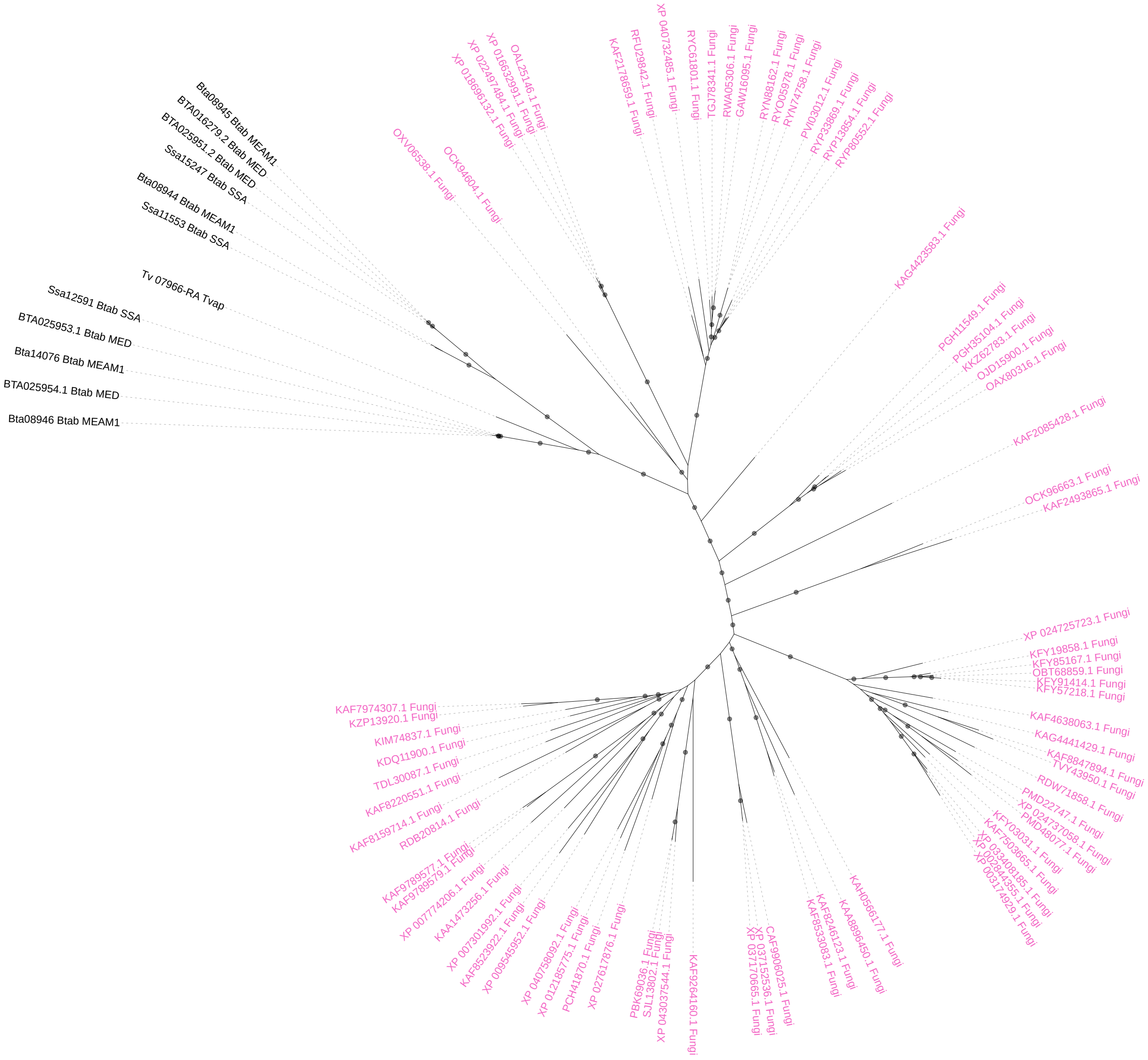

**Donor: Bacteria or Fungi**
**TvaC01_BtaC01**

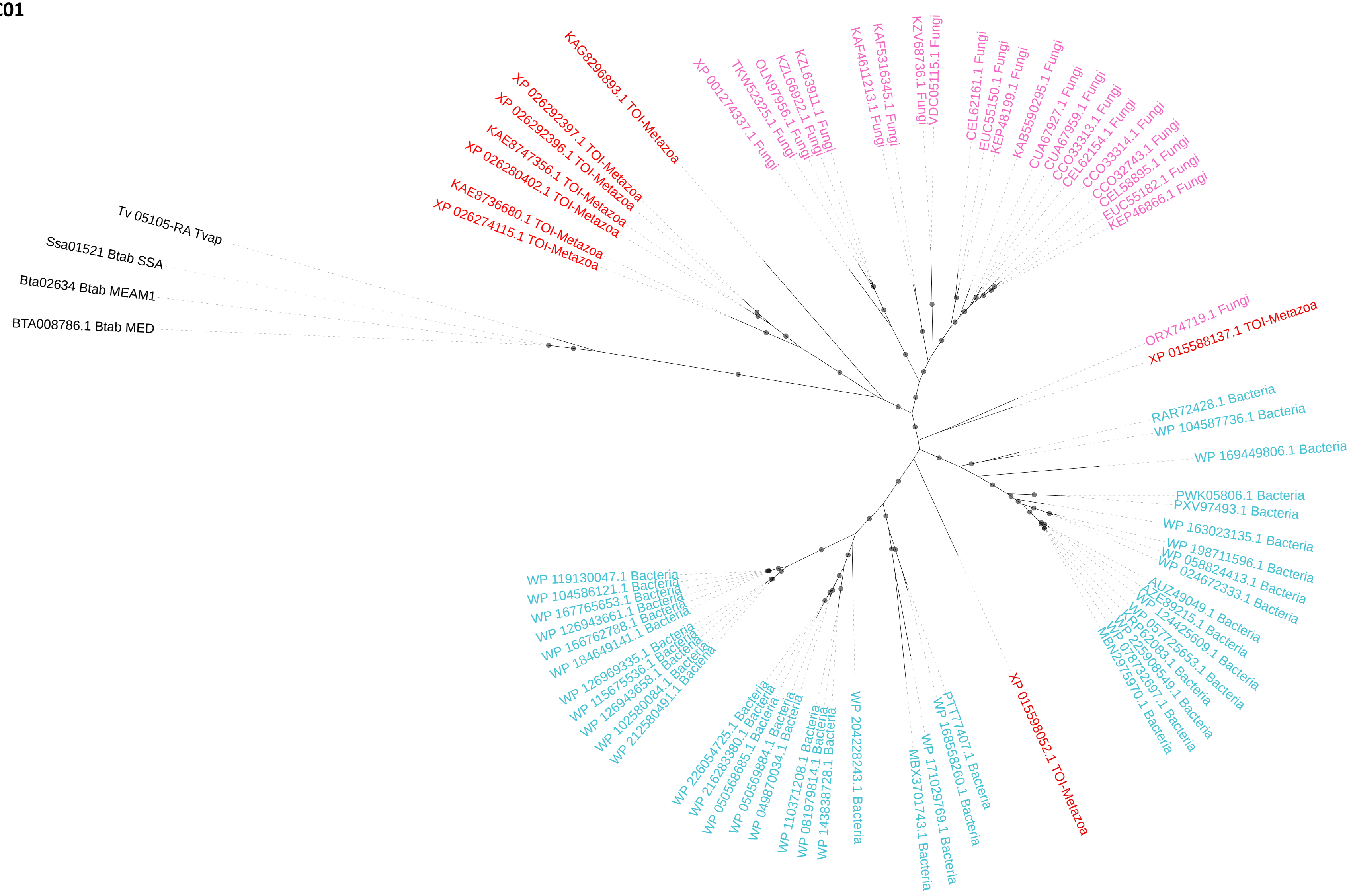

**Supplementary Figure 6.**
Phylogeny inferred for the 12 combined groups in which the grouping of sequences of *B. tabaci* and *T. vaporariorum* was not supported by bootstrap analysis and/or not monophyletic.

**Donor: Bacteria**

**TvaB01_BtaB56**

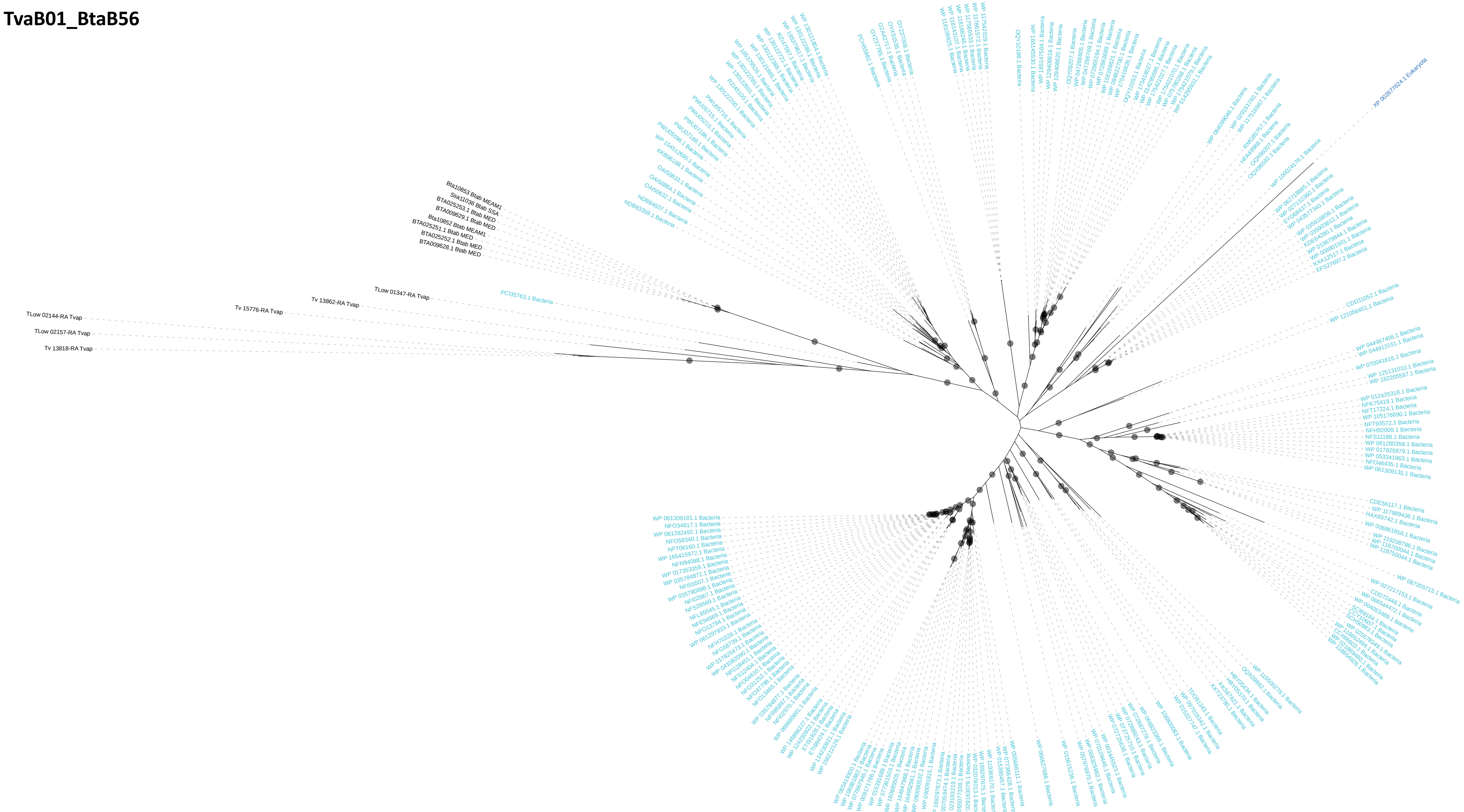

**Donor: Bacteria**
**TvaB02_BtaB12**

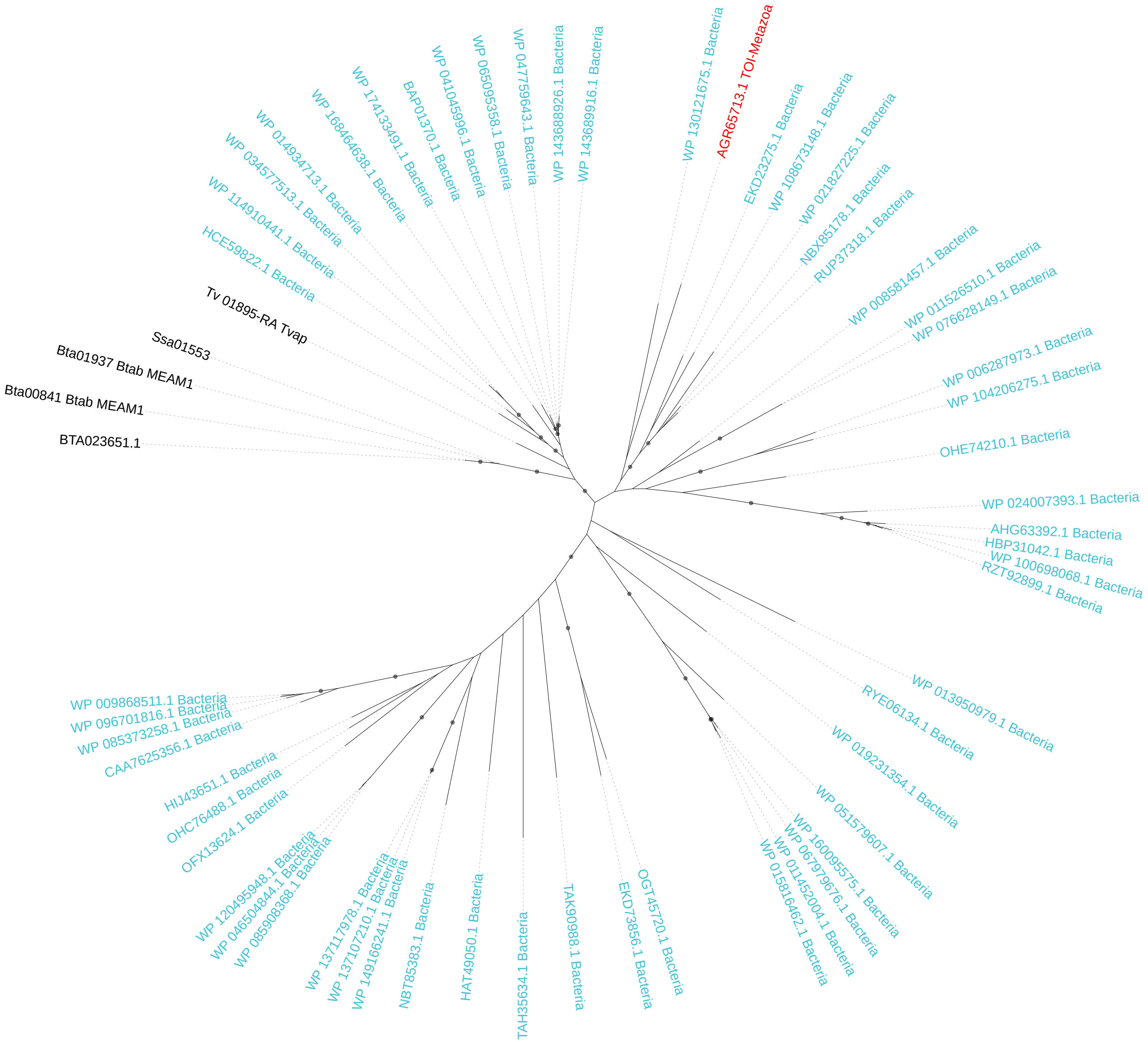

**Donor: Bacteria**
**TvaB03_BtaB68**

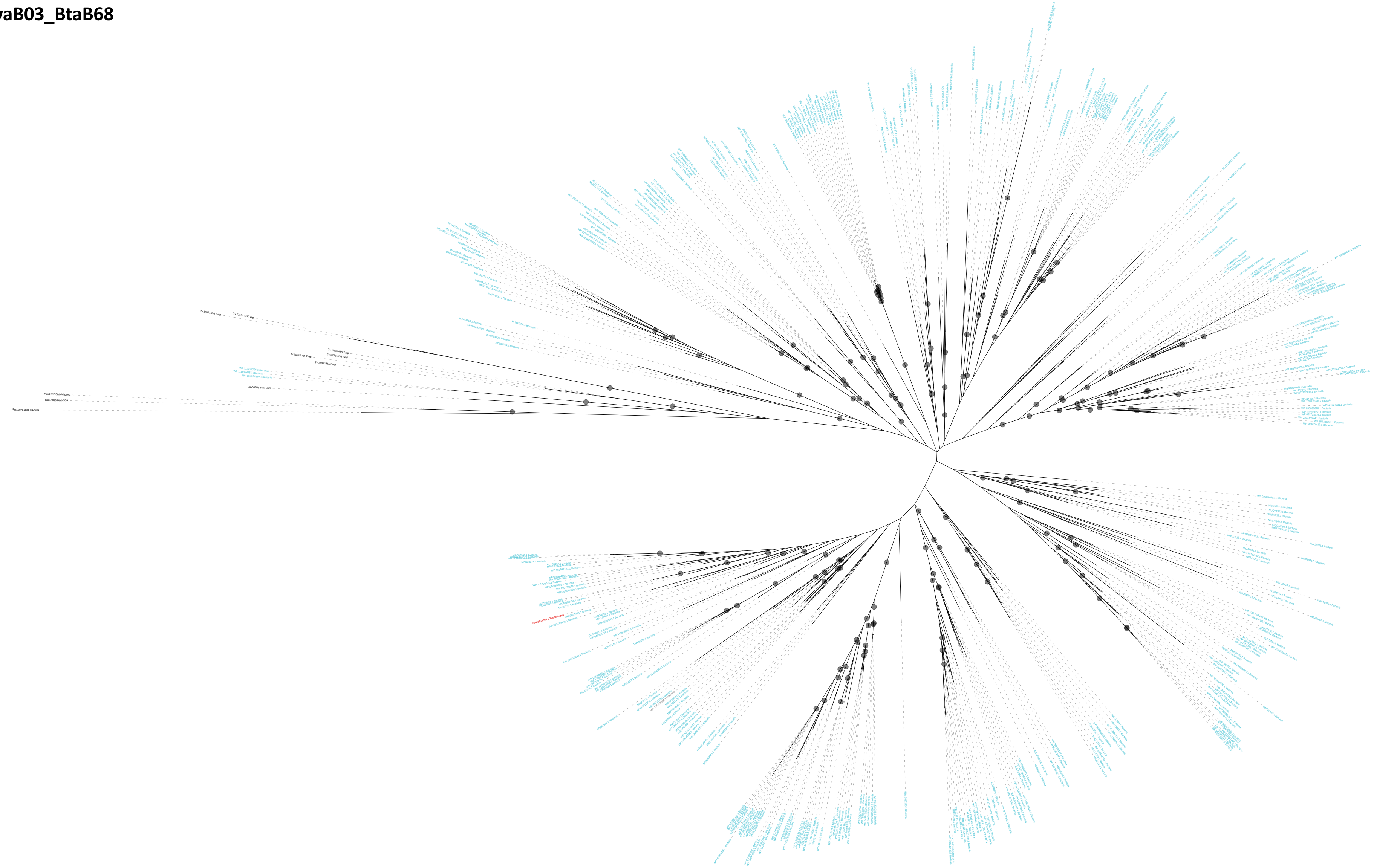

**Donor: Bacteria**

**TvaB04_BtaB44**

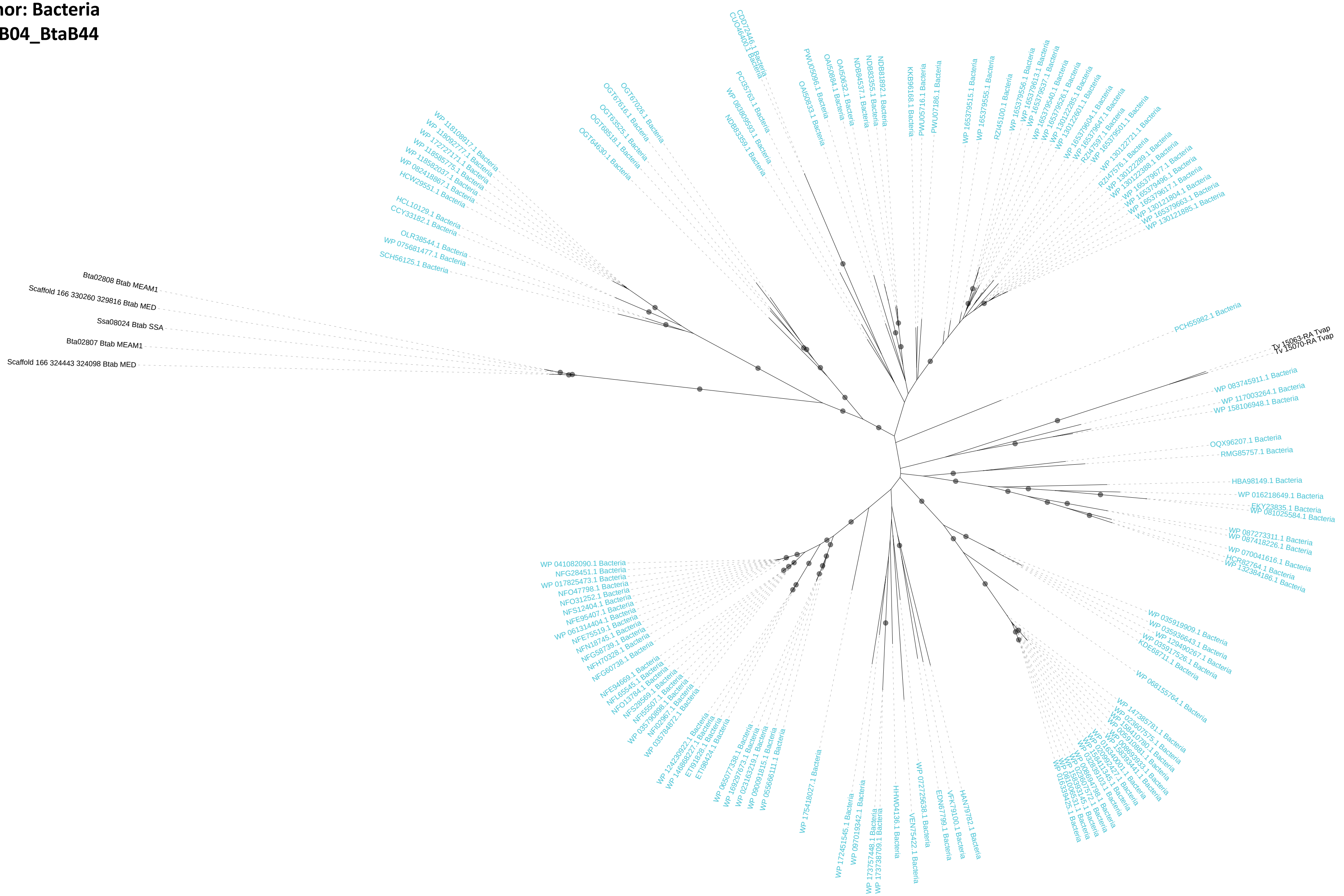

**Donor: Bacteria**
**TvaB05_BtaB02**

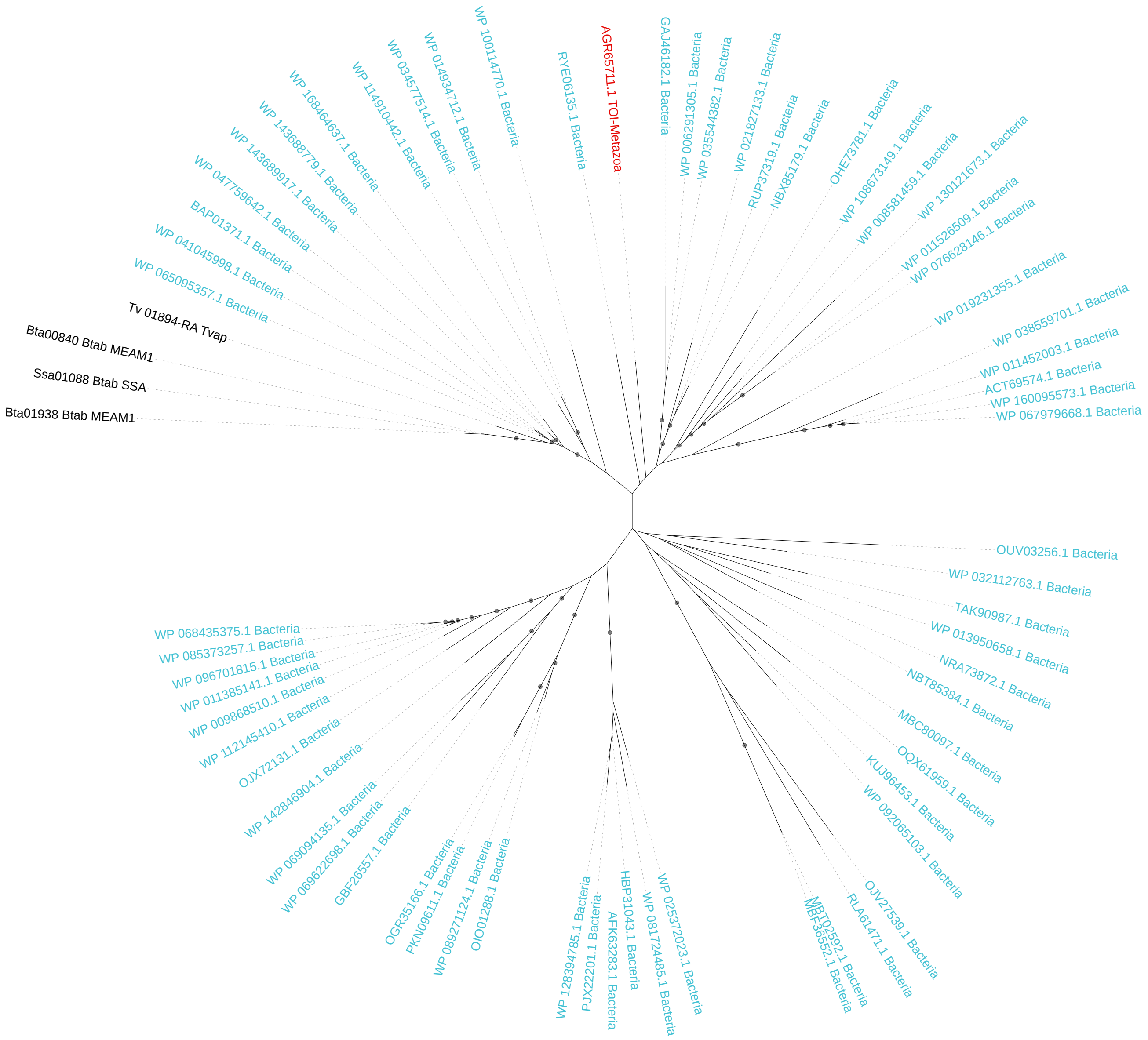

**Donor: Bacteria**

**TvaB08_BtaB31**

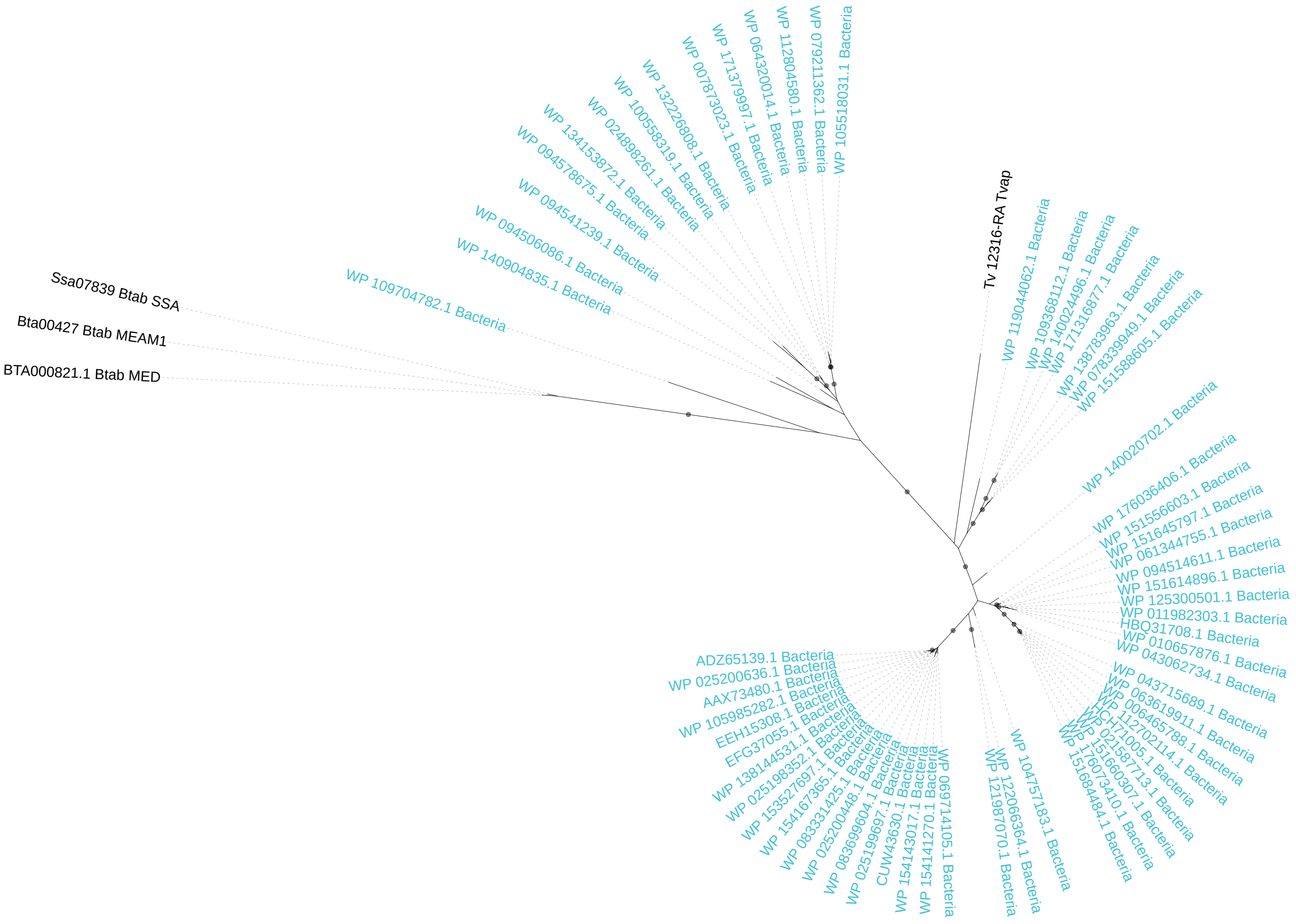

**Donor: Bacteria**
**TvaB11_BtaB03**

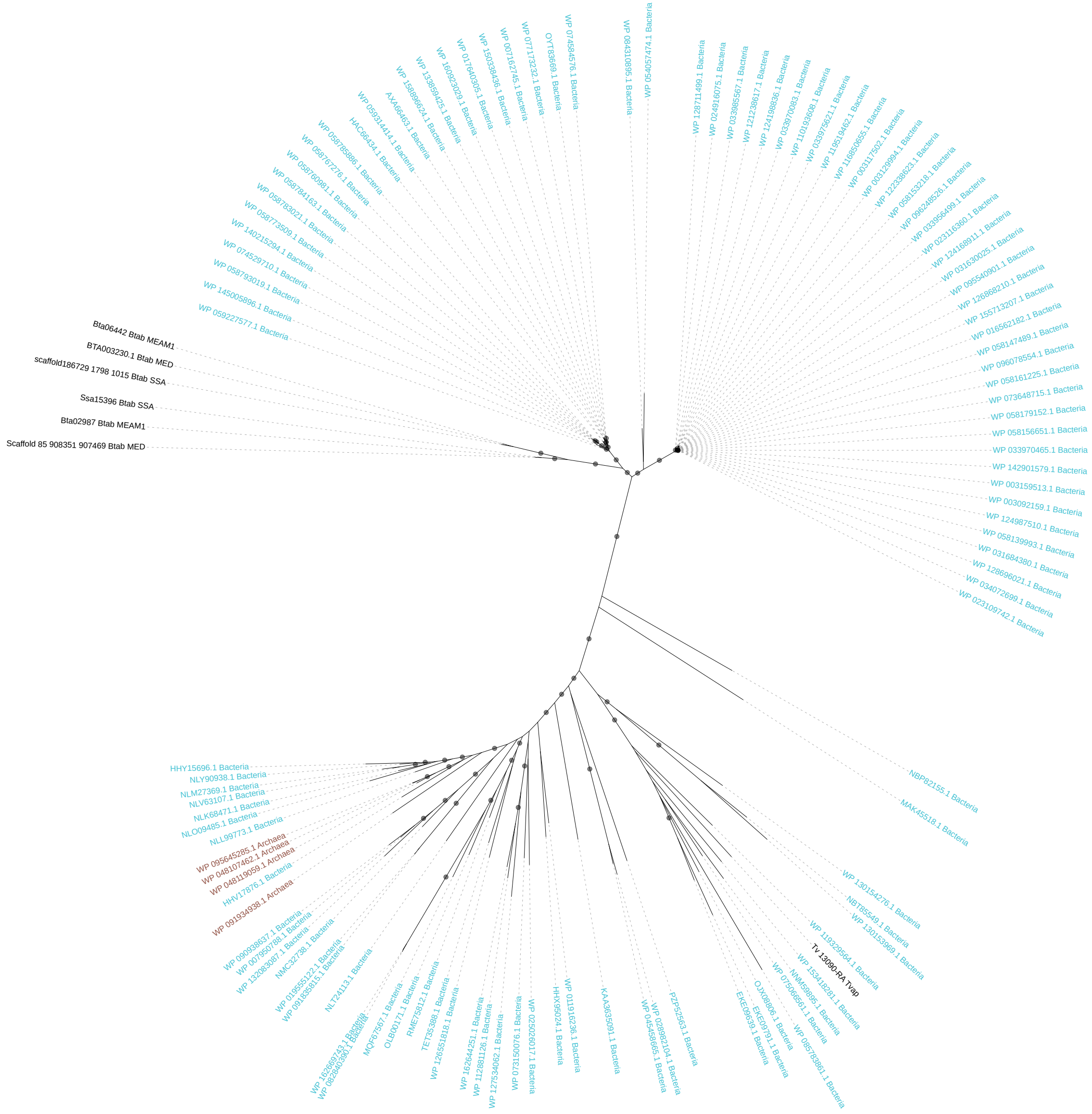

**Donor: Bacteria**
**TvaB15_BtaB33**

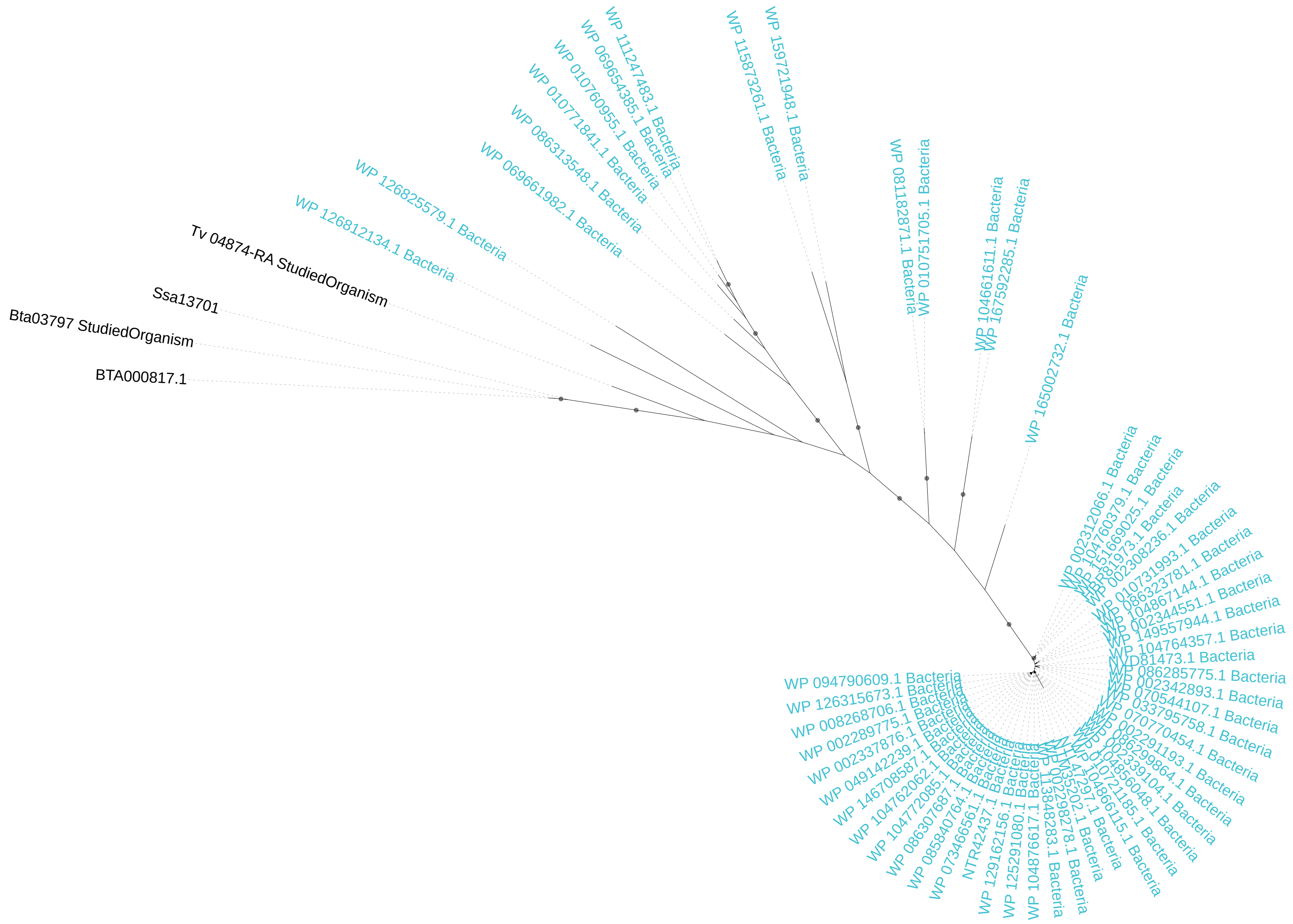

**Donor: Fungi**
**TvaF04_BtaF10**

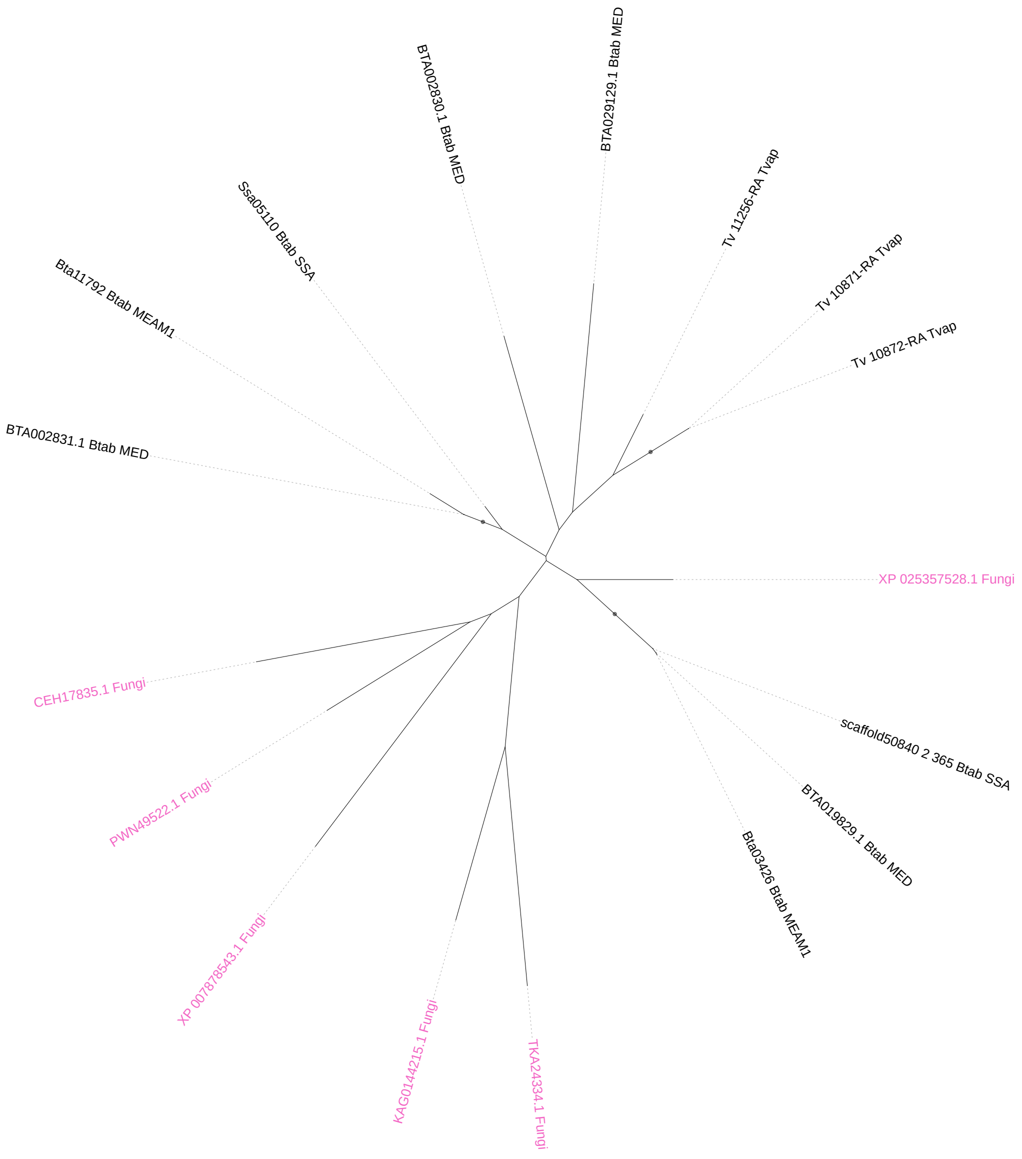

**Donor: Viridiplantae**
**TvaV01_BtaV01**

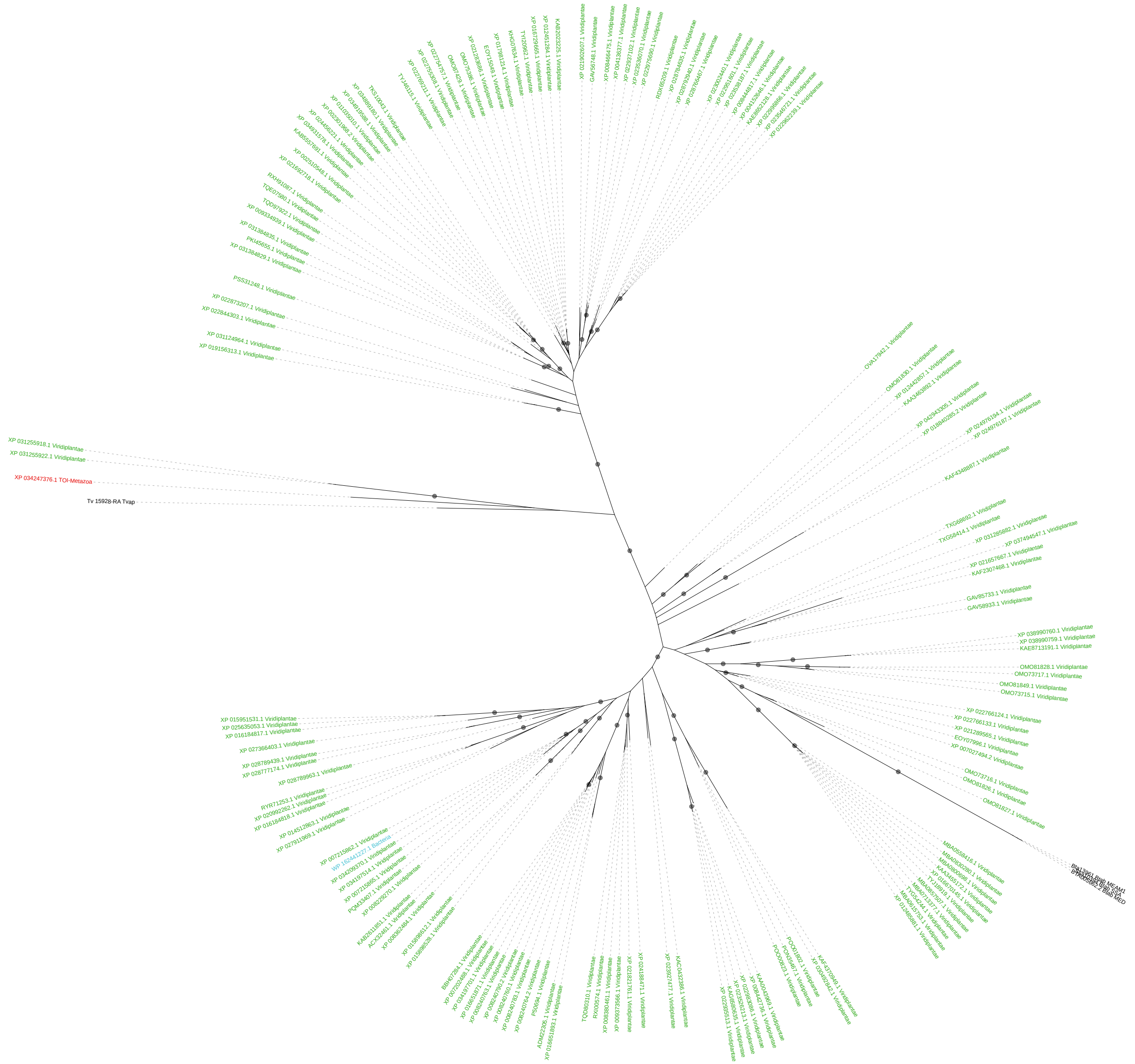

**Donor: Viridiplantae**

**TvaV02_BtaV23**

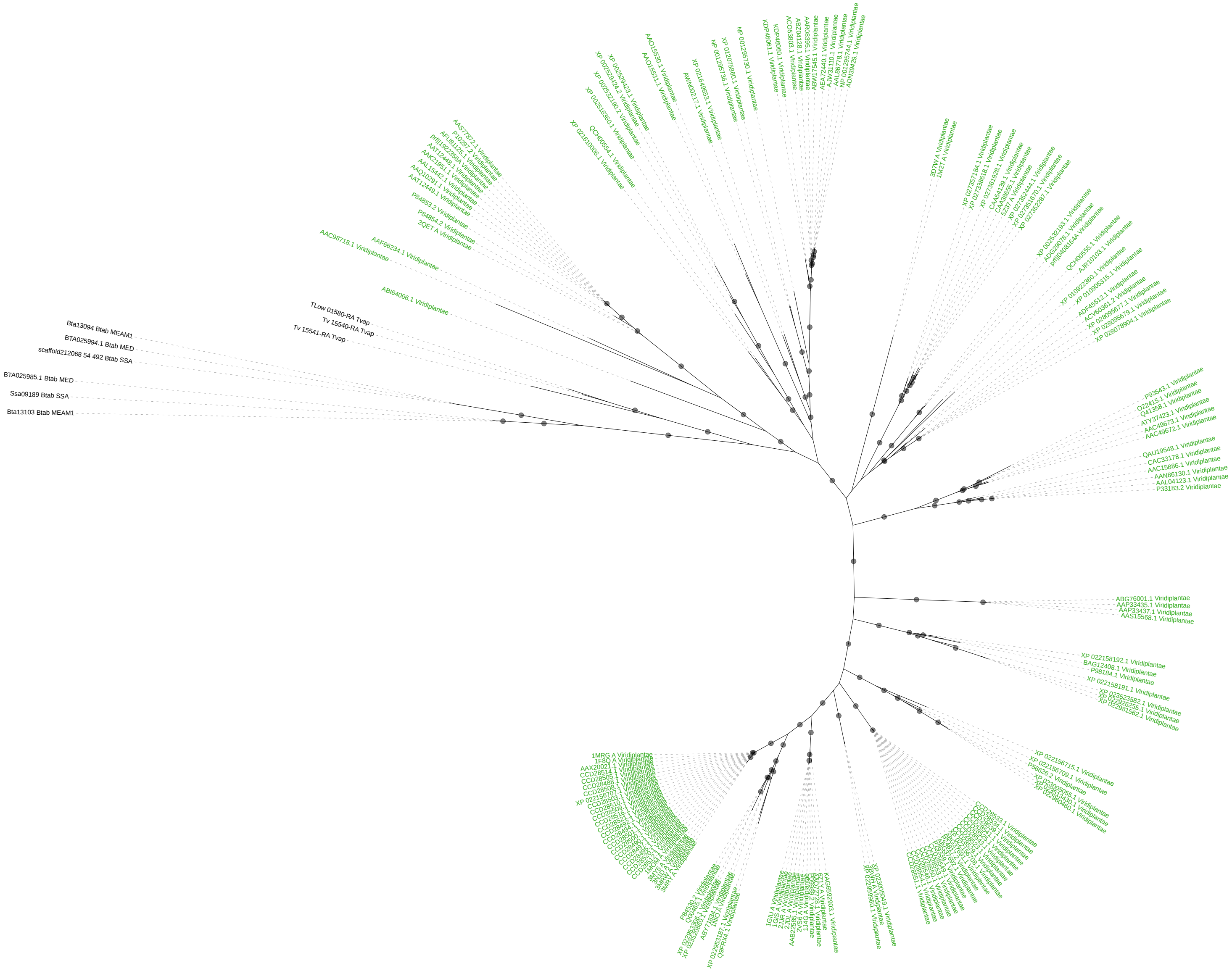

**Donor: Viridiplantae**
**TvaV03_BtaV21**

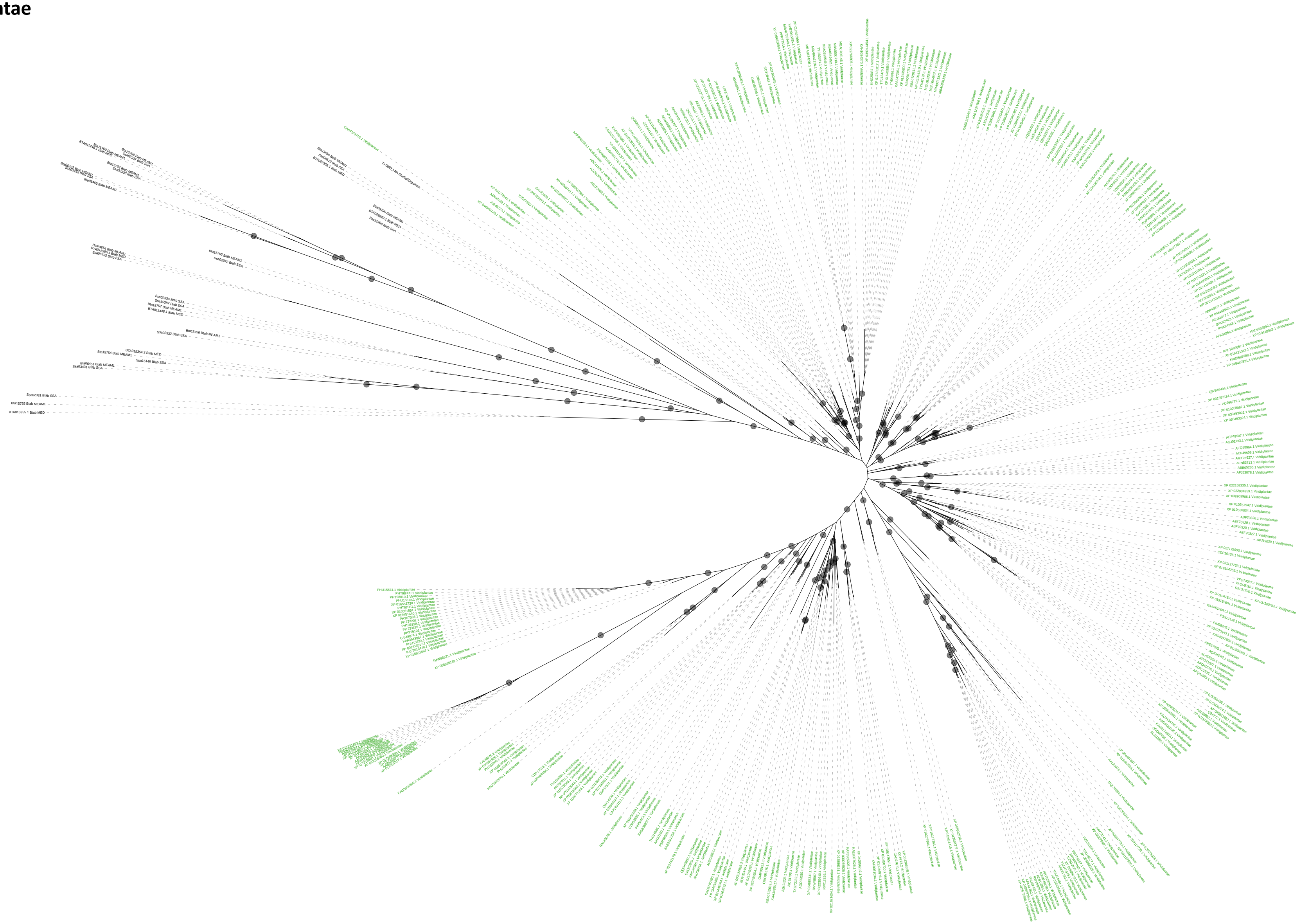

**Supplementary Figure 7.**

Phylogeny inferred for the HGT events corresponding to CAZymes in *F. occidentalis* and *T. palmi*. Black dots correspond to nodes with support values greater than or equal to 80% and 95% for SH-aLRT and UFboot, respectively.

**FocB01_TpaB01: GH32**

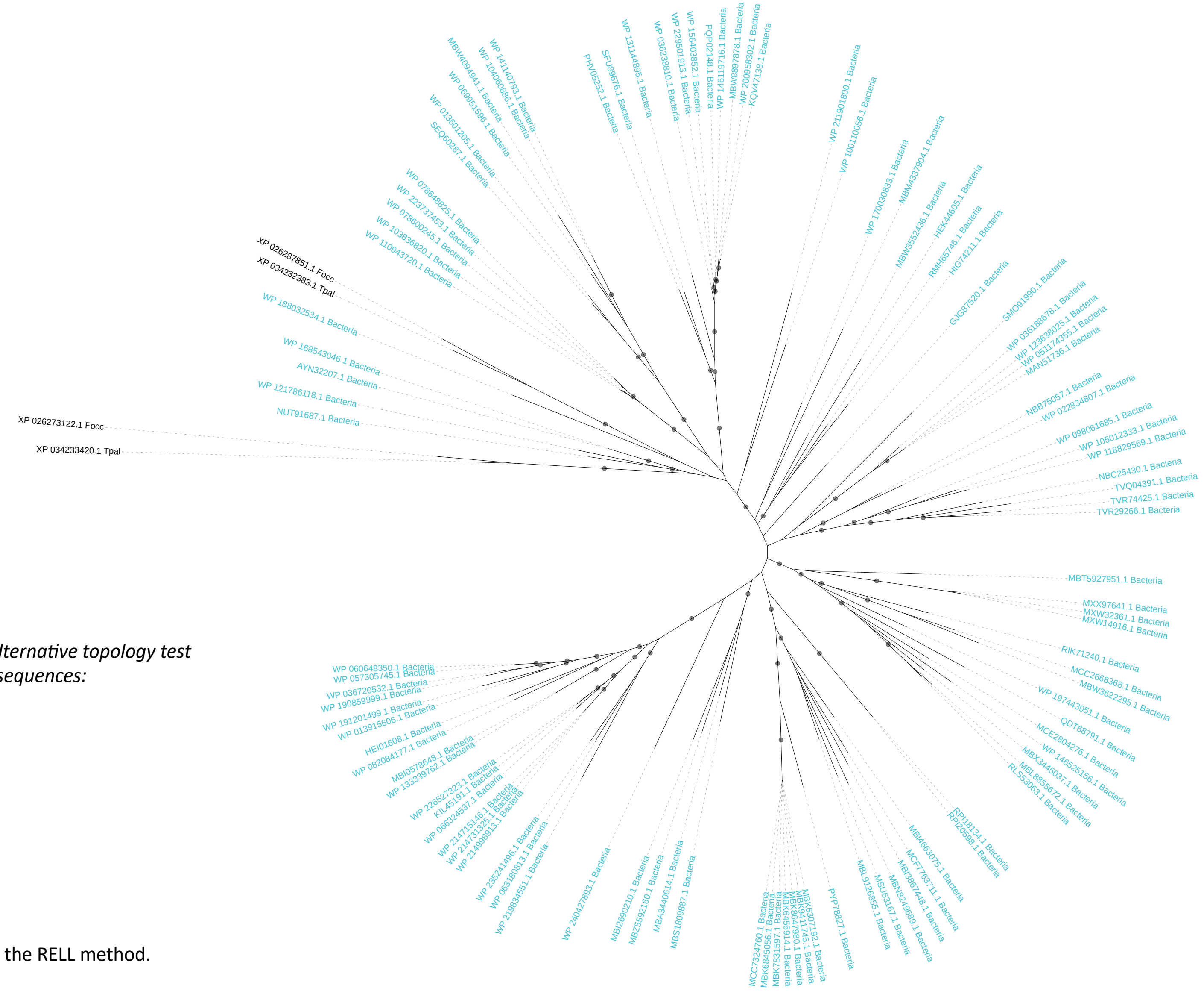

*Results of the approximately unbiased (AU) alternative topology test for monophyly of F. occidentalis and T. palmi sequences:*

| Tree | logL | deltaL | p-AU | |
|---|---|---|---|---|
| 1 | -57479.74173 | 4.2003 | 0.427 | + |
| 2 | -57475.54146 | 0 | 0.573 | + |

Plus signs denote the 95% confidence sets.

Minus signs denote significant exclusion.

All tests performed 10,000 resamplings using the RELL method.

#### FocB01_TpaB01_BtaB53: GH32

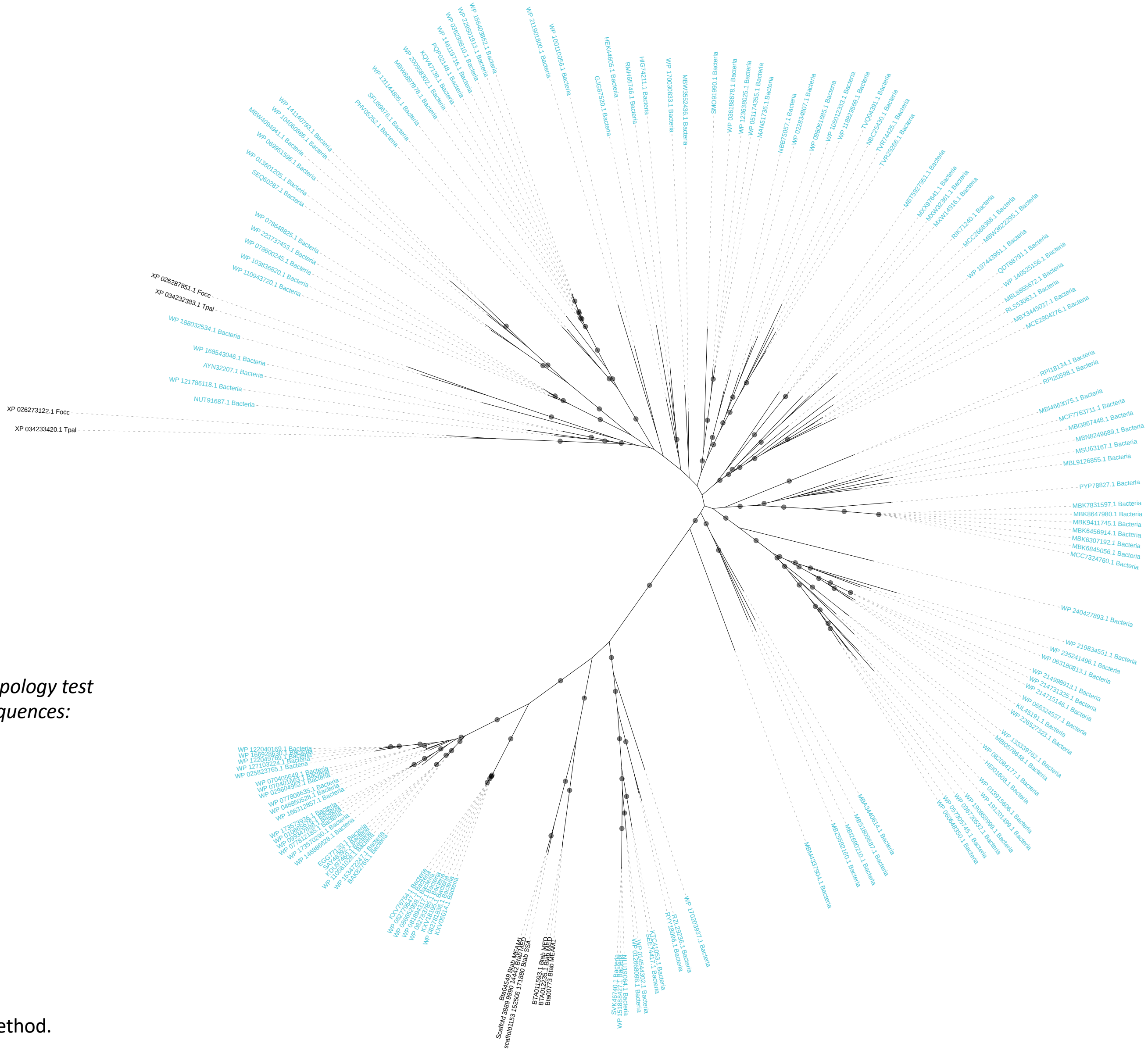

*Results of the approximately unbiased (AU) alternative topology test for monophyly of F. occidentalis, T. palmi and B. tabaci sequences:*

| Tree | logL | deltaL | p-AU | |
|---|---|---|---|---|
| 1 | -44835.73376 | 178.92 | 2.84e-07 | - |
| 2 | -44656.81805 | 0 | 1 | + |

Plus signs denote the 95% confidence sets.

Minus signs denote significant exclusion.

All tests performed 10,000 resamplings using the RELL method.

### FocB02_TpaB02: GH5_8

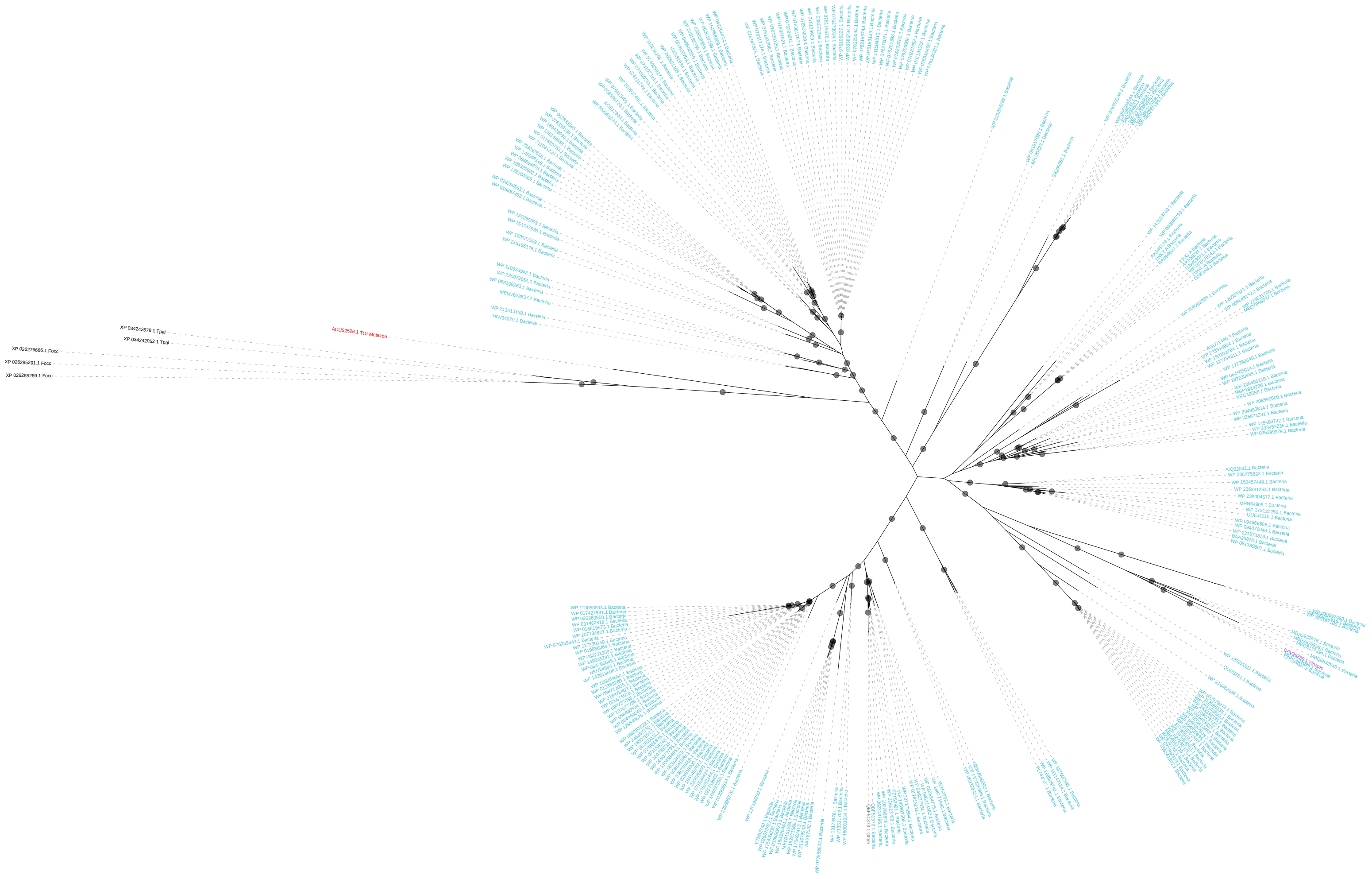

#### FocF01_TpaF01: GH45

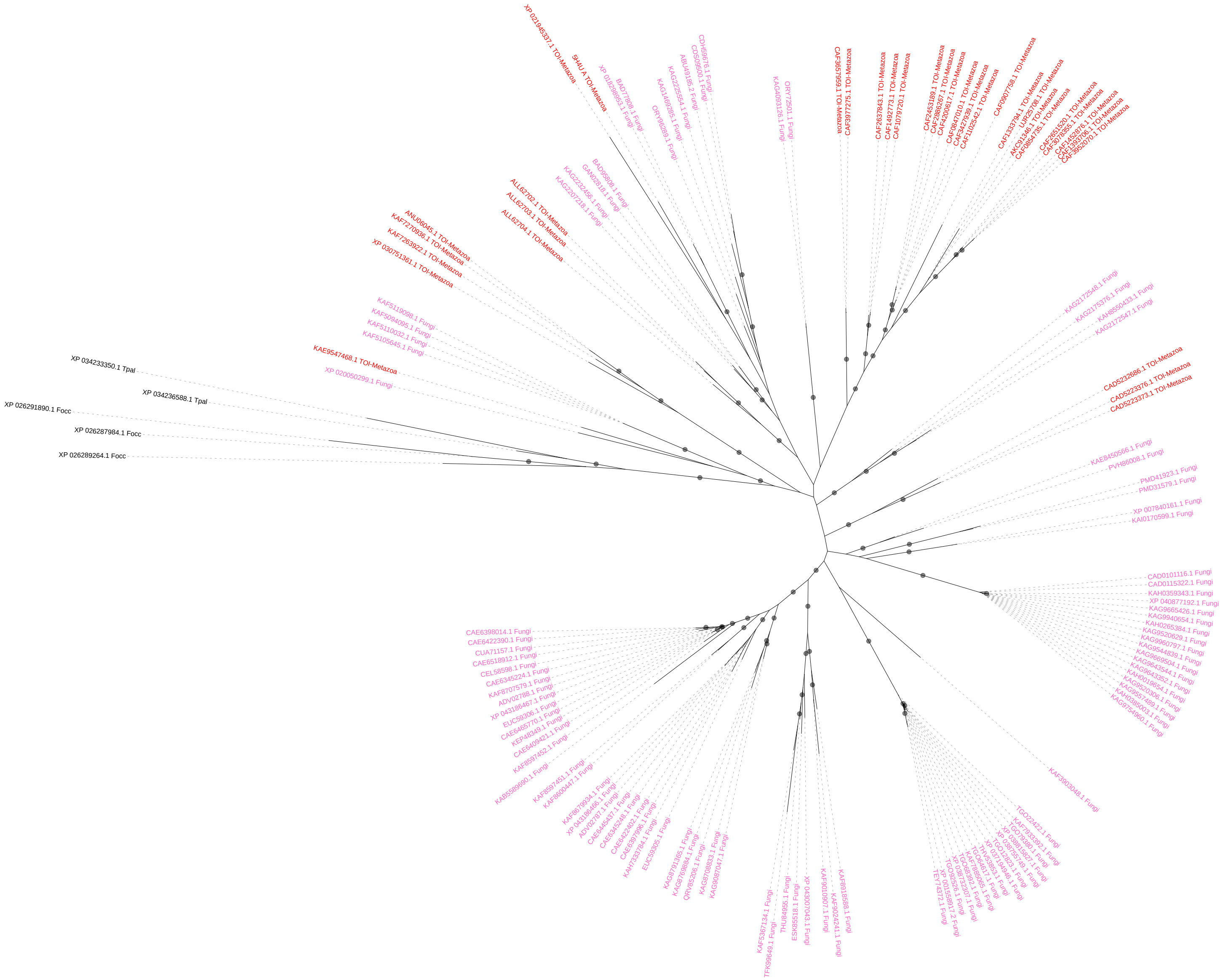

**FocV01_TpaV01: GH152**

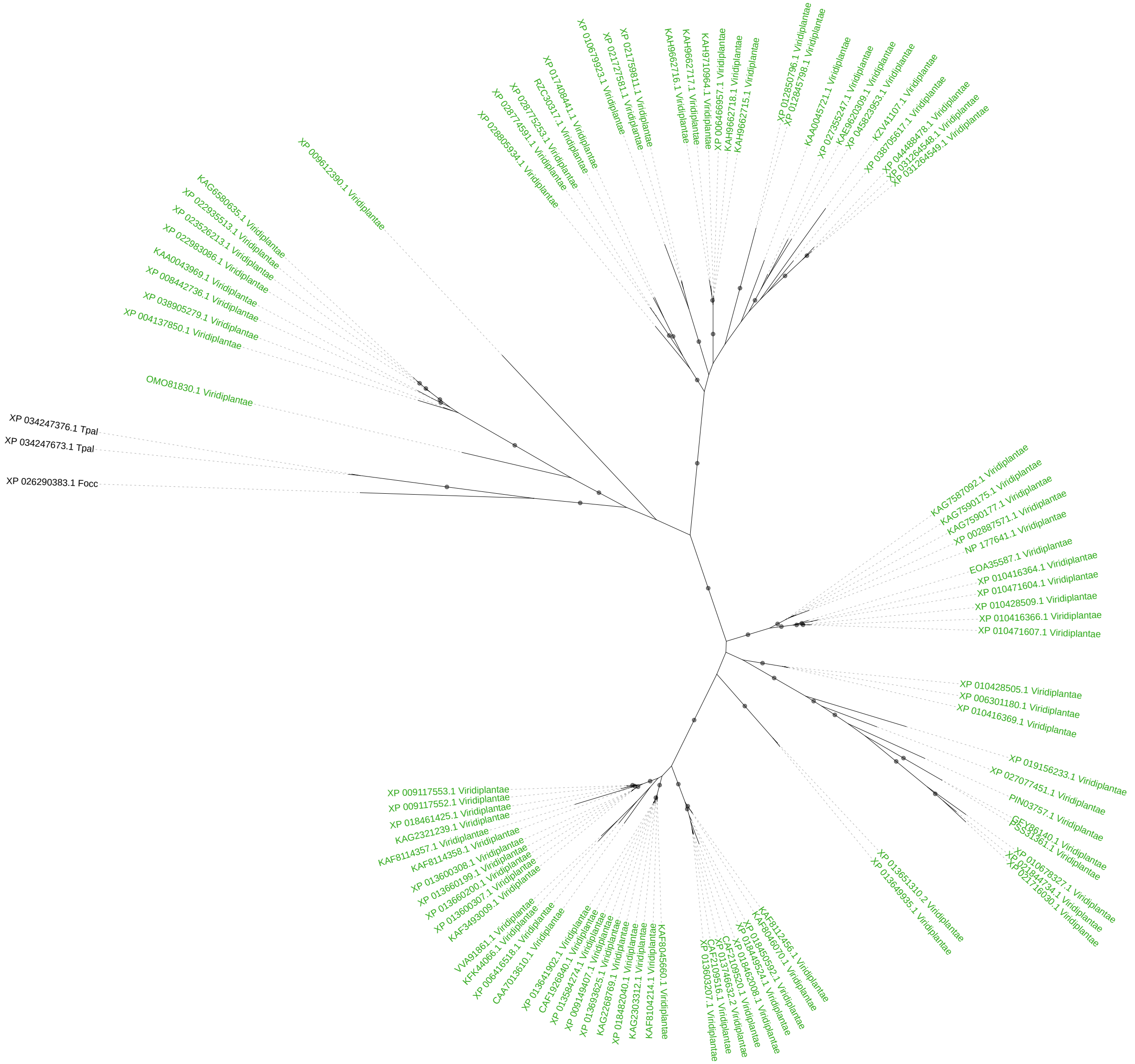

**FocV01_TpaV01_BtaV01_TvaV01: GH152**

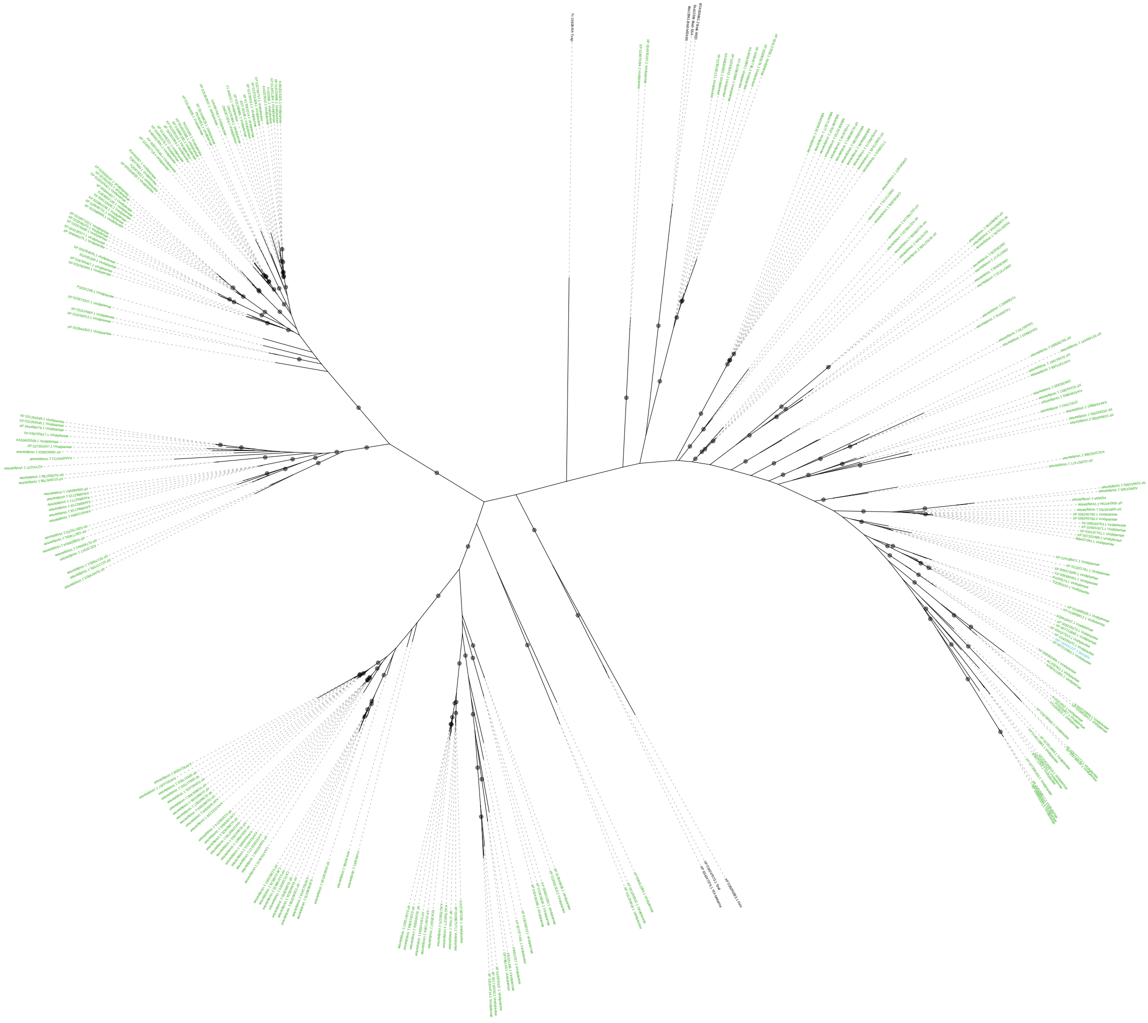

*Results of the approximately unbiased (AU) alternative topology test for monophyly of Thripinae and Aleyrodinae sequences:*

| Tree | logL | deltaL | p-AU | |
|---|---|---|---|---|
| 1 | -22115.50057 | 26.954 | 0.206 | + |
| 2 | -22088.54666 | 0 | 0.794 | + |

Plus signs denote the 95% confidence sets.

Minus signs denote significant exclusion.

All tests performed 10,000 resamplings using the RELL method.

### FocC01_TpaC01: PL1_4

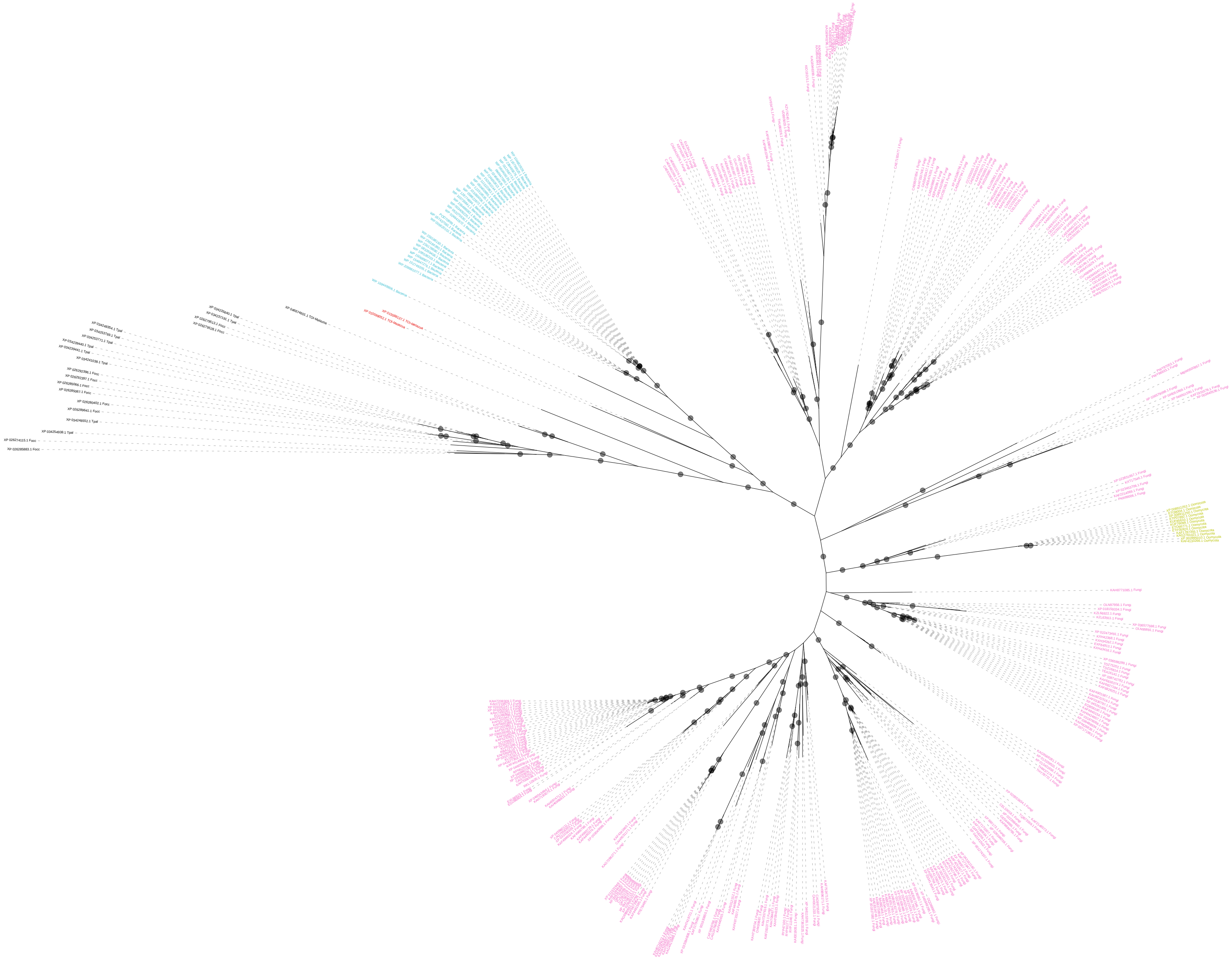

**FocC01_TpaC01_BtaC01_TvaC01: PL1_4**

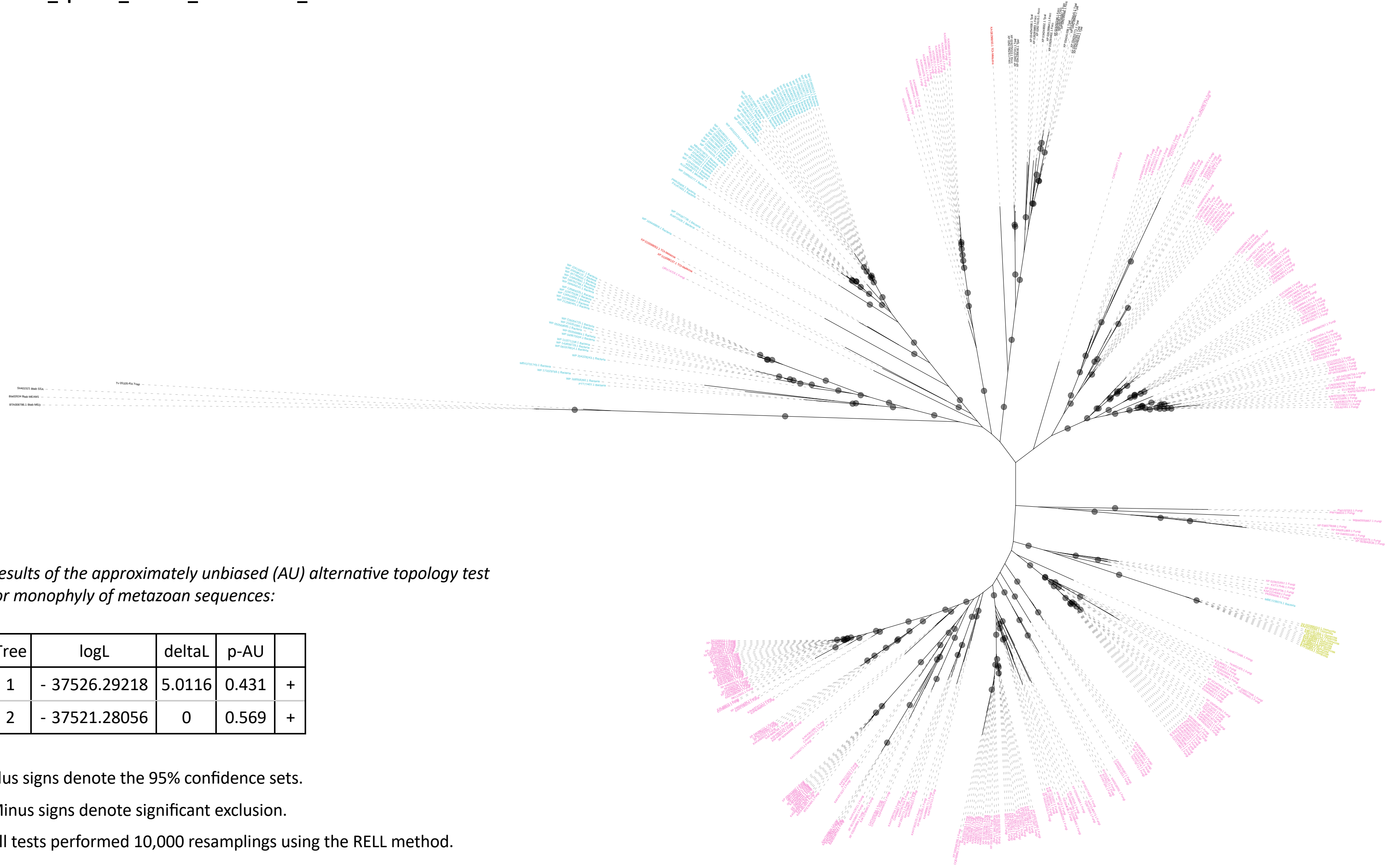

*Results of the approximately unbiased (AU) alternative topology test for monophyly of metazoan sequences:*

| Tree | logL | deltaL | p-AU | |
|---|---|---|---|---|
| 1 | - 37526.29218 | 5.0116 | 0.431 | + |
| 2 | - 37521.28056 | 0 | 0.569 | + |

Plus signs denote the 95% confidence sets.

Minus signs denote significant exclusion.

All tests performed 10,000 resamplings using the RELL method.

### TpaV02: CBM43-CBM43-CBM43-CBM43

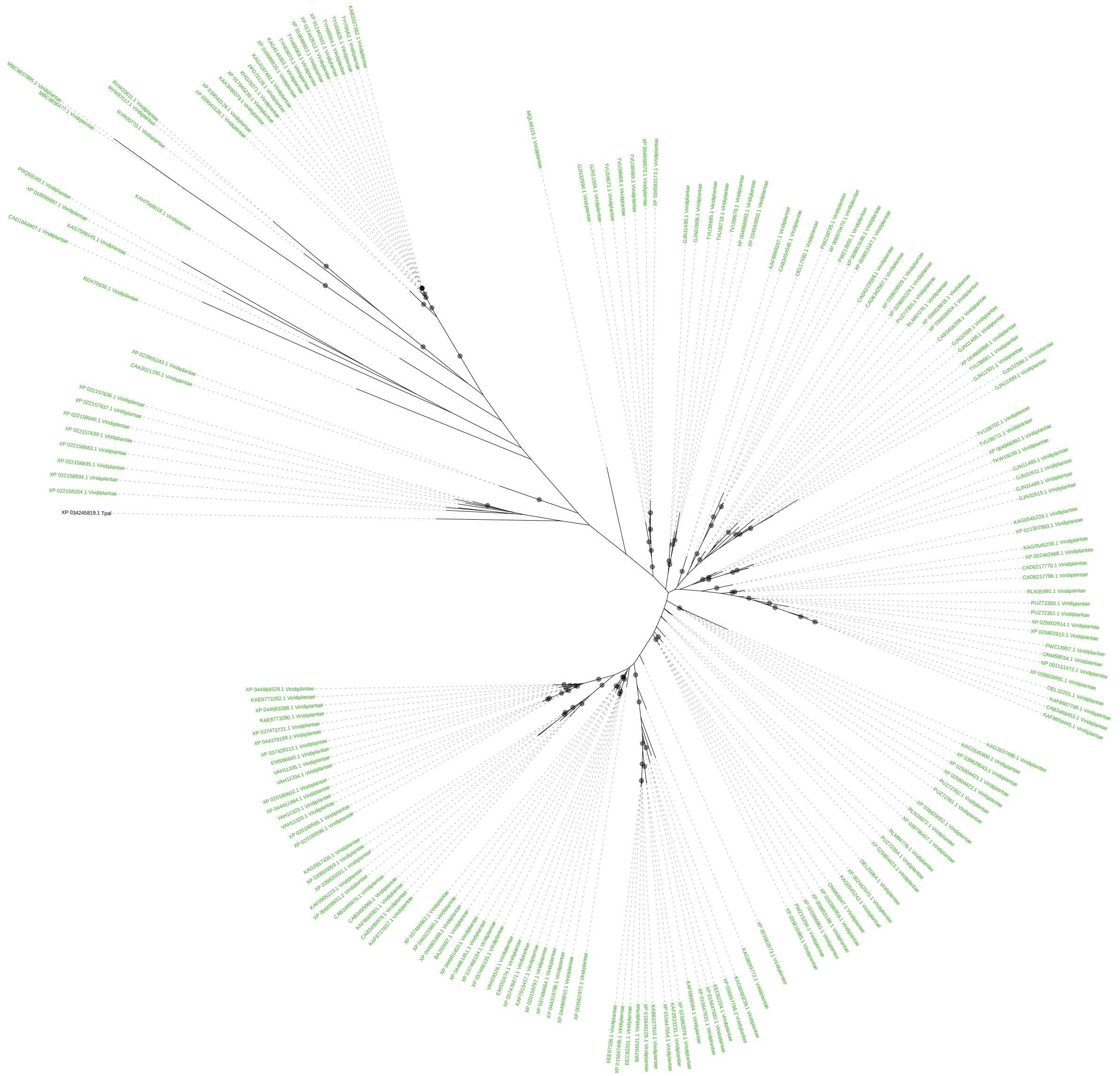
