## Supplementary Tables for "Functional carbohydrate-active enzymes acquired by horizontal gene transfer from plants in the whitefly *Bemisia tabaci*"

**Supplementary Table 1:** Estimation of proteome completeness for *Bemisia tabaci* cryptic species, *Trialeurodes vaporarorium, Frankliniella occidentalis* and *Thrips palmi* using BUSCO according to the Arthropoda Odb10 dataset

| Species | Number of protein-coding genes | BUSCO Arthropoda (1013 groups searched) | | |
|---|---|---|---|---|
| | | Complete[a] | Fragmented | Missing |
| *Bemisia tabaci* MEAM1 | 15,662 | 95.6% [S:94.1%,D:1.5%] | 1.5% | 2.9% |
| *Bemisia tabaci* MED | 20,748 | 86.0% [S:68.8%,D:17.2%] | 3.8% | 10.2% |
| *Bemisia tabaci* SSA-ECA | 15,084 | 89.6% [S:88.0%,D:1.6%] | 3.5% | 6.9% |
| *Trialeurodes vaporarorium* | 18,275 | 93.4% [S:91.6%,D:1.8%] | 1.8% | 4.8% |
| *Frankliniella occidentalis* | 15,678 | 98.8%[S:96.9%,D:1.9%] | 0.5% | 0.7% |
| *Thrips palmi* | 14,332 | 97.5%[S:96.3%,D:1.2%] | 0.1% | 2.4% |

[a] S: Complete and single-copy, D: Complete and duplicated

**Supplementary Table 2:** Validated HGT candidates from potential bacterial, fungal or viridiplantae donors for *B. tabaci* MEAM1

*Bacterial donor:*

| HGT event | Sequence name | Origin of donor sequences | Alternative topology Monophyly of Metazoa | Similarities with the donor sequences | | Length and coverage of the alignment with the donor sequences | | Local score for genomic environment | Homologs found in the two other *B. tabaci* cryptic species | | Homologs found in *T. vaporarorium* | Annotation (WGD version 1.2) | CAZy | Described in literature |
|---|---|---|---|---|---|---|---|---|---|---|---|---|---|---|
| | | | | max id | average id | average aln length | average coverage | | *B. tabaci* MED | *B. tabaci* SSA-ECA | | | | |
| BtaB01 | Bta02653<br>Bta04469<br>Bta05264<br>Bta06818<br>Bta11911<br>Bta11912 | Bacteria | / | 66.1<br>64.9<br>68.5<br>58.1<br>56.3<br>59.3 | 64.7<br>63.7<br>66.4<br>56.4<br>55.2<br>57.1 | 375<br>373<br>273<br>395<br>396<br>394 | 98.5<br>97.8<br>99.6<br>98.2<br>98.3<br>97.5 | 0.67<br>0.9<br>/<br>0.59<br>0.9<br>0.9 | BTA005171.1,<br>BTA005173.1,<br>BTA012513.1,<br>BTA019053.1,<br>BTA023694.1,<br>BTA023695.1,<br>BTA029954.2 | Ssa00993,<br>Ssa02587,<br>Ssa02588,<br>Ssa02589,<br>Ssa09979,<br>Ssa12718 | Tv_08602-RA,<br>Tv_10007-RA | Cyclopropane-fatty-acyl-phospholipid synthase family | | Chen et al. 2016<br>Chen et al. 2016 ; Li et al. 2022<br>No<br>Chen et al. 2016<br>Chen et al. 2016 ; Li et al. 2022<br>Chen et al. 2016 ; Li et al. 2022 |
| BtaB02 | Bta00840<br>Bta01938 | Bacteria | 1 | 72.1<br>70.5 | 54.2<br>52.0 | 200<br>86 | 84.7<br>93.4 | 0.5<br>0.6 | | Ssa01088 | Tv_01894-RA | ATP-dependent dethiobiotin synthetase BioD | | Chen et al. 2016 ; Li et al. 2022<br>Chen et al. 2016 |
| BtaB03 | Bta02987<br>Bta06442 | Bacteria | / | 68.5<br>64.1 | 65.3<br>61.5 | 265<br>260 | 89.1<br>98.6 | 0.96<br>0.8 | BTA003230.1,<br>Scaffold_85_908351_907469 | Ssa15396,<br>scaffold186729_1798_1015 | | Phenazine biosynthesis-like domain-containing protein | | Chen et al. 2016 ; Li et al. 2022<br>Chen et al. 2016 ; Li et al. 2022 |
| BtaB04 | Bta20013 | Bacteria | / | 46.3 | 40.2 | 145 | 65.2 | 0.87 | Scaffold_897_179142_179793 | Ssa06560,<br>Ssa14300,<br>scaffold482_832064_832712_1 | | hypothetical protein | | Chen et al. 2016 |
| BtaB05 | Bta03847 | Bacteria | / | 31.5 | 28.7 | 467 | 71.4 | 0.83 | | | | AAA-ATPase-like domain-containing protein | | Li et al. 2022 |
| BtaB06 | Bta00063 | Bacteria | / | 84.9 | 83.4 | 456 | 98.5 | 0.71 | BTA005853.1 | scaffold95668_2976_6064 | | Argininosuccinate lyase | | Chen et al. 2016 |
| BtaB07 | Bta20014 | Bacteria | / | 61.3 | 59.7 | 703 | 92.3 | 0.8 | BTA005969.2 | Ssa09927 | | methyltransferase | | Chen et al. 2016 |
| BtaB08 | Bta09186 | Bacteria | / | 66.7 | 62.6 | 143 | 92.6 | 0.8 | BTA019104.1 | Ssa07340 | | Ribonuclease H | | Chen et al. 2016 |
| BtaB09 | Bta12386 | Bacteria | / | 40.1 | 32.3 | 433 | 86.0 | 0.79 | BTA006502.1 | Ssa03543,<br>Ssa03544 | Tv_05628-RA | AAA-ATPase-like domain-containing protein | | Li et al. 2022 |
| BtaB10 | Bta01157<br>Bta01158<br>Bta04479<br>Bta05970 | Bacteria | / | 58.1<br>57.7<br>59.4<br>58.5 | 55.3<br>54.9<br>56.8<br>55.4 | 237<br>237<br>240<br>239 | 96.0<br>67.7<br>94.8<br>96.9 | 0.21<br>0.28<br>0.76<br>/ | BTA010421.1,<br>BTA016630.1,<br>scaffold153941_658_1924 | Ssa01004 | | Uncharacterized protein | | No<br>Chen et al. 2016 ; Li et al. 2022<br>Chen et al. 2016 ; Li et al. 2022<br>No |
| BtaB11 | Bta09725 | Bacteria | 1 | 70.6 | 66.2 | 309 | 93.8 | 0.94 | BTA004219.1,<br>BTA028913.1 | Ssa10171 | Tv_15468-RA | Biotin synthase | | Chen et al. 2016 ; Li et al. 2022 |
| BtaB12 | Bta00841<br>Bta01937 | Bacteria | 1 | 70.5<br>73.5 | 55.3<br>59.7 | 273<br>426 | 93.8<br>96.7 | 0.7<br>0.63 | BTA023651.1 | Ssa01553 | Tv_01895-RA | Adenosylmethionine-8-amino-7-oxononanoate aminotransferase BioA | | Chen et al. 2016 ; Li et al. 2022<br>Chen et al. 2016 ; Li et al. 2022 |
| BtaB13 | Bta04607 | Bacteria | / | 30.6 | 27.3 | 451 | 67.5 | 0.61 | BTA010686.1 | Ssa04937 | | AAA-ATPase-like domain-containing protein | | Li et al. 2022 |
| BtaB14 | Bta14802 | Bacteria | / | 68.4 | 63.7 | 398 | 84.1 | 0.92 | BTA007711.2 | Ssa08712 | | Amidohydrolase | | Chen et al. 2016 ; Li et al. 2022 |
| BtaB15 | Bta02416 | Bacteria | / | 46.7 | 41.2 | 66 | 53.0 | 0.83 | | Ssa01016 | | Unknown protein | | No |
| BtaB16 | Bta13770 | Bacteria | / | 32.6 | 29.6 | 101 | 41.6 | 0.91 | BTA022448.2 | Ssa14696 | | Unknown protein | | No |
| BtaB17 | Bta02167 | Bacteria | / | 61.5 | 52.0 | 107 | 10.7 | 0.78 | BTA012693.1 | Ssa02372,<br>Ssa06822 | | Protein NLRC3 | | No |
| BtaB18 | Bta00062 | Bacteria | 1 | 85.9 | 85.0 | 405 | 99.5 | 0.66 | BTA005854.1 | Ssa15432 | | Argininosuccinate synthase | | Chen et al. 2016 |
| BtaB19 | Bta00289<br>Bta02048<br>Bta04448<br>Bta04449<br>Bta07217<br>Bta10169<br>Bta10229<br>Bta10230<br>Bta10232<br>Bta10233<br>Bta14209<br>Bta15534<br>Bta15536 | Bacteria | / | 30<br>31.3<br>31.7<br>31.1<br>30<br>36.8<br>30<br>30.9<br>31.2<br>31.4<br>31.6<br>30.2<br>30.3 | 26.7<br>29.8<br>28.3<br>27.5<br>28.7<br>30.4<br>28.7<br>28.7<br>29.9<br>29.4<br>29.4<br>28.6<br>29.2 | 278<br>471<br>250<br>235<br>337<br>294<br>438<br>430<br>418<br>447<br>431<br>439<br>440 | 27.1<br>72.6<br>73.5<br>91.5<br>98.7<br>63.5<br>67.8<br>71.0<br>69.6<br>71.0<br>71.7<br>73.0<br>66.0 | 0.53<br>0.9<br>0.62<br>0.62<br>/<br>/<br>0.43<br>0.48<br>0.55<br>0.57<br>0.76<br>0.55<br>0.49 | BTA003732.1,<br>BTA004712.1,<br>BTA004713.1,<br>BTA012459.1,<br>BTA014511.1 | Ssa06244,<br>Ssa07626,<br>Ssa07627,<br>Ssa11850,<br>Ssa13138,<br>Ssa14803,<br>Ssa14804,<br>Ssa14962,<br>Ssa15140,<br>Ssa15337,<br>Ssa15382 | | AAA-ATPase-like domain-containing protein | | No<br>Li et al. 2022<br>No<br>No<br>No<br>No<br>No<br>No<br>No<br>Li et al. 2022<br>Li et al. 2022<br>No<br>No |
| BtaB20 | Bta03200 | Bacteria | / | 85.9 | 79.6 | 422 | 96.8 | 0.96 | Scaffold_662_290408_290801 | Ssa03128 | | L-galactonate dehydratase | | Chen et al. 2016 ; Li et al. 2022 |
| BtaB21 | Bta04921 | Bacteria | / | 40.7 | 32.6 | 261 | 47.2 | 0.88 | BTA000806.1,<br>BTA000807.1,<br>BTA008325.1 | Ssa01820 | | Unknown protein | | Chen et al. 2016 ; Li et al. 2022 |
| BtaB22 | Bta03791 | Bacteria | / | 67.3 | 64.7 | 281 | 82.2 | 0.86 | BTA021998.1 | | | Ribosomal RNA small subunit methyltransferase A | | Chen et al. 2016 ; Li et al. 2022 |

| | | | | | | | | | | | | | | |
|---|---|---|---|---|---|---|---|---|---|---|---|---|---|---|
| BtaB23 | Bta03871 | Bacteria | / | 68.1 | 50.7 | 111 | 66.1 | 0.86 | BTA027330.2 | Ssa10244 | | Crossover junction endodeoxyribonuclease rusA | | Chen et al. 2016 ; Li et al. 2022 |
| BtaB24 | Bta06657 | Bacteria | / | 72.0 | 71.0 | 276 | 86.5 | / | BTA007760.1,<br>BTA013028.1 | Ssa11675 | | Diaminopimelate epimerase | | Chen et al. 2016 |
| BtaB25 | Bta15019 | Bacteria | / | 68.1 | 66.6 | 182 | 88.3 | 0.88 | BTA012224.1 | Ssa09419 | | Ribosome recycling factor | | Chen et al. 2016 ; Li et al. 2022 |
| BtaB26 | Bta04508 | Bacteria | / | 73.4 | 72.9 | 1202 | 99.2 | 0.77 | BTA012654.1,<br>BTA017578.1,<br>BTA025646.1 | Ssa06061 | | Urea amidolyase | | Chen et al. 2016 ; Li et al. 2022 |
| BtaB27 | Bta02625<br>Bta04622<br>Bta07024<br>Bta07125<br>Bta07126<br>Bta07870<br>Bta11369<br>Bta11370 | Bacteria | / | 61.1<br>56.5<br>62.1<br>58.0<br>58.5<br>65.4<br>67.4<br>67.8 | 58.8<br>52.8<br>59.5<br>56.2<br>57.3<br>63.9<br>65.9<br>64.8 | 657<br>129<br>642<br>650<br>641<br>640<br>352<br>208 | 98.3<br>70.6<br>96.7<br>97.5<br>89.2<br>98.0<br>97.4<br>93.1 | 0.85<br>0.85<br>0.91<br>0.58<br>0.68<br>0.88<br>0.66<br>0.66 | BTA012629.2,<br>BTA016705.1,<br>BTA016707.1,<br>BTA030125.1,<br>BTA009444.1 | Ssa01508,<br>Ssa08300,<br>Ssa10111,<br>Ssa10113,<br>Ssa10118,<br>Ssa15216 | | Squalene-hopene cyclase | | Chen et al. 2016 ; Li et al. 2022<br>Li et al. 2022<br>Chen et al. 2016 ; Li et al. 2022<br>Chen et al. 2016 ; Li et al. 2022<br>Chen et al. 2016 ; Li et al. 2022<br>Chen et al. 2016 ; Li et al. 2022<br>Chen et al. 2016 ; Li et al. 2022<br>Chen et al. 2016 |
| BtaB28 | Bta11811 | Bacteria | / | 79.7 | 72.9 | 298 | 97.5 | 0.81 | | Ssa05133 | | Plant/T7N9-9 protein | | Chen et al. 2016 ; Li et al. 2022 |
| BtaB29 | Bta20020 | Bacteria | / | 58.9 | 52.4 | 245 | 97.4 | 0.92 | Scaffold_733_232<br>89_24021 | scaffold570_6229<br>85_622253 | | 4-hydroxy-tetrahydrodipicolinate reductase | | Chen et al. 2016 ; Li et al. 2022 |
| BtaB30 | Bta03589<br>Bta03593 | Bacteria | / | 70.1<br>69.3 | 60.8<br>60.3 | 214<br>426 | 83.6<br>94.7 | 0.86<br>0.86 | BTA026180.1 | Ssa05826 | | Diaminopimelate decarboxylase | | Li et al. 2022<br>Chen et al. 2016 ; Li et al. 2022 |
| BtaB31 | Bta00427 | Bacteria | / | 69.3 | 62.5 | 209 | 72.3 | 0.84 | BTA000821.1 | Ssa07839 | | Pantothenate kinase-like protein | | Li et al. 2022 |
| BtaB32 | Bta11965 | Bacteria | / | 46.3 | 38.9 | 130 | 73.4 | 0.81 | BTA017096.1 | Ssa00217 | | Unknown protein | | No |
| BtaB33 | Bta03797 | Bacteria | / | 62.5 | 58.2 | 103 | 34.4 | 0.9 | BTA000817.1 | Ssa13701 | Tv_04874-RA | Methylated-DNA--[protein]-cysteine S-methyltransferase | | Li et al. 2022 |
| BtaB34 | Bta08776 | Bacteria | / | 82.9 | 80.6 | 513 | 95.9 | 1.0 | BTA025302.1,<br>BTA028148.1 | Ssa15043 | | Histidine ammonia-lyase | | Chen et al. 2016 |
| BtaB35 | Bta11531<br>Bta15479 | Bacteria | / | 30.6<br>33 | 28.6<br>30.0 | 526<br>439 | 79.1<br>90.5 | 0.9<br>0.92 | BTA004921.1,<br>BTA018889.1,<br>BTA026605.1 | Ssa12442 | Tv_02105-RA | AAA-ATPase-like domain-containing protein | | No<br>Li et al. 2022 |
| BtaB36 | Bta00829 | Bacteria | / | 33.3 | 30.7 | 526 | 71.4 | 0.45 | BTA007316.1,<br>BTA011402.1 | scaffold554257_3<br>12_1956 | Tv_00146-RA,<br>Tv_00148-RA,<br>Tv_01481-RA,<br>Tv_02102-RA,<br>Tv_02106-RA,<br>Tv_15366-RA | AAA-ATPase-like domain-containing protein | | No |
| BtaB37 | Bta15184<br>Bta15191<br>Bta20012 | Bacteria | / | 68.1<br>70.8<br>78.1 | 66.3<br>69.3<br>75.6 | 255<br>256<br>255 | 98.9<br>99.2<br>99.0 | 0.61<br>0.75<br>0.57 | BTA021540.1,<br>BTA021541.1,<br>BTA030079.2,<br>BTA030081.1,<br>BTA030084.1 | | | Oxidoreductase, 2OG-Fe(II) oxygenase family protein | | Chen et al. 2016 ; Li et al. 2022<br>Chen et al. 2016 ; Li et al. 2022<br>Chen et al. 2016 ; Li et al. 2022 |
| BtaB38 | Bta20015 | Bacteria | / | 56.3 | 47.1 | 443 | 88.0 | 0.9 | BTA012131.1 | | | hypothetical protein | | Chen et al. 2016 ; Li et al. 2022 |
| BtaB39 | Bta06820 | Bacteria | / | 61.6 | 60.7 | 336 | 85.8 | 0.63 | BTA019051.2 | Ssa09977 | | Tryptophan-tRNA ligase | | Chen et al. 2016 |
| BtaB40 | Bta02826 | Bacteria | / | 35.2 | 27.6 | 445 | 70.7 | 0.86 | BTA005203.1 | Ssa08291 | | AAA-ATPase-like domain-containing protein | | No |
| BtaB41 | Bta04431 | Bacteria | / | 75.4 | 70.1 | 183 | 91.2 | 1.0 | BTA007219.2,<br>BTA020661.1 | Ssa00937 | | Acetyltransferase | | Chen et al. 2016 ; Li et al. 2022 |
| BtaB42 | Bta05813 | Bacteria | / | 54.9 | 47.1 | 318 | 45.9 | 0.71 | BTA005145.1 | Ssa12681 | | D-alanine--D-alanine ligase | | No |
| BtaB43 | Bta02812 | Bacteria | / | 70.8 | 69.0 | 264 | 99.2 | 0.68 | BTA006638.1,<br>BTA006641.1 | scaffold413_2442<br>29_245024 | | 4,5 dioxygenase extradiol | | Chen et al. 2016 ; Li et al. 2022 |
| BtaB44 | Bta02807<br>Bta02808 | Bacteria | / | 42.9<br>43.1 | 37.4<br>39.0 | 115<br>111 | 82.4<br>72.6 | 0.55<br>0.6 | Scaffold_166_324<br>443_324098,<br>Scaffold_166_330<br>260_329816 | Ssa08024 | | AAA-ATPase-like domain-containing protein | | No<br>Li et al. 2022 |
| BtaB45 | Bta15103 | Bacteria | / | 50.6 | 44.6 | 151 | 71.6 | 0.88 | BTA020874.1 | Ssa07364 | Tv_06835-RA | Chorismate mutase 1 | | Chen et al. 2016 ; Li et al. 2022 |
| BtaB46 | Bta07556<br>Bta07557 | Bacteria | /<br>/ | 57.1<br>58.3 | 51.4<br>52.6 | 111<br>106 | 46.6<br>44.0 | 0.78<br>0.78 | BTA024248.1 | Ssa13644,<br>Ssa13645,<br>Ssa13646 | | Uncharacterized protein | | No<br>Li et al. 2022 |
| BtaB47 | Bta04809 | Bacteria | / | 30.8 | 29.2 | 412 | 69.2 | 0.56 | BTA006704.1 | Ssa06266 | | AAA-ATPase-like domain-containing protein | | Li et al. 2022 |
| BtaB48 | Bta04987<br>Bta04988 | Bacteria | / | 72.3<br>71.7 | 69.0<br>70.0 | 277<br>285 | 88.7<br>95.4 | 0.82<br>0.86 | BTA020296.1,<br>BTA021770.1,<br>BTA021771.2 | Ssa00739,<br>Ssa00740 | | Amidinotransferase | | Chen et al. 2016 ; Li et al. 2022<br>Chen et al. 2016 ; Li et al. 2022 |
| BtaB49 | Bta13948 | Bacteria | / | 62.6 | 60.9 | 177 | 69.0 | 0.76 | Scaffold_155_308<br>817_300779 | scaffold3967_193<br>336_184613 | | DUF1768-domain-containing protein | | No |
| BtaB50 | Bta15764 | Bacteria | / | 65.0 | 60.3 | 121 | 93.8 | 0.16 | | Ssa02340 | | Uncharacterized protein | | No |
| BtaB51 | Bta05339 | Bacteria | 1 | 72.1 | 66.6 | 285 | 45.3 | 0.91 | BTA017168.3 | Ssa09543 | | 3-methyl-2-oxobutanoate hydroxymethyltransferase | | Chen et al. 2016 ; Li et al. 2022 |

| | | | | | | | | | | | | | | |
|---|---|---|---|---|---|---|---|---|---|---|---|---|---|---|
| BtaB52 | Bta12603 | Bacteria | / | 31.6 | 28.1 | 475 | 75.5 | 0.96 | BTA027018.1 | scaffold190238_1244_716 | | AAA-ATPase-like domain-containing protein | | Li et al. 2022 |
| BtaB53 | Bta00773<br>Bta04549 | Bacteria | / | 49.1<br>46.8 | 45.5<br>44.1 | 476<br>487 | 80.3<br>88.1 | 0.44<br>0.88 | BTA011593.1,<br>BTA012235.1,<br>Scaffold_3889_9990_14442 | scaffold1153_152506_171880 | | Beta-fructofuranosidase | GH32 | Li et al. 2022<br>Li et al. 2022 |
| BtaB54 | Bta07115 | Bacteria | / | 76.4 | 71.8 | 258 | 72.3 | 0.52 | | scaffold333079_2023_1150 | | Meiotic chromosome segregation protein | | Li et al. 2022 |
| BtaB55 | Bta09846 | Bacteria | / | 31.3 | 29.0 | 569 | 87.8 | 0.81 | BTA001371.1 | Ssa14624 | | AAA-ATPase-like domain-containing protein | | Li et al. 2022 |
| BtaB56 | Bta10852<br>Bta10853 | Bacteria | / | 36.8<br>26.1 | 29.5<br>23.5 | 476<br>207 | 80.6<br>74.5 | 0.59<br>0.64 | BTA009628.1,<br>BTA009629.1,<br>BTA025251.1,<br>BTA025252.1,<br>BTA025253.1 | Ssa11036 | TLow_01347-RA,<br>TLow_02144-RA,<br>TLow_02157-RA,<br>Tv_13818-RA,<br>Tv_13862-RA,<br>Tv_15776-RA | AAA-ATPase-like domain-containing protein | | Li et al. 2022<br>No |
| BtaB57 | Bta00871 | Bacteria | / | 53.7 | 50.0 | 226 | 79.2 | 0.95 | BTA021726.1 | Ssa14229 | | Ribonuclease 3 | | Chen et al. 2016 |
| BtaB58 | Bta13949 | Bacteria | / | 72.2 | 70.4 | 597 | 97.3 | 0.76 | BTA005651.1 | Ssa03692 | | Glutamyl-tRNA(Gln) amidotransferase subunit A, putative | | Chen et al. 2016 |
| BtaB59 | Bta15712 | Bacteria | / | 61.7 | 56.8 | 125 | 50.1 | 0.27 | BTA015916.1 | Ssa02297 | | SCP-like extracellular | | Chen et al. 2016 |
| BtaB60 | Bta13118 | Bacteria | / | 79.5 | 76.2 | 219 | 97.8 | 0.77 | BTA014321.1 | | | GNAT family acetyltransferase | | Chen et al. 2016 ; Li et al. 2022 |
| BtaB61 | Bta07442 | Bacteria | / | 73.4 | 65.5 | 190 | 98.0 | 0.73 | BTA023294.1 | scaffold763_1615070_1615649 | | YJL217W-like protein | | Li et al. 2022 |
| BtaB62 | Bta15714 | Bacteria | / | 60.0 | 56.1 | 193 | 87.2 | 0.36 | BTA015913.1 | | | SCP-like extracellular | CBM50 | Chen et al. 2016 ; Li et al. 2022 |
| BtaB63 | Bta00042<br>Bta02417 | Bacteria | / | 62.6<br>38.0 | 57.7<br>27.5 | 405<br>202 | 96.2<br>94.1 | 0.29<br>0.83 | BTA016358.1,<br>Scaffold_5042_3912_2826 | Ssa10934,<br>scaffold26337_34592_35222 | | Ser/Thr protein phosphatase family | | Li et al. 2022<br>Chen et al. 2016 |
| BtaB64 | Bta05687 | Bacteria | / | 50.0 | 44.9 | 145 | 55.8 | 0.48 | Scaffold_2144_73225_73981 | Ssa04068 | | Deoxyuridine 5'-triphosphate nucleotidohydrolase | | No |
| BtaB65 | Bta05013<br>Bta10549<br>Bta14973 | Bacteria | / | 32.1<br>32<br>32.3 | 30.3<br>28.9<br>29.9 | 511<br>546<br>516 | 79.9<br>73.4<br>78.9 | 0.67<br>0.85<br>0.82 | BTA002393.1 | Ssa01599,<br>Ssa10865,<br>Ssa10873,<br>Ssa12918,<br>Ssa13905,<br>Ssa14120,<br>Ssa15393 | Tv_00487-RA,<br>Tv_11451-RA | AAA-ATPase-like domain-containing protein | | No<br>Li et al. 2022<br>No |
| BtaB66 | Bta15713 | Bacteria | / | 63.3 | 50.5 | 176 | 39.0 | 0.32 | BTA015915.1 | Ssa02298 | | SCP-like extracellular | | Chen et al. 2016 ; Li et al. 2022 |
| BtaB67 | Bta04447 | Bacteria | / | 40.1 | 33.2 | 397 | 86.0 | 0.62 | BTA008945.6 | Ssa00953 | | AAA-ATPase-like domain-containing protein | | Li et al. 2022 |
| BtaB68 | Bta00747<br>Bta13975 | Bacteria | / | 49.6<br>52.0 | 44.7<br>45.3 | 146<br>144 | 94.6<br>95.7 | /<br>0.91 | | Ssa08703,<br>Ssa14910 | Tv_02831-RA,<br>Tv_11854-RA,<br>Tv_12101-RA,<br>Tv_15086-RA,<br>Tv_15720-RA,<br>Tv_15881-RA | Deoxyuridine 5'-triphosphate nucleotidohydrolase | | No<br>No |
| BtaB69 | Bta00002 | Bacteria | / | 53.7 | 52.5 | 713 | 99.1 | 0.78 | BTA004354.1,<br>BTA008424.1 | Ssa09173 | | ATP-dependent DNA helicase Q-like 3 | | Chen et al. 2016 ; Li et al. 2022 |
| BtaB70 | Bta05382 | Bacteria | 0 | 46.8 | 43.8 | 137 | 20.4 | 0.76 | BTA000114.1 | Ssa05150 | | Extracellular dioxygenase | | Li et al. 2022 |

*Fungal donor:*

| HGT event | Sequence name | Origin of donor sequences | Alternative topology | Similarities with the donor sequences | | Length and coverage of the alignment with the donor sequences | | Local score for genomic environment | Homologs found in the two other *B. tabaci* cryptic species | | Homologs found in *T. vaporarorium* | Annotation (WGD version 1.2) | CAZy | Described in literature |
|---|---|---|---|---|---|---|---|---|---|---|---|---|---|---|
| | | | | max id | average id | average aln length | average coverage | | *B. tabaci* MED | *B. tabaci* SSA-ECA | | | | |
| BtaF01 | Bta00659<br>Bta00660 | Fungi | / | 54.1<br>49.6 | 49.3<br>45.6 | 377<br>404 | 86.7<br>91.5 | 0.55<br>0.49 | BTA010566.1,<br>BTA010567.1,<br>BTA010568.1 | Ssa03926 | | Aromatic peroxygenase | | Chen et al. 2016 ; Li et al. 2022<br>Chen et al. 2016 ; Li et al. 2022 |
| BtaF02 | Bta12907<br>Bta12908 | Fungi | / | 46.9<br>47.8 | 43.0<br>43.0 | 343<br>348 | 68.8<br>92.4 | 0.82<br>0.82 | BTA009603.1,<br>BTA017757.1,<br>BTA009604.1,<br>BTA017756.1 | Ssa10521,<br>Ssa10522,<br>Ssa10524 | | Late sexual development protein | | Chen et al. 2016 ; Li et al. 2022<br>Chen et al. 2016 ; Li et al. 2022 |
| BtaF03 | Bta02178 | Fungi | / | 43.5 | 40.3 | 438 | 91.9 | 0.92 | BTA007752.1 | Ssa06839 | | Aminotransferase family protein (LolT), putative | | Chen et al. 2016 ; Li et al. 2022 |
| BtaF04 | Bta10926<br>Bta13511 | Fungi | / | 30.8<br>32.4 | 28.8<br>29.3 | 106<br>106 | 38.8<br>45.3 | 0.92<br>0.91 | BTA000754.1,<br>BTA011323.1,<br>BTA016968.1,<br>BTA023763.1,<br>BTA025826.1,<br>BTA026082.1 | Ssa02288 | | Replication factor-a protein 1 | | No<br>No |
| BtaF05 | Bta04178<br>Bta14373<br>Bta14374 | Fungi | / | 46.1<br>43.9<br>48.8 | 27.9<br>28.4<br>36.9 | 911<br>914<br>515 | 84.7<br>60.8<br>73.2 | 1.0<br>0.5<br>0.54 | BTA015808.1,<br>BTA015810.1,<br>BTA026116.1,<br>BTA026115.2 | Ssa02954,<br>Ssa02955,<br>Ssa04524,<br>Ssa04525,<br>Ssa04526 | | 2OG-Fe(II) oxygenase superfamily protein | | Chen et al. 2016 ; Li et al. 2022<br>Chen et al. 2016 ; Li et al. 2022<br>Chen et al. 2016 ; Li et al. 2022 |
| BtaF06 | Bta02658 | Fungi | / | 66.5 | 60.6 | 461 | 95.0 | 0.72 | | Ssa02594 | | Glucosylceramidase, putative | GH30 | Chen et al. 2016 ; Li et al. 2022 |
| BtaF07 | Bta02250 | Fungi | 1 | 63.8 | 51.6 | 580 | 92.6 | 0.77 | BTA018568.2 | Ssa14166 | | Dextranase | GH49 | Chen et al. 2016 ; Li et al. 2022 |
| BtaF08 | Bta14185 | Fungi | / | 42.3 | 39.9 | 677 | 98.8 | 0.92 | | Ssa14298 | | Uracil phosphoribosyltransferase | | Chen et al. 2016 ; Li et al. 2022 |
| BtaF09 | Bta13117 | Fungi | 1 | 59.0 | 48.3 | 402 | 94.5 | 0.74 | BTA014319.1 | scaffold461923_670_139 | | Serine/threonine dehydratase serine racemase | | Chen et al. 2016 |
| BtaF10 | Bta01809<br>Bta01821<br>Bta04681<br>Bta05463<br>Bta05465<br>Bta05467<br>Bta06167<br>Bta07455<br>Bta10729<br>Bta10733<br>Bta10734<br>Bta10735<br>Bta10736<br>Bta10737<br>Bta10760<br>Bta10761<br>Bta14699<br>Bta14700<br>Bta14701<br>Bta14896 | Fungi | / | 45.1<br>47.0<br>50.6<br>40.4<br>41.9<br>37.1<br>44.3<br>43.8<br>44.5<br>38.6<br>39.9<br>39.9<br>38.7<br>40.0<br>53.1<br>47.0<br>41.1<br>41.9<br>48.6<br>48.8 | 41.2<br>45.1<br>48.0<br>38.2<br>40.7<br>35.6<br>41.1<br>41.4<br>40.7<br>36.9<br>35.3<br>34.8<br>37.5<br>38.6<br>50.2<br>43.6<br>37.9<br>38.9<br>46.2<br>45.3 | 400<br>405<br>365<br>403<br>405<br>397<br>248<br>411<br>382<br>405<br>213<br>157<br>402<br>402<br>398<br>416<br>416<br>406<br>410<br>408 | 92.7<br>77.3<br>70.5<br>86.2<br>87.9<br>82.2<br>46.9<br>95.1<br>52.4<br>87.5<br>84.5<br>92.3<br>88.5<br>80.8<br>54.7<br>91.2<br>96.0<br>93.6<br>83.7<br>94.6 | 0.96<br>0.82<br>0.65<br>0.8<br>0.7<br>0.7<br>0.95<br>0.84<br>0.5<br>0.47<br>0.44<br>0.41<br>0.37<br>0.32<br>0.66<br>0.66<br>0.76<br>0.76<br>0.76<br>0.92 | BTA021351.1,<br>BTA016059.1,<br>BTA024446.1,<br>BTA028865.1,<br>BTA020831.1,<br>BTA021213.1,<br>BTA029208.1,<br>BTA029214.1,<br>BTA029213.1,<br>BTA029215.1,<br>BTA029216.1,<br>BTA001574.1,<br>BTA001575.1,<br>BTA022986.1,<br>BTA022987.1,<br>BTA022989.1,<br>BTA022990.1,<br>BTA022991.1 | Ssa10324,<br>Ssa01851,<br>Ssa09478,<br>Ssa09479,<br>Ssa09482,<br>Ssa06288,<br>Ssa00368,<br>Ssa12703,<br>Ssa10743,<br>Ssa10745,<br>Ssa00340,<br>Ssa00341,<br>Ssa09259,<br>Ssa09258,<br>Ssa09257,<br>Ssa10800 | | Major royal jelly-related protein | | Chen et al. 2016<br>Chen et al. 2016<br>Chen et al. 2016 ; Li et al. 2022<br>Chen et al. 2016 ; Li et al. 2022<br>Li et al. 2022<br>Li et al. 2022<br>Chen et al. 2016<br>Chen et al. 2016 ; Li et al. 2022<br>Chen et al. 2016<br>Chen et al. 2016 ; Li et al. 2022<br>No<br>No<br>Chen et al. 2016<br>Chen et al. 2016 ; Li et al. 2022<br>Chen et al. 2016 ; Li et al. 2022<br>Chen et al. 2016 ; Li et al. 2022<br>Chen et al. 2016<br>Chen et al. 2016<br>Chen et al. 2016<br>Chen et al. 2016 |
| BtaF11 | Bta11792<br>Bta03426 | Fungi | / | 47.9<br>54.3 | 38.5<br>40.3 | 140<br>189 | 50.5<br>74.9 | 0.92<br>0.82 | BTA002831.1,<br>BTA002830.1,<br>BTA019829.1,<br>BTA029129.1 | Ssa05110,<br>scaffold50840_2_365 | Tv_10871-RA,<br>Tv_10872-RA,<br>Tv_11256-RA | Unknown protein | | No<br>No |
| BtaF12 | Bta03315 | Fungi | / | 65.9 | 56.2 | 342 | 96.1 | 0.88 | Scaffold_934_82522_83563 | Ssa11275 | | Adenosine deaminase | | Chen et al. 2016 |
| BtaF13 | Bta12326<br>Bta20017 | Fungi | / | 35.3<br>33.7 | 31.4<br>29.6 | 257<br>413 | 89.2<br>88.4 | 0.91<br>0.96 | BTA009468.1 | scaffold5114_691867_692650 | | Activating signal cointegrator 1 complex subunit-like protein | | Chen et al. 2016<br>Chen et al. 2016 |
| BtaF14 | Bta11043 | Fungi | / | 62.3 | 59.7 | 400 | 92.4 | 1.0 | BTA013802.1,<br>BTA015522.1 | Ssa00130,<br>Ssa00131 | | Squalene synthase | | Chen et al. 2016 |
| BtaF15 | Bta03557<br>Bta05545 | Fungi | / | 72.9<br>67.4 | 67.6<br>63.2 | 608<br>609 | 99.4<br>99.6 | 0.74<br>0.84 | BTA001108.1,<br>BTA022973.1,<br>BTA025701.1,<br>BTA027649.1 | Ssa05748,<br>Ssa13062 | | Gamma-glutamyltranspeptidase | | Chen et al. 2016<br>Chen et al. 2016 |
| BtaF16 | Bta02866 | Fungi | / | 52.7 | 42.3 | 426 | 94.8 | 1.0 | BTA013774.1 | scaffold8_3079009_3077710 | | Alpha-1,3-glucanase, putative | GH71 | Chen et al. 2016 ; Li et al. 2022 |
| BtaF17 | Bta14111 | Fungi | 0 | 41.0 | 36.5 | 216 | 93.2 | 1.0 | BTA014094.1 | Ssa09898 | | Transcription factor Rba50, putative | | Chen et al. 2016 ; Li et al. 2022 |

| | | | | | | | | | | | | | | |
|---|---|---|---|---|---|---|---|---|---|---|---|---|---|---|
| BtaF18 | Bta20021 | Fungi | / | 60.1 | 56.0 | 590 | 98.5 | 0.76 | BTA012080.1 | scaffold894_3378<br>74_339566 | | cryptochrome | | Chen et al. 2016 |
| BtaF19 | Bta01072<br>Bta03567<br>Bta05090<br>Bta05091<br>Bta05092<br>Bta05093<br>Bta07691<br>Bta09066<br>Bta12453 | Fungi | 1 | 58.2<br>46.2<br>53.3<br>55.1<br>56.2<br>55.3<br>46.2<br>55.8<br>51.3 | 56.0<br>44.5<br>52.3<br>53.5<br>50.6<br>53.8<br>45.6<br>54.8<br>47.2 | 655<br>651<br>644<br>636<br>375<br>637<br>644<br>650<br>127 | 95.4<br>93.9<br>87.9<br>89.7<br>99.0<br>89.8<br>92.3<br>93.8<br>74.8 | 0.71<br>0.78<br>0.33<br>0.38<br>0.43<br>0.42<br>0.92<br>1.0<br>/ | BTA004602.1,<br>BTA010570.1,<br>BTA009422.2,<br>BTA010036.3,<br>BTA001712.1 | Ssa03924,<br>Ssa14575,<br>Ssa02112,<br>Ssa02113,<br>Ssa07603,<br>Ssa07877 | | Galactose oxidase | AA5+CBM3<br>2 | Chen et al. 2016 ; Li et al. 2022<br>Chen et al. 2016 ; Li et al. 2022<br>Li et al. 2022<br>Chen et al. 2016 ; Li et al. 2022<br>Chen et al. 2016<br>Chen et al. 2016 ; Li et al. 2022<br>Chen et al. 2016 ; Li et al. 2022<br>Chen et al. 2016 ; Li et al. 2022<br>No |
| BtaF20 | Bta15600<br>Bta15601 | Fungi | / | 45.7<br>46.9 | 43.0<br>43.1 | 175<br>233 | 90.6<br>76.3 | /<br>/ | BTA008707.1 | Ssa12367 | | Aromatic peroxygenase | | Chen et al. 2016<br>Chen et al. 2016 ; Li et al. 2022 |
| BtaF21 | Bta01073<br>Bta07721 | Fungi | / | 55.9<br>54.5 | 52.9<br>51.0 | 358<br>389 | 54.4<br>93.2 | 0.66<br>0.77 | BTA004600.1,<br>BTA004601.1,<br>BTA010569.1,<br>BTA017481.1 | Ssa03925,<br>Ssa07646 | | Aromatic peroxygenase | | Chen et al. 2016 ; Li et al. 2022<br>Chen et al. 2016 ; Li et al. 2022 |
| BtaF22 | Bta00932<br>Bta05475<br>Bta07433 | Fungi | / | 33.0<br>32.1<br>32.1 | 30.7<br>30.4<br>30.3 | 1247<br>1257<br>1254 | 99.3<br>99.4<br>98.4 | 0.93<br>0.76<br>0.45 | BTA005597.1,<br>BTA005598.1,<br>BTA005599.1,<br>BTA007606.1,<br>BTA003000.1,<br>BTA004644.1,<br>BTA008032.1,<br>BTA008563.1 | sa06477,<br>Ssa06478,<br>Ssa06479,<br>Ssa04914,<br>Ssa15384 | Tv_08361-RA,<br>Tv_14110-RA,<br>Tv_14111-RA | MYND finger family protein | | Chen et al. 2016<br>Chen et al. 2016 ; Li et al. 2022<br>Chen et al. 2016 ; Li et al. 2022 |
| BtaF23 | Bta05635 | Fungi | / | 64.7 | 60.7 | 243 | 84.3 | 0.91 | Scaffold_31_8000<br>23_801601 | scaffold47637_45<br>_1696 | | Duf1275 domain protein | | Chen et al. 2016 ; Li et al. 2022 |
| BtaF24 | Bta10569 | Fungi | / | 59.0 | 49.3 | 611 | 93.5 | 0.84 | BTA015701.1 | scaffold299226_1<br>298_59 | | Tyrosyl-DNA phosphodiesterase<br>domain protein | | Chen et al. 2016 ; Li et al. 2022 |
| BtaF25 | Bta20016 | Fungi | / | 78.4 | 70.5 | 164 | 90.9 | 0.9 | BTA012132.1,<br>BTA021974.1 | scaffold532168_5<br>9820_60357 | | cyanate hydratase | | Chen et al. 2016 ; Li et al. 2022 |
| BtaF26 | Bta08944<br>Bta08945<br>Bta08946<br>Bta14076 | Fungi | / | 49.0<br>46.8<br>53.0<br>55.5 | 43.6<br>41.9<br>48.9<br>49.5 | 1181<br>1244<br>2102<br>1606 | 94.4<br>88.5<br>99.7<br>99.8 | 0.68<br>0.71<br>0.73<br>/ | BTA016279.2,<br>BTA025951.2,<br>BTA025953.1,<br>BTA025954.1 | Ssa11553,<br>Ssa12591,<br>Ssa15247 | Tv_07966-RA | NFX1-type zinc finger-containing<br>protein 1 | | Chen et al. 2016<br>Chen et al. 2016<br>Chen et al. 2016<br>No |
| BtaF27 | Bta10703 | Fungi | / | 41.4 | 35.5 | 351 | 82.7 | 0.88 | BTA013842.1 | Ssa02002 | | Zinc finger MYND-type protein | | Chen et al. 2016 |
| BtaF28 | Bta00658 | Fungi | / | 53.6 | 50.0 | 387 | 87.9 | 0.6 | BTA010565.1 | Ssa03927 | | Aromatic peroxygenase | | Chen et al. 2016 ; Li et al. 2022 |
| BtaF29 | Bta00041 | Fungi | / | 54.8 | 52.1 | 388 | 87.5 | 0.34 | BTA016362.1 | Ssa10936 | | Aromatic peroxygenase | | Chen et al. 2016 ; Li et al. 2022 |
| BtaF30 | Bta08225 | Fungi | | 44.9 | 41.1 | 270 | 64.6 | 0.9 | BTA009130.1 | Ssa05079 | | Aromatic peroxygenase | | Chen et al. 2016 |
| BtaF31 | Bta08226 | Fungi | / | 56.0 | 46.7 | 111 | 25.8 | 0.9 | Scaffold_520_329<br>177_399713 | scaffold1911_304<br>567_303956 | | Aromatic peroxygenase | | Chen et al. 2016 |
| BtaF32 | Bta05277<br>Bta20018<br>Bta20019 | Fungi | / | 48.4<br>43.6<br>54.5 | 34.0<br>33.8<br>47.4 | 329<br>338<br>218 | 86.9<br>86.5<br>78.0 | 0.56<br>0.8<br>0.8 | BTA017099.3 | Ssa13377,<br>Ssa13378 | | Virginiamycin B lyase protein | | No<br>Chen et al. 2016<br>Chen et al. 2016 |
| BtaF33 | Bta01735 | Fungi | / | 64.4 | 60.9 | 479 | 90.9 | 0.87 | BTA009203.1 | Ssa04872 | | Glutamyl-tRNA(Gln)<br>amidotransferase subunit A | | Li et al. 2022 |
| BtaF34 | Bta11373 | Fungi | / | 45.9 | 43.7 | 377 | 76.8 | 0.76 | Scaffold_424_428<br>395_446441 | Ssa15364 | | Aromatic peroxygenase | | Chen et al. 2016 |
| BtaF35 | Bta02217<br>Bta05754 | Fungi | | 66.4<br>57.4 | 59.6<br>55.1 | 145<br>194 | 83.2<br>90.6 | 0.66<br>0.75 | BTA008151.1 | scaffold1412_248<br>109_245904 | | Aromatic peroxygenase | | Chen et al. 2016 ; Li et al. 2022<br>Chen et al. 2016 |
| BtaF36 | Bta04808 | Fungi | / | 43.9 | 41.3 | 369 | 79.1 | 0.51 | BTA006705.1 | Ssa06267 | | Aromatic peroxygenase | | Chen et al. 2016 ; Li et al. 2022 |
| BtaF37 | Bta07948 | Fungi | / | 30.8 | 29.7 | 1319 | 99.8 | 0.92 | BTA017217.1,<br>BTA017221.1,<br>BTA021006.1 | Ssa00835 | Tv_08362-RA | MYND finger family protein | | Chen et al. 2016 ; Li et al. 2022 |
| BtaF38 | Bta00038<br>Bta00039<br>Bta05684<br>Bta05753<br>Bta05981<br>Bta15602 | Fungi | / | 56.4<br>55.3<br>48.7<br>57.1<br>40.0<br>48.1 | 53.2<br>52.1<br>46.0<br>49.7<br>37.7<br>45.5 | 386<br>388<br>379<br>155<br>371<br>369 | 74.6<br>75.9<br>92.4<br>78.3<br>79.7<br>89.2 | 0.44<br>0.41<br>0.3<br>0.75<br>0.82<br>/ | BTA016364.1,<br>BTA022954.1,<br>BTA008707.1,<br>BTA006705.1,<br>BTA008708.1 | Ssa05078,<br>Ssa10939,<br>Ssa10938,<br>Ssa10588,<br>Ssa10589,<br>Ssa12367,<br>Ssa06267,<br>Ssa12366 | | Aromatic peroxygenase | | Chen et al. 2016 ; Li et al. 2022<br>Chen et al. 2016 ; Li et al. 2022<br>Chen et al. 2016 ; Li et al. 2022<br>Chen et al. 2016<br>Chen et al. 2016 ; Li et al. 2022<br>Chen et al. 2016 ; Li et al. 2022 |

*Viridiplantae donor:*

| HGT event | Sequence name | Origin of donor sequences | Alternative topology | Similarities with the donor sequences | | Length and coverage of the alignment with the donor sequences | | Local score for genomic environment | Homologs found in the two other *B. tabaci* cryptic species | | Homologs found in *T. vaporarorium* | Annotation (WGD version 1.2) | CAZy | Described in literature |
|---|---|---|---|---|---|---|---|---|---|---|---|---|---|---|
| | | | | max id | average id | average aln length | average coverage | | *B. tabaci* MED | *B. tabaci* SSA-ECA | | | | |
| BtaV01 | Bta06115 | Viridiplantae | / | 71.2 | 66.4 | 318 | 91.8 | 0.82 | BTA001223.1, Scaffold_1494_158262_155412 | Ssa15230, scaffold6392_704_1667 | | Glucan endo-1,3-beta-glucosidase | GH17 | Gilbert and Maumus 2022 |
| | Bta06118 | | | 58.8 | 56.8 | 318 | 87.4 | 0.81 | | | | | | Gilbert and Maumus 2022 ; Li et al. 2022 |
| BtaV02 | Bta13961 | Viridiplantae | / | 59.0 | 54.7 | 232 | 93.6 | 0.96 | BTA005662.2 | Ssa10394 | Tv_15928-RA | Thaumatin-like protein 1a | GH152 | Gilbert and Maumus 2022 |
| BtaV03 | Bta02746 | Viridiplantae | / | 50.0 | 47.7 | 636 | 98.1 | 0.83 | BTA017301.1, BTA020608.1 | Ssa06068 | | Plant/T31B5-30 protein | | Gilbert and Maumus 2022 |
| BtaV04 | Bta04871 | Viridiplantae | / | 66.8 | 60.3 | 302 | 86.1 | 0.96 | BTA009091.1 | Ssa10894 | | Nicotianamine synthase, putative | | Gilbert and Maumus 2022 |
| BtaV05 | Bta11576 | Viridiplantae | / | 59.3 | 57.3 | 268 | 96.2 | 0.96 | Scaffold_1194_116759_114636 | scaffold89828_1484_481 | | Sterol desaturase, putative | | Gilbert and Maumus 2022 ; Li et al. 2022 |
| BtaV06 | Bta14387 | Viridiplantae | / | 82.1 | 76.2 | 233 | 95.0 | 0.92 | BTA010079.1 | Ssa02799 , Ssa10266 | | Pathogen-related protein, related | | Gilbert and Maumus 2022 |
| BtaV07 | Bta11842 | Viridiplantae | / | 79.5 | 70.3 | 179 | 74.9 | 0.96 | | | | Transferase, transferring glycosyl groups, putative | | Gilbert and Maumus 2022 ; Li et al. 2022 |
| BtaV08 | Bta02650 | Viridiplantae | / | 42.5 | 39.6 | 455 | 98.5 | 0.59 | BTA005164.2, BTA023005.1 | Ssa02576, Ssa02578 | | Anthocyanin 5-aromatic | | Xia et al. 2021 ; Gilbert and Maumus |
| | Bta07786 | | | 39.7 | 37.5 | 465 | 97.6 | 0.96 | | | | | | Xia et al. 2021 ; Gilbert and Maumus 2022 ; Li et al. 2022 |
| BtaV09 | Bta05651 | Viridiplantae | / | 63.8 | 57.7 | 243 | 97.3 | 0.44 | BTA018079.1, BTA025205.1 | Ssa00364, Ssa03783 Ssa14466 | | HXXXD-type acyl-transferase family | | Gilbert and Maumus 2022 |
| | Bta05662 | | | 66.0 | 55.9 | 148 | 67.4 | 0.82 | | | | | | Gilbert and Maumus 2022 |
| | Bta01560 | | | 57.8 | 53.5 | 465 | 91.8 | 0.8 | | | | | | Gilbert and Maumus 2022 ; Li et al. 2022 |
| | Bta01561 | | | 57.8 | 53.0 | 318 | 93.5 | 0.8 | | | | | | Gilbert and Maumus 2022 |
| BtaV10 | Bta04575 | Viridiplantae | 1 | 59.0 | 54.9 | 174 | 97.5 | 0.88 | BTA022947.1, Scaffold_44_980879_981397 | Ssa09262 | | Ornithine decarboxylase | | Gilbert and Maumus 2022 |
| | Bta05675 | | | 61.8 | 58.9 | 392 | 94.4 | 0.67 | | | | | | Gilbert and Maumus 2022 |
| | Bta05676 | | | 61.4 | 58.6 | 394 | 97.6 | 0.67 | | | | | | Gilbert and Maumus 2022 |
| BtaV11 | Bta08851 | Viridiplantae | / | 73.5 | 69.1 | 401 | 97.6 | 1.0 | BTA001066.1 | Ssa14480 | | Alpha-(1,4)-fucosyltransferase-like protein | GT10 | Gilbert and Maumus 2022 |
| BtaV12 | Bta08104 | Viridiplantae | / | 65.4 | 59.3 | 92 | 71.2 | 0.52 | BTA002345.1 | Ssa14505 | | Subtilisin-like protease | | Gilbert and Maumus 2022 ; Li et al. 2022 |
| BtaV13 | Bta01681 | Viridiplantae | / | 64.5 | 60.3 | 261 | 98.8 | 0.77 | BTA015001.1, BTA022461.1 | Ssa04825, Ssa10246, Ssa12462, Ssa13979 | | S-adenosylmethionine-dependent methyltransferase, putative | | Gilbert and Maumus 2022 ; Li et al. 2022 |
| | Bta13782 | | | 73.6 | 66.3 | 260 | 99.1 | 1.0 | | | | | | Gilbert and Maumus 2022 ; Li et al. 2022 |
| | Bta14307 | | | 69.5 | 62.8 | 222 | 96.4 | 0.96 | | | | | | Gilbert and Maumus 2022 |
| BtaV14 | Bta02345 | Viridiplantae | / | 59.9 | 57.7 | 698 | 97.8 | 0.82 | BTA019832.1 | Ssa01449, Ssa06609 | | Phenylalanine ammonia-lyase | | Li et al. 2022 |
| | Bta09738 | | | 70.3 | 67.5 | 697 | 95.1 | 0.89 | | | | | | Li et al. 2022 |
| BtaV15 | Bta08293 | Viridiplantae | / | 61.9 | 55.4 | 387 | 97.5 | 0.65 | BTA010343.2 | Ssa07619 | | Chalcone synthase | | Gilbert and Maumus 2022 ; Li et al. 2022 |
| BtaV16 | Bta14404 | Viridiplantae | / | 78.1 | 75.6 | 355 | 86.6 | 0.87 | Scaffold_51_1080200_1078090 | Ssa04498 | | Glutelin type-A 1 | | Gilbert and Maumus 2022 ; Li et al. 2022 |
| BtaV17 | Bta07485 | Viridiplantae | / | 67.6 | 64.8 | 365 | 85.5 | 0.81 | BTA008512.1, BTA019386.1, BTA019387.1 | Ssa04263 | | Purple acid phosphatase 29 | | Gilbert and Maumus 2022 ; Li et al. 2022 |
| BtaV18 | Bta05648 | Viridiplantae | / | 68.1 | 64.5 | 120 | 17.1 | 0.56 | Scaffold_1460_66224_62822 | Ssa11172 | | HXXXD-type acyl-transferase family protein | | No |
| BtaV19 | Bta07668 | Viridiplantae | / | 69.6 | 64.4 | 464 | 65.0 | 0.92 | BTA014198.1 | scaffold31037_32915_31767 | | Proteasome-associated ATPase | | Gilbert and Maumus 2022 |
| BtaV20 | Bta04264 | Viridiplantae | / | 52.5 | 49.6 | 370 | 96.0 | 0.88 | BTA015596.1, BTA007356.1, BTA011448.1, BTA015354.2, BTA015355.1, BTA028840.1, BTA011446.1 | Ssa06732, Ssa02331, Ssa02332, Ssa02334, Ssa02342, Ssa03431, Ssa08612, Ssa11969, Ssa15146, Ssa15387, Ssa02337, Ssa02338, Ssa03432 | Tv_08972-RA | Omega-6 fatty acid desaturase | | Gilbert and Maumus 2022 |
| | Bta06451 | | | 52.3 | 51.2 | 366 | 61.1 | 0.45 | | | | | | Gilbert and Maumus 2022 |
| | Bta06452 | | | 49.5 | 47.2 | 364 | 96.1 | 0.39 | | | | | | Gilbert and Maumus 2022 |
| | Bta06453 | | | 50.6 | 47.9 | 364 | 88.4 | 0.49 | | | | | | Gilbert and Maumus 2022 |
| | Bta09295 | | | 76.0 | 73.8 | 383 | 99.4 | 1.0 | | | | | | Gilbert and Maumus 2022 |
| | Bta13659 | | | 62.9 | 61.6 | 393 | 97.3 | 0.87 | | | | | | Gilbert and Maumus 2022 |
| | Bta15754 | | | 57.1 | 53.9 | 212 | 81.3 | 0.1 | | | | | | Gilbert and Maumus 2022 |
| | Bta15755 | | | 65.6 | 63.2 | 380 | 98.9 | 0.2 | | | | | | Gilbert and Maumus 2022 |
| | Bta15756 | | | 60.1 | 58.4 | 362 | 94.3 | 0.2 | | | | | | Gilbert and Maumus 2022 |
| | Bta15757 | | | 57.1 | 55.2 | 363 | 95.9 | 0.2 | | | | | | Gilbert and Maumus 2022 |
| | Bta15759 | | | 55.9 | 52.6 | 166 | 93.4 | 0.2 | | | | | | Gilbert and Maumus 2022 |
| | Bta15760 | | | 54.6 | 52.3 | 145 | 83.8 | 0.2 | | | | | | Gilbert and Maumus 2022 |
| | Bta15762 | | | 51.6 | 49.9 | 368 | 93.5 | 0.16 | | | | | | Gilbert and Maumus 2022 |
| | Bta15765 | | | 65.7 | 63.5 | 363 | 81.8 | 0.16 | | | | | | Gilbert and Maumus 2022 |

| | | | | | | | | | | | | | | |
|---|---|---|---|---|---|---|---|---|---|---|---|---|---|---|
| BtaV21 | Bta14371<br>Bta14372 | Viridiplantae | / | 58.2<br>59.9 | 55.9<br>55.1 | 122<br>488 | 97.5<br>54.3 | 0.39<br>0.45 | Scaffold_688_163109_167894, Scaffold_688_157148_164111 | Ssa04520, Ssa04521, Ssa04522, Ssa04523 | | HXXXD-type acyl-transferase family | | Gilbert and Maumus 2022<br>Gilbert and Maumus 2022 |
| BtaV22 | Bta13094<br>Bta13103 | Viridiplantae | / | 33.0<br>35.6 | 30.7<br>32.9 | 185<br>197 | 95.8<br>69.6 | 0.86<br>0.82 | BTA025985.1, BTA025994.1 | Ssa09189 , scaffold212068_54_492 | TLow_01580-RA, Tv_15540-RA, Tv_15541-RA | rRNA N-glycosidase | | Lapadula et al. 2020<br>Lapadula et al. 2020 ; Gilbert and Maumus 2022 ; Li et al. 2022 |
| BtaV23 | Bta15659 | Viridiplantae | / | 71.8 | 67.7 | 364 | 96.2 | 0.96 | BTA003879.1 | Ssa05099 | | Beta-1,4-N-acetylglucosaminyltransferase family protein | GT17 | Gilbert and Maumus 2022 |
| BtaV24 | Bta14885 | Viridiplantae | / | 60.3 | 55.9 | 507 | 98.1 | 0.94 | BTA018849.1, BTA025653.1 | Ssa09787 | | Beta-1,2-xylosyltransferase | GT61 | Gilbert and Maumus 2022 ; Li et al. 2022 |
| BtaV25 | Bta11221<br>Bta11222 | Viridiplantae | / | 59.4<br>53.5 | 54.8<br>50.8 | 549<br>446 | 66.0<br>86.3 | 0.86<br>0.86 | Scaffold_1904_91816_77676, Scaffold_1904_11 | Ssa12987, scaffold471_39969_63131 | | Pectinesterase | CE8 | Gilbert and Maumus 2022<br>Gilbert and Maumus 2022 |
| BtaV26 | Bta07117 | Viridiplantae | / | 62.0 | 59.1 | 185 | 95.1 | 0.38 | Scaffold_3373_1248_9134 | scaffold168257_14_362 | | Ureidoglycolate hydrolase | | Gilbert and Maumus 2022 ; Li et al. 2022 |
| BtaV27 | Bta15239 | Viridiplantae | / | 69.8 | 64.1 | 119 | 87.6 | 1.0 | BTA026135.1 | Ssa02751 | | Expansin-like EG45 domain-containing protein | | Li et al. 2022 |

*Complex: Bacterial or Fungal donor*

| HGT event | Sequence name | Origin of donor sequences | Alternative topology | Similarities with the donor sequences | | Length and coverage of the alignment with the donor sequences | | Local score for genomic environment | Homologs found in the two other *B. tabaci* cryptic species | | Homologs found in *T. vaporarorium* | Annotation (WGD version 1.2) | CAZy | Described in literature |
|---|---|---|---|---|---|---|---|---|---|---|---|---|---|---|
| | | | | max id | average id | average aln length | average coverage | | *B. tabaci* MED | *B. tabaci* SSA-ECA | | | | |
| BtaC01 | Bta02634 | Bacteria or Fungi | 0 | 34.4 | 31.1 | 350 | 83.9 | 0.6 | BTA008786.1 | Ssa01521 | Tv_05105-RA | Pectin lyase | PL1_4 | No |

**Supplementary Table 3:** HGT candidates described in the literature for *B. tabaci* MEAM1 and not found in our study

| Sequence name | Literature | Reason why it was not found in our study |
|---|---|---|
| Bta00456 | Bacteria (Chen et al. 2016) | No longer found in the 1.2 version of the proteome. |
| Bta02353 | Bacteria (Li et al. 2022) | Identity between the donor sequences and the HGT candidate sequence in the DIAMOND homology search results was less than 30%. |
| Bta02480 | Bacteria (Chen et al. 2016 ; Li et al. 2022) | AHS ≤ 0 |
| Bta07164 | Bacteria (Chen et al. 2016) | AHS ≤ 0 |
| Bta11397 | Bacteria (Chen et al. 2016) | AHS ≤ 0 |
| Bta12968 | Bacteria (Li et al. 2022) | Classified as non-HGT due to the presence of other metazoan sequences in the sister branch and in the ancestral sister branch. The alternative topology constructed by grouping all metazoan sequences was not significantly different from the unconstrained topology, confirming the classification as non-HGT. |
| Bta13226 | Bacteria (Chen et al. 2016) | AHS ≤ 0 |
| Bta13776 | Bacteria (Chen et al. 2016) | AHS ≤ 0 |
| Bta13944 | Bacteria (Li et al. 2022) | AHS ≤ 0 |
| Bta14500 | Bacteria (Chen et al. 2016 ; Li et al. 2022) | AHS ≤ 0 |
| Bta14724 | Bacteria (Li et al. 2022) | AHS ≤ 0 |
| Bta02218 | Fungi (Chen et al. 2016) | No similarity found with NR in the Diamond result. The lack of similarity was confirmed by BLASTP against NR at NCBI. |
| Bta08969 | Fungi (Chen et al. 2016) | HGT_COMPLEX: Unclear whether donor origin was bacterial or fungal. |
| Bta13068 | Viridiplantae (Gilbert and Maumus 2022) | No similarity found with NR in the Diamond result. The lack of similarity was confirmed by BLASTP against NR at NCBI. |

**Supplementary Table 4:** Overrepresented GO terms among validated HGT candidates from potential bacterial, fungal or viridiplantae donors for *B. tabaci* MEAM1

<table>
<tr><th>root_node_name</th><th>node_id</th><th>node_name</th><th>raw_p_overrep</th><th>FWER_overrep</th><th>refined_p_overrep</th><th>FDR_overrep</th><th>nb_genes_refset_root_node</th><th>nb_genes_candidate_in_root</th><th>nb_gene_refset_node</th><th>nb_genes_candidate_in_node</th><th>name_genes_candidate_in_node</th></tr>
<tr><td>HGT Bacteria</td><td></td><td></td><td></td><td></td><td></td><td></td><td></td><td></td><td></td><td></td><td></td></tr>
<tr><td>molecular_function</td><td>GO:0000287</td><td>magnesium ion binding</td><td>4.78709249865742e-08</td><td>0</td><td>4.78709249865742e-08</td><td>1.66776125759678e-07</td><td>6169</td><td>44</td><td>34</td><td>6</td><td>Bta00747|Bta00840|Bta01938|Bta03871|</td></tr>
<tr><td>molecular_function</td><td>GO:0016866</td><td>intramolecular transferase activity</td><td>2.37339959024691e-11</td><td>0</td><td>2.37339959024691e-11</td><td>1.60204472341666e-10</td><td>6169</td><td>44</td><td>21</td><td>7</td><td>Bta02625|Bta07024|Bta07125|Bta07126|Bta07870|Bta11369|Bta11370</td></tr>
<tr><td>molecular_function</td><td>GO:0016879</td><td>ligase activity, forming carbon-nitrogen bonds</td><td>2.27534974684758e-07</td><td>0</td><td>0.0002757252188581</td><td>6.30096852973176e-07</td><td>6169</td><td>44</td><td>23</td><td>5</td><td>Bta00062|Bta00840|Bta01938|Bta05339|Bta05813</td></tr>
<tr><td>molecular_function</td><td>GO:0004170</td><td>dUTP diphosphatase activity</td><td>2.2224899964394e-06</td><td>0.001</td><td>2.2224899964394e-06</td><td>5.33397599145455e-06</td><td>6169</td><td>44</td><td>5</td><td>3</td><td>Bta00747|Bta05687|Bta13975</td></tr>
<tr><td>molecular_function</td><td>GO:0004141</td><td>dethiobiotin synthase activity</td><td>3.77712517891934e-05</td><td>0.013</td><td>3.77712517891934e-05</td><td>5.16366480156062e-05</td><td>6169</td><td>44</td><td>2</td><td>2</td><td>Bta00840|Bta01938</td></tr>
<tr><td>molecular_function</td><td>GO:0008836</td><td>diaminopimelate decarboxylase activity</td><td>3.77712517891934e-05</td><td>0.013</td><td>3.77712517891934e-05</td><td>5.16366480156062e-05</td><td>6169</td><td>44</td><td>2</td><td>2</td><td>Bta03589|Bta03593</td></tr>
<tr><td>cellular_component</td><td>GO:0005811</td><td>lipid droplet</td><td>8.69071715427881e-18</td><td>0</td><td>8.69071715427881e-18</td><td>9.38597452662112e-16</td><td>2247</td><td>10</td><td>8</td><td>7</td><td>Bta02625|Bta07024|Bta07125|Bta07126|</td></tr>
<tr><td>biological_process</td><td>GO:0008610</td><td>lipid biosynthetic process</td><td>2.19831675341128e-12</td><td>0</td><td>0.000189397237390966</td><td>1.8262939182186e-11</td><td>4287</td><td>43</td><td>102</td><td>13</td><td>Bta02625|Bta02653|Bta04469|Bta05264|Bta06818|Bta07024|Bta07125|Bta07126|</td></tr>
<tr><td>biological_process</td><td>GO:0009089</td><td>lysine biosynthetic process via diaminopimelate</td><td>5.97004604987519e-09</td><td>0</td><td>5.97004604987519e-09</td><td>2.47986528225585e-08</td><td>4287</td><td>43</td><td>4</td><td>4</td><td>Bta03589|Bta03593|Bta06657|Bta20020</td></tr>
<tr><td>biological_process</td><td>GO:0009102</td><td>biotin biosynthetic process</td><td>5.97004604987519e-09</td><td>0</td><td>5.97004604987519e-09</td><td>2.47986528225585e-08</td><td>4287</td><td>43</td><td>4</td><td>4</td><td>Bta00840|Bta01937|Bta01938|Bta09725</td></tr>
<tr><td>biological_process</td><td>GO:0016104</td><td>triterpenoid biosynthetic process</td><td>3.12108961795685e-15</td><td>0</td><td>3.12108961795685e-15</td><td>4.81539541056199e-14</td><td>4287</td><td>43</td><td>7</td><td>7</td><td>Bta02625|Bta07024|Bta07125|Bta07126|Bta07870|Bta11369|Bta11370</td></tr>
<tr><td>biological_process</td><td>GO:0006226</td><td>dUMP biosynthetic process</td><td>6.95241478593945e-06</td><td>0.002</td><td>6.95241478593945e-06</td><td>1.31729964365169e-05</td><td>4287</td><td>43</td><td>5</td><td>3</td><td>Bta00747|Bta05687|Bta13975</td></tr>
<tr><td>biological_process</td><td>GO:0046081</td><td>dUTP catabolic process</td><td>6.95241478593945e-06</td><td>0.002</td><td>6.95241478593945e-06</td><td>1.31729964365169e-05</td><td>4287</td><td>43</td><td>5</td><td>3</td><td>Bta00747|Bta05687|Bta13975</td></tr>
<tr><td>biological_process</td><td>GO:0006526</td><td>arginine biosynthetic process</td><td>8.10478963448566e-05</td><td>0.029</td><td>8.10478963448566e-05</td><td>0.000104204438157673</td><td>4287</td><td>43</td><td>2</td><td>2</td><td>Bta00062|Bta00063</td></tr>
<tr><td>HGT Fungi</td><td></td><td></td><td></td><td></td><td></td><td></td><td></td><td></td><td></td><td></td><td></td></tr>
<tr><td>molecular_function</td><td>GO:0004601</td><td>peroxidase activity</td><td>3.66430096659827e-31</td><td>0</td><td>3.66430096659827e-31</td><td>2.19858057995896e-30</td><td>6169</td><td>45</td><td>49</td><td>19</td><td>Bta00038|Bta00039|Bta00041|Bta00658|Bta00659|Bta00660|Bta01073|Bta02217|Bta04808|Bta05684|Bta05753|Bta05754|</td></tr>
<tr><td>HGT Viridiplantae</td><td></td><td></td><td></td><td></td><td></td><td></td><td></td><td></td><td></td><td></td><td></td></tr>
<tr><td>molecular_function</td><td>GO:0016717</td><td>oxidoreductase activity, acting on paired donors, with oxidation of a pair of donors resulting in the reduction of molecular oxygen to two molecules of water</td><td>1.26787439093126e-10</td><td>0</td><td>1.26787439093126e-10</td><td>2.09199274503657e-09</td><td>6169</td><td>29</td><td>20</td><td>6</td><td>Bta06452|Bta09295|Bta13659|Bta15755|Bta15756|Bta15765</td></tr>
<tr><td>molecular_function</td><td>GO:0030598</td><td>rRNA N-glycosylase activity</td><td>1.73683874506334e-05</td><td>0.003</td><td>1.73683874506334e-05</td><td>4.77630654892417e-05</td><td>6169</td><td>29</td><td>2</td><td>2</td><td>Bta13094|Bta13103</td></tr>
<tr><td>molecular_function</td><td>GO:0030599</td><td>pectinesterase activity</td><td>1.73683874506334e-05</td><td>0.003</td><td>1.73683874506334e-05</td><td>4.77630654892417e-05</td><td>6169</td><td>29</td><td>2</td><td>2</td><td>Bta11221|Bta11222</td></tr>
<tr><td>molecular_function</td><td>GO:0016841</td><td>ammonia-lyase activity</td><td>5.19677076179776e-05</td><td>0.005</td><td>5.19677076179776e-05</td><td>0.000107183396962079</td><td>6169</td><td>29</td><td>3</td><td>2</td><td>Bta02345|Bta09738</td></tr>
<tr><td>biological_process</td><td>GO:0006596</td><td>polyamine biosynthetic process</td><td>4.5738072745559e-06</td><td>0</td><td>4.5738072745559e-06</td><td>1.67706266733716e-05</td><td>4408</td><td>26</td><td>6</td><td>3</td><td>Bta04575|Bta05675|Bta05676</td></tr>
<tr><td>biological_process</td><td>GO:0006629</td><td>lipid metabolic process</td><td>1.99651433804412e-12</td><td>0</td><td>1.99651433804412e-12</td><td>6.58849731554559e-11</td><td>4408</td><td>26</td><td>240</td><td>15</td><td>Bta04264|Bta06451|Bta06452|Bta06453|Bta09295|Bta11576|Bta13659|Bta15754|</td></tr>
<tr><td>biological_process</td><td>GO:0042545</td><td>cell wall modification</td><td>3.90430065448644e-05</td><td>0.011</td><td>3.90430065448644e-05</td><td>8.58946143987016e-05</td><td>4408</td><td>26</td><td>2</td><td>2</td><td>Bta11221|Bta11222</td></tr>
</table>

| HGT Bacteria + Fungi + Viridiplantae | | | | | | | | | | | |
|---|---|---|---|---|---|---|---|---|---|---|---|
| molecular_function | GO:0004601 | peroxidase activity | 5.45718553136594e-22 | 0 | 5.45718553136594e-22 | 3.41074095710371e-20 | 6673 | 118 | 49 | 19 | Bta00038\|Bta00039\|Bta00041\|Bta00658\|Bta00659\|Bta00660\|Bta01073\|Bta02217\|Bta04808\|Bta05684\|Bta05753\|Bta05754\|Bta05981\|Bta07721\|Bta08225\|Bta08226\|Bta11373\|Bta15601\|Bta15602 |
| molecular_function | GO:0016717 | oxidoreductase activity, acting on paired donors, with oxidation of a pair of donors resulting in the reduction of molecular oxygen to two molecules of water | 6.3644593518552e-07 | 0 | 6.3644593518552e-07 | 2.20988171939417e-06 | 6673 | 118 | 20 | 6 | Bta06452\|Bta09295\|Bta13659\|Bta15755\|Bta15756\|Bta15765 |
| molecular_function | GO:0016866 | intramolecular transferase activity | 3.04349973779946e-08 | 0 | 3.04349973779946e-08 | 1.40902765638864e-07 | 6673 | 118 | 21 | 7 | Bta02625\|Bta07024\|Bta07125\|Bta07126\|Bta07870\|Bta11369\|Bta11370 |
| molecular_function | GO:0016841 | ammonia-lyase activity | 4.63518691969453e-06 | 0.001 | 4.63518691969453e-06 | 1.25956166296047e-05 | 6673 | 118 | 3 | 3 | Bta02345\|Bta08776\|Bta09738 |
| molecular_function | GO:0000287 | magnesium ion binding | 1.82189865098747e-05 | 0.007 | 1.82189865098747e-05 | 4.46543787006732e-05 | 6673 | 118 | 34 | 6 | Bta00747\|Bta00840\|Bta01938\|Bta03871\|Bta05687\|Bta13975 |
| molecular_function | GO:0016798 | hydrolase activity, acting on glycosyl bonds | 1.95323653630582e-05 | 0.007 | 1.95323653630582e-05 | 4.69528013535052e-05 | 6673 | 118 | 91 | 9 | Bta00773\|Bta02250\|Bta02658\|Bta02866\|Bta04549\|Bta06115\|Bta06118\|Bta13094\|Bta13103 |
| molecular_function | GO:0016879 | ligase activity, forming carbon-nitrogen bonds | 3.26094908050568e-05 | 0.007 | 3.26094908050568e-05 | 7.02790750108982e-05 | 6673 | 118 | 23 | 5 | Bta00062\|Bta00840\|Bta01938\|Bta05339\|Bta05813 |
| molecular_function | GO:0004170 | dUTP diphosphatase activity | 4.52193208416261e-05 | 0.008 | 4.52193208416261e-05 | 8.07487872171894e-05 | 6673 | 118 | 5 | 3 | Bta00747\|Bta05687\|Bta13975 |
| cellular_component | GO:0005811 | lipid droplet | 4.6521100294467e-16 | 0 | 4.6521100294467e-16 | 9.69189589468063e-15 | 1834 | 9 | 8 | 7 | Bta02625\|Bta07024\|Bta07125\|Bta07126\|Bta07870\|Bta11369\|Bta11370 |
| biological_process | GO:0006629 | lipid metabolic process | 1.03172125882258e-19 | 0 | 4.11302896236752e-09 | 2.57930314705646e-18 | 4643 | 77 | 240 | 30 | Bta02625\|Bta02653\|Bta02658\|Bta04264\|Bta04469\|Bta05264\|Bta06451\|Bta06452\|Bta06453\|Bta06818\|Bta07024\|Bta07125\|Bta07126\|Bta07870\|Bta09295\|Bta11043\|Bta11369\|Bta11370\|Bta11576\|Bta11911\|Bta11912\|Bta13659\|Bta15754\|Bta15755\|Bta15756\|Bta15757\|Bta15759\|Bta15760\|Bta15762\|Bta15765 |
| biological_process | GO:0008610 | lipid biosynthetic process | 5.22715083728764e-11 | 0 | 0.000147337329446341 | 4.27575861363102e-10 | 4643 | 77 | 102 | 15 | Bta02625\|Bta02653\|Bta04469\|Bta05264\|Bta06818\|Bta07024\|Bta07125\|Bta07126\|Bta07870\|Bta11043\|Bta11369\|Bta11370\|Bta11576\|Bta11911\|Bta11912 |
| biological_process | GO:0009089 | lysine biosynthetic process via diaminopimelate | 6.90043184238973e-08 | 0 | 6.90043184238973e-08 | 2.61379994029914e-07 | 4643 | 77 | 4 | 4 | Bta03589\|Bta03593\|Bta06657\|Bta20020 |
| biological_process | GO:0009102 | biotin biosynthetic process | 6.90043184238973e-08 | 0 | 6.90043184238973e-08 | 2.61379994029914e-07 | 4643 | 77 | 4 | 4 | Bta00840\|Bta01937\|Bta01938\|Bta09725 |
| biological_process | GO:0016053 | organic acid biosynthetic process | 5.98680141923098e-14 | 0 | 0.000238330962574668 | 9.35437721754841e-13 | 4643 | 77 | 43 | 13 | Bta00062\|Bta00063\|Bta00840\|Bta01937\|Bta01938\|Bta03589\|Bta03593\|Bta04871\|Bta05339\|Bta06657\|Bta09725\|Bta20014\|Bta20020 |
| biological_process | GO:0016104 | triterpenoid biosynthetic process | 2.55883320124494e-13 | 0 | 2.55883320124494e-13 | 2.66545125129681e-12 | 4643 | 77 | 7 | 7 | Bta02625\|Bta07024\|Bta07125\|Bta07126\|Bta07870\|Bta11369\|Bta11370 |
| biological_process | GO:0019752 | carboxylic acid metabolic process | 7.50404068388319e-11 | 0 | 0.000211410856127128 | 5.21113936380777e-10 | 4643 | 77 | 164 | 18 | Bta00062\|Bta00063\|Bta00840\|Bta01937\|Bta01938\|Bta03200\|Bta03589\|Bta03593\|Bta04871\|Bta05339\|Bta06657\|Bta06820\|Bta08776\|Bta09725\|Bta09738\|Bta15103\|Bta20014\|Bta20020 |

| | | | | | | | | | | |
|---|---|---|---|---|---|---|---|---|---|---|
| biological_process | GO:0046394 | carboxylic acid biosynthetic process | 5.98680141923098e-14 | 0 | 0.000238330962574668 | 9.35437721754841e-13 | 4643 | 77 | 43 | 13 | Bta00062\|Bta00063\|Bta00840\|Bta01937\|Bta01938\|Bta03589\|Bta03593\|Bta04871\|Bta05339\|Bta06657\|Bta09725\|Bta20014\|Bta20020 |
| biological_process | GO:0006226 | dUMP biosynthetic process | 4.23797470126717e-05 | 0.01 | 4.23797470126717e-05 | 7.67749040084632e-05 | 4643 | 77 | 5 | 3 | Bta00747\|Bta05687\|Bta13975 |
| biological_process | GO:0046081 | dUTP catabolic process | 4.23797470126717e-05 | 0.01 | 4.23797470126717e-05 | 7.67749040084632e-05 | 4643 | 77 | 5 | 3 | Bta00747\|Bta05687\|Bta13975 |
| biological_process | GO:0006596 | polyamine biosynthetic process | 8.37502425309647e-05 | 0.021 | 8.37502425309647e-05 | 0.000134215132261161 | 4643 | 77 | 6 | 3 | Bta04575\|Bta05675\|Bta05676 |

**Supplementary Table 5:** Predicted CAZymes in *B. tabaci* , *T. vaporariorum* , *F. occidentalis* and *T. palmi*

*B. tabaci*

| Protein ID | Description | Model notes | Definition line | HGT | Donor |
|---|---|---|---|---|---|
| Bta00067 | AA3_2 | | GMC oxidoreductase | | |
| Bta00205 | GT105 | fragment N-term; | Glycosyltransferase Family 105 protein | | |
| Bta00245 | GH18-CBM18-CBM18-CBM18 | | Glycoside Hydrolase Family 18 / Carbohydrate-Binding Module Family 18 protein | | |
| Bta00300 | GT1 | | Glycosyltransferase Family 1 protein | | |
| Bta00304 | CBM21 | | Carbohydrate-Binding Module Family 21 protein | | |
| Bta00306 | AA1 | | Multicopper oxidase | | |
| Bta00307 | GT16 | | Glycosyltransferase Family 16 protein | | |
| Bta00329 | GH37 | | Glycoside Hydrolase Family 37 protein | | |
| Bta00445 | GT27-CBM13 | | Glycosyltransferase Family 27 / Carbohydrate-Binding Module Family 13 protein | | |
| Bta00548 | GT1 | | Glycosyltransferase Family 1 protein | | |
| Bta00633 | GT66 | fragment C-term; | Glycosyltransferase Family 66 protein | | |
| Bta00718 | GT1 | | Glycosyltransferase Family 1 protein | | |
| Bta00731 | GT27-CBM13 | | Glycosyltransferase Family 27 / Carbohydrate-Binding Module Family 13 protein | | |
| Bta00773 | GH32 | | Glycoside Hydrolase Family 32 protein | HGT | Bacteria |
| Bta00822 | GT1 | | Glycosyltransferase Family 1 protein | | |
| Bta00951 | GT23 | (fragment) splicing problem; | Glycosyltransferase Family 23 protein | | |
| Bta00966 | GT31 | | Glycosyltransferase Family 31 protein | | |
| Bta00998 | GH89 | (fragment) splicing problem; | Glycoside Hydrolase Family 89 protein | | |
| Bta01019 | GT31 | | Glycosyltransferase Family 31 protein | | |
| Bta01062 | GT47-GT64 | | Glycosyltransferase Family 47 / Glycosyltransferase Family 64 protein | | |
| Bta01072 | CBM32-AA5_2 | | Carbohydrate-Binding Module Family 32 / Galactose oxidase | HGT | Fungi |
| Bta01126 | GH79 | fragment N-term; | Glycoside Hydrolase Family 79 protein | | |
| Bta01138 | GT1 | fragment C-term; | Glycosyltransferase Family 1 protein | | |
| Bta01168 | GT4 | | Glycosyltransferase Family 4 protein | | |
| Bta01169 | GT8 | | Glycosyltransferase Family 8 protein | | |
| Bta01181 | GT1 | fragment N-term; | Glycosyltransferase Family 1 protein | | |
| Bta01182 | GT1-GT1 | | Glycosyltransferase Family 1 protein | | |
| Bta01183 | GT1 | | Glycosyltransferase Family 1 protein | | |
| Bta01184 | GT1 | | Glycosyltransferase Family 1 protein | | |
| Bta01219 | GH47 | | Glycoside Hydrolase Family 47 protein | | |
| Bta01304 | GT1 | | Glycosyltransferase Family 1 protein | | |
| Bta01342 | GH18-GH18-CBM14 | | Glycoside Hydrolase Family 18 / Carbohydrate-Binding Module Family 14 protein | | |
| Bta01403 | GT27-CBM13 | | Glycosyltransferase Family 27 / Carbohydrate-Binding Module Family 13 protein | | |
| Bta01405 | GH47 | | Glycoside Hydrolase Family 47 protein | | |
| Bta01463 | GH13_17 | fragment N-term; | Glycoside Hydrolase Family 13 protein | | |
| Bta01464 | GH13 | fragment N-term / C-term; | Glycoside Hydrolase Family 13 protein | | |
| Bta01468 | GH63 | | Glycoside Hydrolase Family 63 protein | | |
| Bta01478 | GH13_17 | | Glycoside Hydrolase Family 13 protein | | |
| Bta01485 | GH2 | | Glycoside Hydrolase Family 2 protein | | |
| Bta01758 | GT31 | (fragment) splicing problem; | Glycosyltransferase Family 31 protein | | |
| Bta01766 | GT1 | fragment N-term; | Glycosyltransferase Family 1 protein | | |
| Bta01777 | CBM47 | | Carbohydrate-Binding Module Family 47 protein | | |
| Bta01963 | GT68 | | Glycosyltransferase Family 68 protein | | |
| Bta01975 | CBM14-CBM14-CBM14 | | Carbohydrate-Binding Module Family 14 protein | | |

| | | | | | |
|---|---|---|---|---|---|
| Bta01976 | CBM14-CBM14-CBM14 | | Carbohydrate-Binding Module Family 14 protein | | |
| Bta01977 | CBM14-CBM14-CBM14 | | Carbohydrate-Binding Module Family 14 protein | | |
| Bta01978 | CBM14-CBM14-CBM14 | | Carbohydrate-Binding Module Family 14 protein | | |
| Bta01979 | CBM14-CBM14-CBM14 | | Carbohydrate-Binding Module Family 14 protein | | |
| Bta01980 | CBM14-CBM14-CBM14 | | Carbohydrate-Binding Module Family 14 protein | | |
| Bta01982 | CBM14-CBM14-CBM14 | (fragment) splicing problem; | Carbohydrate-Binding Module Family 14 protein | | |
| Bta02118 | GH30_1 | | Glycoside Hydrolase Family 30 protein | | |
| Bta02176 | GH84 | | Glycoside Hydrolase Family 84 protein | | |
| Bta02228 | GT1 | | Glycosyltransferase Family 1 protein | | |
| Bta02242 | GT2 | | Glycosyltransferase Family 2 protein | | |
| Bta02250 | GH49 | | Glycoside Hydrolase Family 49 protein | HGT | Fungi |
| Bta02273 | GH18 | | Glycoside Hydrolase Family 18 protein | | |
| Bta02274 | GH18 | | Glycoside Hydrolase Family 18 protein | | |
| Bta02288 | GT41 | | Glycosyltransferase Family 41 protein | | |
| Bta02306 | GT1 | | Glycosyltransferase Family 1 protein | | |
| Bta02326 | GT1 | | Glycosyltransferase Family 1 protein | | |
| Bta02348 | GT1 | | Glycosyltransferase Family 1 protein | | |
| Bta02350 | GT32 | | Glycosyltransferase Family 32 protein | | |
| Bta02395 | AA3_2 | | GMC oxidoreductase | | |
| Bta02396 | AA3_2 | (fragment) splicing problem; | GMC oxidoreductase | | |
| Bta02397 | AA3_2 | fragment N-term; | GMC oxidoreductase | | |
| Bta02398 | AA3_2 | | GMC oxidoreductase | | |
| Bta02399 | AA3_2 | (fragment) splicing problem; | GMC oxidoreductase | | |
| Bta02400 | AA3_2 | | GMC oxidoreductase | | |
| Bta02401 | AA3_2 | fragment C-term; | GMC oxidoreductase | | |
| Bta02402 | AA3_2 | fragment N-term; | GMC oxidoreductase | | |
| Bta02405 | AA3_2-AA3_2 | (fragment) splicing problem; | GMC oxidoreductase | | |
| Bta02406 | AA3_2 | | GMC oxidoreductase | | |
| Bta02407 | AA3_2 | | GMC oxidoreductase | | |
| Bta02440 | AA3_2 | | GMC oxidoreductase | | |
| Bta02521 | CBM14 | | Carbohydrate-Binding Module Family 14 protein | | |
| Bta02522 | GH37 | | Glycoside Hydrolase Family 37 protein | | |
| Bta02601 | GT1 | | Glycosyltransferase Family 1 protein | | |
| Bta02602 | GT1 | | Glycosyltransferase Family 1 protein | | |
| Bta02603 | GT1 | | Glycosyltransferase Family 1 protein | | |
| Bta02604 | GT1 | | Glycosyltransferase Family 1 protein | | |
| Bta02622 | CBM47 | fragment N-term; | Carbohydrate-Binding Module Family 47 protein | | |
| Bta02630 | GH13_15 | | Glycoside Hydrolase Family 13 protein | | |
| Bta02634 | PL1_4 | fragment C-term; | Polysaccharide Lyase Family 1 protein | HGT | Complex: Bacteria or Fungi |
| Bta02658 | GH30_3 | | Glycoside Hydrolase Family 30 protein | HGT | Fungi |
| Bta02682 | GH116 | | Glycoside Hydrolase Family 116 protein | | |
| Bta02725 | GT1 | | Glycosyltransferase Family 1 protein | | |
| Bta02726 | GT1 | | Glycosyltransferase Family 1 protein | | |
| Bta02795 | GT7 | | Glycosyltransferase Family 7 protein | | |
| Bta02810 | GH13_17 | | Glycoside Hydrolase Family 13 protein | | |
| Bta02866 | GH71 | | Glycoside Hydrolase Family 71 protein | HGT | Fungi |
| Bta02947 | AA3_2 | fragment C-term; | GMC oxidoreductase | | |
| Bta02948 | AA3_2 | fragment N-term; | GMC oxidoreductase | | |
| Bta02973 | AA3_2 | | GMC oxidoreductase | | |

| | | | | | |
|---|---|---|---|---|---|
| Bta02974 | AA3_2 | | GMC oxidoreductase | | |
| Bta02991 | GT1 | | Glycosyltransferase Family 1 protein | | |
| Bta03001 | AA3_2 | | GMC oxidoreductase | | |
| Bta03112 | GH13_17 | fragment N-term; | Glycoside Hydrolase Family 13 protein | | |
| Bta03142 | GH22 | | Glycoside Hydrolase Family 22 protein | | |
| Bta03157 | GT59 | | Glycosyltransferase Family 59 protein | | |
| Bta03192 | GT1 | | Glycosyltransferase Family 1 protein | | |
| Bta03195 | CBM14 | | Carbohydrate-Binding Module Family 14 protein | | |
| Bta03197 | GH37 | | Glycoside Hydrolase Family 37 protein | | |
| Bta03198 | GH37 | | Glycoside Hydrolase Family 37 protein | | |
| Bta03202 | GH37 | | Glycoside Hydrolase Family 37 protein | | |
| Bta03221 | GH38 | | Glycoside Hydrolase Family 38 protein | | |
| Bta03307 | GH13_17 | | Glycoside Hydrolase Family 13 protein | | |
| Bta03340 | GH47 | | Glycoside Hydrolase Family 47 protein | | |
| Bta03431 | GT1 | | Glycosyltransferase Family 1 protein | | |
| Bta03439 | GH13_17 | | Glycoside Hydrolase Family 13 protein | | |
| Bta03440 | GT1 | fragment N-term; | Glycosyltransferase Family 1 protein | | |
| Bta03567 | CBM32-AA5_2 | | Carbohydrate-Binding Module Family 32 / Galactose oxidase | HGT | Fungi |
| Bta03574 | GT13 | | Glycosyltransferase Family 13 protein | | |
| Bta03601 | GT2 | | Glycosyltransferase Family 2 protein | | |
| Bta03753 | CBM50 | | Carbohydrate-Binding Module Family 50 protein | | |
| Bta03818 | GH13_17 | | Glycoside Hydrolase Family 13 protein | | |
| Bta03820 | GT105 | fragment C-term; | Glycosyltransferase Family 105 protein | | |
| Bta03913 | GT49 | | Glycosyltransferase Family 49 protein | | |
| Bta03920 | GH38 | | Glycoside Hydrolase Family 38 protein | | |
| Bta03960 | GT4 | | Glycosyltransferase Family 4 protein | | |
| Bta03991 | GH13_17 | | Glycoside Hydrolase Family 13 protein | | |
| Bta03992 | GH13_17 | | Glycoside Hydrolase Family 13 protein | | |
| Bta03994 | GH13_17 | | Glycoside Hydrolase Family 13 protein | | |
| Bta03995 | GH13_17 | | Glycoside Hydrolase Family 13 protein | | |
| Bta04042 | GH13_17 | fragment C-term; | Glycoside Hydrolase Family 13 protein | | |
| Bta04297 | GH13_17 | | Glycoside Hydrolase Family 13 protein | | |
| Bta04298 | GH13_17 | | Glycoside Hydrolase Family 13 protein | | |
| Bta04306 | GH13_17 | | Glycoside Hydrolase Family 13 protein | | |
| Bta04331 | GH99 | fragment N-term; | Glycoside Hydrolase Family 99 protein | | |
| Bta04378 | GT65 | | Glycosyltransferase Family 65 protein | | |
| Bta04382 | GH13_17 | fragment C-term; | Glycoside Hydrolase Family 13 protein | | |
| Bta04384 | GH13 | fragment N-term / C-term; | Glycoside Hydrolase Family 13 protein | | |
| Bta04402 | AA3_2 | | GMC oxidoreductase | | |
| Bta04421 | GH13_17 | | Glycoside Hydrolase Family 13 protein | | |
| Bta04425 | GH20 | | Glycoside Hydrolase Family 20 protein | | |
| Bta04444 | GT13 | | Glycosyltransferase Family 13 protein | | |
| Bta04537 | GT1 | fragment N-term; | Glycosyltransferase Family 1 protein | | |
| Bta04539 | GT1 | fragment N-term; | Glycosyltransferase Family 1 protein | | |
| Bta04543 | GT1 | fragment C-term; | Glycosyltransferase Family 1 protein | | |
| Bta04549 | GH32 | | Glycoside Hydrolase Family 32 protein | HGT | Bacteria |
| Bta04553 | GH13_15 | | Glycoside Hydrolase Family 13 protein | | |
| Bta04680 | GT54 | | Glycosyltransferase Family 54 protein | | |
| Bta04683 | GH13 | fragment C-term; | Glycoside Hydrolase Family 13 protein | | |

| | | | | | |
|---|---|---|---|---|---|
| Bta04691 | GT25 | fragment C-term; | Glycosyltransferase Family 25 protein | | |
| Bta04694 | GT54 | | Glycosyltransferase Family 54 protein | | |
| Bta04731 | GH13_15 | fragment N-term; | Glycoside Hydrolase Family 13 protein | | |
| Bta04732 | GH13 | fragment C-term; | Glycoside Hydrolase Family 13 protein | | |
| Bta04739 | GT1 | | Glycosyltransferase Family 1 protein | | |
| Bta05024 | GT31 | | Glycosyltransferase Family 31 protein | | |
| Bta05026 | GT27-CBM13 | | Glycosyltransferase Family 27 / Carbohydrate-Binding Module Family 13 protein | | |
| Bta05090 | CBM32-AA5_2 | | Carbohydrate-Binding Module Family 32 / Galactose oxidase | HGT | Fungi |
| Bta05091 | CBM32-AA5_2 | | Carbohydrate-Binding Module Family 32 / Galactose oxidase | HGT | Fungi |
| Bta05092 | AA5_2 | fragment N-term; | Galactose oxidase | HGT | Fungi |
| Bta05093 | CBM32-AA5_2 | | Carbohydrate-Binding Module Family 32 / Galactose oxidase | HGT | Fungi |
| Bta05237 | GH38 | | Glycoside Hydrolase Family 38 protein | | |
| Bta05251 | GT8 | fragment N-term / C-term; | Glycosyltransferase Family 8 protein | | |
| Bta05309 | GH35 | | Glycoside Hydrolase Family 35 protein | | |
| Bta05340 | GH13_17 | | Glycoside Hydrolase Family 13 protein | | |
| Bta05342 | GH13 | fragment C-term; | Glycoside Hydrolase Family 13 protein | | |
| Bta05343 | GH13_17 | fragment N-term; | Glycoside Hydrolase Family 13 protein | | |
| Bta05386 | GH13_17 | | Glycoside Hydrolase Family 13 protein | | |
| Bta05396 | GH13_17 | | Glycoside Hydrolase Family 13 protein | | |
| Bta05397 | GH13_17 | | Glycoside Hydrolase Family 13 protein | | |
| Bta05466 | GT105 | (fragment) splicing problem; | Glycosyltransferase Family 105 protein | | |
| Bta05518 | GH13_17 | | Glycoside Hydrolase Family 13 protein | | |
| Bta05547 | CBM14 | | Carbohydrate-Binding Module Family 14 protein | | |
| Bta05577 | GT35 | | Glycosyltransferase Family 35 protein | | |
| Bta05604 | GT1 | | Glycosyltransferase Family 1 protein | | |
| Bta05616 | GT27-CBM13 | | Glycosyltransferase Family 27 / Carbohydrate-Binding Module Family 13 protein | | |
| Bta05704 | GH47 | | Glycoside Hydrolase Family 47 protein | | |
| Bta05915 | CBM50 | | Carbohydrate-Binding Module Family 50 protein | | |
| Bta05951 | GT1 | | Glycosyltransferase Family 1 protein | | |
| Bta06017 | GT1 | (fragment) splicing problem; | Glycosyltransferase Family 1 protein | | |
| Bta06059 | GH13_17 | fragment N-term; | Glycoside Hydrolase Family 13 protein | | |
| Bta06072 | CBM14 | | Carbohydrate-Binding Module Family 14 protein | | |
| Bta06073 | CBM14 | (fragment) splicing problem; | Carbohydrate-Binding Module Family 14 protein | | |
| Bta06115 | GH17 | | Glycoside Hydrolase Family 17 protein | HGT | Viridiplantae |
| Bta06118 | GH17 | | Glycoside Hydrolase Family 17 protein | HGT | Viridiplantae |
| Bta06208 | GT92 | | Glycosyltransferase Family 92 protein | | |
| Bta06327 | GH13_17 | (fragment) splicing problem; | Glycoside Hydrolase Family 13 protein | | |
| Bta06329 | GH18 | | Glycoside Hydrolase Family 18 protein | | |
| Bta06441 | GH13 | fragment C-term; | Glycoside Hydrolase Family 13 protein | | |
| Bta06445 | GH13_17 | (fragment) splicing problem; | Glycoside Hydrolase Family 13 protein | | |
| Bta06458 | GH13_17 | | Glycoside Hydrolase Family 13 protein | | |
| Bta06497 | GT90 | (fragment) splicing problem; | Glycosyltransferase Family 90 protein | | |
| Bta06513 | GT31 | | Glycosyltransferase Family 31 protein | | |
| Bta06581 | CBM21 | | Carbohydrate-Binding Module Family 21 protein | | |
| Bta06664 | GT1 | | Glycosyltransferase Family 1 protein | | |
| Bta06665 | GT1 | | Glycosyltransferase Family 1 protein | | |
| Bta06706 | GH20 | (fragment) splicing problem; | Glycoside Hydrolase Family 20 protein | | |
| Bta06849 | GH31 | | Glycoside Hydrolase Family 31 protein | | |
| Bta06905 | GT43 | | Glycosyltransferase Family 43 protein | | |

| | | | | | |
|---|---|---|---|---|---|
| Bta06986 | GH20 | | Glycoside Hydrolase Family 20 protein | | |
| Bta06988 | GT22 | | Glycosyltransferase Family 22 protein | | |
| Bta07185 | GT1 | | Glycosyltransferase Family 1 protein | | |
| Bta07226 | GT22 | | Glycosyltransferase Family 22 protein | | |
| Bta07244 | GT31 | | Glycosyltransferase Family 31 protein | | |
| Bta07272 | AA15 | | Auxilliary Activities Family 15 protein | | |
| Bta07330 | AA3_2 | fragment N-term; | GMC oxidoreductase | | |
| Bta07340 | GT61 | | Glycosyltransferase Family 61 protein | | |
| Bta07348 | GT1 | | Glycosyltransferase Family 1 protein | | |
| Bta07368 | GT1 | | Glycosyltransferase Family 1 protein | | |
| Bta07377 | GH13 | (fragment) splicing problem; | Glycoside Hydrolase Family 13 protein | | |
| Bta07385 | GT31 | | Glycosyltransferase Family 31 protein | | |
| Bta07386 | GT31 | fragment N-term; | Glycosyltransferase Family 31 protein | | |
| Bta07432 | GT105 | fragment N-term; | Glycosyltransferase Family 105 protein | | |
| Bta07452 | GH13_17 | (fragment) splicing problem; | Glycoside Hydrolase Family 13 protein | | |
| Bta07453 | GH13_17 | | Glycoside Hydrolase Family 13 protein | | |
| Bta07478 | GT22 | | Glycosyltransferase Family 22 protein | | |
| Bta07492 | GT110 | | Glycosyltransferase Family 110 protein | | |
| Bta07496 | GT13 | | Glycosyltransferase Family 13 protein | | |
| Bta07516 | GH38 | | Glycoside Hydrolase Family 38 protein | | |
| Bta07589 | GT1 | fragment N-term; | Glycosyltransferase Family 1 protein | | |
| Bta07602 | GT1 | | Glycosyltransferase Family 1 protein | | |
| Bta07603 | GT1 | (fragment) splicing problem; | Glycosyltransferase Family 1 protein | | |
| Bta07604 | GT1 | fragment N-term; | Glycosyltransferase Family 1 protein | | |
| Bta07605 | GT1 | | Glycosyltransferase Family 1 protein | | |
| Bta07606 | GT1-GT1 | (fragment) splicing problem; | Glycosyltransferase Family 1 protein | | |
| Bta07607 | GT1 | fragment C-term; | Glycosyltransferase Family 1 protein | | |
| Bta07608 | GT1 | | Glycosyltransferase Family 1 protein | | |
| Bta07610 | GT1 | fragment N-term; | Glycosyltransferase Family 1 protein | | |
| Bta07623 | GT1 | | Glycosyltransferase Family 1 protein | | |
| Bta07646 | GT1 | fragment N-term; | Glycosyltransferase Family 1 protein | | |
| Bta07691 | CBM32-AA5_2 | | Carbohydrate-Binding Module Family 32 / Galactose oxidase | HGT | Fungi |
| Bta07704 | GT1 | | Glycosyltransferase Family 1 protein | | |
| Bta07715 | GT47-GT64 | | Glycosyltransferase Family 47 / Glycosyltransferase Family 64 protein | | |
| Bta07764 | GH13_17 | | Glycoside Hydrolase Family 13 protein | | |
| Bta08022 | CBM14 | | Carbohydrate-Binding Module Family 14 protein | | |
| Bta08033 | CBM14 | | Carbohydrate-Binding Module Family 14 protein | | |
| Bta08063 | GT7 | | Glycosyltransferase Family 7 protein | | |
| Bta08065 | GT4 | | Glycosyltransferase Family 4 protein | | |
| Bta08213 | GT1 | | Glycosyltransferase Family 1 protein | | |
| Bta08294 | GT1-GT1 | | Glycosyltransferase Family 1 protein | | |
| Bta08372 | GT1 | | Glycosyltransferase Family 1 protein | | |
| Bta08425 | GH13_17 | | Glycoside Hydrolase Family 13 protein | | |
| Bta08426 | GH13_17 | fragment C-term; | Glycoside Hydrolase Family 13 protein | | |
| Bta08427 | GH13_17 | (fragment) splicing problem; | Glycoside Hydrolase Family 13 protein | | |
| Bta08431 | GH13_17 | fragment C-term; | Glycoside Hydrolase Family 13 protein | | |
| Bta08458 | GT20 | (fragment) splicing problem; | Glycosyltransferase Family 20 protein | | |
| Bta08517 | GT8-GT49 | fragment N-term; | Glycosyltransferase Family 8 / Glycosyltransferase Family 49 protein | | |
| Bta08621 | GH27 | | Glycoside Hydrolase Family 27 protein | | |

| | | | | | |
|---|---|---|---|---|---|
| Bta08702 | GT20 | | Glycosyltransferase Family 20 protein | | |
| Bta08716 | CBM20 | | Carbohydrate-Binding Module Family 20 protein | | |
| Bta08743 | GT1 | | Glycosyltransferase Family 1 protein | | |
| Bta08748 | GT27-CBM13 | fragment C-term; | Glycosyltransferase Family 27 / Carbohydrate-Binding Module Family 13 protein | | |
| Bta08851 | GT10 | | Glycosyltransferase Family 10 protein | HGT | Viridiplantae |
| Bta08909 | AA3_2 | | GMC oxidoreductase | | |
| Bta08917 | GH20 | | Glycoside Hydrolase Family 20 protein | | |
| Bta08970 | GH18 | fragment C-term; | Glycoside Hydrolase Family 18 protein | | |
| Bta09009 | GT105 | | Glycosyltransferase Family 105 protein | | |
| Bta09018 | GH47 | | Glycoside Hydrolase Family 47 protein | | |
| Bta09037 | GH18-CBM14-CBM14 | | Glycoside Hydrolase Family 18 / Carbohydrate-Binding Module Family 14 protein | | |
| Bta09066 | CBM32-AA5_2 | | Carbohydrate-Binding Module Family 32 / Galactose oxidase | HGT | Fungi |
| Bta09154 | GT31 | | Glycosyltransferase Family 31 protein | | |
| Bta09170 | GH18 | | Glycoside Hydrolase Family 18 protein | | |
| Bta09254 | GH16_4 | | Glycoside Hydrolase Family 16 protein | | |
| Bta09256 | GH16_4 | | Glycoside Hydrolase Family 16 protein | | |
| Bta09257 | GH16 | (fragment) splicing problem; | Glycoside Hydrolase Family 16 protein | | |
| Bta09258 | GH16_4 | fragment C-term; | Glycoside Hydrolase Family 16 protein | | |
| Bta09259 | GH16_4 | | Glycoside Hydrolase Family 16 protein | | |
| Bta09289 | GT1 | fragment N-term; | Glycosyltransferase Family 1 protein | | |
| Bta09300 | GT32 | | Glycosyltransferase Family 32 protein | | |
| Bta09362 | AA3_2 | fragment N-term; | GMC oxidoreductase | | |
| Bta09385 | GT1 | fragment N-term; | Glycosyltransferase Family 1 protein | | |
| Bta09395 | CBM57 | | Carbohydrate-Binding Module Family 57 protein | | |
| Bta09500 | GT31-GT7 | (fragment) splicing problem; | Glycosyltransferase Family 31 / Glycosyltransferase Family 7 protein | | |
| Bta09508 | AA1 | | Multicopper oxidase | | |
| Bta09533 | GT7 | fragment N-term; | Glycosyltransferase Family 7 protein | | |
| Bta09615 | GT1 | | Glycosyltransferase Family 1 protein | | |
| Bta09633 | GH13_17-GH13_17 | (fragment) splicing problem; | Glycoside Hydrolase Family 13 protein | | |
| Bta09634 | GH13 | | Glycoside Hydrolase Family 13 protein | | |
| Bta09671 | CBM14 | | Carbohydrate-Binding Module Family 14 protein | | |
| Bta09696 | GH13_17 | | Glycoside Hydrolase Family 13 protein | | |
| Bta09701 | GT1 | | Glycosyltransferase Family 1 protein | | |
| Bta09742 | GH85 | | Glycoside Hydrolase Family 85 protein | | |
| Bta09856 | GH37 | | Glycoside Hydrolase Family 37 protein | | |
| Bta09882 | GT49 | | Glycosyltransferase Family 49 protein | | |
| Bta09947 | GT1 | | Glycosyltransferase Family 1 protein | | |
| Bta09948 | GT1 | | Glycosyltransferase Family 1 protein | | |
| Bta10017 | CBM14 | | Carbohydrate-Binding Module Family 14 protein | | |
| Bta10019 | CBM14 | | Carbohydrate-Binding Module Family 14 protein | | |
| Bta10022 | GH13_17 | | Glycoside Hydrolase Family 13 protein | | |
| Bta10045 | CE9 | (fragment) splicing problem; | Carbohydrate Esterase Family 9 protein | | |
| Bta10088 | GH1 | | Glycoside Hydrolase Family 1 protein | | |
| Bta10108 | CBM14 | | Carbohydrate-Binding Module Family 14 protein | | |
| Bta10133 | GT24 | | Glycosyltransferase Family 24 protein | | |
| Bta10152 | GH13_17 | fragment C-term; | Glycoside Hydrolase Family 13 protein | | |
| Bta10275 | GT39 | | Glycosyltransferase Family 39 protein | | |
| Bta10294 | GT57 | | Glycosyltransferase Family 57 protein | | |
| Bta10320 | GH18 | | Glycoside Hydrolase Family 18 protein | | |

| | | | | | |
|---|---|---|---|---|---|
| Bta10399 | GT7 | (fragment) splicing problem; | Glycosyltransferase Family 7 protein | | |
| Bta10525 | GT31 | (fragment) splicing problem; | Glycosyltransferase Family 31 protein | | |
| Bta10551 | GH35 | fragment N-term; | Glycoside Hydrolase Family 35 protein | | |
| Bta10552 | GH35 | fragment C-term; | Glycoside Hydrolase Family 35 protein | | |
| Bta10614 | GH13 | (fragment) splicing problem; | Glycoside Hydrolase Family 13 protein | | |
| Bta10620 | GH18 | | Glycoside Hydrolase Family 18 protein | | |
| Bta10635 | GH22 | fragment C-term; | Glycoside Hydrolase Family 22 protein | | |
| Bta10665 | GT43 | | Glycosyltransferase Family 43 protein | | |
| Bta10684 | CBM39 | | Carbohydrate-Binding Module Family 39 protein | | |
| Bta10764 | GT49 | | Glycosyltransferase Family 49 protein | | |
| Bta10765 | GT49 | (fragment) splicing problem; | Glycosyltransferase Family 49 protein | | |
| Bta10773 | GT23 | | Glycosyltransferase Family 23 protein | | |
| Bta10805 | GH18-CBM14 | | Glycoside Hydrolase Family 18 / Carbohydrate-Binding Module Family 14 protein | | |
| Bta10836 | GH18-CBM14-CBM14-CBM14 | fragment N-term; | Glycoside Hydrolase Family 18 / Carbohydrate-Binding Module Family 14 protein | | |
| Bta10858 | GT31 | (fragment) splicing problem; | Glycosyltransferase Family 31 protein | | |
| Bta10859 | GT31 | fragment N-term; | Glycosyltransferase Family 31 protein | | |
| Bta10860 | GT31-GT31-GT31 | fragment C-term; | Glycosyltransferase Family 31 protein | | |
| Bta10863 | GT1 | | Glycosyltransferase Family 1 protein | | |
| Bta10864 | GT1 | (fragment) splicing problem; | Glycosyltransferase Family 1 protein | | |
| Bta10865 | GT1 | | Glycosyltransferase Family 1 protein | | |
| Bta10911 | AA3_2 | | GMC oxidoreductase | | |
| Bta10981 | CBM14 | | Carbohydrate-Binding Module Family 14 protein | | |
| Bta11035 | AA3_2 | | GMC oxidoreductase | | |
| Bta11045 | GT8 | | Glycosyltransferase Family 8 protein | | |
| Bta11069 | GT1 | fragment N-term; | Glycosyltransferase Family 1 protein | | |
| Bta11080 | GT22 | | Glycosyltransferase Family 22 protein | | |
| Bta11100 | GH22 | | Glycoside Hydrolase Family 22 protein | | |
| Bta11101 | GT1 | fragment N-term; | Glycosyltransferase Family 1 protein | | |
| Bta11108 | GT1 | | Glycosyltransferase Family 1 protein | | |
| Bta11109 | GT1-GT1 | (fragment) splicing problem; | Glycosyltransferase Family 1 protein | | |
| Bta11221 | CE8 | | Carbohydrate Esterase Family 8 protein | HGT | Viridiplantae |
| Bta11222 | CE8 | fragment C-term; | Carbohydrate Esterase Family 8 protein | HGT | Viridiplantae |
| Bta11257 | GH1 | | Glycoside Hydrolase Family 1 protein | | |
| Bta11358 | GH13_17 | | Glycoside Hydrolase Family 13 protein | | |
| Bta11426 | GT66 | | Glycosyltransferase Family 66 protein | | |
| Bta11448 | GH13_15 | | Glycoside Hydrolase Family 13 protein | | |
| Bta11582 | AA3_2 | | GMC oxidoreductase | | |
| Bta11629 | AA3_2 | | GMC oxidoreductase | | |
| Bta11760 | GH13_25-GH133 | | Glycoside Hydrolase Family 13 / Glycoside Hydrolase Family 133 protein | | |
| Bta11780 | GH31 | | Glycoside Hydrolase Family 31 protein | | |
| Bta11831 | GH31 | | Glycoside Hydrolase Family 31 protein | | |
| Bta11864 | GT1 | (fragment) splicing problem; | Glycosyltransferase Family 1 protein | | |
| Bta11878 | AA1 | | Multicopper oxidase | | |
| Bta11896 | AA1 | | Multicopper oxidase | | |
| Bta11905 | AA15 | | Auxilliary Activities Family 15 protein | | |
| Bta11975 | GH13_17 | fragment N-term; | Glycoside Hydrolase Family 13 protein | | |
| Bta11977 | GH13_17 | | Glycoside Hydrolase Family 13 protein | | |
| Bta11978 | GH13_17 | (fragment) splicing problem; | Glycoside Hydrolase Family 13 protein | | |
| Bta11979 | GH13_17 | | Glycoside Hydrolase Family 13 protein | | |

| | | | | | |
|---|---|---|---|---|---|
| Bta12019 | GT2 | | Glycosyltransferase Family 2 protein | | |
| Bta12077 | GT1 | | Glycosyltransferase Family 1 protein | | |
| Bta12101 | GH13_17 | | Glycoside Hydrolase Family 13 protein | | |
| Bta12131 | GH20 | fragment C-term; | Glycoside Hydrolase Family 20 protein | | |
| Bta12153 | CBM14 | | Carbohydrate-Binding Module Family 14 protein | | |
| Bta12213 | GT33 | | Glycosyltransferase Family 33 protein | | |
| Bta12269 | GT1 | | Glycosyltransferase Family 1 protein | | |
| Bta12369 | GT58 | (fragment) splicing problem; | Glycosyltransferase Family 58 protein | | |
| Bta12376 | GT21 | | Glycosyltransferase Family 21 protein | | |
| Bta12433 | GT27-CBM13 | | Glycosyltransferase Family 27 / Carbohydrate-Binding Module Family 13 protein | | |
| Bta12440 | GT1 | | Glycosyltransferase Family 1 protein | | |
| Bta12453 | CBM32 | fragment C-term; | Carbohydrate-Binding Module Family 32 protein | HGT | Fungi |
| Bta12456 | CBM14-CBM14-CBM14-CBM14-CBM14-CBM14-CBM14 | | Carbohydrate-Binding Module Family 14 protein | | |
| Bta12575 | GT31 | | Glycosyltransferase Family 31 protein | | |
| Bta12578 | GH13_17 | | Glycoside Hydrolase Family 13 protein | | |
| Bta12583 | GT32 | fragment C-term; | Glycosyltransferase Family 32 protein | | |
| Bta12585 | GT32 | | Glycosyltransferase Family 32 protein | | |
| Bta12611 | AA3_2 | | GMC oxidoreductase | | |
| Bta12617 | GH31 | | Glycoside Hydrolase Family 31 protein | | |
| Bta12624 | GH13_17 | (fragment) splicing problem; | Glycoside Hydrolase Family 13 protein | | |
| Bta12670 | GH13_17 | | Glycoside Hydrolase Family 13 protein | | |
| Bta12671 | GH13_17 | | Glycoside Hydrolase Family 13 protein | | |
| Bta12678 | GH13_17 | | Glycoside Hydrolase Family 13 protein | | |
| Bta12680 | GH31 | | Glycoside Hydrolase Family 31 protein | | |
| Bta12682 | GH13_17 | | Glycoside Hydrolase Family 13 protein | | |
| Bta12683 | GH13_17 | | Glycoside Hydrolase Family 13 protein | | |
| Bta12684 | GT1 | fragment C-term; | Glycosyltransferase Family 1 protein | | |
| Bta12685 | GT1 | fragment N-term / C-term; | Glycosyltransferase Family 1 protein | | |
| Bta12702 | GT14 | | Glycosyltransferase Family 14 protein | | |
| Bta12717 | GT49 | | Glycosyltransferase Family 49 protein | | |
| Bta12718 | GH27 | | Glycoside Hydrolase Family 27 protein | | |
| Bta12836 | AA3_2 | | GMC oxidoreductase | | |
| Bta12860 | GH37 | | Glycoside Hydrolase Family 37 protein | | |
| Bta12862 | GT92 | | Glycosyltransferase Family 92 protein | | |
| Bta12867 | GH29 | (fragment) splicing problem; | Glycoside Hydrolase Family 29 protein | | |
| Bta12927 | GT10 | (fragment) splicing problem; | Glycosyltransferase Family 10 protein | | |
| Bta13236 | GH13_17 | | Glycoside Hydrolase Family 13 protein | | |
| Bta13237 | GH13_15 | fragment N-term; | Glycoside Hydrolase Family 13 protein | | |
| Bta13238 | GH13_17 | fragment C-term; | Glycoside Hydrolase Family 13 protein | | |
| Bta13239 | GH13_17 | fragment N-term; | Glycoside Hydrolase Family 13 protein | | |
| Bta13432 | GT2 | | Glycosyltransferase Family 2 protein | | |
| Bta13439 | GH2 | (fragment) splicing problem; | Glycoside Hydrolase Family 2 protein | | |
| Bta13494 | GT16 | | Glycosyltransferase Family 16 protein | | |
| Bta13568 | CBM48 | | Carbohydrate-Binding Module Family 48 protein | | |
| Bta13611 | GT7 | | Glycosyltransferase Family 7 protein | | |
| Bta13629 | GT105 | (fragment) splicing problem; | Glycosyltransferase Family 105 protein | | |
| Bta13631 | GT90 | | Glycosyltransferase Family 90 protein | | |
| Bta13660 | GH13_17 | | Glycoside Hydrolase Family 13 protein | | |
| Bta13669 | GH18-CBM18-CBM18-CBM18 | | Glycoside Hydrolase Family 18 / Carbohydrate-Binding Module Family 18 protein | | |

| | | | | | |
|---|---|---|---|---|---|
| Bta13679 | GT3 | | Glycosyltransferase Family 3 protein | | |
| Bta13694 | CBM14 | | Carbohydrate-Binding Module Family 14 protein | | |
| Bta13755 | GT1 | fragment C-term; | Glycosyltransferase Family 1 protein | | |
| Bta13756 | GT1 | | Glycosyltransferase Family 1 protein | | |
| Bta13757 | GT1 | (fragment) splicing problem; | Glycosyltransferase Family 1 protein | | |
| Bta13758 | GT1 | fragment N-term; | Glycosyltransferase Family 1 protein | | |
| Bta13867 | GT47-GT64 | | Glycosyltransferase Family 47 / Glycosyltransferase Family 64 protein | | |
| Bta13871 | CBM48-GH13_8 | | Carbohydrate-Binding Module Family 48 / Glycoside Hydrolase Family 13 protein | | |
| Bta13913 | GH13_17 | fragment N-term / C-term; | Glycoside Hydrolase Family 13 protein | | |
| Bta13914 | GH13_17 | | Glycoside Hydrolase Family 13 protein | | |
| Bta13961 | GH152 | | Glycoside Hydrolase Family 152 protein | HGT | Viridiplantae |
| Bta13982 | GH20 | fragment N-term; | Glycoside Hydrolase Family 20 protein | | |
| Bta14010 | GT7 | fragment N-term; | Glycosyltransferase Family 7 protein | | |
| Bta14019 | GT39 | | Glycosyltransferase Family 39 protein | | |
| Bta14112 | CBM14-CBM14 | | Carbohydrate-Binding Module Family 14 protein | | |
| Bta14119 | GT1 | fragment C-term; | Glycosyltransferase Family 1 protein | | |
| Bta14275 | GT27-CBM13 | | Glycosyltransferase Family 27 / Carbohydrate-Binding Module Family 13 protein | | |
| Bta14305 | GT7 | | Glycosyltransferase Family 7 protein | | |
| Bta14313 | GH13 | fragment C-term; | Glycoside Hydrolase Family 13 protein | | |
| Bta14369 | GH89 | fragment N-term / C-term; | Glycoside Hydrolase Family 89 protein | | |
| Bta14419 | GH13_17 | | Glycoside Hydrolase Family 13 protein | | |
| Bta14422 | GH13_17 | | Glycoside Hydrolase Family 13 protein | | |
| Bta14475 | CBM14-CBM14 | (fragment) splicing problem; | Carbohydrate-Binding Module Family 14 protein | | |
| Bta14509 | GH22-GH22 | | Glycoside Hydrolase Family 22 protein | | |
| Bta14571 | GT31 | (fragment) splicing problem; | Glycosyltransferase Family 31 protein | | |
| Bta14596 | GH18-CBM14-CBM14-CBM14-GH18-GH18-CBM14-GH18 | | Glycoside Hydrolase Family 18 / Carbohydrate-Binding Module Family 14 protein | | |
| Bta14598 | GT29 | (fragment) splicing problem; | Glycosyltransferase Family 29 protein | | |
| Bta14689 | GT66 | (fragment) splicing problem; | Glycosyltransferase Family 66 protein | | |
| Bta14723 | GH20 | | Glycoside Hydrolase Family 20 protein | | |
| Bta14774 | GH20 | | Glycoside Hydrolase Family 20 protein | | |
| Bta14797 | GH56 | | Glycoside Hydrolase Family 56 protein | | |
| Bta14885 | GT61 | | Glycosyltransferase Family 61 protein | HGT | Viridiplantae |
| Bta15044 | GT92 | | Glycosyltransferase Family 92 protein | | |
| Bta15239 | EXPN | | Distantly related to plant expansins | HGT | Viridiplantae |
| Bta15255 | GT1 | | Glycosyltransferase Family 1 protein | | |
| Bta15284 | AA1 | | Multicopper oxidase | | |
| Bta15340 | GH20 | (fragment) splicing problem; | Glycoside Hydrolase Family 20 protein | | |
| Bta15388 | GT32 | | Glycosyltransferase Family 32 protein | | |
| Bta15456 | GH31 | (fragment) splicing problem; | Glycoside Hydrolase Family 31 protein | | |
| Bta15607 | GH35 | fragment C-term; | Glycoside Hydrolase Family 35 protein | | |
| Bta15609 | GH35 | fragment N-term; | Glycoside Hydrolase Family 35 protein | | |
| Bta15610 | GH35 | | Glycoside Hydrolase Family 35 protein | | |
| Bta15649 | GH13_17 | | Glycoside Hydrolase Family 13 protein | | |
| Bta15659 | GT17 | | Glycosyltransferase Family 17 protein | HGT | Viridiplantae |
| Bta15703 | GT20 | | Glycosyltransferase Family 20 protein | | |
| Bta15714 | CBM50 | | Carbohydrate-Binding Module Family 50 protein | HGT | Bacteria |

*T. vaporariorum*

| Protein ID | Description | Model notes | Definition line | HGT | Donor |
|---|---|---|---|---|---|
| TLow_00586-RA | GT2 | fragment N-term; | Glycosyltransferase Family 2 protein | | |
| TLow_00611-RA | GT7 | | Glycosyltransferase Family 7 protein | | |
| TLow_00703-RA | CBM14 | | Carbohydrate-Binding Module Family 14 protein | | |
| TLow_01318-RA | AA3_2 | fragment N-term / C-term; | GMC oxidoreductase | | |
| TLow_01399-RA | GH79 | fragment C-term; | Glycoside Hydrolase Family 79 protein | | |
| TLow_01400-RA | GH79 | fragment C-term; | Glycoside Hydrolase Family 79 protein | | |
| TLow_01636-RA | GH13 | fragment N-term / C-term; | Glycoside Hydrolase Family 13 protein | | |
| TLow_01771-RA | GT61 | fragment N-term; | Glycosyltransferase Family 61 protein | | |
| Tv_00105-RA | CBM14 | | Carbohydrate-Binding Module Family 14 protein | | |
| Tv_00151-RA | AA3_2 | | GMC oxidoreductase | | |
| Tv_00286-RA | GT1 | | Glycosyltransferase Family 1 protein | | |
| Tv_00298-RA | GT1 | | Glycosyltransferase Family 1 protein | | |
| Tv_00316-RA | GT27-CBM13 | | Glycosyltransferase Family 27 / Carbohydrate-Binding Module Family 13 protein | | |
| Tv_00359-RA | GT27-CBM13 | fragment N-term; | Glycosyltransferase Family 27 / Carbohydrate-Binding Module Family 13 protein | | |
| Tv_00385-RA | GT2 | | Glycosyltransferase Family 2 protein | | |
| Tv_00408-RA | GT2 | fragment C-term; | Glycosyltransferase Family 2 protein | | |
| Tv_00469-RA | GT1 | | Glycosyltransferase Family 1 protein | | |
| Tv_00477-RA | GT31 | (fragment) splicing problem; | Glycosyltransferase Family 31 protein | | |
| Tv_00541-RA | GH22 | | Glycoside Hydrolase Family 22 protein | | |
| Tv_00627-RA | GT1 | | Glycosyltransferase Family 1 protein | | |
| Tv_00746-RA | GT105 | fragment N-term / C-term; | Glycosyltransferase Family 105 protein | | |
| Tv_00748-RA | GT105 | fragment N-term; | Glycosyltransferase Family 105 protein | | |
| Tv_00792-RA | CBM14 | | Carbohydrate-Binding Module Family 14 protein | | |
| Tv_00841-RA | GH20 | | Glycoside Hydrolase Family 20 protein | | |
| Tv_00902-RA | GT1 | fragment C-term; | Glycosyltransferase Family 1 protein | | |
| Tv_00903-RA | GT1-GT1-GT1 | fragment N-term; | Glycosyltransferase Family 1 protein | | |
| Tv_00904-RA | GT1-GT1 | | Glycosyltransferase Family 1 protein | | |
| Tv_00906-RA | GT1 | fragment N-term; | Glycosyltransferase Family 1 protein | | |
| Tv_00938-RA | GH13_17 | | Glycoside Hydrolase Family 13 protein | | |
| Tv_00942-RA | GT7 | | Glycosyltransferase Family 7 protein | | |
| Tv_00954-RA | GH37 | | Glycoside Hydrolase Family 37 protein | | |
| Tv_00966-RA | AA3_2 | | GMC oxidoreductase | | |
| Tv_01011-RA | GT8 | fragment C-term; | Glycosyltransferase Family 8 protein | | |
| Tv_01013-RA | GT8 | fragment C-term; | Glycosyltransferase Family 8 protein | | |
| Tv_01066-RA | GH31 | | Glycoside Hydrolase Family 31 protein | | |
| Tv_01108-RA | GH18-CBM14-CBM14-CBM14-GH18-GH18-CBM14 | | Glycoside Hydrolase Family 18 / Carbohydrate-Binding Module Family 14 protein | | |
| Tv_01131-RA | CBM14-CBM14-CBM14 | | Carbohydrate-Binding Module Family 14 protein | | |
| Tv_01133-RA | CBM14-CBM14-CBM14 | | Carbohydrate-Binding Module Family 14 protein | | |
| Tv_01134-RA | CBM14-CBM14-CBM14 | | Carbohydrate-Binding Module Family 14 protein | | |
| Tv_01135-RA | CBM14-CBM14-CBM14 | | Carbohydrate-Binding Module Family 14 protein | | |
| Tv_01140-RA | CBM14 | | Carbohydrate-Binding Module Family 14 protein | | |
| Tv_01194-RA | GT27 | fragment C-term; | Glycosyltransferase Family 27 protein | | |
| Tv_01195-RA | CBM13 | | Carbohydrate-Binding Module Family 13 protein | | |
| Tv_01250-RA | CE9 | (fragment) splicing problem; | Carbohydrate Esterase Family 9 protein | | |
| Tv_01347-RA | GT1 | | Glycosyltransferase Family 1 protein | | |
| Tv_01461-RA | CBM47 | (fragment) splicing problem; | Carbohydrate-Binding Module Family 47 protein | | |

| | | | |
|---|---|---|---|
| Tv_01486-RA | GT29 | | Glycosyltransferase Family 29 protein |
| Tv_01510-RA | GH84 | | Glycoside Hydrolase Family 84 protein |
| Tv_01518-RA | GH37 | fragment N-term; | Glycoside Hydrolase Family 37 protein |
| Tv_01563-RA | GT22 | | Glycosyltransferase Family 22 protein |
| Tv_01595-RA | GT31 | (fragment) splicing problem; | Glycosyltransferase Family 31 protein |
| Tv_01864-RA | GH27 | | Glycoside Hydrolase Family 27 protein |
| Tv_01872-RA | GH20 | | Glycoside Hydrolase Family 20 protein |
| Tv_01873-RA | GH20 | | Glycoside Hydrolase Family 20 protein |
| Tv_02025-RA | CBM14-CBM14 | (fragment) splicing problem; | Carbohydrate-Binding Module Family 14 protein |
| Tv_02051-RA | GT1 | (fragment) splicing problem; | Glycosyltransferase Family 1 protein |
| Tv_02090-RA | GT1 | | Glycosyltransferase Family 1 protein |
| Tv_02098-RA | CBM13 | | Carbohydrate-Binding Module Family 13 protein |
| Tv_02099-RA | CBM13 | (fragment) splicing problem; | Carbohydrate-Binding Module Family 13 protein |
| Tv_02104-RA | GT27 | fragment N-term / C-term; | Glycosyltransferase Family 27 protein |
| Tv_02109-RA | GH47 | fragment N-term; | Glycoside Hydrolase Family 47 protein |
| Tv_02146-RA | CBM14 | fragment N-term; | Carbohydrate-Binding Module Family 14 protein |
| Tv_02147-RA | CBM14 | fragment C-term; | Carbohydrate-Binding Module Family 14 protein |
| Tv_02163-RA | GT105 | | Glycosyltransferase Family 105 protein |
| Tv_02254-RA | GT4 | | Glycosyltransferase Family 4 protein |
| Tv_02275-RA | GT20 | fragment C-term; | Glycosyltransferase Family 20 protein |
| Tv_02277-RA | GT1 | fragment N-term; | Glycosyltransferase Family 1 protein |
| Tv_02342-RA | GH13 | | Glycoside Hydrolase Family 13 protein |
| Tv_02343-RA | GH13 | | Glycoside Hydrolase Family 13 protein |
| Tv_02344-RA | GH13 | | Glycoside Hydrolase Family 13 protein |
| Tv_02358-RA | GT2 | | Glycosyltransferase Family 2 protein |
| Tv_02425-RA | CBM14 | | Carbohydrate-Binding Module Family 14 protein |
| Tv_02426-RA | CBM14 | | Carbohydrate-Binding Module Family 14 protein |
| Tv_02441-RA | GT35 | | Glycosyltransferase Family 35 protein |
| Tv_02581-RA | GT4 | | Glycosyltransferase Family 4 protein |
| Tv_02586-RA | GT7 | | Glycosyltransferase Family 7 protein |
| Tv_03404-RA | GT8-GT49 | (fragment) splicing problem; | Glycosyltransferase Family 8 / Glycosyltransferase Family 49 protein |
| Tv_03505-RA | GH27 | | Glycoside Hydrolase Family 27 protein |
| Tv_03530-RA | GH38 | | Glycoside Hydrolase Family 38 protein |
| Tv_03544-RA | GT20 | fragment N-term; | Glycosyltransferase Family 20 protein |
| Tv_03582-RA | GT27-CBM13 | | Glycosyltransferase Family 27 / Carbohydrate-Binding Module Family 13 protein |
| Tv_03624-RA | GT22 | fragment C-term; | Glycosyltransferase Family 22 protein |
| Tv_03625-RA | GT22 | (fragment) splicing problem; | Glycosyltransferase Family 22 protein |
| Tv_03746-RA | GH13_17 | | Glycoside Hydrolase Family 13 protein |
| Tv_03782-RA | GT43 | fragment C-term; | Glycosyltransferase Family 43 protein |
| Tv_03807-RA | GT47-GT64 | fragment N-term; | Glycosyltransferase Family 47 / Glycosyltransferase Family 64 protein |
| Tv_03973-RA | GT27-CBM13 | (fragment) splicing problem; | Glycosyltransferase Family 27 / Carbohydrate-Binding Module Family 13 protein |
| Tv_03990-RA | GT23 | | Glycosyltransferase Family 23 protein |
| Tv_04069-RA | GH13_17 | | Glycoside Hydrolase Family 13 protein |
| Tv_04071-RA | GH13_17 | | Glycoside Hydrolase Family 13 protein |
| Tv_04072-RA | GH13_17 | | Glycoside Hydrolase Family 13 protein |
| Tv_04073-RA | GH13_17-GH13_17 | | Glycoside Hydrolase Family 13 protein |
| Tv_04075-RA | GH13_17 | | Glycoside Hydrolase Family 13 protein |
| Tv_04077-RA | GH13_17 | | Glycoside Hydrolase Family 13 protein |
| Tv_04078-RA | GH13_17 | | Glycoside Hydrolase Family 13 protein |

| | | | | | |
|---|---|---|---|---|---|
| Tv_04079-RA | GH13 | fragment C-term; | Glycoside Hydrolase Family 13 protein | HGT | Bacteria |
| Tv_04113-RA | GT1 | | Glycosyltransferase Family 1 protein | | |
| Tv_04122-RA | GH13_17 | | Glycoside Hydrolase Family 13 protein | | |
| Tv_04124-RA | GH13_17 | fragment N-term / C-term; | Glycoside Hydrolase Family 13 protein | | |
| Tv_04131-RA | GH13_17 | (fragment) splicing problem; | Glycoside Hydrolase Family 13 protein | | |
| Tv_04132-RA | GH13_17 | | Glycoside Hydrolase Family 13 protein | | |
| Tv_04133-RA | GH13_17 | | Glycoside Hydrolase Family 13 protein | | |
| Tv_04134-RA | GH13_17 | | Glycoside Hydrolase Family 13 protein | | |
| Tv_04172-RA | GT49 | | Glycosyltransferase Family 49 protein | | |
| Tv_04197-RA | GH18 | | Glycoside Hydrolase Family 18 protein | | |
| Tv_04246-RA | GT1-GT1 | | Glycosyltransferase Family 1 protein | | |
| Tv_04295-RA | GH18-CBM14 | | Glycoside Hydrolase Family 18 / Carbohydrate-Binding Module Family 14 protein | | |
| Tv_04305-RA | GT24 | (fragment) splicing problem; | Glycosyltransferase Family 24 protein | | |
| Tv_04489-RA | GH13_17 | (fragment) splicing problem; | Glycoside Hydrolase Family 13 protein | | |
| Tv_04500-RA | GH38 | | Glycoside Hydrolase Family 38 protein | | |
| Tv_04564-RA | GT4 | | Glycosyltransferase Family 4 protein | | |
| Tv_04576-RA | GH13_17 | (fragment) splicing problem; | Glycoside Hydrolase Family 13 protein | | |
| Tv_04643-RA | GT61 | | Glycosyltransferase Family 61 protein | | |
| Tv_04727-RA | GT1 | | Glycosyltransferase Family 1 protein | | |
| Tv_04751-RA | GH13_17 | (fragment) splicing problem; | Glycoside Hydrolase Family 13 protein | | |
| Tv_04752-RA | GH13_17 | (fragment) splicing problem; | Glycoside Hydrolase Family 13 protein | | |
| Tv_04852-RA | CBM14 | | Carbohydrate-Binding Module Family 14 protein | | |
| Tv_04857-RA | GT31 | | Glycosyltransferase Family 31 protein | | |
| Tv_04859-RA | GT1-GT1 | fragment N-term; | Glycosyltransferase Family 1 protein | | |
| Tv_04891-RA | GT65 | (fragment) splicing problem; | Glycosyltransferase Family 65 protein | | |
| Tv_04916-RA | GT1 | fragment N-term; | Glycosyltransferase Family 1 protein | | |
| Tv_04931-RA | GT1 | | Glycosyltransferase Family 1 protein | | |
| Tv_04932-RA | GT1 | | Glycosyltransferase Family 1 protein | | |
| Tv_04934-RA | GT27-CBM13 | (fragment) splicing problem; | Glycosyltransferase Family 27 / Carbohydrate-Binding Module Family 13 protein | | |
| Tv_04957-RA | GT20 | | Glycosyltransferase Family 20 protein | | |
| Tv_04960-RA | GH13_17 | fragment C-term; | Glycoside Hydrolase Family 13 protein | | |
| Tv_04997-RA | GH13_17 | fragment N-term; | Glycoside Hydrolase Family 13 protein | | |
| Tv_04998-RA | GH13 | fragment C-term; | Glycoside Hydrolase Family 13 protein | | |
| Tv_04999-RA | GH13_17 | (fragment) splicing problem; | Glycoside Hydrolase Family 13 protein | | |
| Tv_05000-RA | GH13_17-GH13 | fragment C-term; | Glycoside Hydrolase Family 13 protein | | |
| Tv_05001-RA | GH13_17 | (fragment) splicing problem; | Glycoside Hydrolase Family 13 protein | | |
| Tv_05069-RA | GT49 | (fragment) splicing problem; | Glycosyltransferase Family 49 protein | | |
| Tv_05077-RA | GT21 | fragment N-term; | Glycosyltransferase Family 21 protein | | |
| Tv_05083-RA | GT1 | | Glycosyltransferase Family 1 protein | | |
| Tv_05090-RA | GT58 | | Glycosyltransferase Family 58 protein | | |
| Tv_05105-RA | PL1_4 | fragment C-term; | Polysaccharide Lyase Family 1 protein | HGT | Complex: Bacteria or Fungi |
| Tv_05119-RA | GT1 | | Glycosyltransferase Family 1 protein | | |
| Tv_05121-RA | GT1 | | Glycosyltransferase Family 1 protein | | |
| Tv_05122-RA | GT1 | | Glycosyltransferase Family 1 protein | | |
| Tv_05145-RA | GT1 | | Glycosyltransferase Family 1 protein | | |
| Tv_05165-RA | AA3_2 | | GMC oxidoreductase | | |
| Tv_05314-RA | GT49 | | Glycosyltransferase Family 49 protein | | |
| Tv_05317-RA | GH27 | | Glycoside Hydrolase Family 27 protein | | |
| Tv_05323-RA | GT90 | | Glycosyltransferase Family 90 protein | | |

| | | | |
|---|---|---|---|
| Tv_05391-RA | GT1 | | Glycosyltransferase Family 1 protein |
| Tv_05430-RA | GT1 | fragment C-term; | Glycosyltransferase Family 1 protein |
| Tv_05441-RA | GT1 | (fragment) splicing problem; | Glycosyltransferase Family 1 protein |
| Tv_05533-RA | GT54 | (fragment) splicing problem; | Glycosyltransferase Family 54 protein |
| Tv_05535-RA | GT54 | fragment C-term; | Glycosyltransferase Family 54 protein |
| Tv_05541-RA | GT25 | fragment C-term; | Glycosyltransferase Family 25 protein |
| Tv_05548-RA | GT25 | fragment N-term; | Glycosyltransferase Family 25 protein |
| Tv_05549-RA | GT54 | | Glycosyltransferase Family 54 protein |
| Tv_05587-RA | GT66 | | Glycosyltransferase Family 66 protein |
| Tv_05613-RA | AA3_2 | | GMC oxidoreductase |
| Tv_05624-RA | AA3_2 | | GMC oxidoreductase |
| Tv_05656-RA | GH20 | | Glycoside Hydrolase Family 20 protein |
| Tv_05701-RA | CBM47 | | Carbohydrate-Binding Module Family 47 protein |
| Tv_05908-RA | AA3_2 | | GMC oxidoreductase |
| Tv_05909-RA | GH13_17 | | Glycoside Hydrolase Family 13 protein |
| Tv_05910-RA | GH13_17 | | Glycoside Hydrolase Family 13 protein |
| Tv_05917-RA | GT1-GT1 | | Glycosyltransferase Family 1 protein |
| Tv_05937-RA | GH13 | (fragment) splicing problem; | Glycoside Hydrolase Family 13 protein |
| Tv_05979-RA | GT1 | fragment N-term; | Glycosyltransferase Family 1 protein |
| Tv_05987-RA | GT1 | fragment N-term; | Glycosyltransferase Family 1 protein |
| Tv_06020-RA | GH89 | fragment C-term; | Glycoside Hydrolase Family 89 protein |
| Tv_06021-RA | GH89 | fragment N-term; | Glycoside Hydrolase Family 89 protein |
| Tv_06050-RA | GH13_17 | (fragment) splicing problem; | Glycoside Hydrolase Family 13 protein |
| Tv_06132-RA | GT10 | fragment N-term; | Glycosyltransferase Family 10 protein |
| Tv_06277-RA | GH20 | | Glycoside Hydrolase Family 20 protein |
| Tv_06371-RA | GT10 | | Glycosyltransferase Family 10 protein |
| Tv_06393-RA | GT1 | | Glycosyltransferase Family 1 protein |
| Tv_06471-RA | CBM14 | | Carbohydrate-Binding Module Family 14 protein |
| Tv_06484-RA | GH18-CBM14-CBM14 | | Glycoside Hydrolase Family 18 / Carbohydrate-Binding Module Family 14 protein |
| Tv_06504-RA | GT66 | fragment C-term; | Glycosyltransferase Family 66 protein |
| Tv_06589-RA | CBM50 | | Carbohydrate-Binding Module Family 50 protein |
| Tv_06597-RA | GH20 | | Glycoside Hydrolase Family 20 protein |
| Tv_06654-RA | GT22 | | Glycosyltransferase Family 22 protein |
| Tv_06726-RA | AA1 | | Multicopper oxidase |
| Tv_06734-RA | GT31-GT7 | | Glycosyltransferase Family 31 / Glycosyltransferase Family 7 protein |
| Tv_06794-RA | GT110 | | Glycosyltransferase Family 110 protein |
| Tv_06971-RA | GT47-GT64 | | Glycosyltransferase Family 47 / Glycosyltransferase Family 64 protein |
| Tv_07030-RA | CBM14 | | Carbohydrate-Binding Module Family 14 protein |
| Tv_07082-RA | GH85 | | Glycoside Hydrolase Family 85 protein |
| Tv_07120-RA | AA3_2 | | GMC oxidoreductase |
| Tv_07170-RA | GT13 | | Glycosyltransferase Family 13 protein |
| Tv_07242-RA | GT8 | (fragment) splicing problem; | Glycosyltransferase Family 8 protein |
| Tv_07372-RA | GH2 | | Glycoside Hydrolase Family 2 protein |
| Tv_07391-RA | GT92 | | Glycosyltransferase Family 92 protein |
| Tv_07457-RA | AA3_2 | | GMC oxidoreductase |
| Tv_07510-RA | GH47 | | Glycoside Hydrolase Family 47 protein |
| Tv_07550-RA | GT31 | (fragment) splicing problem; | Glycosyltransferase Family 31 protein |
| Tv_07557-RA | AA1 | | Multicopper oxidase |
| Tv_07565-RA | GT31 | | Glycosyltransferase Family 31 protein |

| | | | |
|---|---|---|---|
| Tv_07716-RA | GT41 | | Glycosyltransferase Family 41 protein |
| Tv_07809-RA | GH18 | (fragment) splicing problem; | Glycoside Hydrolase Family 18 protein |
| Tv_07810-RA | GH18 | | Glycoside Hydrolase Family 18 protein |
| Tv_07905-RA | GH13_17 | | Glycoside Hydrolase Family 13 protein |
| Tv_07947-RA | GT39 | | Glycosyltransferase Family 39 protein |
| Tv_08013-RA | GH56 | | Glycoside Hydrolase Family 56 protein |
| Tv_08063-RA | GT3 | | Glycosyltransferase Family 3 protein |
| Tv_08077-RA | CBM14-CBM14-CBM14-CBM14-CBM14-CBM14-CBM14 | | Carbohydrate-Binding Module Family 14 protein |
| Tv_08081-RA | GT43 | | Glycosyltransferase Family 43 protein |
| Tv_08177-RA | GH31 | | Glycoside Hydrolase Family 31 protein |
| Tv_08210-RA | GH20 | | Glycoside Hydrolase Family 20 protein |
| Tv_08263-RA | CBM14 | | Carbohydrate-Binding Module Family 14 protein |
| Tv_08317-RA | GH30_1 | | Glycoside Hydrolase Family 30 protein |
| Tv_08448-RA | GH47 | (fragment) splicing problem; | Glycoside Hydrolase Family 47 protein |
| Tv_08655-RA | GT31 | fragment C-term; | Glycosyltransferase Family 31 protein |
| Tv_08671-RA | GT1 | | Glycosyltransferase Family 1 protein |
| Tv_08676-RA | GH13-GH13-GH13 | | Glycoside Hydrolase Family 13 protein |
| Tv_08678-RA | GH13_17 | | Glycoside Hydrolase Family 13 protein |
| Tv_08680-RA | GH13_17 | | Glycoside Hydrolase Family 13 protein |
| Tv_08691-RA | GH116 | | Glycoside Hydrolase Family 116 protein |
| Tv_08765-RA | GT1 | | Glycosyltransferase Family 1 protein |
| Tv_08960-RA | GT59 | fragment C-term; | Glycosyltransferase Family 59 protein |
| Tv_08968-RA | CBM14 | | Carbohydrate-Binding Module Family 14 protein |
| Tv_08969-RA | CBM14 | | Carbohydrate-Binding Module Family 14 protein |
| Tv_08979-RA | GH13_17 | | Glycoside Hydrolase Family 13 protein |
| Tv_08981-RA | GH13_17 | | Glycoside Hydrolase Family 13 protein |
| Tv_09051-RA | GT31 | | Glycosyltransferase Family 31 protein |
| Tv_09054-RA | GH18 | | Glycoside Hydrolase Family 18 protein |
| Tv_09074-RA | GT105 | | Glycosyltransferase Family 105 protein |
| Tv_09101-RA | CBM39 | | Carbohydrate-Binding Module Family 39 protein |
| Tv_09204-RA | GH22 | | Glycoside Hydrolase Family 22 protein |
| Tv_09245-RA | GT57 | | Glycosyltransferase Family 57 protein |
| Tv_09445-RA | GT66 | (fragment) splicing problem; | Glycosyltransferase Family 66 protein |
| Tv_09527-RA | GT39 | | Glycosyltransferase Family 39 protein |
| Tv_09545-RA | GT31 | | Glycosyltransferase Family 31 protein |
| Tv_09550-RA | GT31 | fragment N-term; | Glycosyltransferase Family 31 protein |
| Tv_09566-RA | GH1 | (fragment) splicing problem; | Glycoside Hydrolase Family 1 protein |
| Tv_09568-RA | GH22 | | Glycoside Hydrolase Family 22 protein |
| Tv_09581-RA | GH13_17 | | Glycoside Hydrolase Family 13 protein |
| Tv_09582-RA | GH13_17 | fragment C-term; | Glycoside Hydrolase Family 13 protein |
| Tv_09589-RA | GH13-GH13 | (fragment) splicing problem; | Glycoside Hydrolase Family 13 protein |
| Tv_09675-RA | GH13_17 | | Glycoside Hydrolase Family 13 protein |
| Tv_09676-RA | GH13_17 | | Glycoside Hydrolase Family 13 protein |
| Tv_09677-RA | GH13_17 | | Glycoside Hydrolase Family 13 protein |
| Tv_09686-RA | GH13_17 | | Glycoside Hydrolase Family 13 protein |
| Tv_09694-RA | GH39 | | Glycoside Hydrolase Family 39 protein |
| Tv_09747-RA | GH38 | fragment C-term; | Glycoside Hydrolase Family 38 protein |
| Tv_09748-RA | GH38 | fragment N-term; | Glycoside Hydrolase Family 38 protein |
| Tv_09765-RA | GH13 | (fragment) splicing problem; | Glycoside Hydrolase Family 13 protein |

| | | | |
|---|---|---|---|
| Tv_09794-RA | GH37 | fragment C-term; | Glycoside Hydrolase Family 37 protein |
| Tv_09799-RA | GH37-GH37 | fragment N-term / C-term; | Glycoside Hydrolase Family 37 protein |
| Tv_09800-RA | GH37 | fragment C-term; | Glycoside Hydrolase Family 37 protein |
| Tv_09802-RA | GH37 | fragment N-term; | Glycoside Hydrolase Family 37 protein |
| Tv_09868-RA | GT105 | fragment N-term; | Glycosyltransferase Family 105 protein |
| Tv_09920-RA | GT1 | fragment N-term; | Glycosyltransferase Family 1 protein |
| Tv_10002-RA | GH13_17 | (fragment) splicing problem; | Glycoside Hydrolase Family 13 protein |
| Tv_10139-RA | GT105 | fragment N-term / C-term; | Glycosyltransferase Family 105 protein |
| Tv_10141-RA | GT105 | fragment N-term; | Glycosyltransferase Family 105 protein |
| Tv_10332-RA | GH31 | | Glycoside Hydrolase Family 31 protein |
| Tv_10623-RA | GH31 | fragment C-term; | Glycoside Hydrolase Family 31 protein |
| Tv_10709-RA | GT1-GT1-GT1 | | Glycosyltransferase Family 1 protein |
| Tv_10792-RA | AA3_2 | fragment C-term; | GMC oxidoreductase |
| Tv_10793-RA | AA3 | fragment N-term; | GMC oxidoreductase |
| Tv_10808-RA | GT1 | | Glycosyltransferase Family 1 protein |
| Tv_10938-RA | GT31-GT7 | fragment C-term; | Glycosyltransferase Family 31 / Glycosyltransferase Family 7 protein |
| Tv_11122-RA | GH13_17 | fragment N-term / C-term; | Glycoside Hydrolase Family 13 protein |
| Tv_11123-RA | GH13 | fragment C-term; | Glycoside Hydrolase Family 13 protein |
| Tv_11124-RA | GH13_17 | | Glycoside Hydrolase Family 13 protein |
| Tv_11131-RA | CBM21 | | Carbohydrate-Binding Module Family 21 protein |
| Tv_11211-RA | GH13_25-GH133 | (fragment) splicing problem; | Glycoside Hydrolase Family 13 / Glycoside Hydrolase Family 133 protein |
| Tv_11223-RA | GH38 | fragment N-term; | Glycoside Hydrolase Family 38 protein |
| Tv_11224-RA | GH38 | fragment N-term / C-term; | Glycoside Hydrolase Family 38 protein |
| Tv_11262-RA | CBM57 | | Carbohydrate-Binding Module Family 57 protein |
| Tv_11360-RA | AA15 | | Auxilliary Activities Family 15 protein |
| Tv_11375-RA | AA1 | | Multicopper oxidase |
| Tv_11376-RA | AA1 | | Multicopper oxidase |
| Tv_11629-RA | GT13 | | Glycosyltransferase Family 13 protein |
| Tv_11649-RA | GT27-CBM13 | | Glycosyltransferase Family 27 / Carbohydrate-Binding Module Family 13 protein |
| Tv_11809-RA | GH18 | | Glycoside Hydrolase Family 18 protein |
| Tv_11816-RA | CBM48-GH13_8 | | Carbohydrate-Binding Module Family 48 / Glycoside Hydrolase Family 13 protein |
| Tv_11876-RA | GH31 | | Glycoside Hydrolase Family 31 protein |
| Tv_11940-RA | GT1 | | Glycosyltransferase Family 1 protein |
| Tv_12008-RA | GT47-GT64 | | Glycosyltransferase Family 47 / Glycosyltransferase Family 64 protein |
| Tv_12259-RA | GT1 | | Glycosyltransferase Family 1 protein |
| Tv_12473-RA | GT31 | (fragment) splicing problem; | Glycosyltransferase Family 31 protein |
| Tv_12479-RA | GT2 | | Glycosyltransferase Family 2 protein |
| Tv_12497-RA | GT1 | | Glycosyltransferase Family 1 protein |
| Tv_12648-RA | AA3 | fragment N-term; | GMC oxidoreductase |
| Tv_12650-RA | AA3_2 | | GMC oxidoreductase |
| Tv_12651-RA | AA3_2 | | GMC oxidoreductase |
| Tv_12652-RA | AA3_2 | fragment N-term; | GMC oxidoreductase |
| Tv_12653-RA | AA3_2 | fragment C-term; | GMC oxidoreductase |
| Tv_12655-RA | AA3_2 | fragment N-term; | GMC oxidoreductase |
| Tv_12660-RA | AA3_2 | fragment C-term; | GMC oxidoreductase |
| Tv_12661-RA | AA3_2 | | GMC oxidoreductase |
| Tv_12662-RA | AA3_2 | | GMC oxidoreductase |
| Tv_12663-RA | AA3_2 | fragment N-term; | GMC oxidoreductase |
| Tv_12664-RA | AA3_2 | fragment C-term; | GMC oxidoreductase |

| | | | |
|---|---|---|---|
| Tv_12665-RA | AA3_2 | | GMC oxidoreductase |
| Tv_12666-RA | AA3_2 | | GMC oxidoreductase |
| Tv_12766-RA | GT22 | | Glycosyltransferase Family 22 protein |
| Tv_12853-RA | CBM14 | | Carbohydrate-Binding Module Family 14 protein |
| Tv_12865-RA | GT1 | | Glycosyltransferase Family 1 protein |
| Tv_12868-RA | GT1 | fragment N-term; | Glycosyltransferase Family 1 protein |
| Tv_12880-RA | GH1-GH1 | (fragment) splicing problem; | Glycoside Hydrolase Family 1 protein |
| Tv_12890-RA | GH63 | | Glycoside Hydrolase Family 63 protein |
| Tv_12935-RA | GT1 | | Glycosyltransferase Family 1 protein |
| Tv_13034-RA | GT90 | | Glycosyltransferase Family 90 protein |
| Tv_13120-RA | GH13_17 | | Glycoside Hydrolase Family 13 protein |
| Tv_13161-RA | GT33 | | Glycosyltransferase Family 33 protein |
| Tv_13175-RA | GH79 | fragment N-term; | Glycoside Hydrolase Family 79 protein |
| Tv_13281-RA | GT1-GT1 | (fragment) splicing problem; | Glycosyltransferase Family 1 protein |
| Tv_13379-RA | GH20 | fragment C-term; | Glycoside Hydrolase Family 20 protein |
| Tv_13434-RA | AA15 | | Auxilliary Activities Family 15 protein |
| Tv_13463-RA | GT32 | | Glycosyltransferase Family 32 protein |
| Tv_13587-RA | AA3_2 | (fragment) splicing problem; | GMC oxidoreductase |
| Tv_13711-RA | GH35 | | Glycoside Hydrolase Family 35 protein |
| Tv_13737-RA | GH18-CBM18-CBM18-CBM18 | fragment N-term; | Glycoside Hydrolase Family 18 / Carbohydrate-Binding Module Family 18 protein |
| Tv_13767-RA | GH18-CBM14-CBM14 | | Glycoside Hydrolase Family 18 / Carbohydrate-Binding Module Family 14 protein |
| Tv_13768-RA | AA3_2 | | GMC oxidoreductase |
| Tv_13841-RA | GT8 | | Glycosyltransferase Family 8 protein |
| Tv_13954-RA | GT1-GT1 | fragment N-term; | Glycosyltransferase Family 1 protein |
| Tv_14001-RA | CBM21 | | Carbohydrate-Binding Module Family 21 protein |
| Tv_14029-RA | GH22 | | Glycoside Hydrolase Family 22 protein |
| Tv_14032-RA | GT1 | | Glycosyltransferase Family 1 protein |
| Tv_14034-RA | GT1 | fragment N-term; | Glycosyltransferase Family 1 protein |
| Tv_14056-RA | GH16_4 | | Glycoside Hydrolase Family 16 protein |
| Tv_14057-RA | GH16_4 | | Glycoside Hydrolase Family 16 protein |
| Tv_14093-RA | GT14 | | Glycosyltransferase Family 14 protein |
| Tv_14174-RA | AA1 | fragment N-term; | Multicopper oxidase |
| Tv_14176-RA | GT16 | | Glycosyltransferase Family 16 protein |
| Tv_14208-RA | GH37 | fragment N-term; | Glycoside Hydrolase Family 37 protein |
| Tv_14213-RA | GH37 | | Glycoside Hydrolase Family 37 protein |
| Tv_14255-RA | GT105 | fragment N-term; | Glycosyltransferase Family 105 protein |
| Tv_14305-RA | CBM14 | | Carbohydrate-Binding Module Family 14 protein |
| Tv_14306-RA | GH37 | fragment N-term; | Glycoside Hydrolase Family 37 protein |
| Tv_14307-RA | GH37 | fragment C-term; | Glycoside Hydrolase Family 37 protein |
| Tv_14448-RA | GT1 | fragment C-term; | Glycosyltransferase Family 1 protein |
| Tv_14482-RA | CBM48 | | Carbohydrate-Binding Module Family 48 protein |
| Tv_14568-RA | GT1 | fragment N-term; | Glycosyltransferase Family 1 protein |
| Tv_14587-RA | GT1 | | Glycosyltransferase Family 1 protein |
| Tv_14760-RA | GH35 | fragment N-term; | Glycoside Hydrolase Family 35 protein |
| Tv_14790-RA | GT1 | | Glycosyltransferase Family 1 protein |
| Tv_14792-RA | GT1 | fragment N-term; | Glycosyltransferase Family 1 protein |
| Tv_14824-RA | GH13_17 | | Glycoside Hydrolase Family 13 protein |
| Tv_14833-RA | GT1 | | Glycosyltransferase Family 1 protein |
| Tv_14944-RA | GH37 | | Glycoside Hydrolase Family 37 protein |

| | | | | | |
|---|---|---|---|---|---|
| Tv_15005-RA | GH47 | | Glycoside Hydrolase Family 47 protein | | |
| Tv_15020-RA | GT1 | fragment C-term; | Glycosyltransferase Family 1 protein | | |
| Tv_15045-RA | GT31 | | Glycosyltransferase Family 31 protein | | |
| Tv_15081-RA | GT54 | fragment C-term; | Glycosyltransferase Family 54 protein | | |
| Tv_15088-RA | GT54 | fragment N-term; | Glycosyltransferase Family 54 protein | | |
| Tv_15133-RA | GT92 | | Glycosyltransferase Family 92 protein | | |
| Tv_15139-RA | GH29 | | Glycoside Hydrolase Family 29 protein | | |
| Tv_15167-RA | GT1 | | Glycosyltransferase Family 1 protein | | |
| Tv_15503-RA | GT49 | | Glycosyltransferase Family 49 protein | | |
| Tv_15512-RA | GT4 | | Glycosyltransferase Family 4 protein | | |
| Tv_15739-RA | GH13_17 | (fragment) splicing problem; | Glycoside Hydrolase Family 13 protein | | |
| Tv_15755-RA | GH47 | | Glycoside Hydrolase Family 47 protein | | |
| Tv_15844-RA | GH13 | fragment C-term; | Glycoside Hydrolase Family 13 protein | | |
| Tv_15845-RA | GH13_17 | fragment N-term; | Glycoside Hydrolase Family 13 protein | | |
| Tv_15891-RA | CBM14-CBM14-CBM14 | | Carbohydrate-Binding Module Family 14 protein | | |
| Tv_15898-RA | GH99 | | Glycoside Hydrolase Family 99 protein | | |
| Tv_15925-RA | CBM14 | | Carbohydrate-Binding Module Family 14 protein | | |
| Tv_15928-RA | GH152 | (fragment) splicing problem; | Glycoside Hydrolase Family 152 protein | HGT | Viridiplantae |
| Tv_15933-RA | CBM14 | | Carbohydrate-Binding Module Family 14 protein | | |
| Tv_15955-RA | GH18 | fragment C-term; | Glycoside Hydrolase Family 18 protein | | |
| Tv_15956-RA | GH18-GH18 | fragment N-term / C-term; | Glycoside Hydrolase Family 18 protein | | |
| Tv_15957-RA | GH18-CBM14 | fragment N-term; | Glycoside Hydrolase Family 18 / Carbohydrate-Binding Module Family 14 protein | | |
| Tv_16156-RA | GT47 | fragment C-term; | Glycosyltransferase Family 47 protein | | |

*F. occidentalis*

| Protein ID | Description | Model notes | Definition line | HGT | Donor |
|---|---|---|---|---|---|
| XP_026271392.1 | GT1 | | Glycosyltransferase Family 1 protein | | |
| XP_026271551.1 | AA3_2 | | GMC oxidoreductase | | |
| XP_026271901.1 | GH38 | fragment C-term; | Glycoside Hydrolase Family 38 protein | | |
| XP_026272000.1 | GT105 | | Glycosyltransferase Family 105 protein | | |
| XP_026272052.1 | GH16_4 | | Glycoside Hydrolase Family 16 protein | | |
| XP_026272092.1 | GT22 | | Glycosyltransferase Family 22 protein | | |
| XP_026272095.1 | CBM14 | | Carbohydrate-Binding Module Family 14 protein | | |
| XP_026272203.1 | GH1 | | Glycoside Hydrolase Family 1 protein | | |
| XP_026272209.1 | GT10 | | Glycosyltransferase Family 10 protein | | |
| XP_026272388.1 | GH1 | fragment N-term / C-term; | Glycoside Hydrolase Family 1 protein | | |
| XP_026272402.1 | GH16_4 | | Glycoside Hydrolase Family 16 protein | | |
| XP_026272427.1 | GT58 | | Glycosyltransferase Family 58 protein | | |
| XP_026272503.1 | GT31 | | Glycosyltransferase Family 31 protein | | |
| XP_026272515.1 | GT7 | | Glycosyltransferase Family 7 protein | | |
| XP_026272538.1 | GH37 | | Glycoside Hydrolase Family 37 protein | | |
| XP_026272610.1 | GT31 | | Glycosyltransferase Family 31 protein | | |
| XP_026272668.1 | GH22 | | Glycoside Hydrolase Family 22 protein | | |
| XP_026272669.1 | CBM21 | | Carbohydrate-Binding Module Family 21 protein | | |
| XP_026272679.1 | GH22 | | Glycoside Hydrolase Family 22 protein | | |
| XP_026272691.1 | GT20 | | Glycosyltransferase Family 20 protein | | |
| XP_026272815.1 | GT105 | | Glycosyltransferase Family 105 protein | | |
| XP_026272828.1 | GH18-CBM14 | | Glycoside Hydrolase Family 18 / Carbohydrate-Binding Module Family 14 protein | | |

| | | | | | |
|---|---|---|---|---|---|
| XP_026272949.1 | GT39 | | Glycosyltransferase Family 39 protein | | |
| XP_026272953.1 | GT31-GT7 | (fragment) splicing problem; | Glycosyltransferase Family 31 / Glycosyltransferase Family 7 protein | | |
| XP_026272958.1 | GT8-GT49 | | Glycosyltransferase Family 8 / Glycosyltransferase Family 49 protein | | |
| XP_026273012.1 | GT49 | | Glycosyltransferase Family 49 protein | | |
| XP_026273055.1 | GH22 | | Glycoside Hydrolase Family 22 protein | | |
| XP_026273122.1 | GH32 | | Glycoside Hydrolase Family 32 protein | HGT | Bacteria |
| XP_026273138.1 | AA1 | fragment C-term; | Multicopper oxidase | | |
| XP_026273151.1 | GH35 | | Glycoside Hydrolase Family 35 protein | | |
| XP_026273242.1 | GT2 | | Glycosyltransferase Family 2 protein | | |
| XP_026273431.1 | GT47-GT64 | | Glycosyltransferase Family 47 / Glycosyltransferase Family 64 protein | | |
| XP_026273575.1 | GT14 | | Glycosyltransferase Family 14 protein | | |
| XP_026273591.1 | GH13_17 | | Glycoside Hydrolase Family 13 protein | | |
| XP_026273891.1 | GH30_1 | | Glycoside Hydrolase Family 30 protein | | |
| XP_026273905.1 | GT4 | | Glycosyltransferase Family 4 protein | | |
| XP_026273921.1 | GH89 | | Glycoside Hydrolase Family 89 protein | | |
| XP_026273946.1 | CBM14-CBM14-GH1 | | Carbohydrate-Binding Module Family 14 / Glycoside Hydrolase Family 1 protein | | |
| XP_026273952.1 | GT54 | | Glycosyltransferase Family 54 protein | | |
| XP_026273993.1 | GT25 | fragment C-term; | Glycosyltransferase Family 25 protein | | |
| XP_026274115.1 | PL1_4 | fragment C-term; | Polysaccharide Lyase Family 1 protein | HGT | Complex: Bacteria or Fungi |
| XP_026274121.1 | GH1 | | Glycoside Hydrolase Family 1 protein | | |
| XP_026274148.1 | GT25 | fragment N-term; | Glycosyltransferase Family 25 protein | | |
| XP_026274343.1 | GT66 | (fragment) splicing problem; | Glycosyltransferase Family 66 protein | | |
| XP_026274619.1 | CBM14 | | Carbohydrate-Binding Module Family 14 protein | | |
| XP_026274622.1 | CBM14 | | Carbohydrate-Binding Module Family 14 protein | | |
| XP_026274677.1 | CBM14 | | Carbohydrate-Binding Module Family 14 protein | | |
| XP_026274686.1 | CBM14 | | Carbohydrate-Binding Module Family 14 protein | | |
| XP_026274851.1 | CBM14 | | Carbohydrate-Binding Module Family 14 protein | | |
| XP_026274864.1 | GT43 | | Glycosyltransferase Family 43 protein | | |
| XP_026274870.1 | GT27-CBM13 | | Glycosyltransferase Family 27 / Carbohydrate-Binding Module Family 13 protein | | |
| XP_026274904.1 | GT1 | | Glycosyltransferase Family 1 protein | | |
| XP_026274905.1 | GT1 | | Glycosyltransferase Family 1 protein | | |
| XP_026274906.1 | GT1 | | Glycosyltransferase Family 1 protein | | |
| XP_026274911.1 | GT1 | | Glycosyltransferase Family 1 protein | | |
| XP_026274916.1 | GT1-GT1-GT1 | | Glycosyltransferase Family 1 protein | | |
| XP_026275035.1 | CBM14 | | Carbohydrate-Binding Module Family 14 protein | | |
| XP_026275069.1 | CBM14 | | Carbohydrate-Binding Module Family 14 protein | | |
| XP_026275176.1 | GT16 | | Glycosyltransferase Family 16 protein | | |
| XP_026275221.1 | GT1 | | Glycosyltransferase Family 1 protein | | |
| XP_026275317.1 | GH31 | | Glycoside Hydrolase Family 31 protein | | |
| XP_026275448.1 | CBM14 | | Carbohydrate-Binding Module Family 14 protein | | |
| XP_026275456.1 | GT22 | | Glycosyltransferase Family 22 protein | | |
| XP_026275497.1 | GT1 | | Glycosyltransferase Family 1 protein | | |
| XP_026275702.1 | GT43 | | Glycosyltransferase Family 43 protein | | |
| XP_026275815.1 | GT61 | | Glycosyltransferase Family 61 protein | | |
| XP_026275888.1 | AA3_2 | | GMC oxidoreductase | | |
| XP_026276037.1 | GH30_1 | | Glycoside Hydrolase Family 30 protein | | |
| XP_026276202.1 | GT1 | | Glycosyltransferase Family 1 protein | | |
| XP_026276203.1 | GT1 | | Glycosyltransferase Family 1 protein | | |
| XP_026276262.1 | GT1 | | Glycosyltransferase Family 1 protein | | |

| | | | | | |
|---|---|---|---|---|---|
| XP_026276325.1 | GH30_1 | | Glycoside Hydrolase Family 30 protein | | |
| XP_026276440.1 | GH35 | | Glycoside Hydrolase Family 35 protein | | |
| XP_026276449.1 | GT49 | | Glycosyltransferase Family 49 protein | | |
| XP_026276465.1 | GH1 | | Glycoside Hydrolase Family 1 protein | | |
| XP_026276466.1 | GH1 | | Glycoside Hydrolase Family 1 protein | | |
| XP_026276495.1 | GT8 | | Glycosyltransferase Family 8 protein | | |
| XP_026276517.1 | GH20 | | Glycoside Hydrolase Family 20 protein | | |
| XP_026276583.1 | GH18 | fragment C-term; | Glycoside Hydrolase Family 18 protein | | |
| XP_026276646.1 | CBM50 | | Carbohydrate-Binding Module Family 50 protein | | |
| XP_026276666.1 | GH5_8 | | Glycoside Hydrolase Family 5 protein | HGT | Bacteria |
| XP_026276732.1 | AA3_2 | | GMC oxidoreductase | | |
| XP_026276765.1 | GT92 | | Glycosyltransferase Family 92 protein | | |
| XP_026276966.1 | GH56 | | Glycoside Hydrolase Family 56 protein | | |
| XP_026277104.1 | GH1 | | Glycoside Hydrolase Family 1 protein | | |
| XP_026277116.1 | GH1 | | Glycoside Hydrolase Family 1 protein | | |
| XP_026277129.1 | GH1 | | Glycoside Hydrolase Family 1 protein | | |
| XP_026277178.1 | GH1 | | Glycoside Hydrolase Family 1 protein | | |
| XP_026277263.1 | GT90 | | Glycosyltransferase Family 90 protein | | |
| XP_026277346.1 | GH1 | | Glycoside Hydrolase Family 1 protein | | |
| XP_026277353.1 | GH1 | | Glycoside Hydrolase Family 1 protein | | |
| XP_026277354.1 | GH1 | | Glycoside Hydrolase Family 1 protein | | |
| XP_026277504.1 | GT2 | | Glycosyltransferase Family 2 protein | | |
| XP_026277582.1 | GH13_25-GH133 | | Glycoside Hydrolase Family 13 / Glycoside Hydrolase Family 133 protein | | |
| XP_026277756.1 | GT2 | | Glycosyltransferase Family 2 protein | | |
| XP_026277826.1 | GT92 | | Glycosyltransferase Family 92 protein | | |
| XP_026277834.1 | GT57 | | Glycosyltransferase Family 57 protein | | |
| XP_026277861.1 | AA3_2 | | GMC oxidoreductase | | |
| XP_026277927.1 | GT65 | | Glycosyltransferase Family 65 protein | | |
| XP_026277980.1 | CE9 | | Carbohydrate Esterase Family 9 protein | | |
| XP_026277983.1 | GH1 | | Glycoside Hydrolase Family 1 protein | | |
| XP_026278042.1 | GT41 | | Glycosyltransferase Family 41 protein | | |
| XP_026278216.1 | GH1 | | Glycoside Hydrolase Family 1 protein | | |
| XP_026278238.1 | GT90 | | Glycosyltransferase Family 90 protein | | |
| XP_026278264.1 | GT31 | | Glycosyltransferase Family 31 protein | | |
| XP_026278300.1 | CBM21 | | Carbohydrate-Binding Module Family 21 protein | | |
| XP_026278401.1 | GT24 | | Glycosyltransferase Family 24 protein | | |
| XP_026278450.1 | GT27-CBM13 | | Glycosyltransferase Family 27 / Carbohydrate-Binding Module Family 13 protein | | |
| XP_026278473.1 | GT35 | | Glycosyltransferase Family 35 protein | | |
| XP_026278527.1 | GT10 | | Glycosyltransferase Family 10 protein | | |
| XP_026278599.1 | GH22 | | Glycoside Hydrolase Family 22 protein | | |
| XP_026278633.1 | GH2 | | Glycoside Hydrolase Family 2 protein | | |
| XP_026278638.1 | GH20 | (fragment) splicing problem; | Glycoside Hydrolase Family 20 protein | | |
| XP_026278691.1 | GH20 | | Glycoside Hydrolase Family 20 protein | | |
| XP_026278843.1 | CBM14 | | Carbohydrate-Binding Module Family 14 protein | | |
| XP_026278853.1 | GT57 | | Glycosyltransferase Family 57 protein | | |
| XP_026278876.1 | AA15 | | Auxilliary Activities Family 15 protein | | |
| XP_026278917.1 | AA3_2 | | GMC oxidoreductase | | |
| XP_026278952.1 | GH9 | | Glycoside Hydrolase Family 9 protein | | |
| XP_026279015.1 | CBM14-CBM14-CBM14-CBM14-CBM14-CBM14-CBM14 | | Carbohydrate-Binding Module Family 14 protein | | |

| | | | | | |
|---|---|---|---|---|---|
| XP_026279096.1 | GT2 | | Glycosyltransferase Family 2 protein | | |
| XP_026279162.1 | GH84 | | Glycoside Hydrolase Family 84 protein | | |
| XP_026279182.1 | GH31 | | Glycoside Hydrolase Family 31 protein | | |
| XP_026279195.1 | AA1 | | Multicopper oxidase | | |
| XP_026279277.1 | GH1 | fragment N-term; | Glycoside Hydrolase Family 1 protein | | |
| XP_026279283.1 | CBM14-CBM14-GH1 | | Carbohydrate-Binding Module Family 14 / Glycoside Hydrolase Family 1 protein | | |
| XP_026279337.1 | GT54 | (fragment) splicing problem; | Glycosyltransferase Family 54 protein | | |
| XP_026279359.1 | GH18-GH18-CBM14 | | Glycoside Hydrolase Family 18 / Carbohydrate-Binding Module Family 14 protein | | |
| XP_026279375.1 | GT27-CBM13 | | Glycosyltransferase Family 27 / Carbohydrate-Binding Module Family 13 protein | | |
| XP_026279513.1 | PL1_4 | (fragment) splicing problem; | Polysaccharide Lyase Family 1 protein | HGT | Complex: Bacteria or Fungi |
| XP_026279528.1 | PL1_4 | (fragment) splicing problem; | Polysaccharide Lyase Family 1 protein | HGT | Complex: Bacteria or Fungi |
| XP_026279697.1 | GT1 | | Glycosyltransferase Family 1 protein | | |
| XP_026279860.1 | GT1 | | Glycosyltransferase Family 1 protein | | |
| XP_026280023.1 | GT31 | | Glycosyltransferase Family 31 protein | | |
| XP_026280167.1 | GT8 | (fragment) splicing problem; | Glycosyltransferase Family 8 protein | | |
| XP_026280209.1 | GT23 | | Glycosyltransferase Family 23 protein | | |
| XP_026280340.1 | GH13_17 | | Glycoside Hydrolase Family 13 protein | | |
| XP_026280345.1 | GH13_17 | | Glycoside Hydrolase Family 13 protein | | |
| XP_026280363.1 | AA3_2 | | GMC oxidoreductase | | |
| XP_026280366.1 | AA3_2 | | GMC oxidoreductase | | |
| XP_026280374.1 | CBM14-CBM14 | | Carbohydrate-Binding Module Family 14 protein | | |
| XP_026280376.1 | GT66 | (fragment) splicing problem; | Glycosyltransferase Family 66 protein | | |
| XP_026280402.1 | PL1_4 | (fragment) splicing problem; | Polysaccharide Lyase Family 1 protein | HGT | Complex: Bacteria or Fungi |
| XP_026280504.1 | GH20 | | Glycoside Hydrolase Family 20 protein | | |
| XP_026280640.1 | CBM47 | | Carbohydrate-Binding Module Family 47 protein | | |
| XP_026280652.1 | GH20 | | Glycoside Hydrolase Family 20 protein | | |
| XP_026280712.1 | GH79 | | Glycoside Hydrolase Family 79 protein | | |
| XP_026280824.1 | GH31 | (fragment) splicing problem; | Glycoside Hydrolase Family 31 protein | | |
| XP_026280852.1 | AA1 | | Multicopper oxidase | | |
| XP_026280854.1 | GT7 | | Glycosyltransferase Family 7 protein | | |
| XP_026280905.1 | GT98 | | Glycosyltransferase Family 98 protein | | |
| XP_026280965.1 | GH47 | | Glycoside Hydrolase Family 47 protein | | |
| XP_026280995.1 | GT31 | (fragment) splicing problem; | Glycosyltransferase Family 31 protein | | |
| XP_026281016.1 | GT68 | | Glycosyltransferase Family 68 protein | | |
| XP_026281037.1 | GT21 | | Glycosyltransferase Family 21 protein | | |
| XP_026281231.1 | CBM14 | | Carbohydrate-Binding Module Family 14 protein | | |
| XP_026281265.1 | GH1 | fragment C-term; | Glycoside Hydrolase Family 1 protein | | |
| XP_026281402.1 | GH20 | | Glycoside Hydrolase Family 20 protein | | |
| XP_026281449.1 | GT33 | | Glycosyltransferase Family 33 protein | | |
| XP_026281495.1 | GT13 | | Glycosyltransferase Family 13 protein | | |
| XP_026281552.1 | AA3_2 | | GMC oxidoreductase | | |
| XP_026281601.1 | CBM48 | | Carbohydrate-Binding Module Family 48 protein | | |
| XP_026281774.1 | GT4 | | Glycosyltransferase Family 4 protein | | |
| XP_026281838.1 | AA3_2 | | GMC oxidoreductase | | |
| XP_026281850.1 | GT31 | | Glycosyltransferase Family 31 protein | | |
| XP_026281861.1 | GT3 | | Glycosyltransferase Family 3 protein | | |
| XP_026282077.1 | GT105 | | Glycosyltransferase Family 105 protein | | |
| XP_026282345.1 | GT59 | | Glycosyltransferase Family 59 protein | | |
| XP_026282369.1 | GH18-CBM14-CBM14-GH18-CBM14-CBM14-CBM14-GH18-GH18-CBM14-GH18 | | Glycoside Hydrolase Family 18 / Carbohydrate-Binding Module Family 14 protein | | |

| | | | | | |
|---|---|---|---|---|---|
| XP_026282505.1 | GH79 | | Glycoside Hydrolase Family 79 protein | | |
| XP_026282691.1 | GH1 | | Glycoside Hydrolase Family 1 protein | | |
| XP_026282723.1 | GT27-CBM13 | fragment C-term; | Glycosyltransferase Family 27 / Carbohydrate-Binding Module Family 13 protein | | |
| XP_026282959.1 | GT8 | | Glycosyltransferase Family 8 protein | | |
| XP_026283009.1 | GT49 | (fragment) splicing problem; | Glycosyltransferase Family 49 protein | | |
| XP_026283169.1 | GH27 | | Glycoside Hydrolase Family 27 protein | | |
| XP_026283285.1 | GH22 | | Glycoside Hydrolase Family 22 protein | | |
| XP_026283358.1 | GH1 | | Glycoside Hydrolase Family 1 protein | | |
| XP_026283506.1 | CBM14 | | Carbohydrate-Binding Module Family 14 protein | | |
| XP_026283507.1 | CBM14 | | Carbohydrate-Binding Module Family 14 protein | | |
| XP_026283531.1 | GH47 | | Glycoside Hydrolase Family 47 protein | | |
| XP_026283562.1 | GH18-CBM14-CBM14 | (fragment) splicing problem; | Glycoside Hydrolase Family 18 / Carbohydrate-Binding Module Family 14 protein | | |
| XP_026283629.1 | GT76 | | Glycosyltransferase Family 76 protein | | |
| XP_026283655.1 | GH47 | | Glycoside Hydrolase Family 47 protein | | |
| XP_026283824.1 | GT39 | | Glycosyltransferase Family 39 protein | | |
| XP_026283909.1 | GT1 | | Glycosyltransferase Family 1 protein | | |
| XP_026283914.1 | GH1 | | Glycoside Hydrolase Family 1 protein | | |
| XP_026284071.1 | GH1 | | Glycoside Hydrolase Family 1 protein | | |
| XP_026284119.1 | GT31 | (fragment) splicing problem; | Glycosyltransferase Family 31 protein | | |
| XP_026284142.1 | CBM20 | | Carbohydrate-Binding Module Family 20 protein | | |
| XP_026284159.1 | GH13_15 | | Glycoside Hydrolase Family 13 protein | | |
| XP_026284171.1 | GT31 | (fragment) splicing problem; | Glycosyltransferase Family 31 protein | | |
| XP_026284183.1 | CBM47 | | Carbohydrate-Binding Module Family 47 protein | | |
| XP_026284196.1 | AA3_2 | | GMC oxidoreductase | | |
| XP_026284470.1 | AA3_2 | | GMC oxidoreductase | | |
| XP_026284490.1 | GH1 | | Glycoside Hydrolase Family 1 protein | | |
| XP_026284494.1 | GH1 | | Glycoside Hydrolase Family 1 protein | | |
| XP_026284527.1 | GT22 | | Glycosyltransferase Family 22 protein | | |
| XP_026284547.1 | GT4 | | Glycosyltransferase Family 4 protein | | |
| XP_026284668.1 | GH22 | | Glycoside Hydrolase Family 22 protein | | |
| XP_026284670.1 | AA3_2 | | GMC oxidoreductase | | |
| XP_026284729.1 | AA3_2 | | GMC oxidoreductase | | |
| XP_026284910.1 | CBM14 | | Carbohydrate-Binding Module Family 14 protein | | |
| XP_026284999.1 | GH2 | | Glycoside Hydrolase Family 2 protein | | |
| XP_026285006.1 | GH16_4 | | Glycoside Hydrolase Family 16 protein | | |
| XP_026285007.1 | CBM57 | | Carbohydrate-Binding Module Family 57 protein | | |
| XP_026285066.1 | PL1_4 | (fragment) splicing problem; | Polysaccharide Lyase Family 1 protein | HGT | Complex: Bacteria or Fungi |
| XP_026285067.1 | PL1_4 | (fragment) splicing problem; | Polysaccharide Lyase Family 1 protein | HGT | Complex: Bacteria or Fungi |
| XP_026285092.1 | AA3_2 | | GMC oxidoreductase | | |
| XP_026285262.1 | GH1 | | Glycoside Hydrolase Family 1 protein | | |
| XP_026285289.1 | GH5_8 | | Glycoside Hydrolase Family 5 protein | HGT | Bacteria |
| XP_026285291.1 | GH5_8 | | Glycoside Hydrolase Family 5 protein | HGT | Bacteria |
| XP_026285315.1 | GT47-GT64 | | Glycosyltransferase Family 47 / Glycosyltransferase Family 64 protein | | |
| XP_026285319.1 | GH99 | | Glycoside Hydrolase Family 99 protein | | |
| XP_026285326.1 | GH18 | | Glycoside Hydrolase Family 18 protein | | |
| XP_026285426.1 | GT27-CBM13 | | Glycosyltransferase Family 27 / Carbohydrate-Binding Module Family 13 protein | | |
| XP_026285441.1 | GT27-CBM13 | | Glycosyltransferase Family 27 / Carbohydrate-Binding Module Family 13 protein | | |
| XP_026285502.1 | GH1 | | Glycoside Hydrolase Family 1 protein | | |
| XP_026285581.1 | GH1 | | Glycoside Hydrolase Family 1 protein | | |

| | | | | | | |
|---|---|---|---|---|---|---|
| XP_026285608.1 | GT110 | | Glycosyltransferase Family 110 protein | | | |
| XP_026285692.1 | GH1 | | Glycoside Hydrolase Family 1 protein | | | |
| XP_026285787.1 | GH1 | | Glycoside Hydrolase Family 1 protein | | | |
| XP_026285857.1 | GH85 | (fragment) splicing problem; | Glycoside Hydrolase Family 85 protein | | | |
| XP_026285883.1 | PL1_4 | (fragment) splicing problem; | Polysaccharide Lyase Family 1 protein | | HGT | Complex: Bacteria or Fungi |
| XP_026285908.1 | CBM14-CBM14-CBM14 | | Carbohydrate-Binding Module Family 14 protein | | | |
| XP_026285962.1 | GT105 | | Glycosyltransferase Family 105 protein | | | |
| XP_026285994.1 | GH47 | (fragment) splicing problem; | Glycoside Hydrolase Family 47 protein | | | |
| XP_026286003.1 | AA1 | | Multicopper oxidase | | | |
| XP_026286031.1 | GH2 | | Glycoside Hydrolase Family 2 protein | | | |
| XP_026286161.1 | GT2 | | Glycosyltransferase Family 2 protein | | | |
| XP_026286293.1 | GT31 | | Glycosyltransferase Family 31 protein | | | |
| XP_026286356.1 | AA3_2 | | GMC oxidoreductase | | | |
| XP_026286505.1 | GT92 | | Glycosyltransferase Family 92 protein | | | |
| XP_026286599.1 | GH1 | | Glycoside Hydrolase Family 1 protein | | | |
| XP_026286611.1 | AA3_2 | (fragment) splicing problem; | GMC oxidoreductase | | | |
| XP_026286629.1 | GH1 | fragment C-term; | Glycoside Hydrolase Family 1 protein | | | |
| XP_026286630.1 | GH1 | fragment N-term; | Glycoside Hydrolase Family 1 protein | | | |
| XP_026286632.1 | GH1 | fragment N-term / C-term; | Glycoside Hydrolase Family 1 protein | | | |
| XP_026286772.1 | CBM14 | fragment N-term; | Carbohydrate-Binding Module Family 14 protein | | | |
| XP_026286922.1 | GH18-CBM14-CBM14-CBM14 | (fragment) splicing problem; | Glycoside Hydrolase Family 18 / Carbohydrate-Binding Module Family 14 protein | | | |
| XP_026286949.1 | GH16_4 | | Glycoside Hydrolase Family 16 protein | | | |
| XP_026286975.1 | CBM14 | | Carbohydrate-Binding Module Family 14 protein | | | |
| XP_026287231.1 | GH1 | (fragment) splicing problem; | Glycoside Hydrolase Family 1 protein | | | |
| XP_026287312.1 | GT7 | | Glycosyltransferase Family 7 protein | | | |
| XP_026287319.1 | GH22 | | Glycoside Hydrolase Family 22 protein | | | |
| XP_026287364.1 | GH1 | | Glycoside Hydrolase Family 1 protein | | | |
| XP_026287407.1 | GH22 | | Glycoside Hydrolase Family 22 protein | | | |
| XP_026287599.1 | GH1 | fragment C-term; | Glycoside Hydrolase Family 1 protein | | | |
| XP_026287603.1 | GH22-GH22 | | Glycoside Hydrolase Family 22 protein | | | |
| XP_026287675.1 | CBM14 | | Carbohydrate-Binding Module Family 14 protein | | | |
| XP_026287748.1 | GT13 | | Glycosyltransferase Family 13 protein | | | |
| XP_026287789.1 | AA3_2 | (fragment) splicing problem; | GMC oxidoreductase | | | |
| XP_026287851.1 | GH32 | | Glycoside Hydrolase Family 32 protein | | HGT | Bacteria |
| XP_026287983.1 | GH1 | | Glycoside Hydrolase Family 1 protein | | | |
| XP_026287984.1 | GH45 | | Glycoside Hydrolase Family 45 protein | | HGT | Fungi |
| XP_026287990.1 | GT105 | | Glycosyltransferase Family 105 protein | | | |
| XP_026288136.1 | GH1 | | Glycoside Hydrolase Family 1 protein | | | |
| XP_026288193.1 | GH1 | | Glycoside Hydrolase Family 1 protein | | | |
| XP_026288229.1 | GH39 | | Glycoside Hydrolase Family 39 protein | | | |
| XP_026288262.1 | CBM14 | | Carbohydrate-Binding Module Family 14 protein | | | |
| XP_026288499.1 | GH1 | | Glycoside Hydrolase Family 1 protein | | | |
| XP_026288501.1 | GH37 | | Glycoside Hydrolase Family 37 protein | | | |
| XP_026288506.1 | GH37 | | Glycoside Hydrolase Family 37 protein | | | |
| XP_026288543.1 | AA3_2 | fragment N-term; | GMC oxidoreductase | | | |
| XP_026288627.1 | CBM14 | | Carbohydrate-Binding Module Family 14 protein | | | |
| XP_026288628.1 | CBM14-CBM14-CBM14-CBM14-CBM14-CBM14 | | Carbohydrate-Binding Module Family 14 protein | | | |
| XP_026288642.1 | CBM14-CBM14-CBM14 | | Carbohydrate-Binding Module Family 14 protein | | | |
| XP_026288646.1 | CBM14-CBM14-CBM14 | | Carbohydrate-Binding Module Family 14 protein | | | |

| | | | | | |
|---|---|---|---|---|---|
| XP_026288705.1 | CBM14-CBM14-CBM14 | | Carbohydrate-Binding Module Family 14 protein | | |
| XP_026288706.1 | CBM14-CBM14-CBM14 | | Carbohydrate-Binding Module Family 14 protein | | |
| XP_026288707.1 | CBM14-CBM14-CBM14 | | Carbohydrate-Binding Module Family 14 protein | | |
| XP_026288737.1 | GT32 | | Glycosyltransferase Family 32 protein | | |
| XP_026288839.1 | CBM14-CBM14-GH1 | | Carbohydrate-Binding Module Family 14 / Glycoside Hydrolase Family 1 protein | | |
| XP_026288857.1 | GT22 | | Glycosyltransferase Family 22 protein | | |
| XP_026288943.1 | GH1 | (fragment) splicing problem; | Glycoside Hydrolase Family 1 protein | | |
| XP_026288944.1 | GH1 | fragment N-term; | Glycoside Hydrolase Family 1 protein | | |
| XP_026288982.1 | GH1 | | Glycoside Hydrolase Family 1 protein | | |
| XP_026289055.1 | GH20 | | Glycoside Hydrolase Family 20 protein | | |
| XP_026289145.1 | GH22 | fragment N-term; | Glycoside Hydrolase Family 22 protein | | |
| XP_026289264.1 | GH45 | | Glycoside Hydrolase Family 45 protein | HGT | Fungi |
| XP_026289265.1 | GH22 | | Glycoside Hydrolase Family 22 protein | | |
| XP_026289306.1 | AA3_2 | | GMC oxidoreductase | | |
| XP_026289307.1 | AA3_2 | | GMC oxidoreductase | | |
| XP_026289309.1 | AA3_2 | | GMC oxidoreductase | | |
| XP_026289310.1 | AA3_2 | | GMC oxidoreductase | | |
| XP_026289312.1 | AA3_2 | | GMC oxidoreductase | | |
| XP_026289319.1 | AA3_2-AA3_2 | | GMC oxidoreductase | | |
| XP_026289325.1 | AA3_2-AA3_2-AA3_2-AA3_2 | | GMC oxidoreductase | | |
| XP_026289345.1 | GT61 | | Glycosyltransferase Family 61 protein | | |
| XP_026289476.1 | AA3_2 | | GMC oxidoreductase | | |
| XP_026289490.1 | AA3_2 | fragment C-term; | GMC oxidoreductase | | |
| XP_026289493.1 | AA3_2 | | GMC oxidoreductase | | |
| XP_026289521.1 | GH1 | fragment N-term / C-term; | Glycoside Hydrolase Family 1 protein | | |
| XP_026289532.1 | GT31 | | Glycosyltransferase Family 31 protein | | |
| XP_026289562.1 | CBM14 | | Carbohydrate-Binding Module Family 14 protein | | |
| XP_026289622.1 | GH1 | | Glycoside Hydrolase Family 1 protein | | |
| XP_026289641.1 | PL1_4 | (fragment) splicing problem; | Polysaccharide Lyase Family 1 protein | HGT | Complex: Bacteria or Fungi |
| XP_026289642.1 | GH1 | fragment N-term; | Glycoside Hydrolase Family 1 protein | | |
| XP_026289643.1 | AA3_2 | | GMC oxidoreductase | | |
| XP_026289679.1 | CBM39-GH16_4 | (fragment) splicing problem; | Carbohydrate-Binding Module Family 39 / Glycoside Hydrolase Family 16 protein | | |
| XP_026289810.1 | GH1 | | Glycoside Hydrolase Family 1 protein | | |
| XP_026289832.1 | GH1 | fragment N-term / C-term; | Glycoside Hydrolase Family 1 protein | | |
| XP_026289894.1 | GT1 | fragment N-term / C-term; | Glycosyltransferase Family 1 protein | | |
| XP_026289900.1 | GH1 | fragment N-term; | Glycoside Hydrolase Family 1 protein | | |
| XP_026290001.1 | CBM48-GH13_8 | | Carbohydrate-Binding Module Family 48 / Glycoside Hydrolase Family 13 protein | | |
| XP_026290004.1 | AA3_2 | fragment C-term; | GMC oxidoreductase | | |
| XP_026290091.1 | GT31-GT7 | | Glycosyltransferase Family 31 / Glycosyltransferase Family 7 protein | | |
| XP_026290108.1 | GH38 | fragment N-term / C-term; | Glycoside Hydrolase Family 38 protein | | |
| XP_026290110.1 | GT1 | | Glycosyltransferase Family 1 protein | | |
| XP_026290116.1 | GH1 | | Glycoside Hydrolase Family 1 protein | | |
| XP_026290124.1 | GH1 | fragment C-term; | Glycoside Hydrolase Family 1 protein | | |
| XP_026290280.1 | GT1 | | Glycosyltransferase Family 1 protein | | |
| XP_026290293.1 | GH1 | | Glycoside Hydrolase Family 1 protein | | |
| XP_026290315.1 | GT50 | | Glycosyltransferase Family 50 protein | | |
| XP_026290324.1 | GT1 | | Glycosyltransferase Family 1 protein | | |
| XP_026290338.1 | AA3_2 | | GMC oxidoreductase | | |
| XP_026290383.1 | GH152 | | Glycoside Hydrolase Family 152 protein | HGT | Viridiplantae |

| | | | | | |
|---|---|---|---|---|---|
| XP_026290470.1 | AA3_2 | | GMC oxidoreductase | | |
| XP_026290513.1 | AA1 | | Multicopper oxidase | | |
| XP_026290541.1 | CBM14-CBM14 | | Carbohydrate-Binding Module Family 14 protein | | |
| XP_026290559.1 | AA15 | | Auxilliary Activities Family 15 protein | | |
| XP_026290580.1 | GT1 | fragment N-term; | Glycosyltransferase Family 1 protein | | |
| XP_026290752.1 | GH63 | | Glycoside Hydrolase Family 63 protein | | |
| XP_026290776.1 | GH1 | | Glycoside Hydrolase Family 1 protein | | |
| XP_026290795.1 | GH29 | | Glycoside Hydrolase Family 29 protein | | |
| XP_026290983.1 | GH1 | | Glycoside Hydrolase Family 1 protein | | |
| XP_026291029.1 | GH1 | fragment C-term; | Glycoside Hydrolase Family 1 protein | | |
| XP_026291031.1 | GH22 | | Glycoside Hydrolase Family 22 protein | | |
| XP_026291034.1 | GH1 | fragment N-term; | Glycoside Hydrolase Family 1 protein | | |
| XP_026291102.1 | CBM14 | | Carbohydrate-Binding Module Family 14 protein | | |
| XP_026291188.1 | GH1 | fragment C-term; | Glycoside Hydrolase Family 1 protein | | |
| XP_026291249.1 | GT1 | | Glycosyltransferase Family 1 protein | | |
| XP_026291611.1 | GT1 | | Glycosyltransferase Family 1 protein | | |
| XP_026291613.1 | GH1 | | Glycoside Hydrolase Family 1 protein | | |
| XP_026291692.1 | GH18 | | Glycoside Hydrolase Family 18 protein | | |
| XP_026291724.1 | GT16 | fragment N-term; | Glycosyltransferase Family 16 protein | | |
| XP_026291832.1 | AA3_2 | fragment N-term; | GMC oxidoreductase | | |
| XP_026291890.1 | GH45 | (fragment) splicing problem; | Glycoside Hydrolase Family 45 protein | HGT | Fungi |
| XP_026291892.1 | GH18 | fragment N-term; | Glycoside Hydrolase Family 18 protein | | |
| XP_026291972.1 | GH31 | | Glycoside Hydrolase Family 31 protein | | |
| XP_026291995.1 | GT64 | | Glycosyltransferase Family 64 protein | | |
| XP_026292004.1 | CBM14 | | Carbohydrate-Binding Module Family 14 protein | | |
| XP_026292060.1 | GH1 | fragment N-term / C-term; | Glycoside Hydrolase Family 1 protein | | |
| XP_026292146.1 | GT1 | fragment C-term; | Glycosyltransferase Family 1 protein | | |
| XP_026292176.1 | GT1 | | Glycosyltransferase Family 1 protein | | |
| XP_026292251.1 | GT1 | | Glycosyltransferase Family 1 protein | | |
| XP_026292396.1 | PL1_4 | (fragment) splicing problem; | Polysaccharide Lyase Family 1 protein | HGT | Complex: Bacteria or Fungi |
| XP_026292397.1 | PL1_4 | (fragment) splicing problem; | Polysaccharide Lyase Family 1 protein | HGT | Complex: Bacteria or Fungi |
| XP_026292409.1 | GT1 | fragment C-term; | Glycosyltransferase Family 1 protein | | |
| XP_026292436.1 | AA3_2 | fragment C-term; | GMC oxidoreductase | | |
| XP_026292490.1 | CBM39 | fragment N-term; | Carbohydrate-Binding Module Family 39 protein | | |
| XP_026292503.1 | AA1 | fragment N-term; | Multicopper oxidase | | |
| XP_026292528.1 | GH1 | fragment N-term; | Glycoside Hydrolase Family 1 protein | | |
| XP_026292714.1 | CBM14 | | Carbohydrate-Binding Module Family 14 protein | | |
| XP_026292767.1 | GT43 | fragment N-term / C-term; | Glycosyltransferase Family 43 protein | | |
| XP_026292919.1 | GH116 | | Glycoside Hydrolase Family 116 protein | | |
| XP_026292938.1 | GH1 | | Glycoside Hydrolase Family 1 protein | | |
| XP_026292945.1 | GH22 | | Glycoside Hydrolase Family 22 protein | | |
| XP_026293029.1 | GT31 | | Glycosyltransferase Family 31 protein | | |
| XP_026293209.1 | AA1 | | Multicopper oxidase | | |
| XP_026293310.1 | AA3_2 | fragment N-term; | GMC oxidoreductase | | |
| XP_026293399.1 | CE13 | | Carbohydrate Esterase Family 13 protein | | |
| XP_026293405.1 | GH38 | (fragment) splicing problem; | Glycoside Hydrolase Family 38 protein | | |
| XP_026293426.1 | GH18 | | Glycoside Hydrolase Family 18 protein | | |
| XP_026293490.1 | CBM14-CBM14-CBM14-CBM14 | | Carbohydrate-Binding Module Family 14 protein | | |
| XP_026293740.1 | GH1 | | Glycoside Hydrolase Family 1 protein | | |

| | | | |
|---|---|---|---|
| XP_026293776.1 | CBM14 | | Carbohydrate-Binding Module Family 14 protein |
| XP_026293885.1 | AA3_2 | | GMC oxidoreductase |
| XP_026293897.1 | GH47 | | Glycoside Hydrolase Family 47 protein |
| XP_026294185.1 | GT1 | | Glycosyltransferase Family 1 protein |
| XP_026294279.1 | GT29 | | Glycosyltransferase Family 29 protein |
| XP_026294292.1 | GT31 | | Glycosyltransferase Family 31 protein |
| XP_026294370.1 | AA1 | | Multicopper oxidase |
| XP_026294395.1 | GT1 | | Glycosyltransferase Family 1 protein |
| XP_026294449.1 | GT27-CBM13 | (fragment) splicing problem; | Glycosyltransferase Family 27 / Carbohydrate-Binding Module Family 13 protein |
| XP_026294573.1 | GT31 | fragment N-term; | Glycosyltransferase Family 31 protein |
| XP_026294605.1 | GT49 | | Glycosyltransferase Family 49 protein |

*T. palmi*

| **Protein ID** | **Description** | **Model notes** | **Definition line** | **HGT** | **Donor** |
|---|---|---|---|---|---|
| XP_034229989.1 | GH1 | fragment N-term; | Glycoside Hydrolase Family 1 protein | | |
| XP_034230049.1 | CBM14 | | Carbohydrate-Binding Module Family 14 protein | | |
| XP_034230158.1 | GT1 | | Glycosyltransferase Family 1 protein | | |
| XP_034230228.1 | AA3_2 | | GMC oxidoreductase | | |
| XP_034230230.1 | AA3_2 | | GMC oxidoreductase | | |
| XP_034230315.1 | GT105 | | Glycosyltransferase Family 105 protein | | |
| XP_034230380.1 | CBM14-CBM14 | | Carbohydrate-Binding Module Family 14 protein | | |
| XP_034230391.1 | GT13 | | Glycosyltransferase Family 13 protein | | |
| XP_034230419.1 | GT2 | | Glycosyltransferase Family 2 protein | | |
| XP_034230455.1 | CBM57 | | Carbohydrate-Binding Module Family 57 protein | | |
| XP_034230479.1 | GT61 | | Glycosyltransferase Family 61 protein | | |
| XP_034230536.1 | AA15 | | Auxilliary Activities Family 15 protein | | |
| XP_034230692.1 | GH2 | | Glycoside Hydrolase Family 2 protein | | |
| XP_034230730.1 | GH1 | | Glycoside Hydrolase Family 1 protein | | |
| XP_034230753.1 | GH22 | | Glycoside Hydrolase Family 22 protein | | |
| XP_034230762.1 | GT2 | | Glycosyltransferase Family 2 protein | | |
| XP_034230845.1 | AA3_2 | | GMC oxidoreductase | | |
| XP_034230866.1 | GT43 | | Glycosyltransferase Family 43 protein | | |
| XP_034230892.1 | GT65 | | Glycosyltransferase Family 65 protein | | |
| XP_034230949.1 | AA3_2 | | GMC oxidoreductase | | |
| XP_034230950.1 | AA3_2 | | GMC oxidoreductase | | |
| XP_034231003.1 | GT1 | | Glycosyltransferase Family 1 protein | | |
| XP_034231019.1 | GT92 | | Glycosyltransferase Family 92 protein | | |
| XP_034231040.1 | AA3_2-AA3_2 | | GMC oxidoreductase | | |
| XP_034231041.1 | AA3_2 | | GMC oxidoreductase | | |
| XP_034231075.1 | GH22 | | Glycoside Hydrolase Family 22 protein | | |
| XP_034231185.1 | CBM14 | | Carbohydrate-Binding Module Family 14 protein | | |
| XP_034231187.1 | CBM14 | | Carbohydrate-Binding Module Family 14 protein | | |
| XP_034231281.1 | AA1 | | Multicopper oxidase | | |
| XP_034231304.1 | GT1 | | Glycosyltransferase Family 1 protein | | |
| XP_034231365.1 | GH37 | | Glycoside Hydrolase Family 37 protein | | |
| XP_034231424.1 | AA3_2-AA3_2 | | GMC oxidoreductase | | |
| XP_034231426.1 | GT13 | | Glycosyltransferase Family 13 protein | | |
| XP_034231448.1 | GH16_4 | | Glycoside Hydrolase Family 16 protein | | |

| | | | | | |
|---|---|---|---|---|---|
| XP_034231449.1 | AA1 | | Multicopper oxidase | | |
| XP_034231457.1 | GT31 | | Glycosyltransferase Family 31 protein | | |
| XP_034231710.1 | CBM47 | | Carbohydrate-Binding Module Family 47 protein | | |
| XP_034231727.1 | GT41 | | Glycosyltransferase Family 41 protein | | |
| XP_034231998.1 | GT105 | | Glycosyltransferase Family 105 protein | | |
| XP_034232030.1 | GH20 | | Glycoside Hydrolase Family 20 protein | | |
| XP_034232055.1 | CBM47 | | Carbohydrate-Binding Module Family 47 protein | | |
| XP_034232152.1 | GH1 | | Glycoside Hydrolase Family 1 protein | | |
| XP_034232312.1 | GT31 | | Glycosyltransferase Family 31 protein | | |
| XP_034232383.1 | GH32 | | Glycoside Hydrolase Family 32 protein | HGT | Bacteria |
| XP_034232398.1 | GH1 | | Glycoside Hydrolase Family 1 protein | | |
| XP_034232426.1 | AA3_2 | | GMC oxidoreductase | | |
| XP_034232579.1 | AA3_2 | fragment N-term; | GMC oxidoreductase | | |
| XP_034232815.1 | GT1 | | Glycosyltransferase Family 1 protein | | |
| XP_034232830.1 | GH1 | fragment N-term; | Glycoside Hydrolase Family 1 protein | | |
| XP_034232847.1 | GT49 | | Glycosyltransferase Family 49 protein | | |
| XP_034232873.1 | GT31 | | Glycosyltransferase Family 31 protein | | |
| XP_034232917.1 | AA1 | | Multicopper oxidase | | |
| XP_034232935.1 | GH22-GH22 | | Glycoside Hydrolase Family 22 protein | | |
| XP_034233104.1 | GH13_15 | | Glycoside Hydrolase Family 13 protein | | |
| XP_034233350.1 | GH45 | (fragment) splicing problem; | Glycoside Hydrolase Family 45 protein | HGT | Fungi |
| XP_034233351.1 | AA3_2 | fragment N-term; | GMC oxidoreductase | | |
| XP_034233420.1 | GH32 | | Glycoside Hydrolase Family 32 protein | HGT | Bacteria |
| XP_034233461.1 | GH35 | | Glycoside Hydrolase Family 35 protein | | |
| XP_034233657.1 | GT8 | (fragment) splicing problem; | Glycosyltransferase Family 8 protein | | |
| XP_034233979.1 | CBM14-CBM14-CBM14 | | Carbohydrate-Binding Module Family 14 protein | | |
| XP_034233988.1 | GH1 | | Glycoside Hydrolase Family 1 protein | | |
| XP_034234045.1 | CBM14-CBM14 | | Carbohydrate-Binding Module Family 14 protein | | |
| XP_034234061.1 | CBM14 | | Carbohydrate-Binding Module Family 14 protein | | |
| XP_034234070.1 | GT49 | | Glycosyltransferase Family 49 protein | | |
| XP_034234073.1 | GH39 | fragment N-term; | Glycoside Hydrolase Family 39 protein | | |
| XP_034234087.1 | GH18 | | Glycoside Hydrolase Family 18 protein | | |
| XP_034234108.1 | GH20 | | Glycoside Hydrolase Family 20 protein | | |
| XP_034234142.1 | CBM14-CBM14-CBM14 | | Carbohydrate-Binding Module Family 14 protein | | |
| XP_034234196.1 | AA3_2 | | GMC oxidoreductase | | |
| XP_034234238.1 | CBM14-CBM14-CBM14 | | Carbohydrate-Binding Module Family 14 protein | | |
| XP_034234307.1 | GH1 | | Glycoside Hydrolase Family 1 protein | | |
| XP_034234389.1 | GH35 | | Glycoside Hydrolase Family 35 protein | | |
| XP_034234464.1 | GH20 | | Glycoside Hydrolase Family 20 protein | | |
| XP_034234651.1 | GT43 | | Glycosyltransferase Family 43 protein | | |
| XP_034234691.1 | CBM14-CBM14-CBM14 | | Carbohydrate-Binding Module Family 14 protein | | |
| XP_034234754.1 | GT1 | | Glycosyltransferase Family 1 protein | | |
| XP_034234765.1 | CBM14-CBM14-CBM14 | | Carbohydrate-Binding Module Family 14 protein | | |
| XP_034234918.1 | CBM14-CBM14-CBM14 | | Carbohydrate-Binding Module Family 14 protein | | |
| XP_034234920.1 | CBM14-CBM14-CBM14 | | Carbohydrate-Binding Module Family 14 protein | | |
| XP_034234999.1 | AA3_2 | fragment N-term; | GMC oxidoreductase | | |
| XP_034235276.1 | CBM14-CBM14 | | Carbohydrate-Binding Module Family 14 protein | | |
| XP_034235278.1 | GH56 | | Glycoside Hydrolase Family 56 protein | | |
| XP_034235375.1 | GT16 | | Glycosyltransferase Family 16 protein | | |

| | | | | | |
|---|---|---|---|---|---|
| XP_034235409.1 | GH20 | (fragment) splicing problem; | Glycoside Hydrolase Family 20 protein | | |
| XP_034235640.1 | PL1_4 | (fragment) splicing problem; | Polysaccharide Lyase Family 1 protein | HGT | Complex: Bacteria or Fungi |
| XP_034235648.1 | GH1 | | Glycoside Hydrolase Family 1 protein | | |
| XP_034235670.1 | GH1 | fragment C-term; | Glycoside Hydrolase Family 1 protein | | |
| XP_034235714.1 | GT1 | | Glycosyltransferase Family 1 protein | | |
| XP_034235752.1 | GH22 | | Glycoside Hydrolase Family 22 protein | | |
| XP_034235817.1 | GT2 | | Glycosyltransferase Family 2 protein | | |
| XP_034235838.1 | GT90 | | Glycosyltransferase Family 90 protein | | |
| XP_034235996.1 | CBM14 | | Carbohydrate-Binding Module Family 14 protein | | |
| XP_034236588.1 | GH45 | | Glycoside Hydrolase Family 45 protein | HGT | Fungi |
| XP_034236620.1 | AA3_2 | | GMC oxidoreductase | | |
| XP_034236632.1 | GH1 | | Glycoside Hydrolase Family 1 protein | | |
| XP_034236643.1 | GH9 | | Glycoside Hydrolase Family 9 protein | | |
| XP_034236904.1 | GT1 | | Glycosyltransferase Family 1 protein | | |
| XP_034237109.1 | GT61 | | Glycosyltransferase Family 61 protein | | |
| XP_034237121.1 | GH1 | | Glycoside Hydrolase Family 1 protein | | |
| XP_034237190.1 | GT57 | | Glycosyltransferase Family 57 protein | | |
| XP_034237191.1 | PL1_4 | (fragment) splicing problem; | Polysaccharide Lyase Family 1 protein | HGT | Complex: Bacteria or Fungi |
| XP_034237329.1 | AA3_2 | fragment C-term; | GMC oxidoreductase | | |
| XP_034237346.1 | AA3_2 | | GMC oxidoreductase | | |
| XP_034237429.1 | AA15 | | Auxilliary Activities Family 15 protein | | |
| XP_034237478.1 | CBM14 | | Carbohydrate-Binding Module Family 14 protein | | |
| XP_034237553.1 | GH20 | | Glycoside Hydrolase Family 20 protein | | |
| XP_034237725.1 | GH1 | | Glycoside Hydrolase Family 1 protein | | |
| XP_034237726.1 | GH1 | fragment N-term; | Glycoside Hydrolase Family 1 protein | | |
| XP_034237728.1 | GH1 | fragment C-term; | Glycoside Hydrolase Family 1 protein | | |
| XP_034237737.1 | CBM20 | | Carbohydrate-Binding Module Family 20 protein | | |
| XP_034237899.1 | GH1 | | Glycoside Hydrolase Family 1 protein | | |
| XP_034238314.1 | GH13_17 | | Glycoside Hydrolase Family 13 protein | | |
| XP_034238335.1 | GT29 | | Glycosyltransferase Family 29 protein | | |
| XP_034238381.1 | GT31 | | Glycosyltransferase Family 31 protein | | |
| XP_034238486.1 | GT66 | | Glycosyltransferase Family 66 protein | | |
| XP_034238644.1 | GH1 | | Glycoside Hydrolase Family 1 protein | | |
| XP_034238785.1 | GT24 | | Glycosyltransferase Family 24 protein | | |
| XP_034238855.1 | AA3_2 | | GMC oxidoreductase | | |
| XP_034238883.1 | GT22 | | Glycosyltransferase Family 22 protein | | |
| XP_034238963.1 | GH18 | | Glycoside Hydrolase Family 18 protein | | |
| XP_034238972.1 | GT10 | | Glycosyltransferase Family 10 protein | | |
| XP_034239010.1 | GH38 | | Glycoside Hydrolase Family 38 protein | | |
| XP_034239232.1 | GT22 | | Glycosyltransferase Family 22 protein | | |
| XP_034239330.1 | GT23 | | Glycosyltransferase Family 23 protein | | |
| XP_034239395.1 | GH30_1 | fragment N-term; | Glycoside Hydrolase Family 30 protein | | |
| XP_034239440.1 | PL1_4 | fragment N-term / C-term; | Polysaccharide Lyase Family 1 protein | HGT | Complex: Bacteria or Fungi |
| XP_034239465.1 | CBM39-GH16_4 | | Carbohydrate-Binding Module Family 39 / Glycoside Hydrolase Family 16 protein | | |
| XP_034239544.1 | CBM39-GH16_4 | (fragment) splicing problem; | Carbohydrate-Binding Module Family 39 / Glycoside Hydrolase Family 16 protein | | |
| XP_034239569.1 | GH18-CBM14 | | Glycoside Hydrolase Family 18 / Carbohydrate-Binding Module Family 14 protein | | |
| XP_034239606.1 | GT16 | | Glycosyltransferase Family 16 protein | | |
| XP_034239611.1 | GH22 | | Glycoside Hydrolase Family 22 protein | | |
| XP_034239672.1 | GT21 | (fragment) splicing problem; | Glycosyltransferase Family 21 protein | | |

| | | | | | |
|---|---|---|---|---|---|
| XP_034239705.1 | GH27 | | Glycoside Hydrolase Family 27 protein | | |
| XP_034239717.1 | CE9 | (fragment) splicing problem; | Carbohydrate Esterase Family 9 protein | | |
| XP_034239735.1 | GT1 | | Glycosyltransferase Family 1 protein | | |
| XP_034239858.1 | GH2 | | Glycoside Hydrolase Family 2 protein | | |
| XP_034239875.1 | GT57 | | Glycosyltransferase Family 57 protein | | |
| XP_034239912.1 | GH31 | | Glycoside Hydrolase Family 31 protein | | |
| XP_034240246.1 | AA1 | | Multicopper oxidase | | |
| XP_034240255.1 | CBM14 | | Carbohydrate-Binding Module Family 14 protein | | |
| XP_034240262.1 | AA3_2 | (fragment) splicing problem; | GMC oxidoreductase | | |
| XP_034240278.1 | AA3_2 | (fragment) splicing problem; | GMC oxidoreductase | | |
| XP_034240369.1 | GH30_1 | | Glycoside Hydrolase Family 30 protein | | |
| XP_034240415.1 | GH16_4 | | Glycoside Hydrolase Family 16 protein | | |
| XP_034240427.1 | CBM14 | | Carbohydrate-Binding Module Family 14 protein | | |
| XP_034240608.1 | GH1 | | Glycoside Hydrolase Family 1 protein | | |
| XP_034240623.1 | GH1 | | Glycoside Hydrolase Family 1 protein | | |
| XP_034240627.1 | GT22 | | Glycosyltransferase Family 22 protein | | |
| XP_034240632.1 | GT8 | | Glycosyltransferase Family 8 protein | | |
| XP_034240678.1 | GH1 | fragment C-term; | Glycoside Hydrolase Family 1 protein | | |
| XP_034240731.1 | GH1 | fragment N-term; | Glycoside Hydrolase Family 1 protein | | |
| XP_034240850.1 | GH1 | | Glycoside Hydrolase Family 1 protein | | |
| XP_034240865.1 | GT1 | | Glycosyltransferase Family 1 protein | | |
| XP_034240903.1 | GH16_4 | | Glycoside Hydrolase Family 16 protein | | |
| XP_034240904.1 | GH16_4 | | Glycoside Hydrolase Family 16 protein | | |
| XP_034241038.1 | PL1_4 | (fragment) splicing problem; | Polysaccharide Lyase Family 1 protein | HGT | Complex: Bacteria or Fungi |
| XP_034241133.1 | GT105 | (fragment) splicing problem; | Glycosyltransferase Family 105 protein | | |
| XP_034241502.1 | GT1 | | Glycosyltransferase Family 1 protein | | |
| XP_034241515.1 | GT31 | | Glycosyltransferase Family 31 protein | | |
| XP_034241559.1 | GT20 | | Glycosyltransferase Family 20 protein | | |
| XP_034241740.1 | GH13_25-GH133 | | Glycoside Hydrolase Family 13 / Glycoside Hydrolase Family 133 protein | | |
| XP_034241770.1 | CBM14 | | Carbohydrate-Binding Module Family 14 protein | | |
| XP_034241806.1 | GT31 | | Glycosyltransferase Family 31 protein | | |
| XP_034241934.1 | CBM14 | | Carbohydrate-Binding Module Family 14 protein | | |
| XP_034242052.1 | GH5_8 | | Glycoside Hydrolase Family 5 protein | HGT | Bacteria |
| XP_034242065.1 | GH1 | | Glycoside Hydrolase Family 1 protein | | |
| XP_034242067.1 | GH1 | fragment N-term; | Glycoside Hydrolase Family 1 protein | | |
| XP_034242068.1 | GH1 | | Glycoside Hydrolase Family 1 protein | | |
| XP_034242079.1 | CBM14-CBM14-GH1 | | Carbohydrate-Binding Module Family 14 / Glycoside Hydrolase Family 1 protein | | |
| XP_034242082.1 | GT1 | | Glycosyltransferase Family 1 protein | | |
| XP_034242145.1 | GH30_1 | | Glycoside Hydrolase Family 30 protein | | |
| XP_034242173.1 | GT1-GT1 | | Glycosyltransferase Family 1 protein | | |
| XP_034242309.1 | GH1 | | Glycoside Hydrolase Family 1 protein | | |
| XP_034242338.1 | GT54 | | Glycosyltransferase Family 54 protein | | |
| XP_034242400.1 | CBM14-CBM14-CBM14-CBM14-CBM14-CBM14-CBM14 | | Carbohydrate-Binding Module Family 14 protein | | |
| XP_034242473.1 | GT8 | | Glycosyltransferase Family 8 protein | | |
| XP_034242578.1 | GH5_8 | | Glycoside Hydrolase Family 5 protein | HGT | Bacteria |
| XP_034242622.1 | GH47 | | Glycoside Hydrolase Family 47 protein | | |
| XP_034242715.1 | AA3_2 | | GMC oxidoreductase | | |
| XP_034242717.1 | AA3_2 | | GMC oxidoreductase | | |
| XP_034242730.1 | GH1 | | Glycoside Hydrolase Family 1 protein | | |

| | | | | | |
|---|---|---|---|---|---|
| XP_034242821.1 | GT43 | | Glycosyltransferase Family 43 protein | | |
| XP_034242876.1 | GT2 | | Glycosyltransferase Family 2 protein | | |
| XP_034242958.1 | GH18-CBM14-CBM14-GH18-CBM14-CBM14-CBM14-GH18-GH18-CBM14-GH18 | | Glycoside Hydrolase Family 18 / Carbohydrate-Binding Module Family 14 protein | | |
| XP_034243006.1 | GT1 | | Glycosyltransferase Family 1 protein | | |
| XP_034243155.1 | AA3_2 | | GMC oxidoreductase | | |
| XP_034243232.1 | GH20 | | Glycoside Hydrolase Family 20 protein | | |
| XP_034243548.1 | AA3_2 | fragment C-term; | GMC oxidoreductase | | |
| XP_034243618.1 | GT76 | | Glycosyltransferase Family 76 protein | | |
| XP_034243778.1 | GT31 | | Glycosyltransferase Family 31 protein | | |
| XP_034243895.1 | GH2 | | Glycoside Hydrolase Family 2 protein | | |
| XP_034243932.1 | GT1 | | Glycosyltransferase Family 1 protein | | |
| XP_034244054.1 | GT39 | | Glycosyltransferase Family 39 protein | | |
| XP_034244075.1 | GH63 | | Glycoside Hydrolase Family 63 protein | | |
| XP_034244218.1 | GH47 | | Glycoside Hydrolase Family 47 protein | | |
| XP_034244463.1 | GT27-CBM13 | | Glycosyltransferase Family 27 / Carbohydrate-Binding Module Family 13 protein | | |
| XP_034244583.1 | AA3_2 | | GMC oxidoreductase | | |
| XP_034244609.1 | GT27-CBM13 | | Glycosyltransferase Family 27 / Carbohydrate-Binding Module Family 13 protein | | |
| XP_034244814.1 | GH31 | | Glycoside Hydrolase Family 31 protein | | |
| XP_034244844.1 | GT50 | | Glycosyltransferase Family 50 protein | | |
| XP_034244861.1 | CBM21 | | Carbohydrate-Binding Module Family 21 protein | | |
| XP_034244864.1 | CBM14-CBM14 | | Carbohydrate-Binding Module Family 14 protein | | |
| XP_034244872.1 | CBM50 | | Carbohydrate-Binding Module Family 50 protein | | |
| XP_034244875.1 | GH22 | | Glycoside Hydrolase Family 22 protein | | |
| XP_034244880.1 | GH22 | | Glycoside Hydrolase Family 22 protein | | |
| XP_034244905.1 | GH1 | | Glycoside Hydrolase Family 1 protein | | |
| XP_034244999.1 | GH1 | | Glycoside Hydrolase Family 1 protein | | |
| XP_034245055.1 | AA3_2 | | GMC oxidoreductase | | |
| XP_034245074.1 | AA3_2 | | GMC oxidoreductase | | |
| XP_034245312.1 | AA1 | | Multicopper oxidase | | |
| XP_034245357.1 | GH1 | | Glycoside Hydrolase Family 1 protein | | |
| XP_034245606.1 | GH30_1 | | Glycoside Hydrolase Family 30 protein | | |
| XP_034245684.1 | GH1 | fragment N-term / C-term; | Glycoside Hydrolase Family 1 protein | | |
| XP_034245696.1 | GH1 | | Glycoside Hydrolase Family 1 protein | | |
| XP_034245819.1 | CBM43-CBM43-CBM43-CBM43 | | Carbohydrate-Binding Module Family 43 protein | HGT | Viridiplantae |
| XP_034246068.1 | GT27-CBM13 | | Glycosyltransferase Family 27 / Carbohydrate-Binding Module Family 13 protein | | |
| XP_034246129.1 | CBM14-CBM14-GH1 | | Carbohydrate-Binding Module Family 14 / Glycoside Hydrolase Family 1 protein | | |
| XP_034246266.1 | GT49 | | Glycosyltransferase Family 49 protein | | |
| XP_034246495.1 | GH18 | | Glycoside Hydrolase Family 18 protein | | |
| XP_034246754.1 | GT2 | | Glycosyltransferase Family 2 protein | | |
| XP_034246764.1 | CBM14-CBM14-GH1 | (fragment) splicing problem; | Carbohydrate-Binding Module Family 14 / Glycoside Hydrolase Family 1 protein | | |
| XP_034246835.1 | GT105 | fragment N-term; | Glycosyltransferase Family 105 protein | | |
| XP_034246846.1 | GH47 | | Glycoside Hydrolase Family 47 protein | | |
| XP_034246957.1 | GH116 | | Glycoside Hydrolase Family 116 protein | | |
| XP_034247376.1 | GH152 | | Glycoside Hydrolase Family 152 protein | HGT | Viridiplantae |
| XP_034247478.1 | GH38 | fragment C-term; | Glycoside Hydrolase Family 38 protein | | |
| XP_034247610.1 | GT105 | | Glycosyltransferase Family 105 protein | | |
| XP_034247673.1 | GH152 | | Glycoside Hydrolase Family 152 protein | HGT | Viridiplantae |
| XP_034247731.1 | GT31 | | Glycosyltransferase Family 31 protein | | |
| XP_034247819.1 | AA3_2 | | GMC oxidoreductase | | |

| | | | | | |
|---|---|---|---|---|---|
| XP_034247857.1 | AA3_2 | | GMC oxidoreductase | | |
| XP_034247887.1 | GT27-CBM13 | | Glycosyltransferase Family 27 / Carbohydrate-Binding Module Family 13 protein | | |
| XP_034247964.1 | GH1 | | Glycoside Hydrolase Family 1 protein | | |
| XP_034248138.1 | GH85 | (fragment) splicing problem; | Glycoside Hydrolase Family 85 protein | | |
| XP_034248270.1 | GH18-GH18-CBM14 | | Glycoside Hydrolase Family 18 / Carbohydrate-Binding Module Family 14 protein | | |
| XP_034248354.1 | PL1_4 | fragment C-term; | Polysaccharide Lyase Family 1 protein | HGT | Complex: Bacteria or Fungi |
| XP_034248372.1 | GH18-CBM14-CBM14-CBM14 | (fragment) splicing problem; | Glycoside Hydrolase Family 18 / Carbohydrate-Binding Module Family 14 protein | | |
| XP_034248377.1 | GT92 | | Glycosyltransferase Family 92 protein | | |
| XP_034248438.1 | CBM14 | | Carbohydrate-Binding Module Family 14 protein | | |
| XP_034248531.1 | AA15 | | Auxilliary Activities Family 15 protein | | |
| XP_034248552.1 | PL1_4 | fragment C-term; | Polysaccharide Lyase Family 1 protein | HGT | Complex: Bacteria or Fungi |
| XP_034248577.1 | GT3 | | Glycosyltransferase Family 3 protein | | |
| XP_034248646.1 | GH1 | fragment C-term; | Glycoside Hydrolase Family 1 protein | | |
| XP_034248773.1 | CBM14-CBM14-GH1 | | Carbohydrate-Binding Module Family 14 / Glycoside Hydrolase Family 1 protein | | |
| XP_034248903.1 | GT10 | (fragment) splicing problem; | Glycosyltransferase Family 10 protein | | |
| XP_034248925.1 | GH37 | | Glycoside Hydrolase Family 37 protein | | |
| XP_034248976.1 | GH1 | | Glycoside Hydrolase Family 1 protein | | |
| XP_034248983.1 | CBM14 | | Carbohydrate-Binding Module Family 14 protein | | |
| XP_034249033.1 | GT4 | | Glycosyltransferase Family 4 protein | | |
| XP_034249124.1 | GH37 | | Glycoside Hydrolase Family 37 protein | | |
| XP_034249179.1 | GH1 | | Glycoside Hydrolase Family 1 protein | | |
| XP_034249322.1 | GH1 | | Glycoside Hydrolase Family 1 protein | | |
| XP_034249356.1 | GH1 | | Glycoside Hydrolase Family 1 protein | | |
| XP_034249360.1 | GT39 | | Glycosyltransferase Family 39 protein | | |
| XP_034249440.1 | GH38 | | Glycoside Hydrolase Family 38 protein | | |
| XP_034249611.1 | CBM13 | (fragment) splicing problem; | Carbohydrate-Binding Module Family 13 protein | | |
| XP_034249645.1 | GT31-GT7 | | Glycosyltransferase Family 31 / Glycosyltransferase Family 7 protein | | |
| XP_034249763.1 | GT49 | | Glycosyltransferase Family 49 protein | | |
| XP_034249890.1 | GH1 | (fragment) splicing problem; | Glycoside Hydrolase Family 1 protein | | |
| XP_034249940.1 | GH13 | | Glycoside Hydrolase Family 13 protein | | |
| XP_034249956.1 | GT8-GT49 | (fragment) splicing problem; | Glycosyltransferase Family 8 / Glycosyltransferase Family 49 protein | | |
| XP_034250000.1 | GT14 | | Glycosyltransferase Family 14 protein | | |
| XP_034250014.1 | GH31 | | Glycoside Hydrolase Family 31 protein | | |
| XP_034250019.1 | GH18-CBM14 | | Glycoside Hydrolase Family 18 / Carbohydrate-Binding Module Family 14 protein | | |
| XP_034250029.1 | GT47-GT64 | | Glycosyltransferase Family 47 / Glycosyltransferase Family 64 protein | | |
| XP_034250033.1 | GH1 | (fragment) splicing problem; | Glycoside Hydrolase Family 1 protein | | |
| XP_034250131.1 | GT1 | | Glycosyltransferase Family 1 protein | | |
| XP_034250265.1 | GT1 | fragment N-term; | Glycosyltransferase Family 1 protein | | |
| XP_034250391.1 | AA3_2 | | GMC oxidoreductase | | |
| XP_034250679.1 | GT66 | | Glycosyltransferase Family 66 protein | | |
| XP_034250782.1 | GH13 | | Glycoside Hydrolase Family 13 protein | | |
| XP_034250870.1 | GH29 | | Glycoside Hydrolase Family 29 protein | | |
| XP_034250949.1 | GH2 | | Glycoside Hydrolase Family 2 protein | | |
| XP_034251003.1 | CBM21 | | Carbohydrate-Binding Module Family 21 protein | | |
| XP_034251442.1 | CBM48-GH13_8 | | Carbohydrate-Binding Module Family 48 / Glycoside Hydrolase Family 13 protein | | |
| XP_034251591.1 | GT59 | | Glycosyltransferase Family 59 protein | | |
| XP_034251595.1 | GT31 | | Glycosyltransferase Family 31 protein | | |
| XP_034251951.1 | GT7 | | Glycosyltransferase Family 7 protein | | |
| XP_034252032.1 | GH1 | | Glycoside Hydrolase Family 1 protein | | |

| | | | | | |
|---|---|---|---|---|---|
| XP_034252322.1 | GH1 | | Glycoside Hydrolase Family 1 protein | | |
| XP_034252555.1 | GH1 | fragment N-term; | Glycoside Hydrolase Family 1 protein | | |
| XP_034252647.1 | GT31 | | Glycosyltransferase Family 31 protein | | |
| XP_034252780.1 | GT47-GT64 | | Glycosyltransferase Family 47 / Glycosyltransferase Family 64 protein | | |
| XP_034252867.1 | CBM14 | | Carbohydrate-Binding Module Family 14 protein | | |
| XP_034252888.1 | GT22 | | Glycosyltransferase Family 22 protein | | |
| XP_034252975.1 | CBM14 | | Carbohydrate-Binding Module Family 14 protein | | |
| XP_034252989.1 | GH18 | | Glycoside Hydrolase Family 18 protein | | |
| XP_034253001.1 | GH99 | | Glycoside Hydrolase Family 99 protein | | |
| XP_034253060.1 | CBM14 | | Carbohydrate-Binding Module Family 14 protein | | |
| XP_034253076.1 | GH1 | fragment N-term; | Glycoside Hydrolase Family 1 protein | | |
| XP_034253121.1 | GT27-CBM13 | | Glycosyltransferase Family 27 / Carbohydrate-Binding Module Family 13 protein | | |
| XP_034253302.1 | GT47-GT64 | | Glycosyltransferase Family 47 / Glycosyltransferase Family 64 protein | | |
| XP_034253455.1 | GT1 | | Glycosyltransferase Family 1 protein | | |
| XP_034253511.1 | CBM14 | | Carbohydrate-Binding Module Family 14 protein | | |
| XP_034253547.1 | CBM14 | | Carbohydrate-Binding Module Family 14 protein | | |
| XP_034253708.1 | GT4 | | Glycosyltransferase Family 4 protein | | |
| XP_034253709.1 | CBM14 | (fragment) splicing problem; | Carbohydrate-Binding Module Family 14 protein | | |
| XP_034253736.1 | CBM14 | | Carbohydrate-Binding Module Family 14 protein | | |
| XP_034253757.1 | CBM14 | | Carbohydrate-Binding Module Family 14 protein | | |
| XP_034253769.1 | PL1_4 | fragment C-term; | Polysaccharide Lyase Family 1 protein | HGT | Complex: Bacteria or Fungi |
| XP_034253771.1 | PL1_4 | fragment C-term; | Polysaccharide Lyase Family 1 protein | HGT | Complex: Bacteria or Fungi |
| XP_034254025.1 | GT4 | | Glycosyltransferase Family 4 protein | | |
| XP_034254051.1 | GH47 | | Glycoside Hydrolase Family 47 protein | | |
| XP_034254370.1 | GT27-CBM13 | | Glycosyltransferase Family 27 / Carbohydrate-Binding Module Family 13 protein | | |
| XP_034254403.1 | GT27-CBM13 | | Glycosyltransferase Family 27 / Carbohydrate-Binding Module Family 13 protein | | |
| XP_034254538.1 | GH31 | | Glycoside Hydrolase Family 31 protein | | |
| XP_034254592.1 | AA3_2 | | GMC oxidoreductase | | |
| XP_034254627.1 | GH84 | | Glycoside Hydrolase Family 84 protein | | |
| XP_034254640.1 | GH79 | | Glycoside Hydrolase Family 79 protein | | |
| XP_034254683.1 | GH89 | fragment N-term; | Glycoside Hydrolase Family 89 protein | | |
| XP_034254706.1 | GH79 | | Glycoside Hydrolase Family 79 protein | | |
| XP_034254735.1 | GT7 | | Glycosyltransferase Family 7 protein | | |
| XP_034254881.1 | GT68 | | Glycosyltransferase Family 68 protein | | |
| XP_034254926.1 | GH47 | | Glycoside Hydrolase Family 47 protein | | |
| XP_034254936.1 | CBM14-CBM14-GH1 | | Carbohydrate-Binding Module Family 14 / Glycoside Hydrolase Family 1 protein | | |
| XP_034254938.1 | PL1_4 | fragment C-term; | Polysaccharide Lyase Family 1 protein | HGT | Complex: Bacteria or Fungi |
| XP_034255023.1 | AA3_2 | | GMC oxidoreductase | | |
| XP_034255257.1 | GT7 | | Glycosyltransferase Family 7 protein | | |
| XP_034255289.1 | GT25 | | Glycosyltransferase Family 25 protein | | |
| XP_034255299.1 | AA3_2 | | GMC oxidoreductase | | |
| XP_034255300.1 | GH22 | | Glycoside Hydrolase Family 22 protein | | |
| XP_034255301.1 | GH22 | | Glycoside Hydrolase Family 22 protein | | |
| XP_034255332.1 | GH1 | | Glycoside Hydrolase Family 1 protein | | |
| XP_034255409.1 | GT54 | | Glycosyltransferase Family 54 protein | | |
| XP_034255447.1 | GT1 | | Glycosyltransferase Family 1 protein | | |
| XP_034255541.1 | GH22 | | Glycoside Hydrolase Family 22 protein | | |
| XP_034255542.1 | GH22 | | Glycoside Hydrolase Family 22 protein | | |
| XP_034255655.1 | AA3_2 | | GMC oxidoreductase | | |

| | | | |
|---|---|---|---|
| XP_034255658.1 | AA3_2 | | GMC oxidoreductase |
| XP_034255772.1 | GT58 | | Glycosyltransferase Family 58 protein |
| XP_034255852.1 | GT35 | | Glycosyltransferase Family 35 protein |
| XP_034255893.1 | GH20 | | Glycoside Hydrolase Family 20 protein |
| XP_034255982.1 | GT33 | | Glycosyltransferase Family 33 protein |
| XP_034256147.1 | AA1 | | Multicopper oxidase |
| XP_034256234.1 | CBM48 | | Carbohydrate-Binding Module Family 48 protein |
| XP_034256246.1 | AA3_2 | | GMC oxidoreductase |
| XP_034256272.1 | GT31-GT7 | | Glycosyltransferase Family 31 / Glycosyltransferase Family 7 protein |
| XP_034256300.1 | GT31 | (fragment) splicing problem; | Glycosyltransferase Family 31 protein |
| XP_034256316.1 | GT98 | | Glycosyltransferase Family 98 protein |
| XP_034256384.1 | CBM14 | | Carbohydrate-Binding Module Family 14 protein |
| XP_034256564.1 | AA3_2 | | GMC oxidoreductase |
| XP_034256691.1 | GH1 | | Glycoside Hydrolase Family 1 protein |
| XP_034256971.1 | CBM14 | | Carbohydrate-Binding Module Family 14 protein |
| XP_034256998.1 | GT105 | | Glycosyltransferase Family 105 protein |
| XP_034257021.1 | CBM14 | | Carbohydrate-Binding Module Family 14 protein |
| XP_034257023.1 | AA1 | | Multicopper oxidase |
| XP_034257033.1 | AA3_2 | | GMC oxidoreductase |

**Supplementary Table 6:** Validated HGT candidates from potential bacterial, fungal or viridiplantae donors for *T. vaporariorum*

*Bacterial donor:*

| HGT event | Sequence name | Origin of donor sequences | Alternative topology Monophyly of Metazoa | Similarities with the donor sequences | | Length and coverage of the alignment with the donor sequences | | Local score for genomic environment | Homologs found in the *B. tabaci* cryptic species | | | Annotation (BLAST NR) | CAZy |
|---|---|---|---|---|---|---|---|---|---|---|---|---|---|
| | | | | max id | average id | average aln length | average coverage | | Btab_MEAM1 | Btab_MED | Btab_SSA | | |
| TvaB01 | TLow_01347-RA | Bacteria | / | 34.2 | 28.2 | 430 | 81.7 | 0.92 | Bta10852, | BTA009628.1, | Ssa11036 | AAA family ATPase | |
| | TLow_02144-RA | | | 30.8 | 28.6 | 446 | 76.7 | 0.86 | Bta10853 | BTA009629.1, | | | |
| | TLow_02157-RA | | | 31.4 | 27.5 | 353 | 71.7 | 0.88 | | BTA025251.1, | | | |
| | Tv_13818-RA | | | 30.2 | 27.9 | 449 | 76.6 | 0.85 | | BTA025252.1, | | | |
| | Tv_13862-RA | | | 45.3 | 34.6 | 110 | 47.5 | 1.0 | | BTA025253.1 | | | |
| | Tv_15776-RA | | | 31.2 | 26.0 | 438 | 67.1 | 0.83 | | | | | |
| TvaB02 | Tv_01895-RA | Bacteria | 1 | 77.7 | 60.6 | 419 | 49.2 | 0.64 | Bta00841, Bta01937 | BTA023651.1 | Ssa01553 | adenosylmethionine--8-amino-7-oxononanoate transaminase | |
| TvaB03 | Tv_02831-RA | Bacteria | / | 53.9 | 50.5 | 151 | 95.6 | 0.87 | Bta00747, | | Ssa08703, | dUTP diphosphatase | |
| | Tv_11854-RA | | | 53.9 | 50.5 | 151 | 95.8 | 0.71 | Bta13975 | | Ssa14910 | | |
| | Tv_12101-RA | | | 49.0 | 43.6 | 121 | 30.2 | 0.85 | | | | | |
| | Tv_15086-RA | | | 60.8 | 55.7 | 78 | 89.0 | 0.84 | | | | | |
| | Tv_15720-RA | | | 48.1 | 44.1 | 153 | 33.7 | 0.85 | | | | | |
| | Tv_15881-RA | | | 52.0 | 43.8 | 125 | 70.0 | 0.76 | | | | | |
| TvaB04 | Tv_15063-RA | Bacteria | / | 46.3 | 38.6 | 135 | 64.5 | 0.86 | | | | AAA family ATPase | |
| | Tv_15070-RA | | | 34.0 | 32.9 | 405 | 89.1 | 0.86 | | | | | |
| TvaB05 | Tv_01894-RA | Bacteria | 1 | 76.2 | 55.8 | 202 | 93.9 | 0.74 | Bta00840, Bta01938 | | Ssa01088 | dethiobiotin synthase | |
| TvaB06 | Tv_05628-RA | Bacteria | 0 | 40.1 | 32.3 | 433 | 86.0 | 0.79 | Bta12386 | BTA006502.1 | Ssa03543, Ssa03544 | | |
| TvaB07 | Tv_00146-RA | Bacteria | / | 32 | 30.1 | 512 | 72.0 | 0.9 | Bta00829 | BTA007316.1, | scaffold554257_3 | AAA family ATPase | |
| | Tv_00148-RA | | | 30.8 | 28.6 | 546 | 74.5 | 0.9 | | BTA011402.1 | 12_1956 | | |
| | Tv_02102-RA | | | 32 | 28.9 | 498 | 67.1 | 0.62 | | | | | |
| | Tv_02106-RA | | | 28.4 | 25.5 | 213 | 72.9 | 0.62 | | | | | |
| | Tv_01481-RA | | | 30.7 | 28.8 | 550 | 71.2 | 0.96 | | | | | |
| | Tv_15366-RA | | | 36.7 | 31.2 | 399 | 80.2 | 0.95 | | | | | |
| TvaB08 | Tv_12316-RA | Bacteria | / | 69.4 | 67.8 | 216 | 99.5 | 0.88 | | | | nucleoside/nucleotide kinase family protein | |
| TvaB09 | TLow_01825-RA | Bacteria | 1 | 33.1 | 30.2 | 262 | 78.4 | 0.96 | | | | Rpn family recombination- | |
| | Tv_14938-RA | | | 33.7 | 28.2 | 165 | 20.8 | 0.88 | | | | promoting nuclease/putative | |
| TvaB10 | Tv_04079-RA | Bacteria | / | 65.4 | 60.0 | 85 | 72.0 | 0.92 | | | | | GH13 |
| TvaB11 | Tv_13090-RA | Bacteria | / | 56.8 | 47.6 | 259 | 95.9 | 1.0 | | | | PhzF family phenazine biosynthesis isomerase | |
| TvaB12 | TLow_01333-RA | Bacteria | 0 | 31.2 | 29.5 | 279 | 82.6 | 0.96 | | | | Rpn family recombination-promoting nuclease/putative transposase | |
| TvaB13 | Tv_05804-RA | Bacteria | / | 32.1 | 28.9 | 481 | 78.5 | 0.8 | | | | AAA family ATPase | |
| TvaB14 | Tv_01789-RA | Bacteria | / | 34.3 | 31.3 | 463 | 74.8 | 0.92 | | | | AAA family ATPase | |
| | Tv_08710-RA | | | 31.0 | 29.0 | 516 | 85.0 | 0.96 | | | | | |
| TvaB15 | Tv_04874-RA | Bacteria | / | 54.8 | 52.4 | 155 | 84.0 | 0.96 | Bta03797 | BTA000817.1 | Ssa13701 | methylated-DNA--[protein]-cysteine S-methyltransferase | |
| TvaB16 | Tv_06835-RA | Bacteria | / | 45.0 | 41.1 | 158 | 80.0 | 0.88 | Bta15103 | BTA020874.1 | Ssa07364 | chorismate mutase | |
| TvaB17 | Tv_00487-RA | Bacteria | / | 37.6 | 32.1 | 129 | 82.5 | 0.77 | Bta05013, | BTA002393.1 | Ssa01599, | AAA family ATPase | |
| | Tv_11451-RA | | | 31.6 | 29.4 | 554 | 73.0 | 1.0 | Bta10549, | | Ssa10865, | | |
| | | | | | | | | | Bta14973 | | Ssa10873, Ssa12918, Ssa13905, Ssa14120, Ssa15393 | | |

| | | | | | | | | | | | | |
|---|---|---|---|---|---|---|---|---|---|---|---|---|
| TvaB18 | Tv_09126-RA<br>Tv_13900-RA<br>Tv_14921-RA<br>Tv_15127-RA<br>Tv_15631-RA | Bacteria | / | 34.3<br>32.9<br>31.3<br>35.4<br>33.3 | 30.9<br>29.8<br>28.7<br>31.7<br>31.3 | 462<br>408<br>539<br>559<br>558 | 72.1<br>90.2<br>95.1<br>91.6<br>89.2 | 0.88<br>0.86<br>0.64<br>0.84<br>0.78 | | | | AAA family ATPase |
| TvaB19 | Tv_08602-RA<br>Tv_10007-RA | Bacteria | / | 68.2<br>68.4 | 66.9<br>67.0 | 371<br>380 | 65.4<br>86.1 | 0.8<br>0.9 | Bta02653,<br>Bta04469,<br>Bta05264,<br>Bta06818,<br>Bta11911,<br>Bta11912 | BTA005171.1,<br>BTA005173.1,<br>BTA012513.1,<br>BTA019053.1,<br>BTA023694.1,<br>BTA023695.1,<br>BTA029954.2 | Ssa00993,<br>Ssa02587,<br>Ssa02588,<br>Ssa02589,<br>Ssa09979,<br>Ssa12718 | cyclopropane fatty acyl phospholipid |
| TvaB20 | Tv_15468-RA | Bacteria | 1 | 69.8 | 64.1 | 308 | 90.8 | 0.92 | Bta09725 | BTA004219.1,<br>BTA028913.1 | Ssa10171 | Biotin synthase |
| TvaB21 | Tv_04577-RA<br>Tv_08891-RA<br>Tv_08906-RA<br>Tv_08957-RA<br>Tv_08958-RA | Bacteria | / | 36.6<br>35.1<br>35.9<br>36.1<br>34.4 | 31.2<br>30.3<br>31.0<br>30.7<br>31.4 | 416<br>441<br>389<br>428<br>393 | 66.9<br>70.4<br>83.0<br>73.8<br>84.6 | 0.91<br>0.87<br>0.83<br>0.86<br>0.86 | | | | AAA family ATPase |
| TvaB22 | Tv_02105-RA | Bacteria | / | 39.2 | 35.6 | 251 | 95.3 | 0.62 | Bta11531,<br>Bta15479 | BTA004921.1,<br>BTA018889.1,<br>BTA026605.1 | Ssa12442 | AAA family ATPase |
| TvaB23 | TLow_02263-RA<br>Tv_09587-RA<br>Tv_13579-RA<br>Tv_14715-RA<br>Tv_14844-RA | Bacteria | / | 35.4<br>35.8<br>34.2<br>33.4<br>34.9 | 33.2<br>30.5<br>31.5<br>31.1<br>32.1 | 287<br>463<br>482<br>302<br>409 | 94.6<br>71.9<br>81.4<br>73.6<br>94.1 | 0.81<br>0.96<br>0.88<br>0.92<br>0.78 | | | | AAA family ATPase |
| TvaB24 | TLow_01084-RA<br>Tv_01996-RA<br>Tv_11525-RA<br>Tv_15653-RA | Bacteria | / | 60.3<br>55.9<br>57.1<br>57.1 | 54.3<br>46.9<br>50.7<br>50.8 | 66<br>79<br>143<br>143 | 82.8<br>52.2<br>94.8<br>94.7 | 0.96<br>0.76<br>0.76<br>0.91 | | | | dUTP diphosphatase |

*Fungal donor:*

| HGT event | Sequence name | Origin of donor sequences | Alternative topology | Similarities with the donor sequences | | Length and coverage of the alignment with the donor sequences | | Local score for genomic environment | Homologs found in the *B. tabaci* cryptic species | | | Annotation (BLAST NR) | CAZy |
|---|---|---|---|---|---|---|---|---|---|---|---|---|---|
| | | | | max id | average id | average aln length | average coverage | | Btab_MEAM1 | Btab_MED | Btab_SSA | | |
| TvaF01 | Tv_08361-RA<br>Tv_14110-RA<br>Tv_14111-RA | Fungi | / | 32,0<br>34,0<br>34,3 | 30,2<br>31,3<br>31,6 | 910<br>681<br>1274 | 95,3<br>95,0<br>99,0 | 0.7<br>0.68<br>0.68 | Bta00932,<br>Bta05475,<br>Bta07433 | BTA005597.1,<br>BTA005598.1,<br>BTA005599.1,<br>BTA007606.1,<br>BTA003000.1,<br>BTA004644.1,<br>BTA008032.1,<br>BTA008563.1 | sa06477,<br>Ssa06478,<br>Ssa06479,<br>Ssa04914,<br>Ssa15384 | MYND finger family protein | |
| TvaF02 | Tv_08362-RA | Fungi | / | 35,7 | 32,8 | 331 | 96,5 | 0.66 | Bta07948 | BTA017217.1,<br>BTA017221.1,<br>BTA021006.1 | Ssa00835 | MYND finger family protein | |
| TvaF03 | Tv_07966-RA | Fungi | / | 49,9 | 45,9 | 2392 | 97,7 | 0.78 | Bta08944,<br>Bta08945,<br>Bta08946,<br>Bta14076 | BTA016279.2,<br>BTA025951.2,<br>BTA025953.1,<br>BTA025954.1 | Ssa11553,<br>Ssa12591,<br>Ssa15247 | NFX1-type zinc finger-containing protein 1 | |
| TvaF04 | Tv_10871-RA<br>Tv_10872-RA<br>Tv_11256-RA | Fungi | / | 54,8<br>55,7<br>51,6 | 36,9<br>37,1<br>36,8 | 216<br>213<br>212 | 73,9<br>73,1<br>68,7 | 0.64<br>0.64<br>0.91 | Bta03426 | BTA002830.1,<br>BTA019829.1,<br>BTA029129.1,<br>BTA002831.1 | Ssa05110,<br>scaffold50840_2_<br>365 | Unknown protein | |

*Viridiplantae donor:*

| HGT event | Sequence name | Origin of donor sequences | Alternative topology | Similarities with the donor sequences | | Length and coverage of the alignment with the donor sequences | | Local score for genomic environment | Homologs found in the *B. tabaci* cryptic species | | | Annotation (BLAST NR) | CAZy |
|---|---|---|---|---|---|---|---|---|---|---|---|---|---|
| | | | | max id | average id | average aln length | average coverage | | Btab_MEAM1 | Btab_MED | Btab_SSA | | |
| TvaV01 | Tv_15928-RA | Viridiplantae | 0 | 60,2 | 57,7 | 202 | 91,7 | 1.0 | Bta13961 | BTA005662.2 | Ssa10394 | Thaumatin-like protein | GH152 |
| TvaV02 | TLow_01580-RA<br>Tv_15540-RA<br>Tv_15541-RA | Viridiplantae | / | 39,4<br>38,1<br>31,5 | 34,6<br>34,1<br>27,7 | 220<br>223<br>239 | 83,5<br>74,2<br>82,5 | 0.61<br>0.71<br>0.61 | Bta13094,<br>Bta13103 | BTA025985.1,<br>BTA025994.1 | Ssa09189,<br>scaffold212068_54_492 | rRNA N-glycosidase | |
| TvaV03 | Tv_08972-RA | Viridiplantae | / | 71,0 | 68,4 | 379 | 96,9 | 0.96 | Bta04264,<br>Bta06451,<br>Bta06452,<br>Bta06453,<br>Bta09295,<br>Bta13659,<br>Bta15754,<br>Bta15755,<br>Bta15756,<br>Bta15757,<br>Bta15759,<br>Bta15760,<br>Bta15762,<br>Bta15765 | BTA007356.1,<br>BTA011448.1,<br>BTA015354.2,<br>BTA015355.1,<br>BTA028840.1,<br>BTA015596.1,<br>BTA011446.1 | Ssa02331,<br>Ssa02332,<br>Ssa02334,<br>Ssa02342,<br>Ssa03431,<br>Ssa08612,<br>Ssa11969,<br>Ssa15146,<br>Ssa15387,<br>Ssa06732,<br>Ssa02337,<br>Ssa02338,<br>Ssa03432 | delta(12)-fatty-acid desaturase FAD2 | |

*Complex: Bacterial or Fungal donor*

| HGT event | Sequence name | Origin of donor sequences | Alternative topology | Similarities with the donor sequences | | Length and coverage of the alignment with the donor sequences | | Local score for genomic environment | Homologs found in the *B. tabaci* cryptic species | | | Annotation (BLAST NR) | CAZy |
|---|---|---|---|---|---|---|---|---|---|---|---|---|---|
| | | | | max id | average id | average aln length | average coverage | | Btab_MEAM1 | Btab_MED | Btab_SSA | | |
| TvaC01 | Tv_05105-RA | Bacteria or Fungi | 0 | 34.2 | 30.5 | 351 | 80.6 | 0.76 | Bta02634 | BTA005662.2 | Ssa10394 | Pectin lyase | PL1_4 |

**Supplementary Table 7:** Analysis of the 11 combined groups for *B. tabaci* and *T. vaporariorum* in which the grouping of sequences lacked bootstrap support and/or was not monphyletic

*Results of bootstrap analysis for the combined groups for which the grouping of sequences lacked bootstrap support:*

| Combined group | Ultrafast bootstrap support value |
|---|---|
| TvaB05_BtaB02 | 70% |
| TvaV02_BtaV23 | 91% |

Support values were based on an ultrafast bootstrap approximation with 1,000 replicates.
Only support values greater than or equal to 95% were considered.

*Results of the approximately unbiased (AU) alternative topology test for monophyly of Aleyrodinae sequences for the combined groups in which the grouping of sequences was not monophyletic:*

| Combined group | Tree | logL | deltaL | p-AU | |
|---|---|---|---|---|---|
| TvaB01_BtaB56 | 1 | -54869.4139 | 3.9658 | 0.434 | + |
| | 2 | -54865.44813 | 0 | 0.566 | + |
| TvaB02_BtaB12 | 1 | -15665.71625 | 2.8104 | 0.379 | + |
| | 2 | -15662.90584 | 0 | 0.621 | + |
| TvaB03_BtaB68 | 1 | -25152.92208 | 0 | 1 | + |
| | 2 | -25661.69097 | 508.77 | 6.13e-06 | - |
| TvaB04_BtaB44 | 1 | -15393.59309 | 18.658 | 0.129 | + |
| | 2 | -15374.93497 | 0 | 0.871 | + |
| TvaB08_BtaB31 | 1 | -4046.24064 | 18.874 | 0.12 | + |
| | 2 | -4027.366596 | 0 | 0.88 | + |
| TvaB11_BtaB03 | 1 | -18272.4574 | 103.52 | 8.67e-07 | - |
| | 2 | -18168.93243 | 0 | 1 | + |
| TvaF04_BtaF10 | 1 | -3488.191761 | 2.077 | 0.428 | + |
| | 2 | -3486.114778 | 0 | 0.572 | + |
| TvaV01_BtaV01 | 1 | -11952.22823 | 25.552 | 0.177 | + |
| | 2 | -11926.67646 | 0 | 0.823 | + |
| TvaV03_BtaV21 | 1 | -44046.69688 | 5.2108 | 0.46 | + |
| | 2 | -44041.48611 | 0 | 0.54 | + |

Plus signs denote the 95% confidence sets.
Minus signs denote significant exclusion.
All tests performed 10,000 resamplings using the RELL method.

**Supplementary Table 8:** Overrepresented GO terms among validated HGT candidates from potential bacterial, fungal or viridiplantae donors for *T. vaporariorum*

| root_node_name | node_id | node_name | raw_p_overrep | FWER_overrep | refined_p_overrep | FDR_overrep | nb_genes_refset_root_node | nb_genes_candidate_in_root_node | nb_gene_refset_node | nb_genes_candidate_in_node | name_genes_candidate_in_node |
|---|---|---|---|---|---|---|---|---|---|---|---|
| HGT Bacteria | | | | | | | | | | | |
| molecular_function | GO:0000287 | magnesium ion binding | 2.39324290613495e-22 | 0 | 2.39324290613495e-22 | 8.46182313240573e-22 | 6896 | 17 | 42 | 11 | TLow_01084-RA\|Tv_01894-RA\|Tv_02831-RA\|Tv_07802-RA\|Tv_09064-RA\|Tv_11525-RA\|Tv_11854-RA\|Tv_12101-RA\|Tv_15653-RA\|Tv_15720-RA\|Tv_15881-RA |
| molecular_function | GO:0004170 | dUTP diphosphatase activity | 6.51336021681486e-26 | 0 | 6.51336021681486e-26 | 6.44822661464671e-24 | 6896 | 17 | 14 | 10 | TLow_01084-RA\|Tv_02831-RA\|Tv_07802-RA\|Tv_09064-RA\|Tv_11525-RA\|Tv_11854-RA\|Tv_12101-RA\|Tv_15653-RA\|Tv_15720-RA\|Tv_15881-RA |
| biological_process | GO:0006226 | dUMP biosynthetic process | 8.68032002463571e-24 | 0 | 8.68032002463571e-24 | 7.16126402032446e-23 | 3410 | 17 | 14 | 10 | TLow_01084-RA\|Tv_02831-RA\|Tv_07802-RA\|Tv_09064-RA\|Tv_11525-RA\|Tv_11854-RA\|Tv_12101-RA\|Tv_15653-RA\|Tv_15720-RA\|Tv_15881-RA |
| biological_process | GO:0009102 | biotin biosynthetic process | 3.24251943712581e-08 | 0 | 3.24251943712581e-08 | 4.58584891822079e-08 | 3410 | 17 | 3 | 3 | Tv_01894-RA\|Tv_01895-RA\|Tv_15468-RA |
| biological_process | GO:0046081 | dUTP catabolic process | 8.68032002463571e-24 | 0 | 8.68032002463571e-24 | 7.16126402032446e-23 | 3410 | 17 | 14 | 10 | TLow_01084-RA\|Tv_02831-RA\|Tv_07802-RA\|Tv_09064-RA\|Tv_11525-RA\|Tv_11854-RA\|Tv_12101-RA\|Tv_15653-RA\|Tv_15720-RA\|Tv_15881-RA |
| HGT Viridiplantae | | | | | | | | | | | |
| molecular_function | GO:0030598 | rRNA N-glycosylase activity | 4.67171528940521e-11 | 0 | 4.67171528940521e-11 | 7.47474446304833e-10 | 5381 | 3 | 3 | 3 | TLow_01580-RA\|Tv_15540-RA\|Tv_15541-RA |
| biological_process | GO:0017148 | negative regulation of translation | 8.89476306015845e-09 | 0 | 8.89476306015845e-09 | 4.06617739892958e-08 | 445 | 3 | 8 | 3 | TLow_01580-RA\|Tv_15540-RA\|Tv_15541-RA |
| HGT Bacteria + Fungi + Viridiplantae | | | | | | | | | | | |
| molecular_function | GO:0000287 | magnesium ion binding | 1.34007184990538e-20 | 0 | 1.34007184990538e-20 | 4.79832178514505e-20 | 6896 | 22 | 42 | 11 | TLow_01084-RA\|Tv_01894-RA\|Tv_02831-RA\|Tv_07802-RA\|Tv_09064-RA\|Tv_11525-RA\|Tv_11854-RA\|Tv_12101-RA\|Tv_15653-RA\|Tv_15720-RA\|Tv_15881-RA |
| molecular_function | GO:0004170 | dUTP diphosphatase activity | 2.16076981295708e-24 | 0 | 2.16076981295708e-24 | 2.39845449238236e-22 | 6896 | 22 | 14 | 10 | TLow_01084-RA\|Tv_02831-RA\|Tv_07802-RA\|Tv_09064-RA\|Tv_11525-RA\|Tv_11854-RA\|Tv_12101-RA\|Tv_15653-RA\|Tv_15720-RA\|Tv_15881-RA |
| molecular_function | GO:0030598 | rRNA N-glycosylase activity | 1.79861038642101e-08 | 0 | 1.79861038642101e-08 | 2.77285767906572e-08 | 6896 | 22 | 3 | 3 | TLow_01580-RA\|Tv_15540-RA\|Tv_15541-RA |
| biological_process | GO:0006226 | dUMP biosynthetic process | 1.27924660789978e-22 | 0 | 1.27924660789978e-22 | 1.1833031123073e-21 | 3654 | 22 | 14 | 10 | TLow_01084-RA\|Tv_02831-RA\|Tv_07802-RA\|Tv_09064-RA\|Tv_11525-RA\|Tv_11854-RA\|Tv_12101-RA\|Tv_15653-RA\|Tv_15720-RA\|Tv_15881-RA |
| biological_process | GO:0009102 | biotin biosynthetic process | 6.11946070241879e-08 | 0 | 6.11946070241879e-08 | 8.93763339432218e-08 | 3654 | 22 | 3 | 3 | Tv_01894-RA\|Tv_01895-RA\|Tv_15468-RA |
| biological_process | GO:0046081 | dUTP catabolic process | 1.27924660789978e-22 | 0 | 1.27924660789978e-22 | 1.1833031123073e-21 | 3654 | 22 | 14 | 10 | TLow_01084-RA\|Tv_02831-RA\|Tv_07802-RA\|Tv_09064-RA\|Tv_11525-RA\|Tv_11854-RA\|Tv_12101-RA\|Tv_15653-RA\|Tv_15720-RA\|Tv_15881-RA |
| biological_process | GO:0017148 | negative regulation of translation | 3.38127547060968e-06 | 0.001 | 3.38127547060968e-06 | 4.57709240533749e-06 | 3654 | 22 | 8 | 3 | TLow_01580-RA\|Tv_15540-RA\|Tv_15541-RA |

**Supplementary Table 9:** Validated HGT candidates from potential bacterial, fungal or viridiplantae donors for *F. occidentalis*

*Bacterial donor:*

| HGT event | Sequence name | Origin of donor sequences | Alternative topology Monophyly of Metazoa | Similarities with the donor sequences | | Length and coverage of the alignment with the donor sequences | | Local score for genomic environment | Homologs found in *T. palmi* | Annotation (BLAST NR) | CAZy | Described in literature |
|---|---|---|---|---|---|---|---|---|---|---|---|---|
| | | | | max id | average id | average aln length | average coverage | | | | | |
| FocB01 | XP_026273122.1 | Bacteria | / | 53.1 | 47.9 | 494 | 76.6 | 0.92 | XP_034232383.1, | Levanase-like | GH32 | Rotenberg et al. 2020; |
| | XP_026287851.1 | | | 47.6 | 44.2 | 493 | 90.1 | / | XP_034233420.1 | | | Li et al. 2022 |
| FocB02 | XP_026276666.1 | Bacteria | 0 | 55 | 52.8 | 308 | 95.6 | 0.9 | XP_034242052.1, | Mannan endo-1,4-beta- | GH5_8 | Rotenberg et al. 2020; |
| | XP_026285289.1 | | | 55.6 | 52.8 | 311 | 94.2 | / | XP_034242578.1 | mannosidase-like | | Li et al. 2022 |
| | XP_026285291.1 | | | 56 | 53.7 | 306 | 95 | / | | | | |
| FocB03 | XP_026272963.1 | Bacteria | 0 | 54.4 | 50 | 143 | 43.8 | 0.84 | XP_034249514.1 | Riboflavin biosynthesis protein | | No |
| FocB04 | XP_026284437.1 | Bacteria | / | 47.5 | 41.7 | 117 | 83.1 | 0.62 | XP_034246168.1 | Unknown protein | | No |
| FocB05 | XP_026277179.1 | Bacteria | 1 | 52.1 | 46.6 | 220 | 92 | 0.8 | | O-methyltransferase | | Rotenberg et al. 2020; |
| | XP_026279074.1 | | | 52.4 | 46.3 | 214 | 84 | 0.42 | | | | Li et al. 2022 |
| | XP_026279079.1 | | | 50.5 | 45.2 | 212 | 93.1 | 0.5 | | | | |

*Fungal donor:*

| HGT event | Sequence name | Origin of donor sequences | Alternative topology Monophyly of Metazoa | Similarities with the donor sequences | | Length and coverage of the alignment with the donor sequences | | Local score for genomic environment | Homologs found in *T. palmi* | Annotation (BLAST NR) | CAZy | Described in literature |
|---|---|---|---|---|---|---|---|---|---|---|---|---|
| | | | | max id | average id | average aln length | average coverage | | | | | |
| FocF01 | XP_026287984.1 | Fungi | / | 57.8 | 54.2 | 212 | 89.8 | 0.32 | XP_034233350.1, | Endoglucanase | GH45 | No |
| | XP_026289264.1 | | | 56.2 | 52 | 213 | 89.8 | 0.68 | XP_034236588.1 | | | |
| | XP_026291890.1 | | | 62.8 | 57.8 | 133 | 51.4 | / | | | | |

*Viridiplantae donor:*

| HGT event | Sequence name | Origin of donor sequences | Alternative topology Monophyly of Metazoa | Similarities with the donor sequences | | Length and coverage of the alignment with the donor sequences | | Local score for genomic environment | Homologs found in *T. palmi* | Annotation (BLAST NR) | CAZy | Described in literature |
|---|---|---|---|---|---|---|---|---|---|---|---|---|
| | | | | max id | average id | average aln length | average coverage | | | | | |
| FocV01 | XP_026290383.1 | Viridiplantae | / | 58.1 | 55.1 | 227 | 93 | 0.66 | XP_034247376.1, | Pathogenesis-related protein 5-like | GH152 | No |
| | | | | | | | | | XP_034247673.1 | | | |

*Complex (Bacterial or Fungal donor):*

| HGT event | Sequence name | Origin of donor sequences | Alternative topology Monophyly of Metazoa | Similarities with the donor sequences | | Length and coverage of the alignment with the donor sequences | | Local score for genomic environment | Homologs found in *T. palmi* | Annotation (BLAST NR) | CAZy | Described in literature |
|---|---|---|---|---|---|---|---|---|---|---|---|---|
| | | | | max id | average id | average aln length | average coverage | | | | | |
| FocC01 | XP_026274115.1 | Bacteria or | 0 | 42.8 | 39.8 | 336 | 88.2 | 0.56 | XP_034235640.1, | Pectin lyase | PL1_4 | No |
| | XP_026279513.1 | Fungi | | 42.4 | 38.7 | 351 | 87.4 | 0.56 | XP_034237191.1, | | | Li et al. 2022 |
| | XP_026279528.1 | | | 41.9 | 40.4 | 325 | 81.0 | 0.62 | XP_034239440.1, | | | Li et al. 2022 |
| | XP_026280402.1 | | | 42.5 | 39.6 | 337 | 88.7 | 0.8 | XP_034241038.1, | | | No |
| | XP_026285066.1 | | | 42.6 | 40.2 | 328 | 86.8 | 0.73 | XP_034248354.1, | | | No |
| | XP_026285067.1 | | | 42.3 | 40.8 | 330 | 87.0 | 0.71 | XP_034248552.1, | | | Li et al. 2022 |
| | XP_026285883.1 | | | 42.9 | 40.9 | 335 | 86.8 | 0.96 | XP_034253769.1, | | | No |
| | XP_026289641.1 | | | 41.2 | 40.0 | 320 | 84.8 | 0.0 | XP_034253771.1, | | | No |
| | XP_026292396.1 | | | 42.6 | 41.3 | 324 | 85.7 | 0.61 | XP_034254938.1 | | | No |
| | XP_026292397.1 | | | 42.9 | 41.5 | 324 | 85.7 | 0.61 | | | | No |

**Supplementary Table 10:** Validated HGT candidates from potential bacterial, fungal or viridiplantae donors for *T. palmi*

*Bacterial donor:*

| HGT event | Sequence name | Origin of donor sequences | Alternative topology Monophyly of Metazoa | Similarities with the donor sequences | | Length and coverage of the alignment with the donor sequences | | Local score for genomic environment | Homologs found in *F. occidentalis* | Annotation (BLAST NR) | CAZy | Described in literature |
|---|---|---|---|---|---|---|---|---|---|---|---|---|
| | | | | max id | average id | average aln length | average coverage | | | | | |
| TpaB01 | XP_034232383.1<br>XP_034233420.1 | Bacteria | / | 46<br>55 | 43.6<br>48.9 | 511<br>497 | 91.5<br>86 | 0.76<br>0.68 | XP_026273122.1,<br>XP_026287851.1 | Levanase-like | GH32 | Li et al. 2022 |
| TpaB02 | XP_034242052.1<br>XP_034242578.1 | Bacteria | / | 59.2<br>59.5 | 55.4<br>55.3 | 302<br>306 | 93.5<br>94.4 | 0.58<br>0.62 | XP_026276666.1,<br>XP_026285289.1,<br>XP_026285291.1 | Mannan endo-1,4-beta-mannosidase-like | GH5_8 | Li et al. 2022 |
| TpaB03 | XP_034249514.1 | Bacteria | 0 | 57.3 | 49.4 | 145 | 74.8 | 0.72 | XP_026272963.1 | Riboflavin biosynthesis protein | | Li et al. 2022 |
| TpaB04 | XP_034246168.1 | Bacteria | / | 47.8 | 40.3 | 128 | 77.9 | 0.44 | XP_026284437.1 | Unknown protein | | No |

*Fungal donor:*

| HGT event | Sequence name | Origin of donor sequences | Alternative topology Monophyly of Metazoa | Similarities with the donor sequences | | Length and coverage of the alignment with the donor sequences | | Local score for genomic environment | Homologs found in *F. occidentalis* | Annotation (BLAST NR) | CAZy | Described in literature |
|---|---|---|---|---|---|---|---|---|---|---|---|---|
| | | | | max id | average id | average aln length | average coverage | | | | | |
| TpaF01 | XP_034236588.1<br>XP_034233350.1 | Fungi | / | 58.5<br>60 | 54.3<br>52 | 218<br>174 | 89.8<br>54.4 | 0.56<br>0.77 | XP_026287984.1,<br>XP_026289264.1,<br>XP_026291890.1 | Endoglucanase | GH45 | No |

*Viridiplantae donor:*

| HGT event | Sequence name | Origin of donor sequences | Alternative topology Monophyly of Metazoa | Similarities with the donor sequences | | Length and coverage of the alignment with the donor sequences | | Local score for genomic environment | Homologs found in *F. occidentalis* | Annotation (BLAST NR) | CAZy | Described in literature |
|---|---|---|---|---|---|---|---|---|---|---|---|---|
| | | | | max id | average id | average aln length | average coverage | | | | | |
| TpaV01 | XP_034247376.1<br>XP_034247673.1 | Viridiplantae | / | 60.4<br>61.3 | 57.4<br>58 | 226<br>221 | 94<br>92 | 0.3<br>0.5 | XP_026290383.1 | Pathogenesis-related protein 5-like | GH152 | No |
| TpaV02 | XP_034245819.1 | Viridiplantae | / | 57 | 48.9 | 292 | 43.2 | 0.89 | | Glucan endo-1,3-beta-glucosidase 13-like | CBM43-<br>CBM43-<br>CBM43-<br>CBM43 | Li et al. 2022 |

Complex (Bacterial or Fungal donor):

| HGT event | Sequence name | Origin of donor sequences | Alternative topology Monophyly of Metazoa | Similarities with the donor sequences | | Length and coverage of the alignment with the donor sequences | | Local score for genomic environment | Homologs found in *F. occidentalis* | Annotation (BLAST NR) | CAZy | Described in literature |
|---|---|---|---|---|---|---|---|---|---|---|---|---|
| | | | | max id | average id | average aln length | average coverage | | | | | |
| TpaC01 | XP_034235640.1<br>XP_034237191.1<br>XP_034239440.1<br>XP_034241038.1<br>XP_034248354.1<br>XP_034248552.1<br>XP_034253769.1<br>XP_034253771.1<br>XP_034254938.1 | Bacteria or Fungi | 0 | 43.4<br>43.7<br>53.7<br>39.8<br>40.9<br>40.3<br>40.6<br>41.5<br>40.8 | 40.27<br>40.3<br>47.3<br>38.1<br>38.9<br>37.6<br>38.8<br>39.3<br>39.3 | 325<br>326<br>136<br>334<br>339<br>339<br>343<br>334<br>336.6 | 81.7<br>82.0<br>83.9<br>88.2<br>89.3<br>89.0<br>90.6<br>88.6<br>89.9 | 0.58<br>0.8<br>0.4<br>0.36<br>0.38<br>0.48<br>0.44<br>0.36<br>0.68 | XP_026274115.1,<br>XP_026279513.1,<br>XP_026279528.1,<br>XP_026280402.1,<br>XP_026285066.1,<br>XP_026285067.1,<br>XP_026285883.1,<br>XP_026289641.1,<br>XP_026292396.1,<br>XP_026292397.1 | Pectin lyase | PL1_4 | Li et al. 2022 |

**Supplementary Table 11:** Overrepresented GO terms among validated HGT candidates from potential bacterial, fungal or viridiplantae donors for *F. occidentalis* and *T. palmi*

*F. occidentalis:*

| root_node_name | node_id | node_name | raw_p_overrep | FWER_overrep | refined_p_overrep | FDR_overrep | nb_genes_refset_root_node | nb_genes_candidate_in_root_node | nb_gene_refset_node | nb_genes_candidate_in_node | name_genes_candidate_in_node |
|---|---|---|---|---|---|---|---|---|---|---|---|
| HGT Bacteria | | | | | | | | | | | |
| molecular_function | GO:0004553 | hydrolase activity, hydrolyzing O-glycosyl compounds | 7.45253094404549e-08 | 0 | 7.45253094404549e-08 | 2.9810123776182e-07 | 5180 | 8 | 131 | 5 | XP_026273122.1\|XP_026276666.1\|XP_026285289.1\|XP_026285291.1\|XP_026287851.1 |
| molecular_function | GO:0008171 | O-methyltransferase activity | 2.64852256493027e-08 | 0 | 2.64852256493027e-08 | 2.11881805194422e-07 | 5180 | 8 | 7 | 3 | XP_026277179.1\|XP_026279074.1\|XP_026279079.1 |
| HGT Fungi | | | | | | | | | | | |
| molecular_function | GO:0008810 | cellulase activity | 1.353949901403e-11 | 0 | 1.353949901403e-11 | 8.12369940841798e-11 | 5113 | 3 | 3 | 3 | XP_026287984.1\|XP_026289264.1\|XP_026291890.1 |
| biological_process | GO:0005975 | carbohydrate metabolic process | 7.51497630059906e-05 | 0.001 | 7.51497630059906e-05 | 0.000112724644508986 | 4038 | 3 | 212 | 3 | XP_026287984.1\|XP_026289264.1\|XP_026291890.1 |
| HGT Bacteria + Fungi + Viridiplantae | | | | | | | | | | | |
| molecular_function | GO:0004553 | hydrolase activity, hydrolyzing O-glycosyl compounds | 9.67932473173405e-13 | 0 | 5.25368376343683e-07 | 1.35510546244277e-11 | 6668 | 11 | 131 | 8 | XP_026273122.1\|XP_026276666.1\|XP_026285289.1\|XP_026285291.1\|XP_026287851.1\|XP_026287984.1\|XP_026289264.1\|XP_026291890.1 |
| molecular_function | GO:0008810 | cellulase activity | 2.23401733731494e-09 | 0 | 2.23401733731494e-09 | 1.04254142408031e-08 | 6668 | 11 | 3 | 3 | XP_026287984.1\|XP_026289264.1\|XP_026291890.1 |
| molecular_function | GO:0008171 | O-methyltransferase activity | 7.7944673051812e-08 | 0.001 | 7.7944673051812e-08 | 2.72806355681342e-07 | 6668 | 11 | 7 | 3 | XP_026277179.1\|XP_026279074.1\|XP_026279079.1 |
| biological_process | GO:0005975 | carbohydrate metabolic process | 6.59293839201467e-06 | 0 | 6.59293839201467e-06 | 1.84602274976411e-05 | 4569 | 5 | 212 | 5 | XP_026273122.1\|XP_026287851.1\|XP_026287984.1\|XP_026289264.1\|XP_026291890.1 |

*T. palmi:*

| root_node_name | node_id | node_name | raw_p_overrep | FWER_overrep | refined_p_overrep | FDR_overrep | nb_genes_refset_root_node | nb_genes_candidate_in_root_node | nb_gene_refset_node | nb_genes_candidate_in_node | name_genes_candidate_in_node |
|---|---|---|---|---|---|---|---|---|---|---|---|
| HGT Bacteria | | | | | | | | | | | |
| molecular_function | GO:0004553 | hydrolase activity, hydrolyzing O-glycosyl compounds | 6.24920835945864e-08 | 0 | 6.24920835945864e-08 | 2.10908540217664e-07 | 4850 | 4 | 116 | 4 | XP_034232383.1\|XP_034233420.1\|XP_034242052.1\|XP_034242578.1 |
| HGT Fungi | | | | | | | | | | | |
| molecular_function | GO:0008810 | cellulase activity | 3.81287870702843e-08 | 0 | 3.81287870702843e-08 | 1.14386361210853e-07 | 116 | 2 | 2 | 2 | XP_034233350.1\|XP_034236588.1 |
| HGT Bacteria + Fungi + Viridiplantae | | | | | | | | | | | |
| molecular_function | GO:0004553 | hydrolase activity, hydrolyzing O-glycosyl compounds | 1.48275313553821e-11 | 0 | 8.52471690838028e-07 | 1.48275313553821e-10 | 5461 | 6 | 116 | 6 | XP_034232383.1\|XP_034233350.1\|XP_034233420.1\|XP_034236588.1\|XP_034242052.1\|XP_034242578.1 |
| molecular_function | GO:0008810 | cellulase activity | 5.71931806054265e-07 | 0 | 5.71931806054265e-07 | 1.90643935351422e-06 | 5461 | 6 | 2 | 2 | XP_034233350.1\|XP_034236588.1 |
| biological_process | GO:0005975 | carbohydrate metabolic process | 3.67257482566665e-05 | 0.006 | 3.67257482566665e-05 | 7.34514965133331e-05 | 4347 | 4 | 194 | 4 | XP_034232383.1\|XP_034233350.1\|XP_034233420.1\|XP_034236588.1 |
